# The renal inflammatory network of nephronophthisis

**DOI:** 10.1101/2021.01.07.425719

**Authors:** Marceau Quatredeniers, Frank Bienaimé, Giulia Ferri, Pierre Isnard, Esther Porée, Katy Billot, Eléonore Birgy, Salomé Ceccarelli, Flora Silbermann, Simone Braeg, Thao Nguyen-Khoa, Rémi Salomon, Marie-Claire Gubler, E. Wolfgang Kuehn, Sophie Saunier, Amandine Viau

**Author notes:** **Corresponding Author** Amandine Viau Inserm U1163 - Institut Imagine Laboratory of Inherited Kidney Diseases Team “nephronophthisis and hypodysplasia” 24 Boulevard du Montparnasse 75015 Paris France.

## Abstract

**BACKGROUND:** The majority of genetic kidney disease leading to kidney failure is caused by mutations in ciliary genes. How cilia malfunction leads to progressive kidney damage is poorly understood, but recent evidence links ciliopathy genes to CCL2 dependent macrophage recruitment in autosomal dominant polycystic kidney disease (ADPKD), the most studied renal ciliopathy. Whether or not renal inflammation is involved in other renal ciliopathies is unclear.

**METHODS:** We combined mice models with kidney biopsies and renal epithelial cells sampled from human urine to characterize the renal inflammatory network of nephronophthisis (NPH), the most frequent renal ciliopathy in children.

**RESULTS:** In human, mutations in cilia genes involved in NPH enhance urine excretion of the chemokine CCL2, causing abnormal macrophage recruitment in kidney tissues from NPH patients. Differing from ADPKD, inactivating *Ccl2* specifically in mouse tubular cells does not rescue the NPH phenotype, suggesting that other inflammatory mediators are involved. Using transcriptional data from 2 NPH models, we identify a set of pro-inflammatory cytokines upregulated in this disease, independently of CCL2. The majority of detectable transcripts from this set are specifically upregulated in kidney cells from NPH patients. In line with the function of these cytokines, NPH kidneys show disproportionate neutrophils and T cells infiltrates compared to healthy subject or hypertensive and diabetic chronic kidney disease patients.

**CONCLUSIONS:** This study reveals that inflammation is a central aspect in human NPH and delineates a specific set of inflammatory mediators that regulates immune cell recruitment in human NPH.

**SIGNIFICANCE STATEMENT:** Mutations in genes encoding primary cilia proteins are the leading cause of genetic kidney failure. In autosomal dominant polycystic kidney disease (ADPKD), deregulated cilia signaling leads to kidney infiltration by macrophages through the chemokine CCL2. Little is known about renal inflammation in nephronophthisis (NPH), the most frequent pediatric renal ciliopathy. Using NPH mice models, tissues and cells from NPH patients, we unveil renal inflammation as preeminent feature of NPH. Remarkably, the renal inflammatory evoked by ciliary gene mutations in NPH does not overlap with ADPKD: it is CCL2 independent, involves a prominent recruitment of neutrophils and T cells and a specific cytokine signature. This unforeseen findings strengthen the link between primary cilia and renal inflammation.

## INTRODUCTION

Renal ciliopathies (RC) are inherited disorders affecting the kidney caused by mutations in genes encoding proteins that localize in primary cilia^1, 2^. The primary cilium is a solitary antenna-like structure protruding from the apical surface of renal tubular cells, acting as a complex macromolecular sensor and signal transducer^3, 4^.

RC share common features, including tubular dilation, interstitial fibrosis and loss of tubular cell differentiation. On the other hand, the spectrum of RC encompasses genetically distinct and phenotypically heterogeneous diseases that manifest from early-childhood to late-adulthood leading to kidney failure in the majority of cases. Autosomal dominant polycystic kidney disease (ADPKD), the most common ciliopathy manifesting in adulthood, is characterized by the development of multiple renal cysts resulting in progressive enlargement of the kidneys accompanied by kidney function decline. Most ADPKD cases are caused by inactivating mutations in *PKD1*, which encodes polycystin 1 (PC1), a receptor-like transmembrane protein, which localizes to the ciliary membrane. Contrary to ADPKD, nephronophthisis (NPH), the second most prevalent RC, is an autosomal recessive disorder that manifests in childhood by enuresis (bedwetting) due to impaired urine concentrating ability and progressive kidney function decline. Although cysts are frequently observed in NPH, cyst burden remains marginal compared to ADPKD, and the kidneys mostly appear small and fibrotic. NPH is a genetically heterogeneous disorder caused by mutations in more than 20 genes identified so far ^5^. Mutations in *NPHP1* are by far the most common, accounting for 40-50% of identified causative mutations in NPH^6^. Most of the proteins encoded by *NPHP* genes assemble in functional modules that control cilia morphology and gate protein entry and exit to and from the cilia^7, 8^. Yet, how disruption of these complexes translate into pathophysiologic events to cause kidney damage is poorly understood.

Primary cilia of renal tubular cells are required for proper kidney development and maintenance^9^. Yet, none of the mutations reported so far in RC abolish ciliogenesis. While renal tubular cells remain ciliated in RC, they may display either increased or decreased cilia length^10, 11^. It is believed that the genetic defects involved in RC adversely affect cilia composition and signaling which results in modifications of tubular cell behavior that, in turn, alter kidney morphology and function.

Lessons gathered from acquired chronic kidney disease models indicate that progressive renal scarring involves multi-directional communication between tubular, immune and mesenchymal cells that prompts immune cell infiltration, fibroblasts activation and nephron loss^12–14^.

Concordant evidence has pinpointed an important role of renal inflammation in the progression of ADPKD. Indeed, macrophages infiltrate the kidney of ADPKD patients and mice and promote cyst growth^15^. Remarkably, cilia signaling seems to be directly involved in this process as cilia ablation prevents macrophage recruitment in an orthologous model of ADPKD^16^. Mechanistically, cilia positively regulate the expression of the macrophage chemoattractant CCL2 in *Pkd1*-deficient tubular cells both *in vivo* and *in vitro*. Consistently, kidney specific disruption of *Ccl2* or inhibition of the CCL2 receptor reduce cyst burden in *Pkd1* mutant mice^16, 17^.

In contrast to ADPKD, the function of immune cells in NPH has received little attention. Recent evidence however suggests that renal inflammation may be involved in the disease. First, the inactivation of TLR2, a critical receptor of the immune system, prevents renal damage in an orthologous model of a rare form of NPH caused by a mutation in *Glis2/Nphp7* (*Glis2*^lacZ/lacZ^ mice)^18, 19^. We obtained additional evidence supporting an involvement of renal inflammation in NPH while studying the function of LKB1 in tubular cells: LKB1 is a ciliary kinase involved in the control of cell size and metabolism through the regulation of AMPK. We previously reported that LKB1 interacts with NPHP proteins including NPHP1. Strikingly, mice bearing a selective inactivation of *Lkb1* in renal tubules (*Lkb1*^ΔTub^ mice) develop an NPH-like phenotype recapitulating the consequences of *NPHP1* loss in humans. *In vitro*, both NPHP1 and LKB1 repress the expression of CCL2, while CCL2 upregulation and macrophage recruitment are observed *in vivo* at an early time point of the development of the disease in *Lkb1*^ΔTub^ mice^16^. Importantly, CCL2 dependent recruitment of immune cells has been repeatedly shown to promote renal fibrosis in different acquired chronic kidney disease models^14, 20^.

Considering the paucity of data regarding renal inflammation in NPH, we decided to explore the interplay between immune and tubular cells in NPH through a translational approach combining data from transgenic animals and human patients.

## MATERIALS AND METHODS

### Human kidney tissue specimens

Renal kidney tissues from 12 patients suffering from juvenile nephronophthisis (NPH) were collected from 1973 to 1998. Kidneys donated for transplantation (n=5) but unsuitable for implantation (due to damage to the arterial patch or parenchymal sclerosis) or kidney from patient suffering renal disease other than nephronophthisis with minimal histological lesions (n=1) and collected at the same time period were used as controls. Renal biopsies from patients suffering diabetic nephropathy (n=5) or hypertensive nephrosclerosis (n=6) were used to control for non-specific, chronic kidney disease (CKD) related inflammation. Interstitial fibrosis and tubular atrophy (IFTA) on renal biopsy was evaluated by a trained pathologist in a blind fashion. NPH and CKD patient groups were matched according to their IFTA. Patient data are listed in Supplementary Table 1.

The human kidney tissue specimens belonging to the Imagine Biocollection are declared to the French Minister of Research under the number DC-2020-3994 and approved by the French Ethics committee for research at Assistance Publique-Hôpitaux de Paris (CERAPHP) under the IRB registration number #00011928.

### Isolation of urine-derived renal epithelial cells (UREC)

Urine samples were collected from *NPHP1* patients (n=9), healthy relatives and unrelated controls (n=10) and CKD patients (n=6) recruited at Necker Hospital (Paris, France) after written informed consent of the donor. Inclusion criteria for affected patients were to suffer from nephronophthisis with known genetic diagnosis. The relatives were the healthy relatives (father/mother/sister) of an included patient. Healthy controls were unscathed of any chronic kidney disease or suffering renal disease other than nephronophthisis with normal renal function. CKD patients were suffering chronic kidney disease other than nephronophthisis or other renal ciliopathy. The study was approved by the French National Committee for the Protection of Persons (CPP) under the ID-RCB number 2016-A00541-50 and is kept in full accordance with the principles of the Declaration of Helsinki and Good Clinical Practice guidelines.

Urine-derived renal tubular epithelial cells (primary UREC) were isolated and cultured as previously described^21^ with some modifications. Briefly, after centrifugation and washing steps, urine-derived cells were initially cultured for 4 days at 37°C in primary medium containing Dulbecco’s Modified Eagle Medium : Nutrient Mixture F-12 (DMEM/F-12) supplemented with 10% fetal bovine serum (16000-036, Gibco), 10% Penicillin-Streptomycin (15140-122, Gibco), 10% Amphotericin B (15290026, Gibco) and 1X REGM^TM^ SingleQuots^TM^ kit (CC-4127, Lonza) to enhance cell survival and adherence. At day 4, primary medium was replaced by a growth medium containing REBM^TM^ (Basal Medium, CC-3191, Lonza) supplemented with 2% fetal bovine serum, 10% Penicillin-Streptomycin, 10% Amphotericin B, 1X REGM^TM^ SingleQuots^TM^ kit and 10 ng/mL rhEGF (R&D system) and was then changed every 2 days. At 80% confluence (7-30 days), 2x10^4^ cells were seeded for 7-24 days until confluence on 12 well plates (353043, Dutscher) for RNA analysis.

### Mice

All animal experiments were conducted according to the guidelines of the National Institutes of Health *Guide for the Care and Use of Laboratory Animals*, as well as the German law for the welfare of animals, and were approved by regional authorities (Regierungsprasidium Freiburg, 35-9185.81/G-16/27). Mice were housed in a specific pathogen-free facility, fed *ad libitum* and housed at constant ambient temperature in a 12-hour day/night cycle. Breeding and genotyping were done according to standard procedures.

*Lkb1*^flox/flox^ mice (mixed genetic background) and *Ccl2*-RFP^flox/flox^ (B6.Cg-Ccl2tm1.1Pame/J, stock number : 016849, C57BL/6 genetic background) were purchased from The Jackson Laboratories (STOCK *Stk11^tm^*^1^*^.1Sjm^*/J, stock number: 014143) and were crossed to *KspCre* mice (B6.Cg-Tg(Cdh16-cre)91lgr/J; C57Bl/6N background)^22^ to generate a tubule-specific *Lkb1* knockout (further referred to as *Lkb1*^ΔTub^) and *Lkb1*; *Ccl2* knockout (further referred to as *Lkb1*^ΔTub^; *Ccl2*^ΔTub^). Littermates lacking *KspCre* transgene were used as controls. Experiments were conducted on both females and males.

### Quantitative PCR

Total RNAs were obtained from human UREC or mouse kidneys using RNeasy Mini Kit (Qiagen) and reverse transcribed using SuperScript II Reverse Transcriptase (Life Technologies) or High Capacity cDNA Reverse Transcription Kit (Applied Biosystems) according to the manufacturer’s protocol. Quantitative PCR were performed with iTaq™ Universal SYBR® Green Supermix (Bio-Rad) on a CFX384 C1000 Touch (Bio-Rad). *Hprt, Ppia, Rpl13, Sdha and Tbp* were used as normalization controls^23^. Each biological replicate was measured in technical duplicates. The primers used for qRT-PCR are listed in Supplementary Table 2.

### CCL2 ELISA

For CCL2 measurement in urines from *NPHP1* patients (n=9) and controls (n=8), urine specimens were collected and centrifuged at 1,500 x g for 10 minutes at 4°C within 4 hours of collection. Clinical and genetic data of the patients are listed in Supplementary Table 1. The supernatants were collected and stored at -80°C. Frozen aliquots of urine supernatants were thawed at room temperature immediately before the ELISA. The samples were used with a 2-fold dilution and were tested in duplicates. CCL2 levels in urine specimens were quantified using Human CCL2/MCP-1 Quantikine® ELISA Kit (R&D systems, DCP00) according to the manufacturer’s instructions. The plate was read using a Multiskan Sky plate reader set to subtract for wavelength and blank corrections. The optical densities were derived from 4-parameter logistic regression of the standard curve. Measurement of creatinine in urine was performed in the same samples using IDMS-standardized enzymatic method on C16000 Architect analyzer (Abbott Diagnostic). The results were normalized to the urinary creatinine level. Indeed, normalization by the urinary creatinine levels avoids the pitfall of concentration or dilution of urine.

### Morphological Analysis

Human kidney biopsies were fixed in alcohol formalin and acetic acid and paraffin embedded, 4µm sections were stained with periodic acid-Schiff (PAS). PAS-stained full size images were recorded using a whole slide scanner Nanozoomer 560 (Hamamatsu) coupled to NDPview software (Hamamatsu).

Mouse kidneys were fixed in 4% paraformaldehyde, embedded in paraffin, and 4µm sections were stained with PAS or Picrosirius Red. Stained full size images were recorded using a whole slide scanner Nanozoomer 2.0 (Hamamatsu) equipped with a 20x/0.75 NA objective coupled to NDPview software (Hamamatsu). Histology score was evaluated by an independent observer in a blinded fashion assessing the overall lesions comprised of tubular atrophy, tubular basement thickening and cell infiltration of the whole kidney section stained with PAS.

### Urine and Plasma Analyses

8-hour urine samples were obtained from mice housed in individual boxes without access to water and food. Body weight and urine excretion were measured. Urine osmolality was measured with a freezing point depression osmometer (Micro-Osmometer from Knauer or OSMOMAT 3000basic from Gonotec). Retro-orbital blood was collected from anaesthetized mice. Plasma blood urea nitrogen (BUN) was measured using urea kit (LT-UR; Labor&Technik, Eberhard Lehmann GmbH) according to the manufacturer’s instructions.

### Immunohistochemistry

#### For human kidney sections

An automated IHC stainer BOND-III (Leica Biosystems) was used. Briefly, 4µm sections of paraffin-embedded human kidney biopsies were submitted to the appropriate antigen retrieval. Then, sections were incubated with CD68 (Dako, M0814, 1:3,000), CD15 (Beckam Coulter, NIM0165, 1:200) or CD3 (Dako, A0452, 1:200) antibodies. Peroxide blocking, post primary, DAB chromogen and hematoxylin counterstaining was performed automatically using Bond polymer refine detection kit (Leica Biosystems, DS9800). The degree of interstitial cell infiltration was determined using immunostaining targeting macrophages (CD68), neutrophils (CD15) and T lymphocytes (CD3). Slides were scanned with a whole slide scanner Nanozoomer 560 (Hamamatsu). Randomly selected microscopic fields (x200) representative of the entire cortical surface were scored. The degree of cell infiltrate was quantified using ImageJ software. For CD3 and CD68, the area of DAB staining was measured after color deconvolution and intensity thresholding of the images and visualized as the ratio of DAB surface to cortical surface on each microscopic field. For CD15, the number of CD15-positive interstitial cells was quantified manually in a blinded fashion and was expressed as a ratio per mm^2^.

#### For mouse kidney sections

Macrophage staining: 4µm sections of paraffin-embedded mouse kidneys were incubated for 20 min at 95°C in citrate buffer (Zytomed, ZUCD28) followed by avidin/biotin blocking (Vector, SP-2001). Sections were incubated with F4/80 antibody (Clone Cl:A3-1, Bio-Rad, MCA497R, 1:100) followed by biotinylated antibody (Vector, BA-4001, 1:200), HRP-labeled streptavidin (Southern Biotech, 7100-05, 1:2,000) and 3-3′-diamino-benzidine-tetrahydrochloride (DAB, Dako, K3468) revelation.

Neutrophil staining: 4µm sections of paraffin-embedded mouse kidneys were submitted to antigen retrieval for 20 min at 95°C in citrate buffer followed by incubation with Ly-6B.2 antibody (Abcam, ab53457, 1:100). Sections were incubated with HRP-labeled secondary antibody (Vector, PI-9400, 1:200) and DAB revelation. T cell staining: 4µm sections of paraffin-embedded mouse kidneys were incubated for 20 min at 95°C in Tris-EDTA pH9 buffer followed by avidin/biotin blocking. Sections were incubated with CD3 antibody (Abcam, ab16669, 1:100) followed by biotinylated antibody (GE Healthcare, RPN1004V, 1:200), HRP-labeled streptavidin (Southern Biotech, 7100-05, 1:2,000) and DAB revelation.

Full size images were recorded using a whole slide scanner Nanozoomer 2.0 coupled to NDPview software. Stained area was measured with ImageJ software from full size kidney images and visualized as the ratio of stained DAB surface to total kidney section area.

### Comparative microarray data analysis

Processing of data was carried out using R v3.6.0, RStudio and R package dplyr v1.0.1. Renal transcriptomic datasets from 5 weeks old *Lkb1*^ΔTub^ and 4 weeks old *Glis2*^lacZ/lacZ^ mouse models^16, 18^ were compared. While transcriptomic dataset from *Lkb1*^ΔTub^ mice was established by our team (GSE86011)^16^, we downloaded from the GEO database (https://www.ncbi.nlm.nih.gov/geo/) the expression matrix of mRNAs expressions in the kidneys of *Glis2*^lacZ/lacZ^ mice under the accession number GSE6113^18^. Probe IDs in the expression matrix were matched with the corresponding gene IDs in the lookup table (GPL2897) to identify the expression of each mRNA. To assess the common regulated genes between the two mouse models, *Lkb1*^ΔTub^ dataset was filtered out according to false discovery rate (FDR, Benjamini-Hochberg procedure) < 0.05 to obtain a list of 1,991 differentially expressed genes (DEGs). Then, *Glis2*^lacZ/lacZ^ dataset (23,957 genes) was matched with this DEGs list, and the genes whose expression varies the same way were considered the common regulated genes (1,262 genes) listed in Supplementary Table 3.

### GSEA, networking and visualization

Subsequently, Gene Set Enrichment Analysis (GSEA) approach was applied to the 1,262 common regulated genes identified using GSEA software v4.0.0 (Broad Institute) set for 1,000 gene set permutations. The enrichment involved the « biological process » classification of the Gene Ontology downloaded from MSigDB v7.1 (C5-BP). Up- and down-regulated pathways are listed in Supplementary Tables 4-5. Pathways with FDR < 0.05 were considered significant. R package ggplot2 was used to present the more significant up- and down-regulated pathways. Cytoscape software v3.0 and Cytoscape applications EnrichmentMap v3.3.0 (edge cutoff threshold set to 0.6) and WordCloud v3.1.3 were used for pathway clustering and pathway network visualization^24^. Pathway clusters have been hand-annotated.

### Identification of common upregulated cytokines

To identify common upregulated cytokines, the 823 common upregulated genes were matched with lists of genes obtained from the UniProt database (https://www.uniprot.org/) using the following keywords: « cytokine », « secreted » and « immune/inflammation ». Genes that were linked to the 3 keywords are listed in Supplementary Table 6.

Heatmaps of the expression of the identified cytokines from 5 weeks old *Lkb1*^ΔTub^ and 10 weeks old *Pkd2*^ΔTub^ were compared. The *Pkd2*^ΔTub^ log-normalized expression matrix was downloaded from GEO under the accession number GSE149739^25^. Differential expression testing between groups (n=3 per group) was performed using limma v3.42.2. Heatmaps were generated using the R package pheatmap v1.0.12. Expression levels of the 17 identified cytokines in mouse kidney *Lkb1*^ΔTub^ and *Pkd2* ^ΔTub^ datasets are listed in Supplementary Table 7.

### Statistical analysis

Data were expressed as means. Differences between groups were evaluated using Mann-Whitney test when only two groups were compared or when testing more comparisons, one-way ANOVA followed when significant (*P* < 0.05) by the Tukey-Kramer test. The statistical analysis was performed using GraphPad Prism V8 software. All image analysis (immunohistochemistry) and mouse phenotypic analysis were performed in a blinded fashion.

## RESULTS

### Nephronophthisis is associated with macrophage recruitment and enhanced CCL2 expression

To elucidate the contribution of immune cell recruitment in the phenotype of NPH, we first aimed to assess whether, similar to ADPKD, macrophage infiltration occurs in human NPH. Histology inspection of kidney tissues from NPH patients revealed significant infiltration by mononuclear cells (**Figure 1A**). Immunolabelling identified macrophages (CD68-positive cells) as an important contingent of cells infiltrating the kidneys from NPH patients as compared to control individuals (**Figure 1B-C**). As CCL2 has been shown to be the major chemokine responsible for macrophage recruitment in ADPKD, we analyzed primary tubular epithelial cells derived from urine (UREC) of *NPHP1* patients and controls, including both age-matched controls and healthy relatives. UREC derived from patients bearing *NPHP1* mutations showed increased expression of *LCN2* transcript, a marker of tubular injury that positively correlates with the progression of chronic kidney disease (**Supplementary Figure 1A-B**). Remarkably, tubular cells derived from *NPHP1* patients also showed enhanced expression of *CCL2* transcript (**Figure 1D** and **Supplementary Figure 1C**). Consistently, we observed that *NPHP1* patients display a higher urinary excretion rate of CCL2 than controls (**Figure 1E**). Collectively, these data revealed that, similarly to ADPKD, NPH patients show increased renal tubular expression of CCL2 associated with macrophage recruitment.

**Figure 1.**
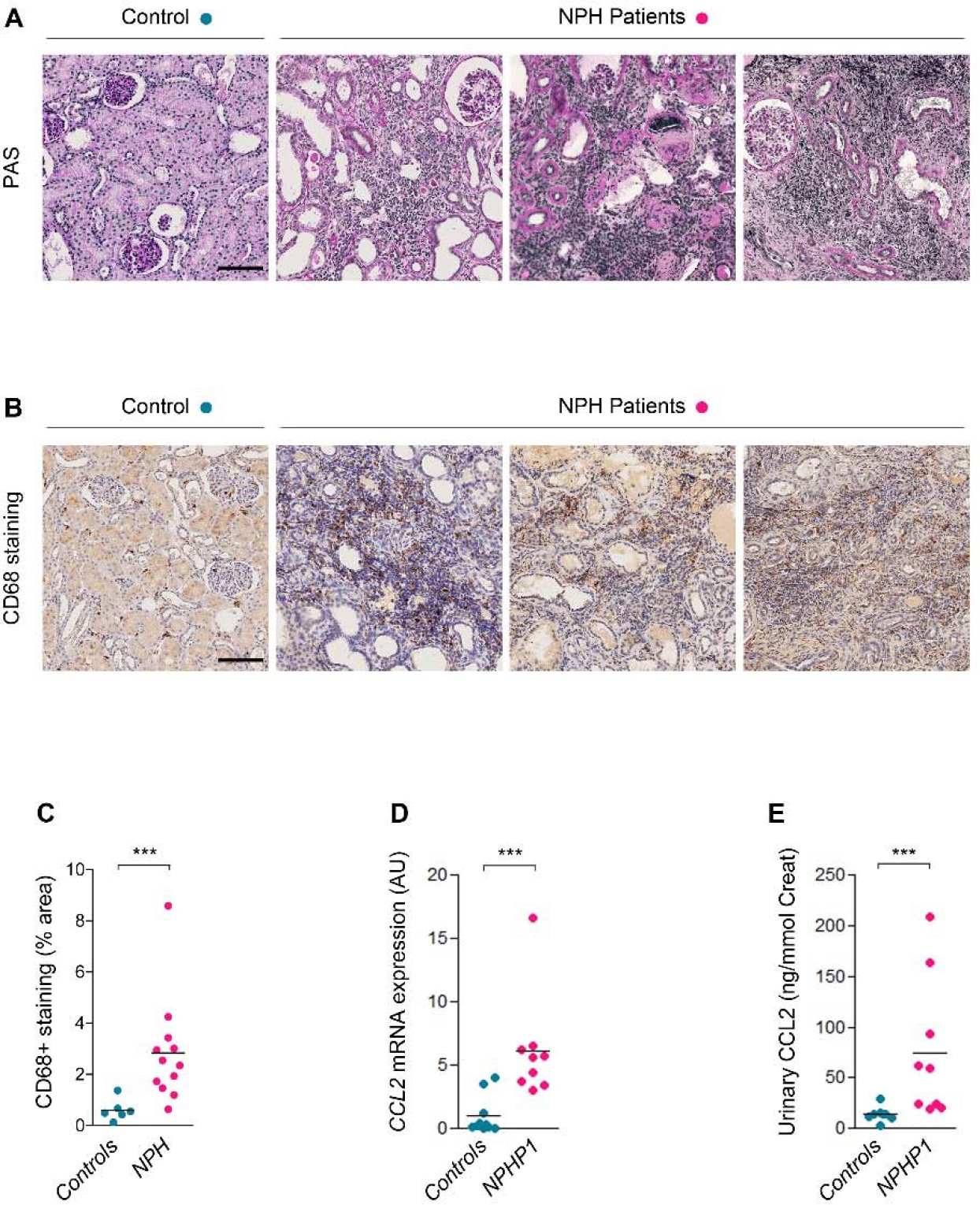
Human nephronophthisis is associated with renal inflammation, macrophage recruitment and CCL2 upregulation. **(A)** Representative images of Periodic Acid–Schiff (PAS) stained kidney biopsies from control and 3 NPH patients. Scale bar: 100µm. **(B)** Representative images of CD68 (macrophages) immunostaining in kidney biopsies from control and 3 NPH patients. Scale bar: 100µm. **(C)** Quantification of CD68-positive staining area in kidney sections from control and NPH patients**. (D)** *CCL2* mRNA expression in primary UREC derived from urine from 10 controls (Controls) and 9 *NPHP1* patients (NPHP1). **(E)** Urinary CCL2 secretion in Controls and *NPHP1* patients. **(C-E)** Each dot represents one individual. Bars indicate mean. Mann-Whitney t test, *** P<0.001. AU: arbitrary unit.

### Tubule-specific *Ccl2* inactivation does not prevent nephronophthisis-like phenotype

Tubular deletion of *Ccl2* in an ADPKD orthologous mouse model dampens renal macrophage recruitment and cyst formation^16, 17^. The deletion of *Lkb1* in the mouse kidney results in an NPH phenotype associated with CCL2 upregulation and macrophage recruitment^16, 26^. To determine if the deletion of *Ccl2* together with *Lkb1* ameliorates the NPH-like renal disease, we crossed tubule-specific *Lkb1*^Δtub^ mice with mice bearing *Ccl2* floxed alleles and compared mice with inactivation of *Lkb1* alone (*Lkb1*^Δtub^) or *Lkb1* and *Ccl2* (*Lkb1*^Δtub^; *Ccl2*^Δtub^) with wild-type controls. At 10 weeks of age, quantitative RT-PCR revealed that *Ccl2* inactivation in tubular cells drastically decreased the overall *Ccl2* expression in *Lkb1* inactivated kidneys (**Figure 2A**). Macroscopic inspection revealed irregular kidneys in both *Lkb1*^Δtub^ and *Lkb1*^Δtub^; *Ccl2*^Δtub^ mice associated with reduced kidney size, which was even more pronounced in *Lkb1*^Δtub^; *Ccl2*^Δtub^ mice (**Figure 2B-C**). *Lkb1*^Δtub^ and *Lkb1*^Δtub^; *Ccl2*^Δtub^ mice both displayed a urine concentration defect and a similar loss of kidney function (**Figure 2D-F**). Histology revealed similar tubular basement membranes thickening, interstitial inflammation, fibrosis, and tubular dilation in kidneys from *Lkb1*^Δtub^ and *Lkb1*^Δtub^; *Ccl2*^Δtub^ mice. Consistently, *Ccl2* inactivation did not affect the upregulation of the tubular injury marker, *Lcn2*, as well as the extracellular matrix deposition markers, *Col1a1* and *Tgfb1* in *Lkb1* deficient kidneys (**Figure 2G-L**).

**Figure 2.**
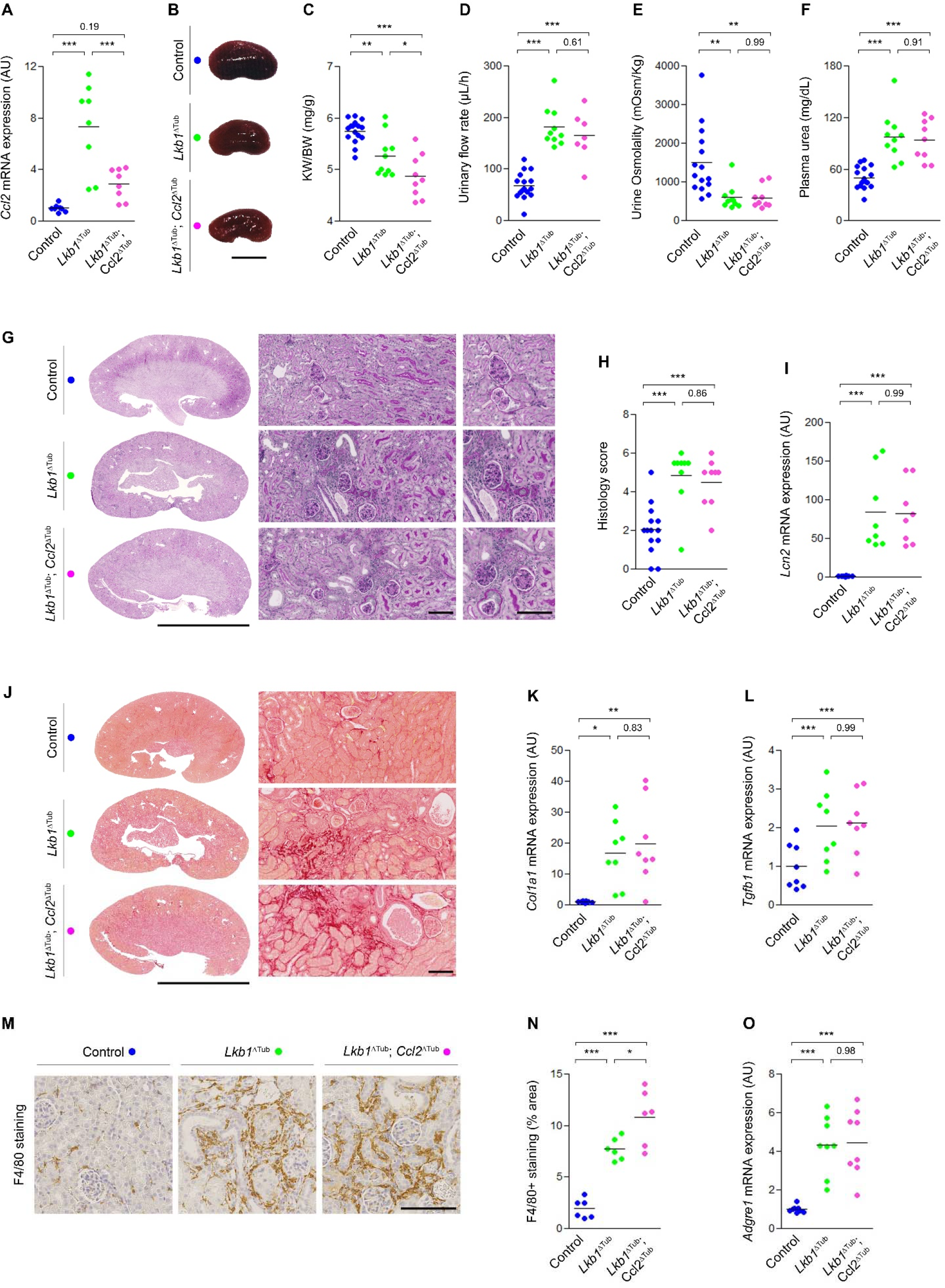
Tubule specific *Ccl2* inactivation does not prevent renal lesion development nor macrophage infiltration of the kidney in response to *Lkb1* disruption. **(A)** *Ccl2* mRNA expression in kidneys from control, *Lkb1*^ΔTub^ and *Lkb1*^ΔTub^; *Ccl2*^ΔTub^ mice at 10 weeks. **(B)** Representative kidneys from 10 weeks old control, *Lkb1*^ΔTub^ and *Lkb1*^ΔTub^; *Ccl2*^ΔTub^ mice. Scale bar: 5 mm. **(C)** Kidney weight to body weight ratio (KW/BW) at 10 weeks. **(D-E)** Urinary flow rate (D) and urine osmolality (E) at 10 weeks. **(F)** Plasma blood urea nitrogen (BUN) at 10 weeks. **(G)** Representative PAS stained kidney sections from 10 weeks old control, *Lkb1*^ΔTub^ and *Lkb1*^ΔTub^; *Ccl2*^ΔTub^ mice. Scale bars: 5mm (left panel), 100µm (middle, right panel). **(H)** Histology score based on PAS-stained sections from control, *Lkb1*^ΔTub^ and *Lkb1*^ΔTub^; *Ccl2*^ΔTub^ mice at 10 weeks. **(I)** *Lcn2* mRNA expression in kidneys from control, *Lkb1*^ΔTub^ and *Lkb1*^ΔTub^; *Ccl2*^ΔTub^ mice at 10 weeks. **(J)** Representative sirius red stained kidney sections from 10 weeks old control, *Lkb1*^ΔTub^ and *Lkb1*^ΔTub^; *Ccl2*^ΔTub^ mice. Scale bars: 5mm (left panel), 100µm (right panel). **(K, L)** *Col1a1* and *Tgfb1* mRNA expression in kidneys from control, *Lkb1*^ΔTub^ and *Lkb1*^ΔTub^; *Ccl2*^ΔTub^ mice at 10 weeks. **(M)** Representative images of F4/80 (macrophages) immunostaining of kidney sections from control, *Lkb1*^ΔTub^ and *Lkb1*^ΔTub^; *Ccl2*^ΔTub^ mice at 10 weeks. Scale bar: 100µm. **(N)** Quantification of F4/80 positive staining area in whole sections from 10 weeks old control, *Lkb1*^ΔTub^ and *Lkb1*^ΔTub^; *Ccl2*^ΔTub^ mice. **(O)** *Adgre1* mRNA expression in kidneys from control, *Lkb1*^ΔTub^ and *Lkb1*^ΔTub^; *Ccl2*^ΔTub^ mice at 10 weeks. **(A, C-F, H-I, K-L, N-O)** Each dot represents one individual mouse. Bars indicate mean. One-way ANOVA followed by Tukey-Kramer test, * P<0.05, ** P<0.01, *** P<0.001. AU: arbitrary unit.

The protective effect of *Ccl2* inactivation in ADPKD and in acquired kidney disease is mediated by reduced macrophage infiltration. Unexpectedly, contrary to what was observed in *Pkd1* deficient mice^16, 17^, *Ccl2* inactivation did not prevent macrophage infiltration of *Lkb1* deficient kidneys as judged by quantification of both F4/80 immunolabelling and F4/80 transcript quantification (*Adgre1*) (**Figure 2M-O**). Collectively, these results demonstrate that, although tubular cells play a major role in the upregulation of CCL2 in *Lkb1* deficient kidneys, this cytokine does not play a significant role in renal macrophage recruitment nor in the induction of NPH-like nephropathy.

### Early NPH renal disease in mice is associated with a prominent immune signature

So far, our findings revealed macrophage infiltration in NPH but, as opposed to ADPKD, this macrophage recruitment appears independent of CCL2. To approach the mechanisms underlying renal inflammation in NPH in an unbiased manner, we compared microarray-based transcriptomes from *Lkb1*^ΔTub^ and *Glis2*^lacZ/lacZ^ kidneys^16, 18^ to control kidneys. Notably, both mouse models recapitulated the features of human NPH: polyuria followed by progressive interstitial fibrosis, tubular basement membrane thickening, tubular dilations and immune cell infiltration^16, 18, 27^. We focused on transcriptome datasets obtained at an early time point of disease development (4 weeks and 5 weeks). Gene overlap analysis identified 1,262 genes that were commonly deregulated in the two models (**Figure 3A-B** and **Supplementary Table 3**). Gene set enrichment analysis (GSEA) of these genes identified a total of 72 enriched pathways (FDR<0.05). Among those pathways, 8 were downregulated, consisting mostly of metabolic processes, while 64 biological processes were upregulated (**Supplementary Tables 4-5**). Among the latter, the GSEA revealed a high enrichment in biological processes linked to immune response and inflammation (**Figure 3C**). Network analysis showed a marked association of the common regulated genes with immune pathways (**Figure 3D** **and Supplementary Figure 2**). This unbiased analysis identified renal inflammation as an early and prominent phenomenon in the course of NPH models in the mouse.

**Figure 3.**
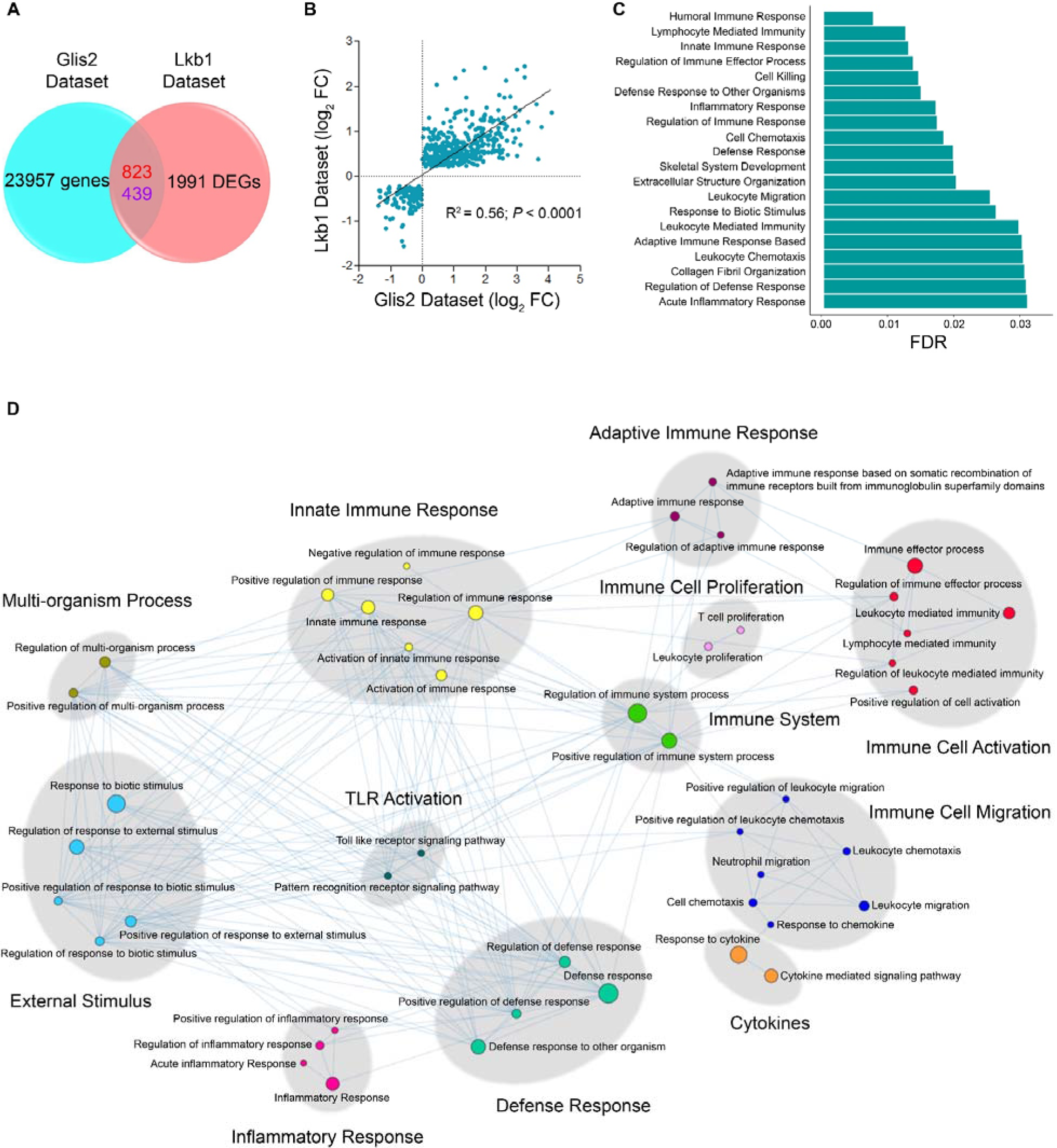
Comparative renal transcriptome analysis in *Glis2* mutant mice and kidney-specific *Lkb1* deficient mice (*Lkb1*^ΔTub^). **(A)** Venn diagram showing intersection between *Glis2*^lacZ/lacZ^ (left; 23,957 genes) and *Lkb1*^ΔTub^ (right; 1,991 differentially expressed genes (DEGs) according to FDR < 0.05) datasets. Red numbers: upregulated; blue numbers: downregulated. **(B)** Jointly up- and down-regulated genes in *Lkb1*^ΔTub^ mice (Lkb1 dataset) and *Glis2*^lacZ/lacZ^ mice (Glis2 dataset). Genes with FDR < 0.01 are represented. Pearson correlation R^2^ = 0.56, P<0.0001. **(C)** Gene set enrichment analysis (GSEA) on the 1,262 common regulated genes revealed 72 pathways overrepresented (FDR < 0.05); the 20 most significantly upregulated are presented. See also Supplementary Tables 4-5. **(D)** Summarized subnetwork representation of significantly enriched Biological Processes GO terms (MSigDB v7.1, C5-BP) linked to immune/inflammatory pathways derived from the GSEA of the 1,262 common regulated genes between *Glis2*^lacZ/lacZ^ and *Lkb1*^ΔTub^ kidney transcriptomes. Nodes are connected according to a similarity score > 0.6, and pathway clusters are hand-annotated according to the most representative GO-BP terms. **(C-D)** Schemes were generated using R v3.6.0 and R studio.

### *Lkb1* tubular inactivation activates multiple chemotactic pathways and drives the recruitment of distinct immune cell populations independently of CCL2

To identify specific cytokines that might illuminate CCL2 independent immune cell recruitment in NPH, we matched the common upregulated genes between *Lkb1*^ΔTub^ and *Glis2*^lacZ/lacZ^ kidneys with UniProt database. This analysis retrieved 17 pro-inflammatory cytokines, other than CCL2 (**Supplementary Figure 3** and **Supplementary Table 6**) that have been implicated in macrophage, neutrophil, T cell and dendritic cell chemotaxis. Reanalyzing RNA-seq data from an orthologous rodent model of ADPKD^25^, we failed to detect a consistent induction of these inflammatory genes in polycystic kidneys. Thus, this inflammatory signature does not appear to be a general feature of renal ciliopathies but is rather specific to NPH (**Supplementary Figure 3** and **Supplementary Table 7**). Quantifying these transcripts in kidneys from *Lkb1*^Δtub^, *Lkb1*^Δtub^; *Ccl2*^Δtub^ and control mice, we observed that *Ccl2* disruption had no impact on the up-regulation of these immune mediators in LKB1 deficient kidneys (**Figure 4**). In line with the different immune cell populations attracted by these cytokines, immunolabelling demonstrated enhanced neutrophils (Ly-6B.2 positive cells) and T cells (CD3 positive cells) recruitment in both kidneys from *Lkb1*^Δtub^ and *Lkb1*^Δtub^; *Ccl2*^Δtub^ mice as compared to control mice (**Figure 5**). Collectively, these results uncover a CCL2 independent inflammatory network in NPH-like renal disease.

**Figure 4.**
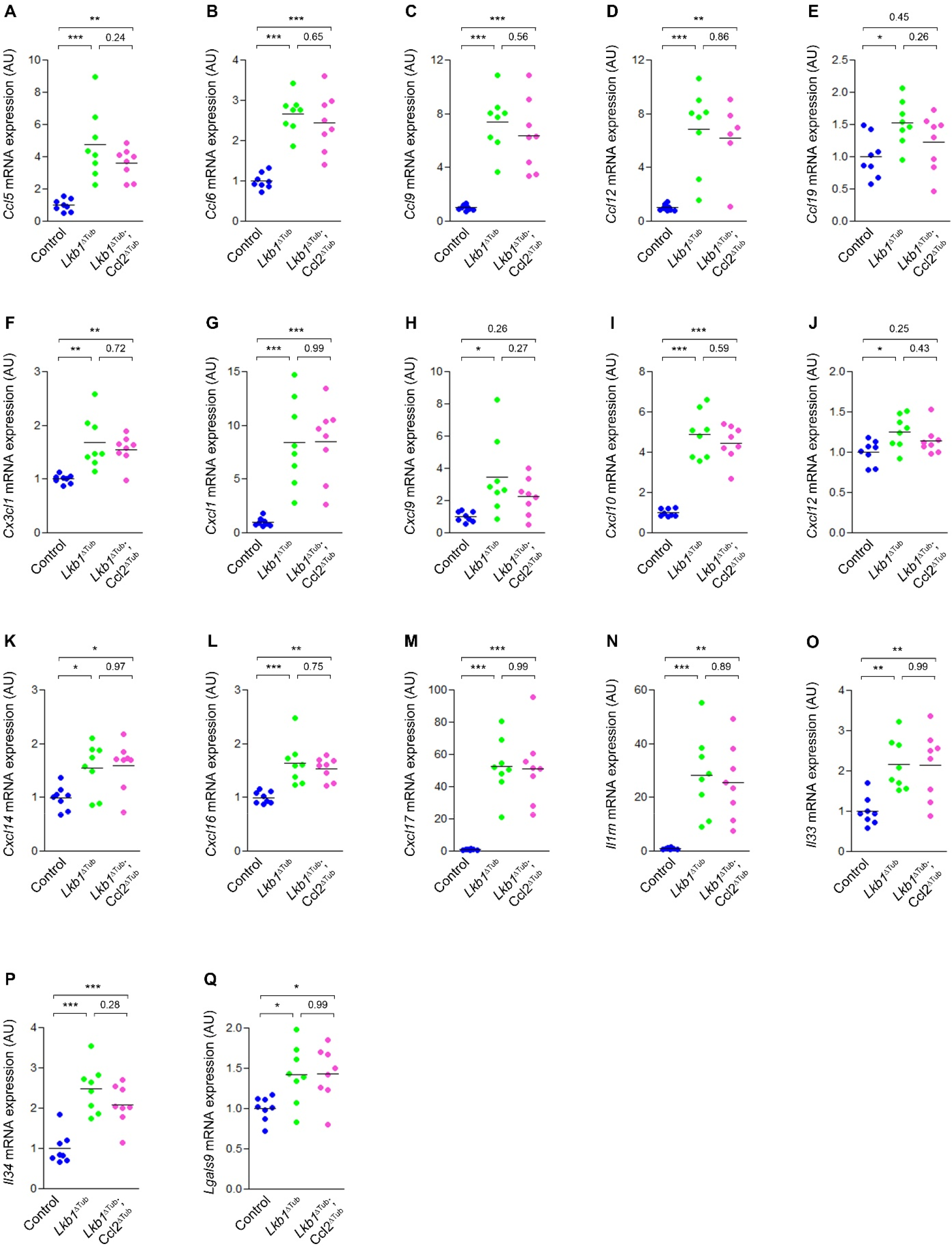
Nephronophthisis inflammatory mediator signature is independent of CCL2. **(A-Q)** Quantification of *Ccl5 (*A), *Ccl6* (B), *Ccl9* (C), *Ccl12* (D), *Ccl19* (E), *Cx3cl1* (F), *Cxcl1* (G), *Cxcl9* (H), *Cxcl10* (I), *Cxcl12* (J), *Cxcl14* (K), *Cxcl16* (L), *Cxcl17* (M), *Il1rn* (N), *Il33* (O), *Il34* (P) and *Lgals9* (Q) mRNA abundance in kidneys from 10 weeks old control, *Lkb1*^ΔTub^ and *Lkb1*^ΔTub^; *Ccl2*^ΔTub^ mice. Each dot represents one individual mouse. Bars indicate mean. One-way ANOVA followed by Tukey-Kramer test, * P<0.05, ** P<0.01, *** P<0.001. AU: arbitrary unit.

**Figure 5.**
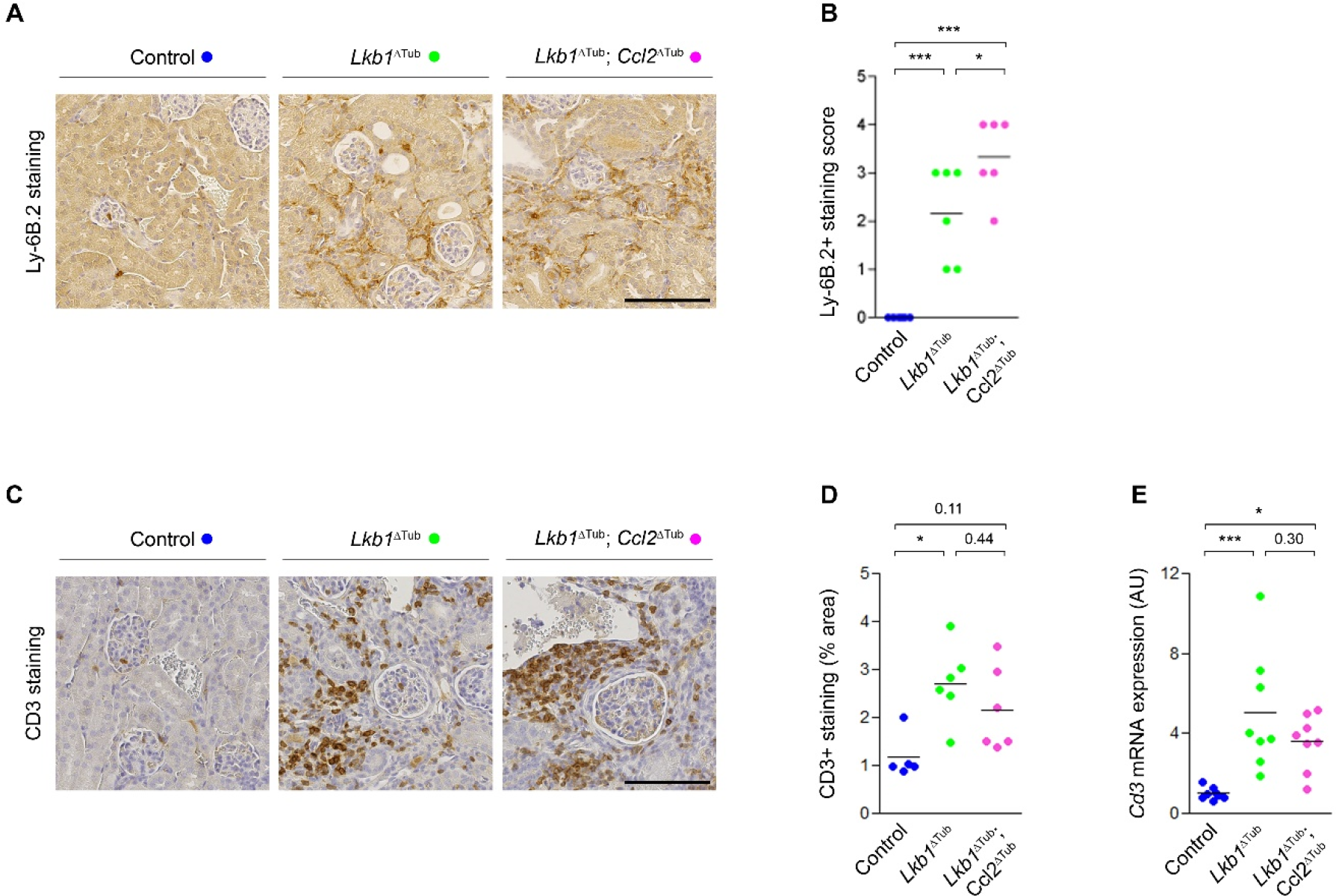
Lymphocytes and neutrophils infiltrate LKB1 deficient kidneys. **(A)** Representative images of Ly-6B.2 (neutrophils) immunostaining of kidney sections from control, *Lkb1*^ΔTub^ and *Lkb1*^ΔTub^; *Ccl2*^ΔTub^ mice at 10 weeks. Scale bar: 100µm. **(B)** Quantification of Ly-6B.2 positive staining area in whole sections from 10 weeks old control, *Lkb1*^ΔTub^ and *Lkb1*^ΔTub^; *Ccl2*^ΔTub^ mice. **(C)** Representative images of CD3 (T lymphocytes) immunostaining of kidney sections from control, *Lkb1*^ΔTub^ and *Lkb1*^ΔTub^; *Ccl2*^ΔTub^ mice at 10 weeks. Scale bar: 100µm. **(D)** Quantification of CD3 positive staining area in whole sections from 10 weeks old control, *Lkb1*^ΔTub^ and *Lkb1*^ΔTub^; *Ccl2*^ΔTub^ mice. **(E)** *Cd3* mRNA expression in kidneys from control, *Lkb1*^ΔTub^ and *Lkb1*^ΔTub^; *Ccl2*^ΔTub^ mice at 10 weeks. **(B, D-E)** Each dot represents one individual mouse. Bars indicate mean. One-way ANOVA followed by Tukey-Kramer test, * P<0.05, *** P<0.001. AU: arbitrary unit.

### Identifying the inflammatory network associated with loss of *NPHP1* function in humans

Considering the distinct immune cell populations infiltrating the kidneys of *Lkb1*^Δtub^ mice, we aimed to determine if such phenomena were also found in NPH patients. CD15 and CD3 immunostaining revealed increased neutrophils and T cells recruitment to the renal parenchyma of NPH patients compared to controls (**Figure 6A-B**). In contrast, these immune cell populations were less abundant in kidney biopsies from patients suffering from acquired non-immune chronic kidney disease caused by diabetes or hypertension (**Figure 6A-B**), supporting the notion that a specific inflammatory network delineates human NPH. To determine if the cytokine network that we identified as features of both *Lkb1*^Δtub^ and *Glis2*^lacZ/lacZ^ mouse models of NPH was relevant to human NPH, we quantified the mRNA abundance of its components in UREC from *NPHP1* patients and controls. As there are no human orthologs for CCL6, 9 and 12, we focused our analysis on the 14 other inflammatory mediators. Indeed, we found 8 transcripts upregulated in human tubular cells with *NPHP1* mutations (*CCL5*, *CXCL1*, *CXCL10*, *CXCL16*, *CXCL17*, *CX3CL1*, *IL1RN* and *LGALS9),* while two were not differentially regulated (*IL33* and *CCL19*) and 4 were not detected (*CXCL9*, *CXCL12*, *CXCL14*, *IL34)* (**Figure 6C-L**). No difference of these cytokines mRNA expression was observed between age-matched controls and relatives with normal kidney function (**Supplementary Figure 4**). In addition, the mRNA level of CXCL1, CXCL17, IL1RN and LGALS9 were significantly higher in UREC derived from NPH patients than from patients suffering from other CKD. CX3CL1, CXCL10 and CXCL16 show the same trend, approaching but not reaching significance (**Figure 6C-L**). These data reveal that, in addition to previously described CCL2 upregulation and macrophage infiltration, NPH is characterized by a complex and specific cytokine signature, which is associated with kidney infiltration by T cells and neutrophils.

**Figure 6.**
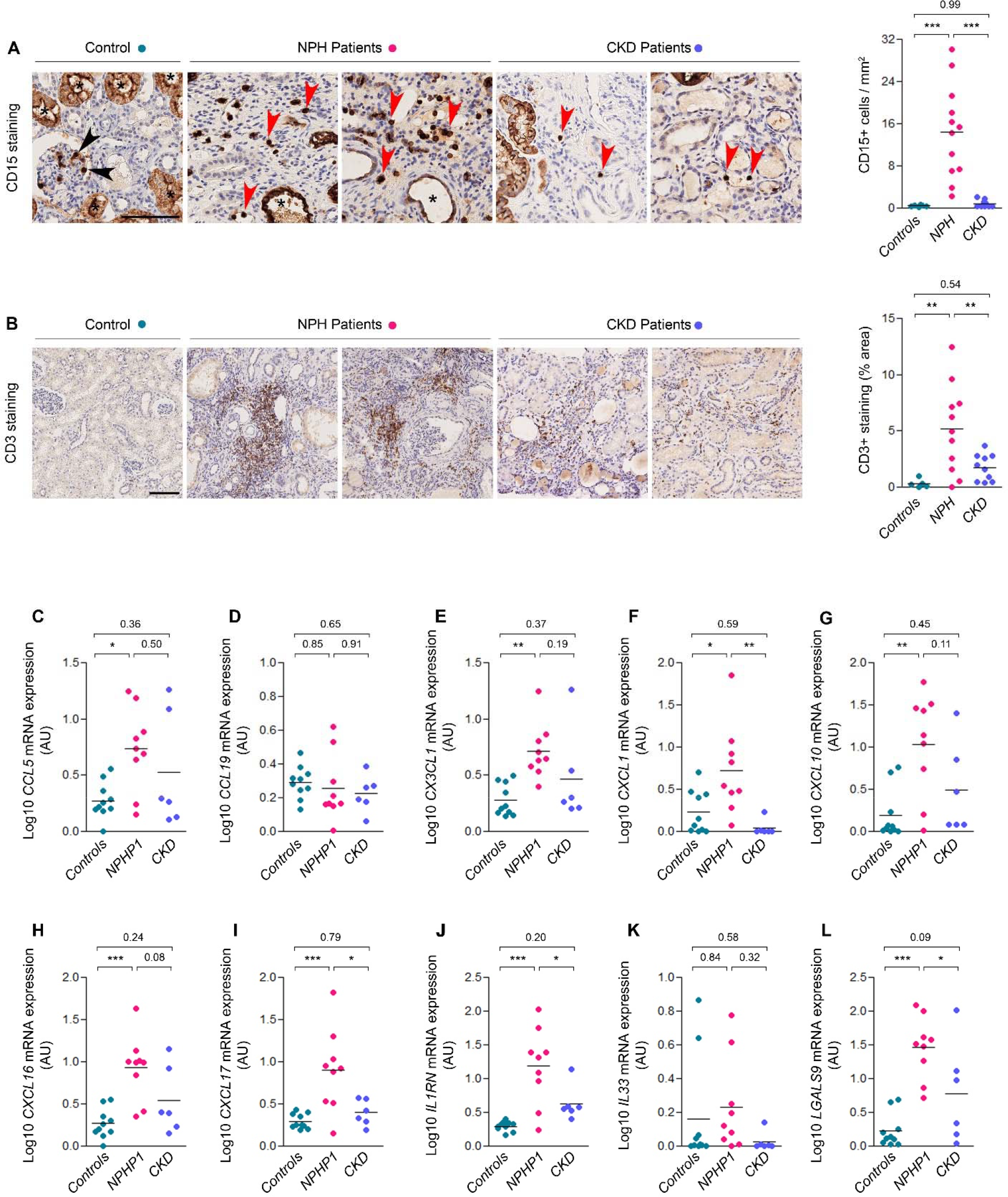
T cells and neutrophils kidney infiltration and inflammatory mediator induction in tubular cells are conserved features of patients suffering from nephronophthisis. **(A)** Representative images and quantification of CD15 (neutrophils) immunostaining of kidney biopsies from control, NPH and chronic kidney disease (CKD) patients. Scale bar: 100µm. Asterisk: unspecific tubular staining, black arrow: intravascular neutrophils in a glomerulus, red arrow, interstitial neutrophils. **(B)** Representative images and quantification of CD3 (T lymphocytes) immunostaining of kidney biopsies from control, NPH and CKD patients. Scale bar: 100µm. (C-L) *CCL5* (C), *CCL19* (D), *CX3CL1* (E), *CXCL1* (F), *CXCL10* (G), *CXCL16* (H), *CXCL17* (I), *IL1RN* (J), *IL33* (K), *LGALS9* (L) mRNA expression expressed in Log10 in primary UREC from controls (Controls), *NPHP1* patients (NPHP1) and non-ciliopathy CKD patients (CKD). **(A-L)** Each dot represents one individual. Bars indicate mean. One-way ANOVA followed by Tukey-Kramer test, * P<0.05, ** P < 0.01, *** P<0.001. AU: arbitrary unit.

## DISCUSSION

In sharp contrast with the abundant literature regarding the molecular mechanisms of disease progression in ADPKD, insights into the pathophysiology of NPH remain scarce. Despite a growing list of causative gene defects, it is unknown how dysfunction of ciliary NPHP complexes results in the unique renal manifestations of NPH. Some data point to a defect in renal development but a prominent feature is severe progressive fibrosis, resulting in renal failure during the second decade of life. The study of the molecular pathogenesis of NPH is hampered by the lack of orthologous mouse models recapitulating the fibrotic disease observed in most NPH patients. Indeed, neither *Nphp1*^-/-^ nor *Nphp4*^-/-^ mice^28, 29^ develop renal fibrosis and only few mouse models orthologous to rare forms of the disease phenocopy the human pathology^18, 27, 30^. An additional factor limiting insights into the disease is the reduced availability of kidney tissues from NPH patients. As genetic testing has become more widely available, kidney biopsies are less and less performed in NPH.

Renal inflammation has emerged as an important mediator of renal fibrosis in acquired chronic kidney disease^12^. In the renal ciliopathy ADPKD, CCL2 dependent macrophage recruitment promotes disease progression^16, 17^. Having previously shown that both NPHP1 and LKB1 repress CCL2 expression *in vitro* and that *Lkb1* inactivation in mice results in an NPH-like phenotype preceded by CCL2 upregulation and macrophage recruitment^16^, we first sought to determine if CCL2 induction and macrophage recruitment were also hallmarks of human NPH. In line with the data gathered from *Lkb1*^Δtub^ mice, we observed significant macrophage infiltration in kidney biopsies from NPH patients and enhanced CCL2 levels in the urine. We then assessed the role of CCL2 in the *Lkb1*^Δtub^ phenotype by inactivating *Ccl2* specifically in tubular cells. Unexpectedly, and contrary to its effect in ADPKD models, *Ccl2* invalidation had no detectable impact on the NPH phenotype of *Lkb1*^Δtub^ mice. Strikingly, CCL2 inactivation did not reduce macrophage infiltration, nor inflammatory cytokine expression in LKB1 deficient kidneys. Beyond ADPKD, CCL2 is instrumental to macrophage recruitment and kidney damage in a range of experimental renal disease including unilateral ureteric obstruction, acute kidney injury, diabetic nephropathy, subtotal nephrectomy^31^ or in response to renal infection^32^. Thus, the fact that renal inflammation and macrophage recruitment in *Lkb1*^Δtub^ mice is independent of tubular CCL2 suggests that the mechanisms driving renal inflammation in NPH are distinct from those implicated in most chronic kidney disease models. To get further insight into the nature of renal inflammation in NPH, we analyzed the population of immune cells infiltrating the kidneys in human NPH and *Lkb1*^Δtub^ mice. In both case, macrophage infiltration was associated with the recruitment of T cells and neutrophils, by opposition to ADPKD where macrophages are the prominent drivers of inflammation^15–17^. To assess the mediators driving early NPH pathology, we analyzed common regulated genes in the transcriptome of *Glis2*^lacZ/lacZ^ and *Lkb1*^Δtub^ kidneys, two genetically distinct mouse models of NPH. Unbiased pathway enrichment analysis of their common upregulated genes revealed a striking preponderance of inflammatory processes, pinpointing that early inflammation is a prominent feature of experimental NPH. Indeed, we found a large number of soluble mediators of immune activation in urinary tubular cells derived from NPH patients. A subset of 7 pro-inflammatory cytokines were specifically upregulated in *NPHP1* UREC and NPH mouse models as compared to urinary tubular cells derived from non-ciliopathy CKD patients : CX3CL1, CXCL1, CXCL10, CXCL16 and CXCL17, IL1RN and LGALS9. Notably, CXCL10, CXCL16 and IL1RN have also been found to be induced in a rodent orthologous model of ADPKD^25^, suggesting that they are part of a broader signature of early renal ciliopathies. In contrast, differential expression of CX3CL1, CXCL1, CXCL17 and LGALS9 transcripts were not. Thus, these cytokines appear more specific of the early renal inflammation observed in NPH.

Fractalkine/CX3CL1, which binds to CX3CR1, plays a major role in the recruitment of resident macrophage during kidney development^33^. However, it has also been shown to support renal monocytes recruitment in the context of renal injury and CX3CL1/CX3CR1 inhibition has been generally shown to reduce renal fibrosis in different models of renal injury^34–36^. In particular, knocking out *Cx3cr1* in mice, Peng and colleagues achieved to decrease renal fibrosis in a model of unilateral ureteral obstruction-induced kidney injury, which was due to a lower infiltration of pro-inflammatory macrophages^36^. In addition, they showed that CX3CL1/CX3CR1 signaling enhanced macrophage survival, which is consistent with previous reports^36, 37^. Ischemia-reperfusion injury-mediated renal fibrosis was also decreased using anti-CX3CR1 antibody in mice, and *Cx3cr1* inactivation reduced diabetic nephropathy-associated fibrosis in mice through a downregulation of a TGF-β/CX3CR1 positive feedback loop and a lower macrophage infiltration in diabetic kidneys^35^. CXCL1 induces the recruitment of various immune cell types, especially neutrophils, through its receptor CXCR2. The CXCL1/CXCR2/neutrophil axis plays an important role in the pathology of acute kidney inflammation in different mouse models^38–40^. Interestingly, we observed concordant neutrophil infiltration in human NPH kidneys and mice, even at an early stage of the disease. In contrast, neutrophil recruitment does not occur in ADPKD^16^. As neutrophils are known to promote renal fibrosis^41^ it is plausible that CXCL1 mediated neutrophil recruitment contributes to phenotypic changes in NPH.

Our findings add to the recently identified expression of CXCL17 in the kidney^42^, the last identified member of the CXC chemokine family in mammals^43, 44^. CXCL17 attracts professional antigen presenting cells^45^, including macrophages, through its suspected receptor GPR35/CXCR8^46^, even though a second unidentified G protein-coupled receptor may mediate CXCL17-dependent signaling^47^. Although very little is known about the role of CXCL17 in the kidney, single-cell RNA sequencing analysis revealed that its expression is enhanced at an early time point in murine tubulointerstitial fibrosis model^42^. It is consistent with the increased levels of CXCL17 observed in interstitial pulmonary fibrosis, where it is thought to recruit immune cells that in turn produce pro-inflammatory cytokines^47^. Further research may determine if this axis could participate in the pathogenesis of NPH and its fibrotic features.

LGALS9, a mammalian β-galactoside binding lectin (Gal-9), was first isolated from murine embryonic kidney^48^. LGALS9 is known to participate in numerous cellular processes including the induction of apoptosis of different immune cells, particularly cytotoxic T lymphocytes when bound to its surface receptor TIM3/HAVCR2^49^. The precise role of LGALS9 in the kidney has not been described so far, however a recent study showed that anti-TIM3 antibody ameliorates kidney injury and decreased macrophage infiltration in ischemic mouse model^50^. This is consistent with a concomitant study that showed increased levels of soluble TIM3 and soluble Gal-9 in the blood of patients with kidney transplantation-related renal dysfunction^51^. In addition, upregulation of serum Gal-9 is closely related to glomerular filtration rate decrease in patients with type 2 diabetes^52^. Thus, circulating Gal-9 and TIM3 may be useful biomarkers to monitor predict GFR decline in nephronophthisis. Further studies are needed to assess their function and interplay in the disease. Besides, Gal-9-dependent signaling may participates in switching macrophages from pro-inflammatory M1 phenotype to its anti-inflammatory M2 counterpart, raising the possibility that Gal-9/TIM3 is involved in the polarization of macrophages in NPH^53, 54^.

Thus, this work identified a specific network of commonly regulated cytokines that represent plausible mediators of immune cell recruitment to NPH kidneys beyond CCL2. Unexpectedly, we identified inflammation as the predominant signature in two independent models of NPH and we find that this holds true in human patients. The parallelism between the inflammatory pathways activated in the early diseased kidneys from *Lkb1*^ΔTub^ or *Glis2*^lacZ/lacZ^ mice in one hand and those found in the UREC from NPH patients in the other hand, along with the protective effect of TLR2/MYD88 inhibition in *Glis2*^lacZ/lacZ^ mice, suggest causality. However, we cannot exclude that chemokine expression and immune cell recruitment are innocent bystanders of an undefined process leading to kidney damage in NPH. In addition certain immune cells may also be protective such as T lymphocytes in the context of ADPKD^55^.

This work establishes renal inflammation as a prominent feature of human NPH and identifies specific mediators of this process that are common to NPH mouse models and patients. Addressing the precise function of these mediators in NPH in the future will help to characterize the underlying processes responsible for renal deterioration in this orphan disease and to evaluate inflammation as a potential therapeutic target.

## ACKNOWLEDGEMENTS

We thank the technicians from the mouse histology facility (S.F.R Necker INSERM US24, Paris, France) and the department of Pathology (Necker Hospital, Paris, France) for technical assistance. We are grateful to the patients and their families for their participation. We thank Pauline Krug, Olivia Boyer, Nathalie Biebuyck, Saoussen Krid, Marina Charbit, Romain Berthaud and Guillaume Lezmi (Pediatric Nephrology, Necker Hospital, AP-HP, Paris, France), Aurélie Hummel (Adult Nephrology, Necker Hospital, AP-HP, Paris, France), Amélie Lezmi Ryckewaert and Sophie Taque (Pediatric Hematology and Oncology, Hôpital Universitaire, Rennes, France), Odile Boespflug-Tanguy (Centre de Compétence des Leucodystrophies et Leucoencéphalopathies de Cause Rare, Pôle Femme et Enfant, Hôpital Estaing, Centre Hospitalier Universitaire de Clermont-Ferrand, Clermont-Ferrand, France), Jérôme Harambat and Brigitte Llanas (Department of Pediatrics, Bordeaux University Hospital, Bordeaux, France), Bruno Ranchin (Pediatric Nephrology, Centre Hospitalier Universitaire de Lyon, Bron, France), Elodie Merieau (Centre Hospitalier Régional Universitaire de Tours, Tours, France), Marc Fila (Centre Hospitalier Universitaire de Montpellier, Montpellier, France) and Tory Kalman (Semmelweis University, Budapest, Hungary) who helped the follow-up of patients. We thank Corinne Antignac and Laurence Heidet (Department of Genetics, Necker Hospital, AP-HP, Paris, France) for genetic diagnostic. We thank the Department of Clinical Research at Imagine Institute with the sponsorship team that facilitates and structures the set-up of the clinical research projects and the investigation team that prepares and ensures the follow-up of the clinical trials.

Marceau Quatredeniers and Amandine Viau were supported by by a public grant “RHU-C’IL-LICO” overseen by the French National Research Agency (ANR) as part of the second “Investissements d’Avenir” program (reference: ANR-17-RHUS-0002), Frank Bienaimé was supported by EMBO ALTF 927-2013, E. Wolfgang Kuehn was supported by Deutsche Forschungsgemeinschaft KU1504/7-1 and KU1504/8-1, Sophie Saunier was supported by the Institut National de la Santé et de la Recherche Médicale (INSERM), the Ministère de l’Education Nationale de la Recherche et de la Technologie (MRT), by a State funding from the Agence Nationale de la Recherche under “Investissements d’avenir” program (ANR-10-IAHU-01) and by a public grant “RHU-C’IL-LICO” overseen by the French National Research Agency (ANR) as part of the second “Investissements d’Avenir” program (reference: ANR-17-RHUS-0002).

## AUTHOR CONTRIBUTIONS

A.V designed the study; M.Q, F.B, G.F, P.I, E.P, K.B, E.B, S.C, F.L, S.B, T.NK and A.V performed experiments and analyzed data; S.S, R.S and M-C.G participated to human material collection; M.Q, F.B, E.W.K, S.S and A.V drafted and revised the manuscript; all authors approved the final version of the manuscript.

