## Supplementary Material for "The renal inflammatory network of nephronophthisis"

### Table of contents

**Supplementary Figure 1**: Primary UREC derived from control individuals.

**Supplementary Figure 2**: GSEA-enrichment map of 1,262 common regulated genes.

**Supplementary Figure 3**: Heatmaps of the 17 pro-inflammatory cytokines identified.

**Supplementary Figure 4**: Pro-inflammatory cytokine levels in primary UREC derived from control individuals.

**Supplementary Table 1**: Genetic and clinical data from patients suffering from juvenile nephronopthisis, chronic kidney disease and healthy controls and relatives.

**Supplementary Table 2**: Primer used for qRT-PCR.

**Supplementary Table 3**: Common up- and down-regulated genes in *Glis2*$^{lacZ/lacZ}$ mouse kidney and *Lkb1*$^{\Delta Tub}$ mouse kidney with FDR<0.05 in *Lkb1*$^{\Delta Tub}$ dataset.

**Supplementary Table 4**: GSEA pathways (biological processes) upregulated for the 1,262 common regulated genes between *Glis2*$^{lacZ/lacZ}$ and *Lkb1*$^{\Delta Tub}$ mouse kidney datasets.

**Supplementary Table 5**: GSEA pathways (biological processes) downregulated for the 1,262 common regulated genes between *Glis2*$^{lacZ/lacZ}$ and *Lkb1*$^{\Delta Tub}$ mouse kidney datasets.

**Supplementary Table 6**: Common secreted cytokine-coding genes linked to immune response/inflammation among the 823 common upregulated genes between *Glis2*$^{lacZ/lacZ}$ and *Lkb1*$^{\Delta Tub}$ mouse kidney datasets.

**Supplementary Table 7**: Expression matrix of the 17 identified cytokines from *Lkb1*$^{\Delta Tub}$ and *Pkd2*$^{\Delta Tub}$ mouse kidney datasets.

**Supplementary Figure 1**

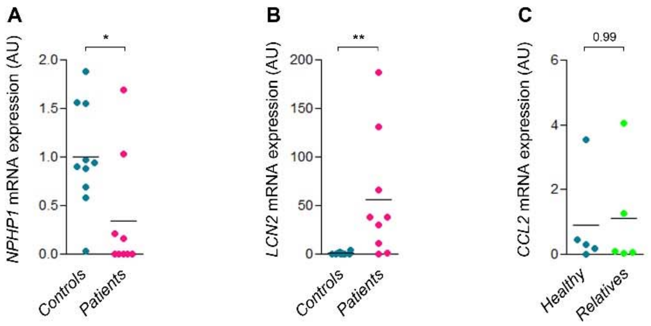


**Supplementary Figure 1. Primary UREC derived from control individuals.**

**(A-B)** *NPHP1* (A) and *LCN2* (B) mRNA expression in primary UREC derived from urine from controls (Controls) and NPHP1 patients (Patients). **(C)** *CCL2* mRNA expression in primary UREC derived from urine from controls either healthy controls (Healthy) or relatives from NPHP1 patients (Relatives). **(A-C)** Each dot represents one individual. Bars indicate mean. Mann-Whitney t test, * P<0.05, **P < 0.01. AU: arbitrary unit.

**Supplementary Figure 2**

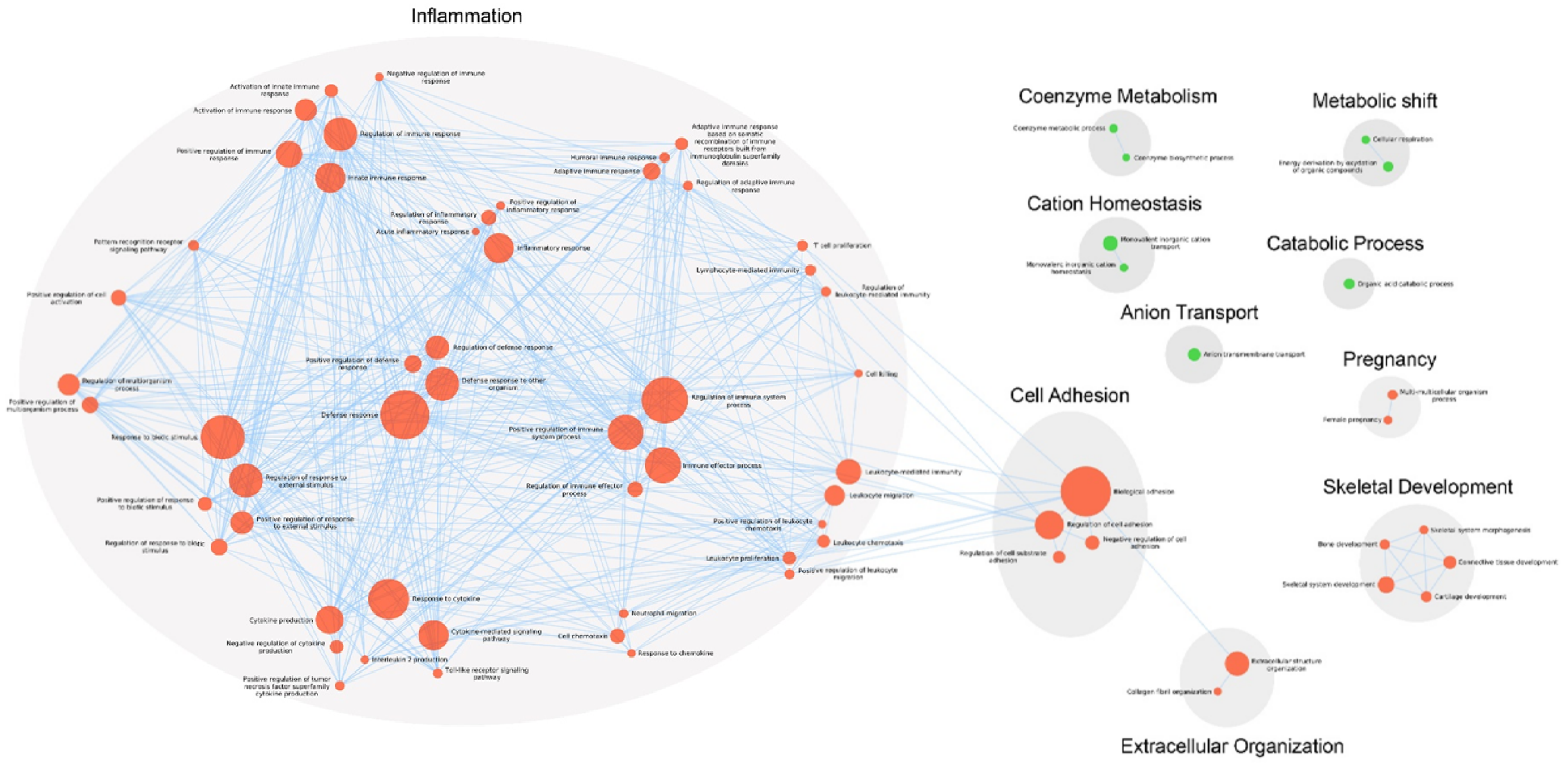

**Supplementary Figure 2. GSEA-enrichment map of 1,262 common regulated genes.**

Enrichment map of pathways enriched in the 1,262 common regulated genes between *Glis2*$^{lacZ/lacZ}$ and *Lkb1*$^{\Delta Tub}$ kidney transcriptomes. Each node represents an enriched pathway; node internal color reflects the normalized enrichment score (NES) for significant upregulated (shade of red) and downregulated (shade of green) pathways, and external color reflects the FDR (shade of purple). Edge width represent the number of genes overlapping between two pathways, according to similarity coefficient.

Supplementary Figure 3

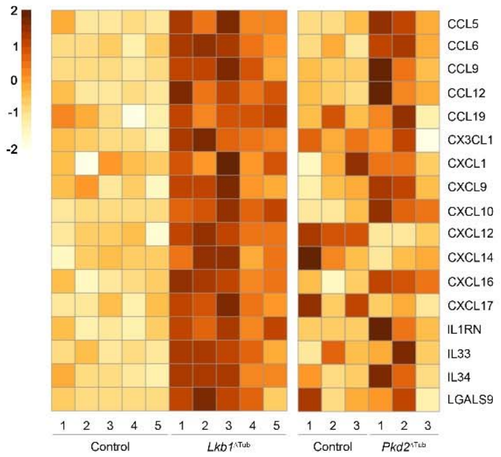


**Supplementary Figure 3. Heatmaps of the 17 pro-inflammatory cytokines identified.**

Heatmaps show Z-scores computed on log-normalized expression of these cytokines in the *Lkb1*$^{\Delta\text{Tub}}$ kidney dataset comparing control (n=5) and *Lkb1*$^{\Delta\text{Tub}}$ (n=5) mice at 5 weeks of age, and in the *Pkd2*$^{\Delta\text{Tub}}$ kidney dataset comparing control (n=3) and *Pkd2*$^{\Delta\text{Tub}}$ (n=3) mice at 10 weeks of age. Each row indicates individual gene and each column shows individual mouse. Heatmaps were generated using the package pheatmap v1.0.12.

**Supplementary Figure 4**

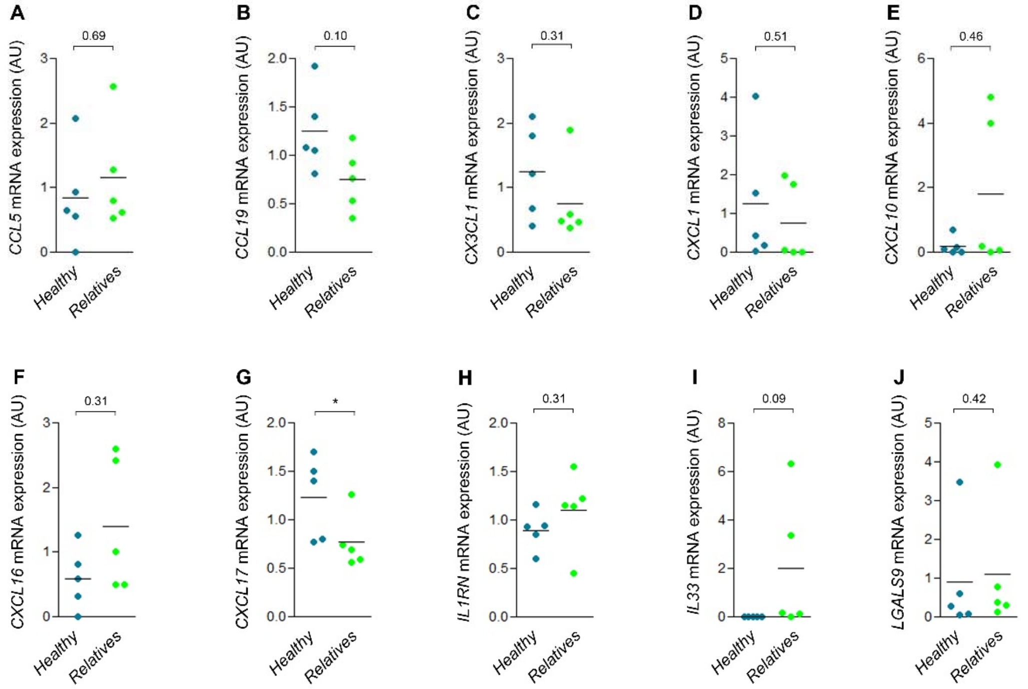


**Supplementary Figure 4. Pro-inflammatory cytokine levels in primary UREC derived from control individuals. (A-J)** *CCL5* (A), *CCL19* (B), *CX3CL1* (C), *CXCL1* (D), *CXCL10* (E), *CXCL16* (F), *CXCL17* (G), *IL1RN* (H), *IL33* (I), *LGALS9* (J) mRNA expression in primary UREC derived from urine from controls either healthy controls (Healthy) or relatives from NPHP1 patients (Relatives). **(A-J)** Each dot represents one individual. Bars indicate mean. Mann-Whitney t test, * P<0.05. AU: arbitrary unit.

**Supplementary Table 1: Genetic and clinical data from patients suffering from juvenile nephronopthisis, chronic kidney disease and healthy controls and relatives.**

NPH: nephronophthisis, CKD: chronic kidney disease, CAKUT: congenital anomalies of the kidney and urinary tract, F: female, M: male, Het: heterozygous, Hom: homozygous, Del: deletion, eGFR: estimated glomerular filtration rate, NA: not available.
GFR is estimated according to Schwartz formula.

**a. Human kidney sections**

| Patient ID | Group ID | Gender | Clinical Status | Causal gene | Genetic Status | IFTA (Interstitial Fibrosis Tubular Atrophy) % |
|---|---|---|---|---|---|---|
| **Control 1** | Control | M | Kidneys donated for transplantation but unsuitable for implantation | / | / | 5% |
| **Control 2** | Control | NA | Kidneys donated for transplantation but unsuitable for implantation | / | / | 0% |
| **Control 3** | Control | NA | Kidneys donated for transplantation but unsuitable for implantation | / | / | 0% |
| **Control 4** | Control | M | Kidneys donated for transplantation but unsuitable for implantation | / | / | NA |
| **Control 5** | Control | NA | Kidneys donated for transplantation but unsuitable for implantation | / | / | 5% |
| **Control 6** | Control | M | Minimal change disease | / | / | NA |
| **NPH 1** | NPH Patient | M | Juvenile Nephronophthisis | *NPHP1* | Hom Del | NA |
| **NPH 2** | NPH Patient | F | Juvenile Nephronophthisis | *NPHP1* | Hom Del | NA |
| **NPH 3** | NPH Patient | F | Juvenile Nephronophthisis | *NPHP1* | Hom Del | 40% |
| **NPH 4** | NPH Patient | M | Juvenile Nephronophthisis | *NPHP1* | Hom Del | 20% |
| **NPH 5** | NPH Patient | M | Juvenile Nephronophthisis | *NPHP1* | Hom Del | 40% |
| **NPH 6** | NPH Patient | M | Juvenile Nephronophthisis | *NPHP1* | Hom Del | 50% |
| **NPH 7** | NPH Patient | M | Juvenile Nephronophthisis | *NPHP1* | Hom Del | 30% |
| **NPH 8** | NPH Patient | F | Juvenile Nephronophthisis | *NPHP1* | Hom Del | 30% |
| **NPH 9** | NPH Patient | F | Juvenile Nephronophthisis | *NPHP4* | Hom c.1972 C>T p.R658* STOP | 30% |
| **NPH 10** | NPH Patient | M | Juvenile Nephronophthisis | *NPHP6* | compound c.5649InsA p.L1884Tfs*23 Frameshift + c.5850delT p.F1950Lfs*15 Frameshift | 50% |
| **NPH 11** | NPH Patient | F | Juvenile Nephronophthisis | *NPHP8* | Hom 2083 G>C p.A695P Missense | NA |
| **NPH 12** | NPH Patient | F | Juvenile Nephronophthisis | *NPHP8* | Hom 2083 G>C p.A695P Missense | 0% |
| **CKD-1** | CKD Patient | M | Diabetic Nephropathy | / | / | 20% |
| **CKD-2** | CKD Patient | M | Diabetic Nephropathy | / | / | NA |
| **CKD-3** | CKD Patient | F | Diabetic Nephropathy | / | / | 15% |
| **CKD-4** | CKD Patient | F | Diabetic Nephropathy | / | / | 10% |
| **CKD-5** | CKD Patient | F | Diabetic Nephropathy | / | / | 20% |
| **CKD-6** | CKD Patient | M | Hypertensive Nephrosclerosis | / | / | 20% |
| **CKD-7** | CKD Patient | M | Hypertensive Nephrosclerosis | / | / | 50% |
| **CKD-8** | CKD Patient | M | Hypertensive Nephrosclerosis | / | / | 40% |
| **CKD-9** | CKD Patient | M | Hypertensive Nephrosclerosis | / | / | 25% |
| **CKD-10** | CKD Patient | M | Hypertensive Nephrosclerosis | / | / | 50% |
| **CKD-11** | CKD Patient | F | Hypertensive Nephrosclerosis | / | / | 50% |

**b. Primary human UREC**

| Patient ID | Group ID | Gender | Clinical Status | Causal gene | Genetic Status | eGFR (ml/min/1.73m2) |
|---|---|---|---|---|---|---|
| **Control 7** | Healthy control | F | Pediatric patient without kidney disease | / | / | 121 |
| **Control 8** | Healthy control | F | Pediatric patient without kidney disease | / | / | NA |
| **Control 9** | Healthy control | M | Pediatric patient without kidney disease | / | / | NA |
| **Control 10** | Healthy control | M | CAKUT with normal kidney function | / | / | 93 |
| **Control 11** | Healthy control | M | Solitary functioning kidney with normal kidney function | / | / | 122 |
| **Control 12** | Relative control | M | / | *NPHP1* | Het Del | NA |
| **Control 13** | Relative control | F | / | *NPHP1* | Het c.1078C>T - p.Q360* | NA |
| **Control 14** | Relative control | F | / | *NPHP1* | / | NA |
| **Control 15** | Relative control | F | / | *NPHP1* | Het c.1326+1G>A - p.? | NA |
| **Control 16** | Relative control | F | / | *NPHP1* | Het Del | NA |
| **NPHP1-1** | NPHP1 Patient | M | Juvenile Nephronophthisis | *NPHP1* | Hom Del | 21 |
| **NPHP1-2** | NPHP1 Patient | F | Juvenile Nephronophthisis | *NPHP1* | Hom Del | 54 |
| **NPHP1-3** | NPHP1 Patient | M | Juvenile Nephronophthisis | *NPHP1* | Het Del + Het c.143G>A - p.R48K | 25 |
| **NPHP1-4** | NPHP1 Patient | F | Juvenile Nephronophthisis | *NPHP1* | Het c.1884+1G>T - p.? + Het c.1252-2A>G - p.? | 19.1 |
| **NPHP1-5** | NPHP1 Patient | F | Juvenile Nephronophthisis | *NPHP1* | Het Del + Het c.1326+1G>A - p.? | 16 |
| **NPHP1-6** | NPHP1 Patient | F | Juvenile Nephronophthisis | *NPHP1* | Het Del + Het c.70-1G>A - p.? | 24 |
| **NPHP1-7** | NPHP1 Patient | F | Juvenile Nephronophthisis | *NPHP1* | NA | 17 |
| **NPHP1-8** | NPHP1 Patient | F | Juvenile Nephronophthisis | *NPHP1* | Hom Del | 19 |
| **NPHP1-9** | NPHP1 Patient | F | Juvenile Nephronophthisis | *NPHP1* | Hom Del | 38 |
| **CKD-12** | CKD Patient | M | Solitary functioning kidney with reduced GFR | / | / | 67 |
| **CKD-13** | CKD Patient | M | Hemolytic and uremic syndrome | / | / | 12 |
| **CKD-14** | CKD Patient | F | IgA nephropathy | / | / | 23 |
| **CKD-15** | CKD Patient | M | CAKUT | / | / | 23.4 |
| **CKD-16** | CKD Patient | M | CAKUT | / | / | 32 |
| **CKD-17** | CKD Patient | F | Solitary functioning kidney with reduced GFR | / | / | 68 |

**c. Human urines**

| Patient ID | Group ID | Gender | Clinical Status | Causal gene | Genetic Status | eGFR (ml/min/1.73m2) |
|---|---|---|---|---|---|---|
| **Control 17** | Healthy control | F | Pediatric patient without kidney disease | / | / | NA |
| **Control 18** | Healthy control | M | Pediatric patient without kidney disease | / | / | NA |
| **Control 19** | Healthy control | F | Pediatric patient without kidney disease | / | / | NA |
| **Control 20** | Healthy control | F | Pediatric patient without kidney disease | / | / | NA |
| **Control 21** | Healthy control | F | Pediatric patient without kidney disease | / | / | NA |
| **Control 14** | Relative control | F | / | *NPHP1* | / | NA |
| **Control 22** | Relative control | M | / | *NPHP1* | / | NA |
| **Control 23** | Relative control | M | / | *NPHP1* | Het Del | 118 |
| **NPHP1-1** | NPHP1 Patient | M | Juvenile Nephronophthisis | *NPHP1* | Hom Del | 18.3 |
| **NPHP1-2** | NPHP1 Patient | F | Juvenile Nephronophthisis | *NPHP1* | Hom Del | 48 |
| **NPHP1-3** | NPHP1 Patient | M | Juvenile Nephronophthisis | *NPHP1* | Het Del + Het c.143G>A - p.R48K | 25 |
| **NPHP1-4** | NPHP1 Patient | F | Juvenile Nephronophthisis | *NPHP1* | Het c.1884+1G>T - p.? + Het c.1252-2A>G - p.? | 19.1 |
| **NPHP1-5** | NPHP1 Patient | F | Juvenile Nephronophthisis | *NPHP1* | Het Del + Het c.1326+1G>A - p.? | 16 |
| **NPHP1-6** | NPHP1 Patient | F | Juvenile Nephronophthisis | *NPHP1* | Het Del + Het c.70-1G>A - p.? | 18 |
| **NPHP1-10** | NPHP1 Patient | F | Juvenile Nephronophthisis | *NPHP1* | Hom Del | 17 |
| **NPHP1-11** | NPHP1 Patient | M | Juvenile Nephronophthisis | *NPHP1* | Hom Del | 139 |
| **NPHP1-12** | NPHP1 Patient | F | Juvenile Nephronophthisis | *NPHP1* | Hom Del | 22 |

**Supplementary Table 2: Primer pairs used for qRT-PCR.**

| Gene name | Specie | Forward Primer (5' to 3') | Reverse Primer (5' to 3') |
|---|---|---|---|
| *CCL2* | Human | TCATAGCAGCCACCTTCATTC | CTCTGCACTGAGATCTTCCTATTG |
| *CCL5* | Human | TGCCCACATCAAGGAGTATTT | GATGTACTCCCGAACCCATTT |
| *CCL19* | Human | GGAACTTCCACTACCTTCTCATC | GTCTCTGGATGATGCGTTCTAC |
| *CX3CL1* | Human | GTGCAGCAAGATGACATCAAAG | CTGTCTCGTCTCCAAGATGATTG |
| *CXCL1* | Human | ACTCAAGAATGGGCGGAAAG | CCCTTCTGGTCAGTTGGATTT |
| *CXCL10* | Human | CCATTCTGATTTGCTGCCTTATC | TACTAATGCTGATGCAGGTACAG |
| *CXCL16* | Human | TATGTGCTGTGCAAGAGGAG | TCCATTCTTGGCTCAGGTATTAG |
| *CXCL17* | Human | ACCAAAGGCACCACAGAAAG | CTCCTACAAAGGCAGAGCAAAG |
| *HPRT* | Human | TCTTTGCTGACCTGCTGGATT | GTTGAGAGATCATCTCCACCAATTACT |
| *IL1RN* | Human | CTGTGTCAAGTCTGGTGATGAG | CACTGTCTGAGCGGATGAAG |
| *IL33* | Human | TACTTTATGAAGCTCCGCTCTG | CAGCGAGTACCAGATGTCTTT |
| *LCN2* | Human | GGAGCTGACTTCGGAACTAAAG | CGTCGATACACTGGTCGATTG |
| *LGALS9* | Human | GCTCAGAGGTTCCACATCAA | GGACCACAGCATTCTCATCA |
| *NPHP1* | Human | GGATGCTACAGAAGGCACTATT | ATGTCTGCTGAGAACCTGTATG |
| *PPIA* | Human | GGTCCCAAAGACAGCAGAAA | GTCACCACCCTGACACATAAA |
| *SDHA* | Human | GCCGTGGTCGAGCTAGAAAA | CACGCTGATAAATCTTCCCATCT |
| *TBP* | Human | TCCACAGTGAATCTTGGTTGT | GTGGTTCGTGGCTCTCTTATC |

| Gene name | Specie | Forward Primer (5' to 3') | Reverse Primer (5' to 3') |
|---|---|---|---|
| *Adgre1* | Mouse | CGTCAGGTACGGGATGAATATAAG | ATCTTGGAAGTGGATGGCATAG |
| *Ccl2* | Mouse | AGTAGGCTGGAGAGCTACAA | GTATGTCTGGACCCATTCCTTC |
| *Ccl5* | Mouse | CCAATCTTGCAGTCGTGTTTG | ACCCTCTATCCTAGCTCATCTC |
| *Ccl6* | Mouse | GGCTTTGGAATGTGTCTGGT | CTGGCCCCGTAGTTCTATGA |
| *Ccl9* | Mouse | AGTGGTCTGTGGGACTTTGG | CAGACCTGTGGCTGCATAGA |
| *Ccl12* | Mouse | GATCTTCAGGACCATACTGGATAAG | GAAGGTTCAAGGATGAAGGTTTG |
| *Ccl19* | Mouse | GCCTTCCGCTACCTTCTTAAT | GAGGTGCACAGAGCTGATAG |
| *Cd3* | Mouse | CTGTTCCCAACCCAGACTATG | AAGGCGATGTCTCTCCTATCT |
| *Col1a1* | Mouse | GCCGCAAAGAGTCTACATGTCTAG | TGGCAGATACAGATCAAGCATACC |
| *Cx3cl1* | Mouse | GCTTTGCTCATCCGCTATCA | GTCTTGGACCCATTTCTCCTTC |
| *Cxcl1* | Mouse | CGAAGTCATAGCCACACTCAA | GAGCAGTCTGTCTTCTTTCTCC |
| *Cxcl9* | Mouse | GTTCGAGGAACCCTAGTGATAAG | GTTTGAGGTCTTTGAGGGATTTG |
| *Cxcl10* | Mouse | GGCCATAGGGAAGCTTGAAA | CAGACATCTCTGCTCATCATTCT |
| *Cxcl12* | Mouse | CTCTGCATCAGTGACGGTAAA | CACAGTTTGGAGTGTTGAGGA |
| *Cxcl14* | Mouse | CTGCGAGGAGAAGATGGTTATC | CTTCTCGTTCCAGGCATTGTA |
| *Cxcl16* | Mouse | ATCAGGTTCCAGTTGCAGTC | CATGACCAGTTCCACACTCTT |
| *Cxcl17* | Mouse | CCTTCCTTCTGTTGCTTCCA | TTCCAAGAGCCACCTCCTA |
| *Hprt* | Mouse | GTTAAGCAGTACAGCCCCAAA | AGGGCATATCCAACAACAAACTT |
| *Il1rn* | Mouse | TTGTGCCAAGTCTGGAGATG | CTCAGAGCGGATGAAGGTAAAG |
| *Il33* | Mouse | TGCCTCCCTGAGTACATACA | CTGGTCTTGCTCTTGGTCTTT |
| *Il34* | Mouse | GATATGGACTCTGACCCAAGATAAG | AGCAATCCTGTAGTTGATGGG |
| *Lcn2* | Mouse | GGACCAGGGCTGTCGCTACT | GGTGGCCACTTGCACATTGT |
| *Lgals9* | Mouse | CAGATGCTACGAGGTTCCATATC | GTGTTTCGGACAACAGCATTC |
| *Ppia* | Mouse | GGCTATAAGGGTTCCTCCTTTC | TTTCTCTCCGTAGATGGACCT |
| *Rpl13* | Mouse | GCTCCAAGCTCATCCTGTT | GGTGGCCAGCTTAAGTTCT |
| *Sdha* | Mouse | AGAACATCAGAACTACGCCTAAACATG | CCATTCCCCTGTCGAATGTCT |
| *Tgfb1* | Mouse | GGGAAGCAGTGCCCGAACCC | TGGGGGTCAGCAGCCGGTTA |

**Supplementary Table 3: Common up- and down-regulated genes in *Glis2* $^{lacZ/lacZ}$ mouse kidney and *Lkb1* $^{\Delta Tub}$ mouse kidney with FDR<0.05 in *Lkb1* $^{\Delta Tub}$ dataset.**

FDR: False discovery rate.

| | | Dataset *Glis2* $^{lacZ/lacZ}$ | Dataset *Lkb1* $^{\Delta Tub}$ | | |
|---|---|---|---|---|---|
| **Gene_ID** | **Gene-Name** | **Fold change** | **Fold change** | **p value** | **FDR** |
| **C3** | complement component 3 | 9.43 | 5.48 | 1.46E-08 | 4.06E-05 |
| **Dpt** | dermatopontin | 3.91 | 5.30 | 5.02E-08 | 8.30E-05 |
| **Lyz2** | lysozyme 2 | 7.97 | 5.19 | 5.09E-08 | 8.30E-05 |
| **Grem2** | gremlin 2 homolog, cysteine knot superfamily (Xenopus laevis) | 2.47 | 5.13 | 3.40E-06 | 6.76E-04 |
| **Serpina3n** | serine (or cysteine) peptidase inhibitor, clade A, member 3N | 9.41 | 4.58 | 3.69E-05 | 2.59E-03 |
| **Bmp3** | bone morphogenetic protein 3 | 2.06 | 4.6 | 3.69E-08 | 7.68E-05 |
| **Mmp7** | matrix metallopeptidase 7 | 6.23 | 4.36 | 3.06E-06 | 6.35E-04 |
| **Stc1** | stanniocalcin 1 | 1.36 | 4.08 | 1.00E-09 | 8.32E-06 |
| **Vcam1** | vascular cell adhesion molecule 1 | 5.45 | 3.48 | 6.05E-07 | 2.65E-04 |
| **Lyz1** | lysozyme 1 | 6.85 | 3.39 | 5.61E-07 | 2.59E-04 |
| **Mgp** | matrix Gla protein | 6.08 | 3.38 | 1.06E-05 | 1.26E-03 |
| **Smoc2** | SPARC related modular calcium binding 2 | 2.63 | 3.36 | 1.08E-06 | 3.54E-04 |
| **Aoc1** | amine oxidase, copper-containing 1 | 12.19 | 3.02 | 3.70E-07 | 2.14E-04 |
| **Lum** | lumican | 4.07 | 2.88 | 2.00E-04 | 7.47E-03 |
| **Sprr1a** | small proline-rich protein 1A | 7.68 | 2.86 | 7.73E-06 | 1.03E-03 |
| **Krt19** | keratin 19 | 1.71 | 2.80 | 3.27E-09 | 1.48E-05 |
| **Havcr1** | hepatitis A virus cellular receptor 1 | 45.19 | 2.78 | 3.76E-04 | 1.14E-02 |
| **Fgg** | fibrinogen gamma chain | 8.38 | 2.78 | 6.39E-07 | 2.70E-04 |
| **Mpeg1** | macrophage expressed gene 1 | 2.57 | 2.78 | 2.76E-06 | 5.91E-04 |
| **Ms4a6c** | membrane-spanning 4-domains, subfamily A, member 6C | 5.34 | 2.76 | 7.64E-05 | 4.05E-03 |
| **Fam129a** | family with sequence similarity 129, member A | 2.31 | 2.74 | 2.77E-06 | 5.91E-04 |
| **Serpina10** | serine (or cysteine) peptidase inhibitor, clade A (alpha-1 antiproteinase, antitrypsin), member 10 | 14.34 | 2.73 | 1.24E-03 | 2.40E-02 |
| **Igsf6** | immunoglobulin superfamily, member 6 | 3.01 | 2.70 | 3.39E-06 | 6.76E-04 |
| **Pappa** | pregnancy-associated plasma protein A | 2.21 | 2.68 | 1.05E-06 | 3.54E-04 |
| **Cfi** | complement component factor i | 2.13 | 2.67 | 3.73E-06 | 7.10E-04 |
| **Krt20** | keratin 20 | 17.09 | 2.66 | 1.24E-07 | 1.15E-04 |
| **Mrc1** | mannose receptor, C type 1 | 2.15 | 2.65 | 7.40E-06 | 1.01E-03 |
| **Aldh1a1** | aldehyde dehydrogenase family 1, subfamily A1 | 2.05 | 2.62 | 3.13E-05 | 2.29E-03 |
| **Fn1** | fibronectin 1 | 3.97 | 2.58 | 2.56E-05 | 2.06E-03 |
| **Gabrp** | gamma-aminobutyric acid (GABA) A receptor, pi | 3.00 | 2.58 | 1.01E-05 | 1.21E-03 |
| **Postn** | periostin, osteoblast specific factor | 3.91 | 2.57 | 8.05E-06 | 1.05E-03 |
| **Aldh1a2** | aldehyde dehydrogenase family 1, subfamily A2 | 7.64 | 2.54 | 7.73E-07 | 3.00E-04 |
| **Dcn** | decorin | 3.86 | 2.52 | 1.11E-06 | 3.54E-04 |
| **Scd1** | stearoyl-Coenzyme A desaturase 1 | 1.17 | 2.50 | 7.31E-04 | 1.71E-02 |
| **Ctss** | cathepsin S | 5.92 | 2.48 | 2.49E-05 | 2.03E-03 |
| **Igsf10** | immunoglobulin superfamily, member 10 | 1.15 | 2.47 | 1.63E-06 | 4.03E-04 |
| **Tgfbi** | transforming growth factor, beta induced | 4.51 | 2.47 | 4.36E-06 | 7.78E-04 |
| **Cp** | ceruloplasmin | 2.02 | 2.46 | 4.00E-06 | 7.45E-04 |
| **Col3a1** | collagen, type III, alpha 1 | 3.88 | 2.43 | 4.33E-06 | 7.78E-04 |
| **C1qb** | complement component 1, q subcomponent, beta polypeptide | 3.65 | 2.41 | 9.41E-06 | 1.17E-03 |
| **Mmp2** | matrix metallopeptidase 2 | 3.57 | 2.41 | 5.22E-05 | 3.18E-03 |
| **Gsdmc2** | gasdermin C2 | 2.63 | 2.39 | 1.03E-04 | 4.86E-03 |
| **Edn1** | endothelin 1 | 5.31 | 2.39 | 6.96E-07 | 2.80E-04 |
| **Casp4** | caspase 4, apoptosis-related cysteine peptidase | 3.68 | 2.38 | 3.91E-05 | 2.66E-03 |
| **D17H6S56E-5** | DNA segment, Chr 17, human D6S56E 5 | 3.86 | 2.36 | 5.90E-07 | 2.63E-04 |
| **Tmem173** | transmembrane protein 173 | 4.87 | 2.36 | 3.06E-07 | 2.12E-04 |
| **Cpxm1** | carboxypeptidase X 1 (M14 family) | 4.37 | 2.34 | 3.86E-05 | 2.66E-03 |
| **Serping1** | serine (or cysteine) peptidase inhibitor, clade G, member 1 | 4.39 | 2.33 | 1.47E-05 | 1.50E-03 |

| | | | | | |
|---|---|---|---|---|---|
| **Car15** | carbonic anhydrase 15 | 1.85 | 2.33 | 4.75E-10 | 5.93E-06 |
| **Col1a1** | collagen, type I, alpha 1 | 4.67 | 2.31 | 1.14E-06 | 3.55E-04 |
| **Serpina3g** | serine (or cysteine) peptidase inhibitor, clade A, member 3G | 10.24 | 2.30 | 3.97E-05 | 2.68E-03 |
| **Samd5** | sterile alpha motif domain containing 5 | 5.65 | 2.29 | 3.63E-10 | 5.93E-06 |
| **Scg5** | secretogranin V | 1.60 | 2.28 | 1.18E-05 | 1.32E-03 |
| **Osmr** | oncostatin M receptor | 6.02 | 2.26 | 4.87E-06 | 8.07E-04 |
| **Gsdmc3** | gasdermin C3 | 1.37 | 2.26 | 4.71E-04 | 1.31E-02 |
| **Dcdc2a** | doublecortin domain containing 2a | 2.97 | 2.23 | 1.18E-06 | 3.55E-04 |
| **Ly86** | lymphocyte antigen 86 | 2.96 | 2.21 | 2.36E-05 | 1.94E-03 |
| **Lox** | lysyl oxidase | 3.19 | 2.19 | 9.31E-04 | 2.01E-02 |
| **H2-Aa** | histocompatibility 2, class II antigen A, alpha | 6.40 | 2.17 | 4.93E-06 | 8.09E-04 |
| **Fgb** | fibrinogen beta chain | 1.89 | 2.14 | 1.72E-05 | 1.62E-03 |
| **Clec4a1** | C-type lectin domain family 4, member a1 | 2.84 | 2.12 | 1.45E-06 | 3.82E-04 |
| **Ccl9** | chemokine (C-C motif) ligand 9 | 13.65 | 2.12 | 1.17E-05 | 1.31E-03 |
| **Abi3bp** | ABI gene family, member 3 (NESH) binding protein | 3.42 | 2.10 | 4.58E-05 | 2.94E-03 |
| **Col1a2** | collagen, type I, alpha 2 | 3.58 | 2.10 | 4.96E-06 | 8.10E-04 |
| **Ms4a7** | membrane-spanning 4-domains, subfamily A, member 7 | 5.31 | 2.09 | 4.73E-07 | 2.40E-04 |
| **Spink8** | serine peptidase inhibitor, Kazal type 8 | 1.36 | 2.09 | 4.02E-04 | 1.18E-02 |
| **Mmp14** | matrix metallopeptidase 14 (membrane-inserted) | 3.80 | 2.09 | 1.68E-05 | 1.61E-03 |
| **Pde3b** | phosphodiesterase 3B, cGMP-inhibited | 1.04 | 2.09 | 2.94E-07 | 2.10E-04 |
| **Clca3a2** | chloride channel calcium activated 3A2 | 6.11 | 2.08 | 1.00E-06 | 3.52E-04 |
| **Arg2** | arginase type II | 1.13 | 2.07 | 7.90E-06 | 1.04E-03 |
| **Cd163** | CD163 antigen | 1.32 | 2.06 | 1.52E-04 | 6.18E-03 |
| **Spp1** | secreted phosphoprotein 1 | 3.24 | 2.04 | 1.05E-06 | 3.54E-04 |
| **Lgi2** | leucine-rich repeat LGI family, member 2 | 8.30 | 2.04 | 2.85E-05 | 2.15E-03 |
| **Ms4a6b** | membrane-spanning 4-domains, subfamily A, member 6B | 7.34 | 2.01 | 1.69E-05 | 1.61E-03 |
| **Pld4** | phospholipase D family, member 4 | 4.52 | 1.98 | 1.09E-05 | 1.27E-03 |
| **Lipg** | lipase, endothelial | 1.20 | 1.98 | 1.04E-04 | 4.88E-03 |
| **Tubb2b** | tubulin, beta 2B class IIB | 1.76 | 1.98 | 3.69E-07 | 2.14E-04 |
| **Lbp** | lipopolysaccharide binding protein | 1.28 | 1.97 | 4.68E-05 | 2.96E-03 |
| **Mfap5** | microfibrillar associated protein 5 | 3.68 | 1.97 | 7.70E-05 | 4.07E-03 |
| **Pltp** | phospholipid transfer protein | 3.71 | 1.97 | 1.63E-05 | 1.60E-03 |
| **Arhgap36** | Rho GTPase activating protein 36 | 1.52 | 1.97 | 3.87E-05 | 2.66E-03 |
| **Alox5ap** | arachidonate 5-lipoxygenase activating protein | 5.26 | 1.96 | 4.52E-05 | 2.94E-03 |
| **Tlr7** | toll-like receptor 7 | 1.46 | 1.96 | 2.16E-05 | 1.83E-03 |
| **Cdh3** | cadherin 3 | 1.46 | 1.94 | 1.06E-07 | 1.15E-04 |
| **Krt18** | keratin 18 | 2.55 | 1.94 | 1.87E-05 | 1.69E-03 |
| **Cd4** | CD4 antigen | 1.92 | 1.94 | 3.11E-05 | 2.29E-03 |
| **H2-Ab1** | histocompatibility 2, class II antigen A, beta 1 | 5.82 | 1.93 | 3.11E-04 | 1.01E-02 |
| **Fcer1g** | Fc receptor, IgE, high affinity I, gamma polypeptide | 5.00 | 1.93 | 1.81E-06 | 4.36E-04 |
| **H2-Eb1** | histocompatibility 2, class II antigen E beta | 5.48 | 1.93 | 7.36E-05 | 3.97E-03 |
| **Aqp2** | aquaporin 2 | 1.02 | 1.92 | 3.87E-04 | 1.16E-02 |
| **Slc25a24** | solute carrier family 25 (mitochondrial carrier, phosphate carrier), member 24 | 3.83 | 1.91 | 3.19E-06 | 6.48E-04 |
| **Cd53** | CD53 antigen | 1.58 | 1.90 | 1.53E-04 | 6.20E-03 |
| **Akr1b8** | aldo-keto reductase family 1, member B8 | 5.56 | 1.90 | 5.62E-06 | 8.65E-04 |
| **Sprr2f** | small proline-rich protein 2F | 2.66 | 1.89 | 4.91E-07 | 2.40E-04 |
| **Fga** | fibrinogen alpha chain | 3.42 | 1.88 | 7.09E-06 | 9.78E-04 |
| **Casp12** | caspase 12 | 3.37 | 1.88 | 1.61E-05 | 1.59E-03 |
| **Tnfsf13b** | tumor necrosis factor (ligand) superfamily, member 13b | 4.24 | 1.87 | 5.18E-06 | 8.37E-04 |
| **Fbn1** | fibrillin 1 | 4.45 | 1.87 | 1.46E-04 | 6.07E-03 |
| **Cyp1b1** | cytochrome P450, family 1, subfamily b, polypeptide 1 | 3.82 | 1.87 | 3.25E-04 | 1.04E-02 |
| **Antxr1** | anthrax toxin receptor 1 | 1.27 | 1.86 | 2.74E-04 | 9.23E-03 |
| **Lilrb4a** | leukocyte immunoglobulin-like receptor, subfamily B, member 4A | 2.03 | 1.85 | 4.61E-05 | 2.95E-03 |
| **C1s2** | complement component 1, s subcomponent 2 | 3.20 | 1.85 | 1.15E-05 | 1.30E-03 |

| | | | | | |
|---|---|---|---|---|---|
| **Col15a1** | collagen, type XV, alpha 1 | 3.67 | 1.85 | 8.32E-05 | 4.22E-03 |
| **Cd84** | CD84 antigen | 3.22 | 1.85 | 1.41E-05 | 1.47E-03 |
| **Itgam** | integrin alpha M | 1.41 | 1.84 | 2.57E-05 | 2.06E-03 |
| **A730054J21Rik** | RIKEN cDNA A730054J21 gene | 3.43 | 1.84 | 4.69E-05 | 2.96E-03 |
| **Thbs2** | thrombospondin 2 | 4.07 | 1.84 | 3.97E-05 | 2.68E-03 |
| **C3ar1** | complement component 3a receptor 1 | 3.74 | 1.84 | 7.63E-06 | 1.03E-03 |
| **Il33** | interleukin 33 | 2.02 | 1.84 | 3.17E-05 | 2.29E-03 |
| **Mfsd7b** | major facilitator superfamily domain containing 7B | 2.73 | 1.83 | 2.94E-06 | 6.21E-04 |
| **Scara3** | scavenger receptor class A, member 3 | 5.67 | 1.83 | 6.29E-07 | 2.70E-04 |
| **Pcdh9** | protocadherin 9 | 1.01 | 1.83 | 4.30E-05 | 2.82E-03 |
| **Nckap1l** | NCK associated protein 1 like | 1.16 | 1.82 | 4.76E-06 | 8.07E-04 |
| **Ddr2** | discoidin domain receptor family, member 2 | 1.71 | 1.82 | 4.57E-05 | 2.94E-03 |
| **Clec7a** | C-type lectin domain family 7, member a | 1.77 | 1.82 | 2.35E-05 | 1.94E-03 |
| **Btc** | betacellulin, epidermal growth factor family member | 1.32 | 1.80 | 8.18E-07 | 3.05E-04 |
| **Cfh** | complement component factor h | 1.44 | 1.80 | 2.64E-04 | 9.04E-03 |
| **Il1f6** | interleukin 1 family, member 6 | 7.72 | 1.80 | 1.58E-05 | 1.57E-03 |
| **Cdh11** | cadherin 11 | 4.83 | 1.79 | 1.01E-04 | 4.78E-03 |
| **Aldh1a7** | aldehyde dehydrogenase family 1, subfamily A7 | 1.39 | 1.78 | 1.92E-04 | 7.26E-03 |
| **Ccr2** | chemokine (C-C motif) receptor 2 | 1.26 | 1.77 | 3.91E-05 | 2.66E-03 |
| **Loxl1** | lysyl oxidase-like 1 | 4.68 | 1.76 | 8.52E-06 | 1.08E-03 |
| **Ptprc** | protein tyrosine phosphatase, receptor type, C | 4.27 | 1.76 | 8.20E-06 | 1.05E-03 |
| **Hspb8** | heat shock protein 8 | 3.59 | 1.76 | 5.50E-08 | 8.30E-05 |
| **Muc4** | mucin 4 | 1.21 | 1.76 | 1.58E-05 | 1.57E-03 |
| **Adamts1** | a disintegrin-like and metallopeptidase (reprolysin type) with thrombospondin type 1 motif, 1 | 2.03 | 1.76 | 1.33E-06 | 3.67E-04 |
| **Col6a1** | collagen, type VI, alpha 1 | 5.09 | 1.75 | 5.62E-06 | 8.65E-04 |
| **Svep1** | sushi, von Willebrand factor type A, EGF and pentraxin domain containing 1 | 2.75 | 1.75 | 1.73E-04 | 6.75E-03 |
| **Clu** | clusterin | 3.14 | 1.75 | 3.58E-07 | 2.14E-04 |
| **Tnc** | tenascin C | 4.03 | 1.74 | 1.14E-03 | 2.27E-02 |
| **Ms4a6d** | membrane-spanning 4-domains, subfamily A, member 6D | 3.21 | 1.74 | 1.70E-04 | 6.69E-03 |
| **Colec12** | collectin sub-family member 12 | 1.40 | 1.73 | 7.30E-04 | 1.71E-02 |
| **Slit3** | slit homolog 3 (Drosophila) | 3.68 | 1.73 | 1.98E-05 | 1.75E-03 |
| **Man1c1** | mannosidase, alpha, class 1C, member 1 | 2.15 | 1.73 | 7.85E-05 | 4.11E-03 |
| **Akap12** | A kinase (PRKA) anchor protein (gravin) 12 | 1.87 | 1.72 | 4.34E-04 | 1.24E-02 |
| **Pyhin1** | pyrin and HIN domain family, member 1 | 6.34 | 1.72 | 2.80E-05 | 2.14E-03 |
| **Apobec3** | apolipoprotein B mRNA editing enzyme, catalytic polypeptide 3 | 1.70 | 1.72 | 4.83E-04 | 1.32E-02 |
| **C1ra** | complement component 1, r subcomponent A | 3.14 | 1.70 | 2.13E-05 | 1.82E-03 |
| **Kcnt2** | potassium channel, subfamily T, member 2 | 1.72 | 1.70 | 2.61E-05 | 2.06E-03 |
| **Pde10a** | phosphodiesterase 10A | 1.97 | 1.70 | 5.96E-06 | 8.70E-04 |
| **Ifitm3** | interferon induced transmembrane protein 3 | 2.32 | 1.70 | 8.10E-06 | 1.05E-03 |
| **Ogn** | osteoglycin | 4.61 | 1.70 | 6.96E-04 | 1.66E-02 |
| **Ctsk** | cathepsin K | 4.83 | 1.69 | 2.13E-04 | 7.81E-03 |
| **Steap4** | STEAP family member 4 | 2.46 | 1.69 | 2.77E-05 | 2.14E-03 |
| **Adgrg2** | adhesion G protein-coupled receptor G2 | 2.32 | 1.69 | 5.81E-07 | 2.63E-04 |
| **Tff2** | trefoil factor 2 (spasmolytic protein 1) | 1.73 | 1.69 | 5.33E-05 | 3.21E-03 |
| **Ccdc80** | coiled-coil domain containing 80 | 3.35 | 1.68 | 1.48E-04 | 6.11E-03 |
| **Sytl2** | synaptotagmin-like 2 | 2.42 | 1.68 | 7.97E-05 | 4.13E-03 |
| **Ifi204** | interferon activated gene 204 | 2.23 | 1.68 | 7.40E-05 | 3.97E-03 |
| **Prelp** | proline arginine-rich end leucine-rich repeat | 1.59 | 1.68 | 8.94E-05 | 4.39E-03 |
| **Ch25h** | cholesterol 25-hydroxylase | 2.36 | 1.68 | 1.19E-03 | 2.34E-02 |
| **Fbln1** | fibulin 1 | 7.37 | 1.68 | 4.45E-06 | 7.87E-04 |
| **Slamf7** | SLAM family member 7 | 1.57 | 1.67 | 1.07E-05 | 1.26E-03 |
| **Masp1** | mannan-binding lectin serine peptidase 1 | 1.72 | 1.66 | 4.15E-04 | 1.20E-02 |
| **Cpe** | carboxypeptidase E | 3.17 | 1.66 | 1.20E-05 | 1.33E-03 |
| **Tmem45a** | transmembrane protein 45a | 3.10 | 1.66 | 1.77E-03 | 3.03E-02 |

| | | | | | |
|---|---|---|---|---|---|
| **Trf** | transferrin | 2.08 | 1.65 | 3.67E-06 | 7.04E-04 |
| **Cd14** | CD14 antigen | 5.41 | 1.65 | 7.82E-04 | 1.79E-02 |
| **Klf6** | Kruppel-like factor 6 | 3.86 | 1.65 | 4.79E-06 | 8.07E-04 |
| **Hgf** | hepatocyte growth factor | 1.56 | 1.65 | 4.68E-04 | 1.30E-02 |
| **Cbr3** | carbonyl reductase 3 | 2.45 | 1.65 | 1.72E-04 | 6.70E-03 |
| **Foxa1** | forkhead box A1 | 1.35 | 1.65 | 3.86E-05 | 2.66E-03 |
| **F3** | coagulation factor III | 2.14 | 1.64 | 3.17E-07 | 2.14E-04 |
| **Vim** | vimentin | 3.01 | 1.64 | 3.52E-05 | 2.49E-03 |
| **Myof** | myoferlin | 2.04 | 1.64 | 1.13E-05 | 1.29E-03 |
| **Il34** | interleukin 34 | 1.85 | 1.64 | 3.02E-05 | 2.24E-03 |
| **Matn2** | matrilin 2 | 3.39 | 1.63 | 3.05E-04 | 1.00E-02 |
| **Plac8** | placenta-specific 8 | 3.74 | 1.63 | 1.32E-05 | 1.41E-03 |
| **Rcn1** | reticulocalbin 1 | 2.60 | 1.63 | 5.20E-06 | 8.37E-04 |
| **Laptm5** | lysosomal-associated protein transmembrane 5 | 1.22 | 1.62 | 1.87E-05 | 1.69E-03 |
| **Ccl5** | chemokine (C-C motif) ligand 5 | 8.54 | 1.62 | 1.05E-04 | 4.91E-03 |
| **Adamts2** | a disintegrin-like and metallopeptidase (reprolysin type) with thrombospondin type 1 motif, 2 | 4.27 | 1.62 | 3.31E-05 | 2.37E-03 |
| **Tacstd2** | tumor-associated calcium signal transducer 2 | 3.31 | 1.62 | 7.77E-06 | 1.03E-03 |
| **Havcr2** | hepatitis A virus cellular receptor 2 | 1.11 | 1.62 | 5.89E-06 | 8.65E-04 |
| **Cxcl9** | chemokine (C-X-C motif) ligand 9 | 3.87 | 1.61 | 2.44E-03 | 3.70E-02 |
| **Bgn** | biglycan | 4.75 | 1.61 | 1.01E-04 | 4.78E-03 |
| **Cd48** | CD48 antigen | 3.74 | 1.61 | 8.37E-05 | 4.22E-03 |
| **Birc3** | baculoviral IAP repeat-containing 3 | 1.70 | 1.60 | 1.05E-05 | 1.25E-03 |
| **Cd209a** | CD209a antigen | 1.73 | 1.60 | 2.55E-05 | 2.06E-03 |
| **Slc9a9** | solute carrier family 9 (sodium/hydrogen exchanger), member 9 | 2.13 | 1.60 | 2.27E-06 | 5.17E-04 |
| **Cybb** | cytochrome b-245, beta polypeptide | 1.57 | 1.60 | 7.60E-05 | 4.04E-03 |
| **Dapp1** | dual adaptor for phosphotyrosine and 3-phosphoinositides 1 | 5.41 | 1.59 | 3.93E-06 | 7.38E-04 |
| **Crispld2** | cysteine-rich secretory protein LCCL domain containing 2 | 1.40 | 1.59 | 4.06E-04 | 1.19E-02 |
| **Col14a1** | collagen, type XIV, alpha 1 | 3.46 | 1.59 | 2.28E-05 | 1.90E-03 |
| **Cebpd** | CCAAT/enhancer binding protein (C/EBP), delta | 1.48 | 1.58 | 1.33E-04 | 5.79E-03 |
| **Fbln5** | fibulin 5 | 1.83 | 1.58 | 3.80E-04 | 1.14E-02 |
| **Gpr65** | G-protein coupled receptor 65 | 1.26 | 1.58 | 1.41E-04 | 5.94E-03 |
| **Mrc2** | mannose receptor, C type 2 | 1.89 | 1.58 | 4.08E-05 | 2.72E-03 |
| **Ctsc** | cathepsin C | 2.90 | 1.58 | 2.13E-05 | 1.82E-03 |
| **Art4** | ADP-ribosyltransferase 4 | 5.55 | 1.57 | 6.82E-04 | 1.64E-02 |
| **Upk1a** | uroplakin 1A | 1.21 | 1.57 | 3.43E-03 | 4.57E-02 |
| **Adgrg6** | adhesion G protein-coupled receptor G6 | 1.08 | 1.57 | 3.95E-04 | 1.17E-02 |
| **Nid1** | nidogen 1 | 2.39 | 1.57 | 5.11E-04 | 1.36E-02 |
| **Dock2** | dedicator of cyto-kinesis 2 | 1.27 | 1.56 | 3.73E-05 | 2.60E-03 |
| **Cd33** | CD33 antigen | 1.20 | 1.56 | 7.87E-07 | 3.00E-04 |
| **P2rx7** | purinergic receptor P2X, ligand-gated ion channel, 7 | 1.40 | 1.56 | 2.88E-05 | 2.15E-03 |
| **Serpina1b** | serine (or cysteine) preptidase inhibitor, clade A, member 1B | 2.24 | 1.55 | 3.25E-03 | 4.44E-02 |
| **Rras** | Harvey rat sarcoma oncogene, subgroup R | 2.29 | 1.54 | 1.40E-06 | 3.79E-04 |
| **Emp2** | epithelial membrane protein 2 | 3.38 | 1.54 | 7.66E-06 | 1.03E-03 |
| **Sparcl1** | SPARC-like 1 | 2.28 | 1.54 | 5.25E-05 | 3.18E-03 |
| **Slc34a2** | solute carrier family 34 (sodium phosphate), member 2 | 1.33 | 1.54 | 6.91E-07 | 2.80E-04 |
| **Gpc6** | glypican 6 | 2.04 | 1.54 | 5.25E-05 | 3.18E-03 |
| **Pros1** | protein S (alpha) | 3.35 | 1.54 | 1.71E-04 | 6.70E-03 |
| **Apobec1** | apolipoprotein B mRNA editing enzyme, catalytic polypeptide 1 | 1.43 | 1.53 | 1.18E-04 | 5.28E-03 |
| **Gprc5a** | G protein-coupled receptor, family C, group 5, member A | 1.29 | 1.53 | 2.22E-03 | 3.52E-02 |
| **Mal2** | mal, T cell differentiation protein 2 | 1.07 | 1.53 | 2.35E-03 | 3.63E-02 |
| **Rrad** | Ras-related associated with diabetes | 2.29 | 1.53 | 4.83E-06 | 8.07E-04 |
| **Ly6e** | lymphocyte antigen 6 complex, locus E | 2.45 | 1.53 | 2.85E-05 | 2.15E-03 |
| **Cmtm3** | CKLF-like MARVEL transmembrane domain containing 3 | 3.36 | 1.53 | 1.30E-04 | 5.72E-03 |
| **B4galnt1** | beta-1,4-N-acetyl-galactosaminyl transferase 1 | 1.07 | 1.53 | 2.35E-05 | 1.94E-03 |

| | | | | | |
|---|---|---|---|---|---|
| **Sprr2b** | small proline-rich protein 2B | 1.03 | 1.53 | 1.40E-03 | 2.62E-02 |
| **Hpgds** | hematopoietic prostaglandin D synthase | 1.23 | 1.53 | 1.30E-05 | 1.41E-03 |
| **Arhgap15** | Rho GTPase activating protein 15 | 1.64 | 1.53 | 3.71E-05 | 2.60E-03 |
| **Myo1b** | myosin IB | 1.92 | 1.52 | 2.60E-03 | 3.86E-02 |
| **Rarres2** | retinoic acid receptor responder (tazarotene induced) 2 | 1.99 | 1.52 | 4.57E-05 | 2.94E-03 |
| **Serpina1e** | serine (or cysteine) peptidase inhibitor, clade A, member 1E | 1.89 | 1.52 | 3.22E-03 | 4.41E-02 |
| **Bst2** | bone marrow stromal cell antigen 2 | 2.51 | 1.52 | 2.02E-05 | 1.77E-03 |
| **Epsti1** | epithelial stromal interaction 1 (breast) | 1.47 | 1.52 | 5.89E-05 | 3.41E-03 |
| **Plp2** | proteolipid protein 2 | 2.41 | 1.51 | 8.34E-06 | 1.06E-03 |
| **Timp2** | tissue inhibitor of metalloproteinase 2 | 2.97 | 1.51 | 1.14E-04 | 5.17E-03 |
| **Rpl12** | ribosomal protein L12 | 1.16 | 1.51 | 7.58E-06 | 1.03E-03 |
| **Ms4a4c** | membrane-spanning 4-domains, subfamily A, member 4C | 3.05 | 1.51 | 5.66E-05 | 3.33E-03 |
| **Cxcl16** | chemokine (C-X-C motif) ligand 16 | 4.73 | 1.51 | 2.74E-05 | 2.14E-03 |
| **Gria4** | glutamate receptor, ionotropic, AMPA4 (alpha 4) | 1.57 | 1.51 | 1.36E-04 | 5.88E-03 |
| **Ano1** | anoctamin 1, calcium activated chloride channel | 2.39 | 1.51 | 1.32E-04 | 5.76E-03 |
| **Cldn4** | claudin 4 | 2.13 | 1.50 | 9.00E-05 | 4.41E-03 |
| **Sparc** | secreted acidic cysteine rich glycoprotein | 2.12 | 1.50 | 2.25E-04 | 8.09E-03 |
| **C1qc** | complement component 1, q subcomponent, C chain | 5.37 | 1.50 | 9.59E-05 | 4.66E-03 |
| **Apoe** | apolipoprotein E | 1.43 | 1.50 | 2.06E-05 | 1.79E-03 |
| **Upk2** | uroplakin 2 | 1.17 | 1.50 | 2.42E-04 | 8.54E-03 |
| **Dpysl3** | dihydropyrimidinase-like 3 | 5.82 | 1.49 | 1.22E-05 | 1.34E-03 |
| **F2r** | coagulation factor II (thrombin) receptor | 1.43 | 1.49 | 2.85E-03 | 4.08E-02 |
| **Tmsb4x** | thymosin, beta 4, X chromosome | 1.96 | 1.49 | 4.00E-05 | 2.69E-03 |
| **Tyrobp** | TYRO protein tyrosine kinase binding protein | 4.01 | 1.49 | 4.53E-04 | 1.28E-02 |
| **Cxcl10** | chemokine (C-X-C motif) ligand 10 | 5.08 | 1.48 | 1.78E-07 | 1.53E-04 |
| **Cnn2** | calponin 2 | 2.07 | 1.48 | 1.14E-04 | 5.15E-03 |
| **Axl** | AXL receptor tyrosine kinase | 2.92 | 1.48 | 1.09E-04 | 5.03E-03 |
| **Nipal1** | NIPA-like domain containing 1 | 1.31 | 1.47 | 2.58E-03 | 3.84E-02 |
| **Gdpd1** | glycerophosphodiester phosphodiesterase domain containing 1 | 1.37 | 1.47 | 8.35E-05 | 4.22E-03 |
| **Gstp1** | glutathione S-transferase, pi 1 | 1.05 | 1.47 | 3.56E-06 | 6.95E-04 |
| **Cbr2** | carbonyl reductase 2 | 2.27 | 1.47 | 5.68E-06 | 8.65E-04 |
| **Bdkrb2** | bradykinin receptor, beta 2 | 3.67 | 1.47 | 3.41E-07 | 2.14E-04 |
| **Lgals9** | lectin, galactose binding, soluble 9 | 2.68 | 1.46 | 3.05E-04 | 1.00E-02 |
| **Fcgr1** | Fc receptor, IgG, high affinity I | 6.86 | 1.46 | 1.43E-03 | 2.65E-02 |
| **Sla** | src-like adaptor | 1.22 | 1.46 | 3.35E-04 | 1.06E-02 |
| **Rab31** | RAB31, member RAS oncogene family | 2.63 | 1.46 | 4.73E-05 | 2.97E-03 |
| **Gm4956** | predicted gene 4956 | 1.31 | 1.46 | 1.59E-04 | 6.40E-03 |
| **Cxcl12** | chemokine (C-X-C motif) ligand 12 | 1.69 | 1.46 | 3.19E-05 | 2.30E-03 |
| **C1qa** | complement component 1, q subcomponent, alpha polypeptide | 4.82 | 1.46 | 9.85E-05 | 4.73E-03 |
| **Mgl2** | macrophage galactose N-acetyl-galactosamine specific lectin 2 | 1.58 | 1.46 | 1.09E-04 | 5.03E-03 |
| **Spaca7** | sperm acrosome associated 7 | 2.69 | 1.45 | 4.14E-05 | 2.74E-03 |
| **Aqp3** | aquaporin 3 | 1.38 | 1.45 | 6.73E-06 | 9.38E-04 |
| **Casp1** | caspase 1 | 3.29 | 1.45 | 5.81E-05 | 3.38E-03 |
| **Serpine2** | serine (or cysteine) peptidase inhibitor, clade E, member 2 | 3.76 | 1.45 | 1.27E-03 | 2.45E-02 |
| **P2ry6** | pyrimidinergic receptor P2Y, G-protein coupled, 6 | 3.90 | 1.45 | 2.57E-04 | 8.86E-03 |
| **Fat4** | FAT tumor suppressor homolog 4 (Drosophila) | 2.32 | 1.45 | 4.84E-04 | 1.32E-02 |
| **Col4a2** | collagen, type IV, alpha 2 | 2.09 | 1.45 | 6.42E-05 | 3.61E-03 |
| **Pde3a** | phosphodiesterase 3A, cGMP inhibited | 1.61 | 1.45 | 4.05E-04 | 1.19E-02 |
| **Icam1** | intercellular adhesion molecule 1 | 3.48 | 1.45 | 6.06E-06 | 8.74E-04 |
| **Apbb1ip** | amyloid beta (A4) precursor protein-binding, family B, member 1 interacting protein | 2.83 | 1.45 | 6.25E-04 | 1.55E-02 |
| **Cd52** | CD52 antigen | 4.23 | 1.45 | 1.69E-04 | 6.66E-03 |
| **Ccdc109b** | coiled-coil domain containing 109B | 3.30 | 1.45 | 2.76E-05 | 2.14E-03 |
| **Fam46a** | family with sequence similarity 46, member A | 1.52 | 1.45 | 9.56E-05 | 4.66E-03 |
| **Fyb** | FYN binding protein | 4.56 | 1.45 | 4.65E-05 | 2.95E-03 |

| | | | | | |
|---|---|---|---|---|---|
| **Col4a1** | collagen, type IV, alpha 1 | 1.84 | 1.45 | 2.66E-05 | 2.09E-03 |
| **Ccl6** | chemokine (C-C motif) ligand 6 | 4.27 | 1.45 | 3.61E-05 | 2.54E-03 |
| **Ifngr1** | interferon gamma receptor 1 | 2.08 | 1.45 | 2.52E-06 | 5.53E-04 |
| **Stab1** | stabilin 1 | 2.36 | 1.44 | 9.63E-05 | 4.67E-03 |
| **Lhfp** | lipoma HMGIC fusion partner | 3.17 | 1.44 | 3.84E-03 | 4.90E-02 |
| **Spink12** | serine peptidase inhibitor, Kazal type 12 | 1.84 | 1.44 | 2.67E-03 | 3.93E-02 |
| **Rac2** | RAS-related C3 botulinum substrate 2 | 5.69 | 1.44 | 4.22E-05 | 2.79E-03 |
| **Gxylt2** | glucoside xylosyltransferase 2 | 3.05 | 1.44 | 1.29E-06 | 3.67E-04 |
| **Hmox1** | heme oxygenase 1 | 1.76 | 1.44 | 6.94E-05 | 3.86E-03 |
| **Frzb** | frizzled-related protein | 1.35 | 1.44 | 2.80E-03 | 4.04E-02 |
| **Apod** | apolipoprotein D | 1.64 | 1.44 | 9.62E-04 | 2.05E-02 |
| **Neurl3** | neuralized E3 ubiquitin protein ligase 3 | 1.97 | 1.43 | 5.29E-06 | 8.39E-04 |
| **Epha3** | Eph receptor A3 | 1.58 | 1.43 | 7.05E-05 | 3.86E-03 |
| **Lama2** | laminin, alpha 2 | 3.81 | 1.43 | 2.75E-03 | 4.00E-02 |
| **Themis2** | thymocyte selection associated family member 2 | 2.65 | 1.43 | 1.86E-04 | 7.09E-03 |
| **Inpp5d** | inositol polyphosphate-5-phosphatase D | 4.88 | 1.43 | 7.52E-04 | 1.74E-02 |
| **Itgb2** | integrin beta 2 | 4.90 | 1.43 | 4.95E-04 | 1.34E-02 |
| **Il7r** | interleukin 7 receptor | 3.42 | 1.43 | 4.87E-04 | 1.32E-02 |
| **Plek** | pleckstrin | 2.66 | 1.43 | 1.90E-04 | 7.21E-03 |
| **Amica1** | adhesion molecule, interacts with CXADR antigen 1 | 1.92 | 1.43 | 3.34E-03 | 4.51E-02 |
| **Fam105a** | family with sequence similarity 105, member A | 3.32 | 1.42 | 2.33E-03 | 3.63E-02 |
| **Cacna2d1** | calcium channel, voltage-dependent, alpha2/delta subunit 1 | 1.35 | 1.42 | 1.27E-04 | 5.64E-03 |
| **Rerg** | RAS-like, estrogen-regulated, growth-inhibitor | 1.68 | 1.42 | 2.39E-03 | 3.67E-02 |
| **Anxa1** | annexin A1 | 3.59 | 1.42 | 2.96E-03 | 4.17E-02 |
| **Cygb** | cytoglobin | 5.50 | 1.42 | 1.71E-05 | 1.62E-03 |
| **Fstl1** | follistatin-like 1 | 2.44 | 1.42 | 2.40E-03 | 3.68E-02 |
| **Golm1** | golgi membrane protein 1 | 1.32 | 1.42 | 8.08E-06 | 1.05E-03 |
| **Col6a2** | collagen, type VI, alpha 2 | 3.89 | 1.42 | 2.55E-05 | 2.06E-03 |
| **Nav3** | neuron navigator 3 | 1.64 | 1.42 | 2.66E-03 | 3.92E-02 |
| **Penk** | preproenkephalin | 2.41 | 1.42 | 6.12E-04 | 1.53E-02 |
| **Ak5** | adenylate kinase 5 | 1.98 | 1.42 | 4.76E-05 | 2.98E-03 |
| **Lamc2** | laminin, gamma 2 | 1.55 | 1.42 | 1.01E-05 | 1.21E-03 |
| **Adamtsl1** | ADAMTS-like 1 | 1.81 | 1.42 | 4.54E-05 | 2.94E-03 |
| **Clip3** | CAP-GLY domain containing linker protein 3 | 2.27 | 1.41 | 5.42E-05 | 3.24E-03 |
| **Unc93b1** | unc-93 homolog B1 (C. elegans) | 1.83 | 1.41 | 2.05E-05 | 1.78E-03 |
| **Mxra8** | matrix-remodelling associated 8 | 3.63 | 1.41 | 3.26E-04 | 1.04E-02 |
| **Adamts5** | a disintegrin-like and metallopeptidase (reprolysin type) with thrombospondin type 1 motif, 5 (aggrecanase-2) | 2.45 | 1.41 | 2.66E-04 | 9.05E-03 |
| **Lrp1** | low density lipoprotein receptor-related protein 1 | 4.40 | 1.41 | 1.02E-03 | 2.13E-02 |
| **Plbd1** | phospholipase B domain containing 1 | 1.78 | 1.41 | 5.05E-04 | 1.35E-02 |
| **Pik3ap1** | phosphoinositide-3-kinase adaptor protein 1 | 2.94 | 1.41 | 1.06E-04 | 4.92E-03 |
| **Lgals3bp** | lectin, galactoside-binding, soluble, 3 binding protein | 2.21 | 1.41 | 2.23E-04 | 8.06E-03 |
| **Ifitm1** | interferon induced transmembrane protein 1 | 2.04 | 1.41 | 5.76E-04 | 1.45E-02 |
| **Adamts16** | a disintegrin-like and metallopeptidase (reprolysin type) with thrombospondin type 1 motif, 16 | 1.10 | 1.41 | 1.81E-05 | 1.68E-03 |
| **Fcrls** | Fc receptor-like S, scavenger receptor | 7.63 | 1.41 | 2.23E-03 | 3.54E-02 |
| **Cxcl1** | chemokine (C-X-C motif) ligand 1 | 10.29 | 1.40 | 2.83E-03 | 4.06E-02 |
| **Gabra3** | gamma-aminobutyric acid (GABA) A receptor, subunit alpha 3 | 2.00 | 1.40 | 6.57E-04 | 1.59E-02 |
| **Fblim1** | filamin binding LIM protein 1 | 2.32 | 1.40 | 4.42E-04 | 1.25E-02 |
| **Ucp2** | uncoupling protein 2 (mitochondrial, proton carrier) | 1.57 | 1.40 | 1.30E-03 | 2.48E-02 |
| **Sulf1** | sulfatase 1 | 2.42 | 1.40 | 2.06E-04 | 7.63E-03 |
| **Olfml1** | olfactomedin-like 1 | 1.79 | 1.40 | 6.21E-06 | 8.88E-04 |
| **Nipal2** | NIPA-like domain containing 2 | 2.25 | 1.40 | 7.95E-07 | 3.00E-04 |
| **Aebp1** | AE binding protein 1 | 3.25 | 1.40 | 1.23E-03 | 2.39E-02 |
| **Arpc1b** | actin related protein 2/3 complex, subunit 1B | 2.91 | 1.40 | 6.82E-06 | 9.46E-04 |
| **Adcy7** | adenylate cyclase 7 | 6.55 | 1.40 | 7.23E-05 | 3.91E-03 |

| | | | | | |
|---|---|---|---|---|---|
| **Tubb5** | tubulin, beta 5 class I | 2.97 | 1.39 | 3.43E-04 | 1.08E-02 |
| **Dock11** | dedicator of cytokinesis 11 | 1.97 | 1.39 | 1.78E-05 | 1.66E-03 |
| **Ptgfrn** | prostaglandin F2 receptor negative regulator | 1.64 | 1.39 | 1.38E-04 | 5.92E-03 |
| **Siglec1** | sialic acid binding Ig-like lectin 1, sialoadhesin | 1.08 | 1.39 | 1.36E-04 | 5.88E-03 |
| **Scarf2** | scavenger receptor class F, member 2 | 1.63 | 1.39 | 6.02E-05 | 3.45E-03 |
| **Pdgfrb** | platelet derived growth factor receptor, beta polypeptide | 2.56 | 1.39 | 1.97E-04 | 7.39E-03 |
| **Ncam1** | neural cell adhesion molecule 1 | 2.26 | 1.39 | 8.51E-05 | 4.27E-03 |
| **Pi15** | peptidase inhibitor 15 | 2.01 | 1.38 | 3.59E-04 | 1.11E-02 |
| **Cd24a** | CD24a antigen | 1.50 | 1.38 | 7.26E-04 | 1.71E-02 |
| **Adgra1** | adhesion G protein-coupled receptor A1 | 1.99 | 1.38 | 2.00E-03 | 3.29E-02 |
| **Tuba1c** | tubulin, alpha 1C | 1.66 | 1.38 | 6.97E-05 | 3.86E-03 |
| **Slfn2** | schlafen 2 | 1.75 | 1.38 | 2.39E-03 | 3.67E-02 |
| **Ednra** | endothelin receptor type A | 2.33 | 1.38 | 3.61E-03 | 4.73E-02 |
| **Ccl12** | chemokine (C-C motif) ligand 12 | 7.93 | 1.38 | 1.24E-05 | 1.35E-03 |
| **Sox9** | SRY (sex determining region Y)-box 9 | 1.05 | 1.38 | 1.10E-06 | 3.54E-04 |
| **Ubtd2** | ubiquitin domain containing 2 | 1.87 | 1.37 | 2.38E-03 | 3.66E-02 |
| **Cyth4** | cytohesin 4 | 1.52 | 1.37 | 2.90E-04 | 9.66E-03 |
| **Kdelr3** | KDEL (Lys-Asp-Glu-Leu) endoplasmic reticulum protein retention receptor 3 | 2.39 | 1.37 | 1.32E-05 | 1.41E-03 |
| **Cldn16** | claudin 16 | 1.30 | 1.37 | 4.32E-06 | 7.78E-04 |
| **Runx1** | runt related transcription factor 1 | 1.05 | 1.37 | 1.45E-05 | 1.49E-03 |
| **Lpo** | lactoperoxidase | 1.72 | 1.37 | 8.02E-05 | 4.14E-03 |
| **Col18a1** | collagen, type XVIII, alpha 1 | 2.35 | 1.37 | 1.41E-04 | 5.94E-03 |
| **Pof1b** | premature ovarian failure 1B | 1.89 | 1.37 | 8.30E-04 | 1.85E-02 |
| **Sfrp4** | secreted frizzled-related protein 4 | 2.88 | 1.37 | 3.26E-04 | 1.04E-02 |
| **Sult5a1** | sulfotransferase family 5A, member 1 | 1.33 | 1.37 | 5.70E-04 | 1.45E-02 |
| **Abca8a** | ATP-binding cassette, sub-family A (ABC1), member 8a | 1.42 | 1.36 | 2.18E-04 | 7.94E-03 |
| **Arhgdib** | Rho, GDP dissociation inhibitor (GDI) beta | 2.37 | 1.36 | 2.00E-04 | 7.47E-03 |
| **Tmem98** | transmembrane protein 98 | 2.58 | 1.36 | 3.49E-04 | 1.10E-02 |
| **Lxn** | latexin | 3.20 | 1.36 | 1.47E-05 | 1.50E-03 |
| **Gulp1** | GULP, engulfment adaptor PTB domain containing 1 | 1.88 | 1.36 | 5.11E-05 | 3.13E-03 |
| **Chrnb1** | cholinergic receptor, nicotinic, beta polypeptide 1 (muscle) | 2.77 | 1.36 | 3.97E-04 | 1.17E-02 |
| **Pik3cg** | phosphoinositide-3-kinase, catalytic, gamma polypeptide | 1.15 | 1.36 | 3.89E-05 | 2.66E-03 |
| **P2ry13** | purinergic receptor P2Y, G-protein coupled 13 | 1.11 | 1.36 | 6.69E-04 | 1.61E-02 |
| **B2m** | beta-2 microglobulin | 1.94 | 1.36 | 2.12E-04 | 7.79E-03 |
| **Mgst1** | microsomal glutathione S-transferase 1 | 1.21 | 1.36 | 5.34E-06 | 8.39E-04 |
| **Spi1** | spleen focus forming virus (SFFV) proviral integration oncogene | 1.14 | 1.36 | 1.56E-05 | 1.56E-03 |
| **Sdk1** | sidekick homolog 1 (chicken) | 2.08 | 1.36 | 1.59E-05 | 1.57E-03 |
| **Ccr1** | chemokine (C-C motif) receptor 1 | 1.66 | 1.36 | 2.36E-03 | 3.65E-02 |
| **Rasa3** | RAS p21 protein activator 3 | 2.78 | 1.35 | 3.85E-05 | 2.66E-03 |
| **Fkbp7** | FK506 binding protein 7 | 2.63 | 1.35 | 1.30E-04 | 5.72E-03 |
| **Ccdc3** | coiled-coil domain containing 3 | 1.08 | 1.35 | 1.99E-04 | 7.44E-03 |
| **Fcgr3** | Fc receptor, IgG, low affinity III | 3.01 | 1.35 | 3.93E-03 | 4.95E-02 |
| **Dab1** | disabled 1 | 2.30 | 1.35 | 1.96E-04 | 7.35E-03 |
| **Zfp36l1** | zinc finger protein 36, C3H type-like 1 | 1.92 | 1.35 | 1.14E-05 | 1.30E-03 |
| **Stk17b** | serine/threonine kinase 17b (apoptosis-inducing) | 1.93 | 1.35 | 1.40E-04 | 5.94E-03 |
| **Cep170** | centrosomal protein 170 | 1.33 | 1.35 | 1.33E-04 | 5.79E-03 |
| **Tagln** | transgelin | 3.40 | 1.35 | 6.57E-04 | 1.59E-02 |
| **Srpx** | sushi-repeat-containing protein | 3.18 | 1.35 | 6.43E-04 | 1.57E-02 |
| **Gbp2** | guanylate binding protein 2 | 1.99 | 1.35 | 1.25E-03 | 2.42E-02 |
| **Bmp1** | bone morphogenetic protein 1 | 2.15 | 1.35 | 5.99E-05 | 3.45E-03 |
| **Myo1f** | myosin IF | 3.63 | 1.35 | 2.77E-04 | 9.31E-03 |
| **Dtx4** | deltex 4 homolog (Drosophila) | 1.10 | 1.35 | 3.79E-05 | 2.63E-03 |
| **Il1rn** | interleukin 1 receptor antagonist | 1.93 | 1.35 | 1.83E-05 | 1.69E-03 |
| **Plcxd3** | phosphatidylinositol-specific phospholipase C, X domain containing 3 | 1.44 | 1.35 | 4.41E-04 | 1.25E-02 |

| | | | | | |
|---|---|---|---|---|---|
| **Cpne8** | copine VIII | 2.69 | 1.34 | 3.05E-03 | 4.24E-02 |
| **Vav1** | vav 1 oncogene | 4.39 | 1.34 | 3.28E-04 | 1.05E-02 |
| **Bcat1** | branched chain aminotransferase 1, cytosolic | 1.20 | 1.34 | 2.64E-03 | 3.90E-02 |
| **Peak1** | pseudopodium-enriched atypical kinase 1 | 1.37 | 1.34 | 2.89E-03 | 4.11E-02 |
| **Rela** | v-rel reticuloendotheliosis viral oncogene homolog A (avian) | 3.08 | 1.34 | 1.90E-05 | 1.70E-03 |
| **Gpr34** | G protein-coupled receptor 34 | 1.08 | 1.34 | 7.91E-05 | 4.13E-03 |
| **Krt5** | keratin 5 | 2.39 | 1.34 | 2.57E-04 | 8.86E-03 |
| **Il10ra** | interleukin 10 receptor, alpha | 1.39 | 1.34 | 9.63E-06 | 1.18E-03 |
| **Klrd1** | killer cell lectin-like receptor, subfamily D, member 1 | 2.65 | 1.33 | 4.04E-05 | 2.70E-03 |
| **Akr1b7** | aldo-keto reductase family 1, member B7 | 1.49 | 1.33 | 1.39E-03 | 2.61E-02 |
| **H6pd** | hexose-6-phosphate dehydrogenase (glucose 1-dehydrogenase) | 1.40 | 1.33 | 3.08E-04 | 1.01E-02 |
| **Wfdc2** | WAP four-disulfide core domain 2 | 1.56 | 1.33 | 7.61E-06 | 1.03E-03 |
| **Fcgr4** | Fc receptor, IgG, low affinity IV | 2.05 | 1.33 | 1.34E-03 | 2.54E-02 |
| **Naip5** | NLR family, apoptosis inhibitory protein 5 | 1.21 | 1.33 | 6.23E-05 | 3.55E-03 |
| **Lgals1** | lectin, galactose binding, soluble 1 | 2.40 | 1.33 | 2.44E-04 | 8.55E-03 |
| **Tpm1** | tropomyosin 1, alpha | 5.44 | 1.33 | 5.94E-05 | 3.42E-03 |
| **Prr15** | proline rich 15 | 1.73 | 1.33 | 5.94E-04 | 1.49E-02 |
| **Pmaip1** | phorbol-12-myristate-13-acetate-induced protein 1 | 2.47 | 1.32 | 5.60E-05 | 3.31E-03 |
| **Sepn1** | selenoprotein N, 1 | 1.73 | 1.32 | 7.39E-04 | 1.72E-02 |
| **Cd109** | CD109 antigen | 1.45 | 1.32 | 2.97E-04 | 9.84E-03 |
| **Csrp1** | cysteine and glycine-rich protein 1 | 2.99 | 1.32 | 1.12E-04 | 5.10E-03 |
| **Samhd1** | SAM domain and HD domain, 1 | 1.56 | 1.32 | 1.70E-04 | 6.68E-03 |
| **Elf3** | E74-like factor 3 | 2.01 | 1.32 | 1.08E-04 | 4.98E-03 |
| **Itga4** | integrin alpha 4 | 1.44 | 1.32 | 1.23E-05 | 1.35E-03 |
| **Cyp2f2** | cytochrome P450, family 2, subfamily f, polypeptide 2 | 1.14 | 1.32 | 1.49E-03 | 2.70E-02 |
| **Sorbs2** | sorbin and SH3 domain containing 2 | 2.09 | 1.32 | 6.92E-04 | 1.66E-02 |
| **Mxra7** | matrix-remodelling associated 7 | 1.79 | 1.32 | 3.35E-04 | 1.06E-02 |
| **Cd86** | CD86 antigen | 3.11 | 1.32 | 1.69E-04 | 6.66E-03 |
| **Gpm6b** | glycoprotein m6b | 1.96 | 1.32 | 1.80E-03 | 3.06E-02 |
| **Dap** | death-associated protein | 1.55 | 1.32 | 6.00E-06 | 8.71E-04 |
| **Ptafr** | platelet-activating factor receptor | 4.21 | 1.32 | 8.86E-04 | 1.94E-02 |
| **Gpc3** | glypican 3 | 1.75 | 1.32 | 2.56E-03 | 3.81E-02 |
| **Clec12a** | C-type lectin domain family 12, member a | 1.33 | 1.32 | 6.48E-04 | 1.58E-02 |
| **Cldn3** | claudin 3 | 1.75 | 1.32 | 8.15E-05 | 4.19E-03 |
| **Cxcl14** | chemokine (C-X-C motif) ligand 14 | 3.97 | 1.31 | 8.58E-04 | 1.89E-02 |
| **Tmem47** | transmembrane protein 47 | 1.17 | 1.31 | 6.94E-04 | 1.66E-02 |
| **Vtcn1** | V-set domain containing T cell activation inhibitor 1 | 1.34 | 1.31 | 4.32E-04 | 1.24E-02 |
| **Kif1a** | kinesin family member 1A | 1.34 | 1.31 | 8.60E-04 | 1.89E-02 |
| **Pip4k2a** | phosphatidylinositol-5-phosphate 4-kinase, type II, alpha | 1.06 | 1.31 | 1.22E-03 | 2.38E-02 |
| **Ncf2** | neutrophil cytosolic factor 2 | 2.49 | 1.31 | 5.78E-06 | 8.65E-04 |
| **Rnasel** | ribonuclease L (2', 5'-oligoisoadenylate synthetase-dependent) | 1.44 | 1.31 | 1.15E-04 | 5.18E-03 |
| **Dact1** | dapper homolog 1, antagonist of beta-catenin (xenopus) | 3.39 | 1.31 | 5.76E-04 | 1.45E-02 |
| **Myl9** | myosin, light polypeptide 9, regulatory | 1.78 | 1.31 | 5.31E-05 | 3.21E-03 |
| **Ms4a4b** | membrane-spanning 4-domains, subfamily A, member 4B | 5.84 | 1.31 | 3.32E-05 | 2.37E-03 |
| **C4a** | complement component 4A (Rodgers blood group) | 4.47 | 1.31 | 8.31E-04 | 1.85E-02 |
| **Wdfy4** | WD repeat and FYVE domain containing 4 | 3.15 | 1.31 | 3.12E-04 | 1.02E-02 |
| **AW112010** | expressed sequence AW112010 | 2.06 | 1.31 | 1.73E-03 | 2.98E-02 |
| **Lbh** | limb-bud and heart | 3.19 | 1.31 | 1.86E-05 | 1.69E-03 |
| **Ubd** | ubiquitin D | 12.67 | 1.31 | 1.75E-05 | 1.64E-03 |
| **Tenm3** | teneurin transmembrane protein 3 | 3.49 | 1.31 | 1.69E-04 | 6.66E-03 |
| **Gm21188** | predicted gene, 21188 | 1.03 | 1.30 | 5.20E-04 | 1.37E-02 |
| **Sytl5** | synaptotagmin-like 5 | 1.06 | 1.30 | 5.16E-05 | 3.15E-03 |
| **Emp3** | epithelial membrane protein 3 | 1.36 | 1.30 | 8.60E-04 | 1.89E-02 |
| **Csf2rb** | colony stimulating factor 2 receptor, beta, low-affinity (granulocyte-macrophage) | 1.65 | 1.30 | 9.01E-05 | 4.41E-03 |

| | | | | | |
|---|---|---|---|---|---|
| **Ogfrl1** | opioid growth factor receptor-like 1 | 1.07 | 1.30 | 7.18E-05 | 3.90E-03 |
| **Itih5** | inter-alpha (globulin) inhibitor H5 | 1.76 | 1.30 | 4.39E-04 | 1.25E-02 |
| **Efemp2** | epidermal growth factor-containing fibulin-like extracellular matrix protein 2 | 4.12 | 1.30 | 4.25E-04 | 1.23E-02 |
| **1810022K09Rik** | RIKEN cDNA 1810022K09 gene | 1.13 | 1.30 | 3.42E-03 | 4.57E-02 |
| **Camk1d** | calcium/calmodulin-dependent protein kinase ID | 1.01 | 1.30 | 1.18E-04 | 5.28E-03 |
| **Rhou** | ras homolog gene family, member U | 3.41 | 1.30 | 2.89E-04 | 9.62E-03 |
| **Rcn3** | reticulocalbin 3, EF-hand calcium binding domain | 3.25 | 1.30 | 3.24E-04 | 1.04E-02 |
| **Stat3** | signal transducer and activator of transcription 3 | 2.28 | 1.30 | 1.07E-05 | 1.26E-03 |
| **Oas2** | 2'-5' oligoadenylate synthetase 2 | 3.27 | 1.30 | 7.90E-04 | 1.79E-02 |
| **Pea15a** | phosphoprotein enriched in astrocytes 15A | 2.61 | 1.30 | 4.10E-05 | 2.72E-03 |
| **Cldn1** | claudin 1 | 2.08 | 1.29 | 2.03E-03 | 3.32E-02 |
| **Cx3cl1** | chemokine (C-X3-C motif) ligand 1 | 3.55 | 1.29 | 7.06E-05 | 3.86E-03 |
| **Gstm1** | glutathione S-transferase, mu 1 | 1.06 | 1.29 | 1.08E-04 | 4.99E-03 |
| **Luzp2** | leucine zipper protein 2 | 1.25 | 1.29 | 3.49E-03 | 4.60E-02 |
| **Ccl2** | chemokine (C-C motif) ligand 2 | 2.40 | 1.29 | 1.65E-04 | 6.58E-03 |
| **Adamts17** | a disintegrin-like and metallopeptidase (reprolysin type) with thrombospondin type 1 motif, 17 | 1.66 | 1.29 | 2.07E-03 | 3.37E-02 |
| **Pdpn** | podoplanin | 1.94 | 1.29 | 9.08E-04 | 1.97E-02 |
| **Timp1** | tissue inhibitor of metalloproteinase 1 | 21.84 | 1.29 | 2.41E-03 | 3.68E-02 |
| **Tpm4** | tropomyosin 4 | 1.86 | 1.29 | 3.61E-04 | 1.11E-02 |
| **Pdcd4** | programmed cell death 4 | 1.42 | 1.29 | 4.61E-04 | 1.29E-02 |
| **Cxcr4** | chemokine (C-X-C motif) receptor 4 | 3.46 | 1.29 | 6.45E-04 | 1.58E-02 |
| **Tgfbr2** | transforming growth factor, beta receptor II | 1.15 | 1.29 | 1.14E-04 | 5.15E-03 |
| **Slc7a2** | solute carrier family 7 (cationic amino acid transporter, y+ system), member 2 | 1.17 | 1.29 | 8.72E-05 | 4.32E-03 |
| **Cysltr1** | cysteinyl leukotriene receptor 1 | 2.58 | 1.29 | 6.89E-04 | 1.65E-02 |
| **Trpv6** | transient receptor potential cation channel, subfamily V, member 6 | 2.02 | 1.28 | 1.41E-03 | 2.62E-02 |
| **Tnfrsf12a** | tumor necrosis factor receptor superfamily, member 12a | 3.40 | 1.28 | 3.20E-03 | 4.39E-02 |
| **Smim3** | small integral membrane protein 3 | 1.48 | 1.28 | 7.42E-04 | 1.73E-02 |
| **Pgm5** | phosphoglucomutase 5 | 1.44 | 1.28 | 3.59E-03 | 4.71E-02 |
| **Sh3bgrl3** | SH3 domain binding glutamic acid-rich protein-like 3 | 2.27 | 1.28 | 3.68E-04 | 1.12E-02 |
| **Stim2** | stromal interaction molecule 2 | 1.55 | 1.28 | 2.64E-04 | 9.02E-03 |
| **B3galt1** | UDP-Gal:betaGlcNAc beta 1,3-galactosyltransferase, polypeptide 1 | 8.45 | 1.28 | 4.45E-04 | 1.26E-02 |
| **Cers3** | ceramide synthase 3 | 1.16 | 1.28 | 1.93E-03 | 3.23E-02 |
| **Pcdhb14** | protocadherin beta 14 | 1.15 | 1.28 | 4.78E-04 | 1.31E-02 |
| **Sept8** | septin 8 | 1.67 | 1.28 | 1.01E-03 | 2.13E-02 |
| **Mmp11** | matrix metallopeptidase 11 | 2.67 | 1.28 | 2.46E-03 | 3.73E-02 |
| **Rftn1** | raftlin lipid raft linker 1 | 1.13 | 1.28 | 4.75E-04 | 1.31E-02 |
| **Cd300ld** | CD300 molecule-like family member d | 3.83 | 1.28 | 9.86E-04 | 2.08E-02 |
| **Parp14** | poly (ADP-ribose) polymerase family, member 14 | 5.19 | 1.28 | 3.71E-04 | 1.13E-02 |
| **Cdh6** | cadherin 6 | 1.25 | 1.28 | 1.40E-04 | 5.94E-03 |
| **Pira6** | paired-Ig-like receptor A6 | 1.30 | 1.28 | 4.27E-04 | 1.23E-02 |
| **Robo1** | roundabout homolog 1 (Drosophila) | 1.95 | 1.27 | 7.63E-04 | 1.76E-02 |
| **Rpl35** | ribosomal protein L35 | 2.68 | 1.27 | 3.16E-05 | 2.29E-03 |
| **Ttc39b** | tetratricopeptide repeat domain 39B | 1.28 | 1.27 | 2.99E-05 | 2.23E-03 |
| **Tmeff1** | transmembrane protein with EGF-like and two follistatin-like domains 1 | 1.12 | 1.27 | 1.55E-03 | 2.79E-02 |
| **Rnf150** | ring finger protein 150 | 1.31 | 1.27 | 2.72E-04 | 9.21E-03 |
| **Bcl3** | B cell leukemia/lymphoma 3 | 1.58 | 1.27 | 1.49E-04 | 6.13E-03 |
| **Rpl39** | ribosomal protein L39 | 1.15 | 1.27 | 1.48E-04 | 6.11E-03 |
| **Map3k8** | mitogen-activated protein kinase kinase kinase 8 | 1.50 | 1.27 | 5.48E-04 | 1.42E-02 |
| **Nrg1** | neuregulin 1 | 2.21 | 1.27 | 2.94E-03 | 4.15E-02 |
| **Zmynd15** | zinc finger, MYND-type containing 15 | 2.68 | 1.27 | 8.62E-05 | 4.30E-03 |
| **Lrrn1** | leucine rich repeat protein 1, neuronal | 1.80 | 1.26 | 2.73E-03 | 3.99E-02 |
| **Tlr2** | toll-like receptor 2 | 3.79 | 1.26 | 9.74E-05 | 4.70E-03 |
| **Ifitm2** | interferon induced transmembrane protein 2 | 1.62 | 1.26 | 3.13E-03 | 4.32E-02 |
| **Krt8** | keratin 8 | 2.44 | 1.26 | 9.25E-06 | 1.15E-03 |

| | | | | | |
|---|---|---|---|---|---|
| **Mpp1** | membrane protein, palmitoylated | 1.11 | 1.26 | 1.15E-03 | 2.29E-02 |
| **Plekhh2** | pleckstrin homology domain containing, family H (with MyTH4 domain) member 2 | 1.16 | 1.26 | 1.11E-03 | 2.24E-02 |
| **Cblb** | Casitas B-lineage lymphoma b | 1.40 | 1.26 | 7.32E-04 | 1.71E-02 |
| **Smpdl3b** | sphingomyelin phosphodiesterase, acid-like 3B | 4.73 | 1.26 | 2.07E-03 | 3.37E-02 |
| **Gsn** | gelsolin | 2.56 | 1.26 | 2.53E-03 | 3.79E-02 |
| **Kirrel** | kin of IRRE like (Drosophila) | 1.25 | 1.26 | 2.69E-03 | 3.95E-02 |
| **Cxcl17** | chemokine (C-X-C motif) ligand 17 | 1.39 | 1.26 | 1.67E-04 | 6.64E-03 |
| **Phf11b** | PHD finger protein 11B | 1.98 | 1.26 | 9.03E-04 | 1.96E-02 |
| **Acat3** | acetyl-Coenzyme A acetyltransferase 3 | 1.35 | 1.26 | 6.46E-04 | 1.58E-02 |
| **Cd3e** | CD3 antigen, epsilon polypeptide | 1.11 | 1.26 | 1.10E-03 | 2.22E-02 |
| **Fam49b** | family with sequence similarity 49, member B | 1.48 | 1.26 | 1.46E-03 | 2.67E-02 |
| **Hip1** | huntingtin interacting protein 1 | 4.03 | 1.26 | 2.52E-03 | 3.79E-02 |
| **Gm2115** | predicted gene 2115 | 1.49 | 1.26 | 1.54E-04 | 6.23E-03 |
| **Clca3b** | chloride channel calcium activated 3B | 3.78 | 1.26 | 2.02E-04 | 7.50E-03 |
| **Gnai2** | guanine nucleotide binding protein (G protein), alpha inhibiting 2 | 1.77 | 1.26 | 1.93E-05 | 1.72E-03 |
| **Plekho1** | pleckstrin homology domain containing, family O member 1 | 2.27 | 1.26 | 4.56E-05 | 2.94E-03 |
| **Sprr2g** | small proline-rich protein 2G | 1.38 | 1.26 | 1.48E-03 | 2.70E-02 |
| **Glipr1** | GLI pathogenesis-related 1 (glioma) | 2.75 | 1.25 | 4.74E-04 | 1.31E-02 |
| **Tgfb1** | transforming growth factor, beta 1 | 2.32 | 1.25 | 3.55E-04 | 1.11E-02 |
| **Cyp2d22** | cytochrome P450, family 2, subfamily d, polypeptide 22 | 1.22 | 1.25 | 5.11E-04 | 1.36E-02 |
| **Atf3** | activating transcription factor 3 | 2.25 | 1.25 | 1.89E-05 | 1.70E-03 |
| **E230029C05Rik** | RIKEN cDNA E230029C05 gene | 1.53 | 1.25 | 1.60E-04 | 6.43E-03 |
| **Ccr5** | chemokine (C-C motif) receptor 5 | 1.50 | 1.25 | 3.82E-03 | 4.89E-02 |
| **Nfam1** | Nfat activating molecule with ITAM motif 1 | 2.65 | 1.25 | 3.92E-04 | 1.16E-02 |
| **Mfap2** | microfibrillar-associated protein 2 | 9.24 | 1.25 | 1.29E-03 | 2.47E-02 |
| **Cd9** | CD9 antigen | 2.02 | 1.25 | 4.79E-05 | 2.98E-03 |
| **Ywhah** | tyrosine 3-monooxygenase/tryptophan 5-monooxygenase activation protein, eta polypeptide | 1.59 | 1.25 | 3.56E-06 | 6.95E-04 |
| **Irf1** | interferon regulatory factor 1 | 1.96 | 1.25 | 1.23E-04 | 5.46E-03 |
| **Thsd7a** | thrombospondin, type I, domain containing 7A | 1.14 | 1.25 | 4.67E-04 | 1.30E-02 |
| **Des** | desmin | 2.75 | 1.25 | 1.21E-04 | 5.39E-03 |
| **Esyt1** | extended synaptotagmin-like protein 1 | 2.31 | 1.25 | 1.34E-03 | 2.53E-02 |
| **4632428N05Rik** | RIKEN cDNA 4632428N05 gene | 1.55 | 1.25 | 3.86E-03 | 4.91E-02 |
| **Rgs6** | regulator of G-protein signaling 6 | 1.48 | 1.25 | 2.51E-03 | 3.78E-02 |
| **Isyna1** | myo-inositol 1-phosphate synthase A1 | 1.95 | 1.25 | 1.66E-03 | 2.92E-02 |
| **Vcl** | vinculin | 1.08 | 1.25 | 7.01E-04 | 1.67E-02 |
| **Inhbb** | inhibin beta-B | 1.55 | 1.25 | 6.00E-04 | 1.50E-02 |
| **Slc13a1** | solute carrier family 13 (sodium/sulfate symporters), member 1 | 1.16 | 1.25 | 4.61E-04 | 1.29E-02 |
| **Tubb4a** | tubulin, beta 4A class IVA | 3.09 | 1.25 | 9.69E-04 | 2.07E-02 |
| **Islr** | immunoglobulin superfamily containing leucine-rich repeat | 1.40 | 1.25 | 1.12E-03 | 2.25E-02 |
| **H2-DMa** | histocompatibility 2, class II, locus DMa | 3.91 | 1.24 | 1.40E-03 | 2.62E-02 |
| **Tuba1b** | tubulin, alpha 1B | 1.29 | 1.24 | 2.17E-03 | 3.47E-02 |
| **Sobp** | sine oculis-binding protein homolog (Drosophila) | 2.52 | 1.24 | 2.19E-04 | 7.97E-03 |
| **Alpk1** | alpha-kinase 1 | 1.31 | 1.24 | 1.91E-04 | 7.24E-03 |
| **Ctnnd2** | catenin (cadherin associated protein), delta 2 | 1.73 | 1.24 | 9.45E-04 | 2.03E-02 |
| **B4galt1** | UDP-Gal:betaGlcNAc beta 1,4- galactosyltransferase, polypeptide 1 | 1.68 | 1.24 | 4.15E-04 | 1.20E-02 |
| **Tbc1d2b** | TBC1 domain family, member 2B | 2.12 | 1.24 | 4.57E-04 | 1.28E-02 |
| **Pear1** | platelet endothelial aggregation receptor 1 | 1.75 | 1.24 | 3.48E-03 | 4.60E-02 |
| **Epdr1** | ependymin related protein 1 (zebrafish) | 1.82 | 1.24 | 1.41E-04 | 5.94E-03 |
| **Cald1** | caldesmon 1 | 1.85 | 1.24 | 2.28E-03 | 3.59E-02 |
| **Sulf2** | sulfatase 2 | 1.29 | 1.24 | 9.90E-05 | 4.74E-03 |
| **Sik1** | salt inducible kinase 1 | 1.23 | 1.24 | 3.05E-05 | 2.25E-03 |
| **Hck** | hemopoietic cell kinase | 2.03 | 1.23 | 1.80E-03 | 3.06E-02 |
| **Gpr153** | G protein-coupled receptor 153 | 3.75 | 1.23 | 1.76E-03 | 3.02E-02 |
| **Ciita** | class II transactivator | 4.18 | 1.23 | 1.31E-03 | 2.50E-02 |

| | | | | | |
|---|---|---|---|---|---|
| **Trim30a** | tripartite motif-containing 30A | 1.93 | 1.23 | 1.69E-03 | 2.94E-02 |
| **Mcoln3** | mucolipin 3 | 1.73 | 1.23 | 2.38E-03 | 3.66E-02 |
| **Capg** | capping protein (actin filament), gelsolin-like | 1.70 | 1.23 | 3.01E-04 | 9.93E-03 |
| **Arhgap25** | Rho GTPase activating protein 25 | 1.65 | 1.23 | 2.11E-04 | 7.79E-03 |
| **Csf2ra** | colony stimulating factor 2 receptor, alpha, low-affinity (granulocyte-macrophage) | 2.28 | 1.23 | 6.96E-04 | 1.66E-02 |
| **Htra3** | HtrA serine peptidase 3 | 6.45 | 1.23 | 9.22E-04 | 1.99E-02 |
| **Ddit4l** | DNA-damage-inducible transcript 4-like | 1.66 | 1.23 | 1.40E-03 | 2.62E-02 |
| **Cd276** | CD276 antigen | 2.02 | 1.23 | 2.75E-03 | 4.00E-02 |
| **Egr2** | early growth response 2 | 16.20 | 1.23 | 1.04E-03 | 2.15E-02 |
| **Fbln7** | fibulin 7 | 1.71 | 1.23 | 2.07E-03 | 3.37E-02 |
| **Tes** | testis derived transcript | 2.23 | 1.23 | 2.38E-03 | 3.66E-02 |
| **Lrrc26** | leucine rich repeat containing 26 | 1.24 | 1.23 | 3.13E-03 | 4.32E-02 |
| **Tmem176a** | transmembrane protein 176A | 1.34 | 1.23 | 2.38E-03 | 3.66E-02 |
| **Klf10** | Kruppel-like factor 10 | 1.80 | 1.23 | 2.41E-04 | 8.53E-03 |
| **Fermt3** | fermitin family homolog 3 (Drosophila) | 2.91 | 1.23 | 2.85E-03 | 4.08E-02 |
| **Gucy1a2** | guanylate cyclase 1, soluble, alpha 2 | 2.55 | 1.23 | 7.16E-04 | 1.69E-02 |
| **Cdo1** | cysteine dioxygenase 1, cytosolic | 1.10 | 1.23 | 1.14E-03 | 2.27E-02 |
| **Celf2** | CUGBP, Elav-like family member 2 | 2.02 | 1.23 | 4.88E-04 | 1.32E-02 |
| **Arhgap11a** | Rho GTPase activating protein 11A | 1.32 | 1.23 | 1.44E-04 | 6.02E-03 |
| **Sdk2** | sidekick homolog 2 (chicken) | 1.94 | 1.23 | 2.60E-03 | 3.86E-02 |
| **Clec10a** | C-type lectin domain family 10, member A | 1.32 | 1.23 | 4.66E-04 | 1.30E-02 |
| **1700025G04Rik** | RIKEN cDNA 1700025G04 gene | 1.54 | 1.23 | 8.24E-05 | 4.21E-03 |
| **Cd300a** | CD300A antigen | 1.18 | 1.23 | 3.85E-03 | 4.91E-02 |
| **Ptn** | pleiotrophin | 4.35 | 1.23 | 3.60E-03 | 4.71E-02 |
| **Sntg1** | syntrophin, gamma 1 | 1.34 | 1.23 | 2.96E-03 | 4.17E-02 |
| **Sh3kbp1** | SH3-domain kinase binding protein 1 | 1.75 | 1.23 | 1.11E-03 | 2.24E-02 |
| **Serpinb6b** | serine (or cysteine) peptidase inhibitor, clade B, member 6b | 2.80 | 1.23 | 5.00E-04 | 1.34E-02 |
| **Zc3hav1** | zinc finger CCCH type, antiviral 1 | 3.00 | 1.23 | 2.83E-04 | 9.49E-03 |
| **Ptger4** | prostaglandin E receptor 4 (subtype EP4) | 2.87 | 1.23 | 5.50E-04 | 1.42E-02 |
| **Ostc** | oligosaccharyltransferase complex subunit | 1.23 | 1.22 | 7.50E-04 | 1.74E-02 |
| **Tgif1** | TGFB-induced factor homeobox 1 | 1.59 | 1.22 | 2.51E-03 | 3.78E-02 |
| **Hoxd11** | homeobox D11 | 1.66 | 1.22 | 3.17E-03 | 4.37E-02 |
| **Irf4** | interferon regulatory factor 4 | 4.66 | 1.22 | 2.27E-03 | 3.58E-02 |
| **Sat1** | spermidine/spermine N1-acetyl transferase 1 | 1.04 | 1.22 | 8.69E-05 | 4.32E-03 |
| **Il2rg** | interleukin 2 receptor, gamma chain | 2.76 | 1.22 | 2.20E-03 | 3.51E-02 |
| **Il18r1** | interleukin 18 receptor 1 | 3.44 | 1.22 | 1.19E-03 | 2.34E-02 |
| **Rps20** | ribosomal protein S20 | 1.17 | 1.22 | 7.89E-04 | 1.79E-02 |
| **Wbp5** | WW domain binding protein 5 | 1.48 | 1.22 | 1.46E-03 | 2.67E-02 |
| **Nr4a2** | nuclear receptor subfamily 4, group A, member 2 | 1.02 | 1.22 | 3.67E-04 | 1.12E-02 |
| **Adam19** | a disintegrin and metallopeptidase domain 19 (meltrin beta) | 2.22 | 1.22 | 7.99E-04 | 1.81E-02 |
| **Ifngr2** | interferon gamma receptor 2 | 1.68 | 1.22 | 6.27E-05 | 3.55E-03 |
| **Eef1b2** | eukaryotic translation elongation factor 1 beta 2 | 1.04 | 1.22 | 7.95E-05 | 4.13E-03 |
| **Naip2** | NLR family, apoptosis inhibitory protein 2 | 2.29 | 1.22 | 1.67E-03 | 2.93E-02 |
| **Rgs5** | regulator of G-protein signaling 5 | 2.86 | 1.22 | 1.95E-03 | 3.24E-02 |
| **Cdk6** | cyclin-dependent kinase 6 | 1.78 | 1.21 | 8.26E-05 | 4.21E-03 |
| **Cacnb3** | calcium channel, voltage-dependent, beta 3 subunit | 2.46 | 1.21 | 1.87E-03 | 3.15E-02 |
| **Efhd2** | EF hand domain containing 2 | 2.37 | 1.21 | 2.16E-05 | 1.83E-03 |
| **Tshz2** | teashirt zinc finger family member 2 | 1.05 | 1.21 | 7.22E-04 | 1.70E-02 |
| **Col5a2** | collagen, type V, alpha 2 | 2.14 | 1.21 | 1.75E-04 | 6.80E-03 |
| **Adamtsl2** | ADAMTS-like 2 | 3.32 | 1.21 | 1.44E-03 | 2.65E-02 |
| **C1qtnf6** | C1q and tumor necrosis factor related protein 6 | 1.78 | 1.21 | 3.22E-04 | 1.04E-02 |
| **Ifltd1** | intermediate filament tail domain containing 1 | 1.64 | 1.21 | 1.44E-04 | 6.02E-03 |
| **Ier3** | immediate early response 3 | 1.76 | 1.21 | 7.96E-04 | 1.80E-02 |
| **Cttnbp2nl** | CTTNBP2 N-terminal like | 1.82 | 1.21 | 2.34E-03 | 3.63E-02 |

| | | | | | |
|---|---|---|---|---|---|
| **Nsg2** | neuron specific gene family member 2 | 1.04 | 1.21 | 7.44E-04 | 1.73E-02 |
| **Adamts10** | a disintegrin-like and metallopeptidase (reprolysin type) with thrombospondin type 1 motif, 10 | 2.22 | 1.21 | 7.26E-04 | 1.71E-02 |
| **Trim30d** | tripartite motif-containing 30D | 2.64 | 1.21 | 3.56E-03 | 4.68E-02 |
| **Dclk1** | doublecortin-like kinase 1 | 2.33 | 1.21 | 2.32E-03 | 3.62E-02 |
| **Lacc1** | laccase (multicopper oxidoreductase) domain containing 1 | 2.28 | 1.21 | 2.22E-03 | 3.52E-02 |
| **Fat1** | FAT tumor suppressor homolog 1 (Drosophila) | 1.63 | 1.21 | 2.69E-03 | 3.95E-02 |
| **AA414768** | expressed sequence AA414768 | 1.62 | 1.21 | 5.47E-04 | 1.42E-02 |
| **Grm7** | glutamate receptor, metabotropic 7 | 1.28 | 1.20 | 3.03E-04 | 9.98E-03 |
| **Zyx** | zyxin | 1.78 | 1.20 | 1.51E-05 | 1.53E-03 |
| **Grb10** | growth factor receptor bound protein 10 | 1.87 | 1.20 | 1.00E-03 | 2.11E-02 |
| **Msn** | moesin | 3.07 | 1.20 | 2.86E-04 | 9.58E-03 |
| **Crem** | cAMP responsive element modulator | 1.09 | 1.20 | 2.87E-04 | 9.59E-03 |
| **Oasl2** | 2'-5' oligoadenylate synthetase-like 2 | 3.22 | 1.20 | 3.44E-03 | 4.57E-02 |
| **Man2a1** | mannosidase 2, alpha 1 | 1.32 | 1.20 | 9.08E-04 | 1.97E-02 |
| **Plce1** | phospholipase C, epsilon 1 | 1.16 | 1.20 | 1.83E-04 | 7.01E-03 |
| **P2ry12** | purinergic receptor P2Y, G-protein coupled 12 | 1.32 | 1.20 | 1.61E-03 | 2.88E-02 |
| **Vat1** | vesicle amine transport protein 1 homolog (T californica) | 1.91 | 1.20 | 7.59E-05 | 4.04E-03 |
| **Ngfrap1** | nerve growth factor receptor (TNFRSF16) associated protein 1 | 1.31 | 1.20 | 9.83E-04 | 2.08E-02 |
| **Bhlhe40** | basic helix-loop-helix family, member e40 | 1.28 | 1.20 | 1.15E-03 | 2.29E-02 |
| **Rpl10a** | ribosomal protein L10A | 1.06 | 1.20 | 1.07E-03 | 2.19E-02 |
| **Basp1** | brain abundant, membrane attached signal protein 1 | 3.75 | 1.20 | 5.15E-04 | 1.37E-02 |
| **Fgf12** | fibroblast growth factor 12 | 1.39 | 1.20 | 3.68E-04 | 1.12E-02 |
| **Mkl1** | MKL (megakaryoblastic leukemia)/myocardin-like 1 | 1.68 | 1.20 | 2.04E-03 | 3.33E-02 |
| **Nek6** | NIMA (never in mitosis gene a)-related expressed kinase 6 | 2.30 | 1.20 | 1.88E-04 | 7.16E-03 |
| **Dnm1** | dynamin 1 | 1.65 | 1.20 | 7.80E-05 | 4.10E-03 |
| **Creb3l1** | cAMP responsive element binding protein 3-like 1 | 1.12 | 1.20 | 5.81E-05 | 3.38E-03 |
| **Trp53** | transformation related protein 53 | 1.40 | 1.20 | 1.37E-04 | 5.88E-03 |
| **Smc2** | structural maintenance of chromosomes 2 | 1.24 | 1.20 | 1.29E-03 | 2.47E-02 |
| **Fads1** | fatty acid desaturase 1 | 1.14 | 1.20 | 5.30E-04 | 1.39E-02 |
| **Pdgfrl** | platelet-derived growth factor receptor-like | 3.36 | 1.20 | 9.54E-04 | 2.04E-02 |
| **Myo1e** | myosin IE | 2.22 | 1.20 | 8.94E-04 | 1.95E-02 |
| **Ap3s1** | adaptor-related protein complex 3, sigma 1 subunit | 1.10 | 1.19 | 7.10E-04 | 1.68E-02 |
| **Tmod3** | tropomodulin 3 | 1.46 | 1.19 | 2.52E-04 | 8.78E-03 |
| **Capsl** | calcyphosine-like | 1.38 | 1.19 | 2.21E-03 | 3.52E-02 |
| **Itk** | IL2 inducible T cell kinase | 2.03 | 1.19 | 3.73E-03 | 4.82E-02 |
| **Serpinb6a** | serine (or cysteine) peptidase inhibitor, clade B, member 6a | 2.12 | 1.19 | 3.27E-04 | 1.04E-02 |
| **Guca2b** | guanylate cyclase activator 2b (retina) | 1.15 | 1.19 | 1.84E-03 | 3.12E-02 |
| **Jund** | jun D proto-oncogene | 1.64 | 1.19 | 4.97E-04 | 1.34E-02 |
| **Gucy1b3** | guanylate cyclase 1, soluble, beta 3 | 1.96 | 1.19 | 1.55E-03 | 2.79E-02 |
| **Gbp7** | guanylate binding protein 7 | 1.29 | 1.19 | 4.12E-04 | 1.20E-02 |
| **Syk** | spleen tyrosine kinase | 2.63 | 1.19 | 1.33E-03 | 2.52E-02 |
| **Gpx7** | glutathione peroxidase 7 | 2.46 | 1.19 | 7.41E-04 | 1.73E-02 |
| **Mthfd1l** | methylenetetrahydrofolate dehydrogenase (NADP+ dependent) 1-like | 2.24 | 1.19 | 2.42E-04 | 8.54E-03 |
| **Hmha1** | histocompatibility (minor) HA-1 | 1.31 | 1.19 | 2.42E-03 | 3.68E-02 |
| **Nfkb2** | nuclear factor of kappa light polypeptide gene enhancer in B cells 2, p49/p100 | 3.36 | 1.19 | 7.39E-05 | 3.97E-03 |
| **Hsbp1** | heat shock factor binding protein 1 | 1.16 | 1.19 | 1.13E-03 | 2.26E-02 |
| **Btf3** | basic transcription factor 3 | 1.15 | 1.19 | 3.91E-04 | 1.16E-02 |
| **Tagln2** | transgelin 2 | 1.97 | 1.19 | 2.11E-05 | 1.81E-03 |
| **Sfrp1** | secreted frizzled-related protein 1 | 1.76 | 1.19 | 2.30E-03 | 3.61E-02 |
| **Apaf1** | apoptotic peptidase activating factor 1 | 1.57 | 1.19 | 6.63E-05 | 3.72E-03 |
| **Cst3** | cystatin C | 1.76 | 1.19 | 1.06E-04 | 4.92E-03 |
| **Slc41a2** | solute carrier family 41, member 2 | 1.30 | 1.19 | 7.96E-05 | 4.13E-03 |
| **Actg2** | actin, gamma 2, smooth muscle, enteric | 2.27 | 1.19 | 9.15E-04 | 1.98E-02 |
| **Rtp4** | receptor transporter protein 4 | 3.25 | 1.19 | 3.81E-03 | 4.89E-02 |

| | | | | | |
|---|---|---|---|---|---|
| **Ntn4** | netrin 4 | 1.34 | 1.19 | 2.34E-03 | 3.63E-02 |
| **Cast** | calpastatin | 1.74 | 1.19 | 2.93E-03 | 4.15E-02 |
| **Twf2** | twinfilin, actin-binding protein, homolog 2 (Drosophila) | 1.72 | 1.19 | 4.46E-04 | 1.26E-02 |
| **Samd14** | sterile alpha motif domain containing 14 | 1.36 | 1.19 | 1.48E-03 | 2.69E-02 |
| **Steap1** | six transmembrane epithelial antigen of the prostate 1 | 1.70 | 1.19 | 3.43E-04 | 1.08E-02 |
| **1500009L16Rik** | RIKEN cDNA 1500009L16 gene | 3.54 | 1.19 | 1.24E-03 | 2.40E-02 |
| **Limd2** | LIM domain containing 2 | 1.63 | 1.19 | 2.14E-04 | 7.82E-03 |
| **Snx10** | sorting nexin 10 | 2.44 | 1.19 | 1.14E-03 | 2.28E-02 |
| **Scn1b** | sodium channel, voltage-gated, type I, beta | 1.77 | 1.18 | 3.03E-04 | 9.98E-03 |
| **Rrbp1** | ribosome binding protein 1 | 1.47 | 1.18 | 1.88E-04 | 7.16E-03 |
| **Slamf9** | SLAM family member 9 | 2.13 | 1.18 | 1.11E-03 | 2.24E-02 |
| **Ctps** | cytidine 5'-triphosphate synthase | 1.74 | 1.18 | 1.82E-04 | 6.99E-03 |
| **Rps29** | ribosomal protein S29 | 1.14 | 1.18 | 5.53E-04 | 1.42E-02 |
| **Piezo1** | piezo-type mechanosensitive ion channel component 1 | 1.50 | 1.18 | 1.86E-03 | 3.14E-02 |
| **Tmem123** | transmembrane protein 123 | 1.34 | 1.18 | 3.22E-04 | 1.04E-02 |
| **Cfp** | complement factor properdin | 2.87 | 1.18 | 3.23E-03 | 4.42E-02 |
| **Akna** | AT-hook transcription factor | 2.60 | 1.18 | 1.05E-03 | 2.16E-02 |
| **Fam114a1** | family with sequence similarity 114, member A1 | 2.94 | 1.18 | 1.71E-03 | 2.96E-02 |
| **Emilin1** | elastin microfibril interfacer 1 | 1.05 | 1.18 | 2.68E-03 | 3.94E-02 |
| **Klf3** | Kruppel-like factor 3 (basic) | 1.63 | 1.18 | 7.84E-04 | 1.79E-02 |
| **Dctn2** | dynactin 2 | 1.22 | 1.18 | 9.28E-04 | 2.00E-02 |
| **Anxa5** | annexin A5 | 1.85 | 1.18 | 2.71E-03 | 3.96E-02 |
| **Irf9** | interferon regulatory factor 9 | 3.04 | 1.18 | 8.26E-04 | 1.85E-02 |
| **Gsdmc** | gasdermin C | 1.39 | 1.18 | 2.40E-03 | 3.67E-02 |
| **Plekhg2** | pleckstrin homology domain containing, family G (with RhoGef domain) member 2 | 2.97 | 1.18 | 1.08E-03 | 2.20E-02 |
| **Dock7** | dedicator of cytokinesis 7 | 1.57 | 1.18 | 3.26E-03 | 4.45E-02 |
| **Clic1** | chloride intracellular channel 1 | 1.11 | 1.18 | 1.02E-03 | 2.13E-02 |
| **Gsdmd** | gasdermin D | 1.87 | 1.18 | 3.26E-03 | 4.44E-02 |
| **Unc13b** | unc-13 homolog B (C. elegans) | 1.96 | 1.18 | 1.24E-04 | 5.49E-03 |
| **Kdelc2** | KDEL (Lys-Asp-Glu-Leu) containing 2 | 1.14 | 1.17 | 1.62E-03 | 2.88E-02 |
| **Pcdhb17** | protocadherin beta 17 | 2.28 | 1.17 | 1.19E-03 | 2.34E-02 |
| **Rpl19** | ribosomal protein L19 | 1.00 | 1.17 | 7.72E-04 | 1.77E-02 |
| **Hebp2** | heme binding protein 2 | 1.53 | 1.17 | 3.64E-03 | 4.75E-02 |
| **Cd80** | CD80 antigen | 1.61 | 1.17 | 6.49E-04 | 1.58E-02 |
| **Dopey2** | dopey family member 2 | 1.53 | 1.17 | 1.05E-03 | 2.16E-02 |
| **Myh9** | myosin, heavy polypeptide 9, non-muscle | 1.63 | 1.17 | 7.37E-04 | 1.72E-02 |
| **Nlgn2** | neuroligin 2 | 2.64 | 1.17 | 2.08E-04 | 7.70E-03 |
| **Sept7** | septin 7 | 1.05 | 1.17 | 1.89E-03 | 3.17E-02 |
| **Fxyd1** | FXYD domain-containing ion transport regulator 1 | 2.41 | 1.17 | 3.90E-04 | 1.16E-02 |
| **Ptbp3** | polypyrimidine tract binding protein 3 | 1.06 | 1.17 | 3.77E-04 | 1.14E-02 |
| **Mmrn1** | multimerin 1 | 1.04 | 1.17 | 7.42E-04 | 1.73E-02 |
| **Nkd2** | naked cuticle 2 homolog (Drosophila) | 1.19 | 1.17 | 2.55E-03 | 3.81E-02 |
| **Actg1** | actin, gamma, cytoplasmic 1 | 1.54 | 1.17 | 1.36E-03 | 2.56E-02 |
| **C030034L19Rik** | RIKEN cDNA C030034L19 gene | 1.31 | 1.17 | 1.68E-03 | 2.93E-02 |
| **Pla2r1** | phospholipase A2 receptor 1 | 3.47 | 1.17 | 2.22E-03 | 3.52E-02 |
| **Prelid1** | PRELI domain containing 1 | 1.16 | 1.17 | 4.02E-04 | 1.18E-02 |
| **Tcerg1l** | transcription elongation regulator 1-like | 3.90 | 1.17 | 2.35E-03 | 3.63E-02 |
| **Pld2** | phospholipase D2 | 1.28 | 1.17 | 3.64E-04 | 1.12E-02 |
| **Psmb10** | proteasome (prosome, macropain) subunit, beta type 10 | 2.67 | 1.17 | 1.17E-03 | 2.32E-02 |
| **Psme1** | proteasome (prosome, macropain) activator subunit 1 (PA28 alpha) | 1.29 | 1.17 | 1.03E-04 | 4.85E-03 |
| **Mob3a** | MOB kinase activator 3A | 1.89 | 1.17 | 1.91E-03 | 3.21E-02 |
| **Slc35g1** | solute carrier family 35, member G1 | 1.24 | 1.17 | 1.83E-03 | 3.10E-02 |
| **Plin2** | perilipin 2 | 1.52 | 1.17 | 2.89E-03 | 4.11E-02 |
| **Pfn1** | profilin 1 | 1.38 | 1.17 | 2.67E-04 | 9.05E-03 |

| | | | | | |
|---|---|---|---|---|---|
| **Maged1** | melanoma antigen, family D, 1 | 1.27 | 1.17 | 3.18E-04 | 1.03E-02 |
| **Carhsp1** | calcium regulated heat stable protein 1 | 1.35 | 1.17 | 1.75E-03 | 3.01E-02 |
| **Elk3** | ELK3, member of ETS oncogene family | 1.41 | 1.16 | 7.62E-04 | 1.76E-02 |
| **Ybx3** | Y box protein 3 | 1.14 | 1.16 | 5.30E-04 | 1.39E-02 |
| **Tcf12** | transcription factor 12 | 2.59 | 1.16 | 1.29E-03 | 2.47E-02 |
| **Il6st** | interleukin 6 signal transducer | 2.08 | 1.16 | 1.04E-03 | 2.15E-02 |
| **Trp63** | transformation related protein 63 | 1.22 | 1.16 | 7.92E-04 | 1.79E-02 |
| **Nfkb1** | nuclear factor of kappa light polypeptide gene enhancer in B cells 1, p105 | 2.01 | 1.16 | 2.33E-03 | 3.63E-02 |
| **Nrcam** | neuronal cell adhesion molecule | 1.12 | 1.16 | 1.38E-03 | 2.59E-02 |
| **Dnase1l1** | deoxyribonuclease 1-like 1 | 1.19 | 1.16 | 4.76E-04 | 1.31E-02 |
| **Ly6a** | lymphocyte antigen 6 complex, locus A | 1.25 | 1.16 | 2.13E-04 | 7.81E-03 |
| **Fhl2** | four and a half LIM domains 2 | 3.11 | 1.16 | 3.28E-03 | 4.47E-02 |
| **Nfatc1** | nuclear factor of activated T cells, cytoplasmic, calcineurin dependent 1 | 1.08 | 1.16 | 3.77E-04 | 1.14E-02 |
| **Rps3a1** | ribosomal protein S3A1 | 1.14 | 1.16 | 1.01E-03 | 2.12E-02 |
| **Sirpa** | signal-regulatory protein alpha | 1.23 | 1.16 | 7.24E-05 | 3.91E-03 |
| **4930503E14Rik** | RIKEN cDNA 4930503E14 gene | 6.66 | 1.16 | 3.10E-03 | 4.29E-02 |
| **Ccl19** | chemokine (C-C motif) ligand 19 | 3.42 | 1.16 | 1.79E-03 | 3.06E-02 |
| **Rsu1** | Ras suppressor protein 1 | 1.12 | 1.16 | 3.51E-03 | 4.63E-02 |
| **Rnf213** | ring finger protein 213 | 3.76 | 1.16 | 1.07E-03 | 2.18E-02 |
| **Itih3** | inter-alpha trypsin inhibitor, heavy chain 3 | 1.50 | 1.16 | 2.32E-03 | 3.62E-02 |
| **Morf4l2** | mortality factor 4 like 2 | 1.69 | 1.16 | 3.47E-03 | 4.59E-02 |
| **Map4** | microtubule-associated protein 4 | 1.77 | 1.16 | 1.05E-04 | 4.90E-03 |
| **Ptpn12** | protein tyrosine phosphatase, non-receptor type 12 | 1.66 | 1.16 | 1.68E-03 | 2.93E-02 |
| **Fau** | Finkel-Biskis-Reilly murine sarcoma virus (FBR-MuSV) ubiquitously expressed (fox derived) | 1.13 | 1.16 | 2.25E-03 | 3.56E-02 |
| **Rhoc** | ras homolog gene family, member C | 1.51 | 1.16 | 1.01E-03 | 2.12E-02 |
| **Grhl3** | grainyhead-like 3 (Drosophila) | 1.21 | 1.16 | 1.35E-03 | 2.55E-02 |
| **Map7d1** | MAP7 domain containing 1 | 2.38 | 1.16 | 3.61E-04 | 1.11E-02 |
| **Ppp1r14b** | protein phosphatase 1, regulatory (inhibitor) subunit 14B | 1.50 | 1.16 | 2.11E-03 | 3.41E-02 |
| **Klf4** | Kruppel-like factor 4 (gut) | 1.50 | 1.15 | 3.47E-03 | 4.59E-02 |
| **Ddx58** | DEAD (Asp-Glu-Ala-Asp) box polypeptide 58 | 1.08 | 1.15 | 2.25E-04 | 8.09E-03 |
| **Mapk10** | mitogen-activated protein kinase 10 | 1.21 | 1.15 | 2.06E-03 | 3.36E-02 |
| **Clint1** | clathrin interactor 1 | 1.30 | 1.15 | 2.49E-03 | 3.76E-02 |
| **Atp1b2** | ATPase, Na+/K+ transporting, beta 2 polypeptide | 2.75 | 1.15 | 2.11E-03 | 3.41E-02 |
| **Ifnar1** | interferon (alpha and beta) receptor 1 | 1.28 | 1.15 | 8.01E-04 | 1.81E-02 |
| **Tcaf1** | TRPM8 channel-associated factor 1 | 1.54 | 1.15 | 9.79E-04 | 2.08E-02 |
| **Vasp** | vasodilator-stimulated phosphoprotein | 1.84 | 1.15 | 6.36E-04 | 1.56E-02 |
| **Hn1** | hematological and neurological expressed sequence 1 | 1.77 | 1.15 | 3.51E-03 | 4.63E-02 |
| **Flrt2** | fibronectin leucine rich transmembrane protein 2 | 1.80 | 1.15 | 3.07E-03 | 4.27E-02 |
| **Tnxb** | tenascin XB | 1.38 | 1.15 | 1.83E-03 | 3.10E-02 |
| **Capn5** | calpain 5 | 1.76 | 1.15 | 1.55E-03 | 2.79E-02 |
| **Eef2k** | eukaryotic elongation factor-2 kinase | 1.01 | 1.15 | 7.58E-04 | 1.75E-02 |
| **Tmem43** | transmembrane protein 43 | 1.72 | 1.15 | 1.47E-03 | 2.69E-02 |
| **Myo9b** | myosin IXb | 1.80 | 1.15 | 8.70E-04 | 1.91E-02 |
| **Prrc2a** | proline-rich coiled-coil 2A | 1.83 | 1.15 | 2.87E-03 | 4.10E-02 |
| **Stat1** | signal transducer and activator of transcription 1 | 2.12 | 1.15 | 6.95E-04 | 1.66E-02 |
| **Gorasp2** | golgi reassembly stacking protein 2 | 1.08 | 1.15 | 1.08E-03 | 2.21E-02 |
| **Dync1i2** | dynein cytoplasmic 1 intermediate chain 2 | 1.14 | 1.15 | 2.35E-03 | 3.63E-02 |
| **Tnip1** | TNFAIP3 interacting protein 1 | 1.47 | 1.15 | 2.02E-03 | 3.32E-02 |
| **Cd302** | CD302 antigen | 1.28 | 1.15 | 3.43E-03 | 4.57E-02 |
| **Rps26** | ribosomal protein S26 | 1.18 | 1.14 | 1.26E-03 | 2.42E-02 |
| **Adam10** | a disintegrin and metallopeptidase domain 10 | 1.27 | 1.14 | 1.06E-03 | 2.17E-02 |
| **Csmd1** | CUB and Sushi multiple domains 1 | 1.83 | 1.14 | 8.65E-04 | 1.90E-02 |
| **Ugcg** | UDP-glucose ceramide glucosyltransferase | 1.01 | 1.14 | 7.61E-04 | 1.76E-02 |
| **Sbno2** | strawberry notch homolog 2 (Drosophila) | 1.30 | 1.14 | 9.32E-04 | 2.01E-02 |

| | | | | | |
|---|---|---|---|---|---|
| **Eif3k** | eukaryotic translation initiation factor 3, subunit K | 1.06 | 1.14 | 1.28E-03 | 2.46E-02 |
| **Psmd8** | proteasome (prosome, macropain) 26S subunit, non-ATPase, 8 | 1.08 | 1.14 | 1.13E-03 | 2.26E-02 |
| **Sh3bp1** | SH3-domain binding protein 1 | 2.64 | 1.14 | 7.93E-04 | 1.79E-02 |
| **Pon2** | paraoxonase 2 | 1.30 | 1.14 | 1.24E-03 | 2.40E-02 |
| **Foxc2** | forkhead box C2 | 1.39 | 1.14 | 1.36E-03 | 2.57E-02 |
| **Sec24d** | Sec24 related gene family, member D (S. cerevisiae) | 1.53 | 1.14 | 1.44E-03 | 2.65E-02 |
| **Fam111a** | family with sequence similarity 111, member A | 2.92 | 1.14 | 3.77E-03 | 4.86E-02 |
| **Sh3gl1** | SH3-domain GRB2-like 1 | 1.50 | 1.14 | 1.05E-03 | 2.16E-02 |
| **Slco2a1** | solute carrier organic anion transporter family, member 2a1 | 1.85 | 1.14 | 1.45E-03 | 2.66E-02 |
| **Rac3** | RAS-related C3 botulinum substrate 3 | 1.56 | 1.14 | 3.22E-03 | 4.41E-02 |
| **Vwa5a** | von Willebrand factor A domain containing 5A | 1.47 | 1.14 | 3.00E-03 | 4.19E-02 |
| **Shtn1** | shootin 1 | 1.67 | 1.14 | 9.46E-04 | 2.03E-02 |
| **Tapbpl** | TAP binding protein-like | 1.38 | 1.14 | 1.71E-03 | 2.96E-02 |
| **Traf1** | TNF receptor-associated factor 1 | 3.72 | 1.14 | 9.92E-04 | 2.09E-02 |
| **Plec** | plectin | 2.01 | 1.14 | 1.35E-03 | 2.55E-02 |
| **Rps10** | ribosomal protein S10 | 1.23 | 1.14 | 5.71E-04 | 1.45E-02 |
| **Ikbke** | inhibitor of kappaB kinase epsilon | 2.45 | 1.14 | 1.31E-03 | 2.49E-02 |
| **Sdc1** | syndecan 1 | 2.57 | 1.13 | 5.35E-04 | 1.40E-02 |
| **G6pdx** | glucose-6-phosphate dehydrogenase X-linked | 1.43 | 1.13 | 2.09E-03 | 3.40E-02 |
| **Rps11** | ribosomal protein S11 | 1.35 | 1.13 | 4.58E-04 | 1.28E-02 |
| **Slc50a1** | solute carrier family 50 (sugar transporter), member 1 | 1.35 | 1.13 | 2.39E-03 | 3.67E-02 |
| **Flot1** | flotillin 1 | 1.82 | 1.13 | 1.01E-03 | 2.12E-02 |
| **Amph** | amphiphysin | 2.02 | 1.13 | 1.90E-03 | 3.19E-02 |
| **Bmp2** | bone morphogenetic protein 2 | 6.99 | 1.13 | 3.07E-03 | 4.27E-02 |
| **Mylk** | myosin, light polypeptide kinase | 1.66 | 1.13 | 3.43E-03 | 4.57E-02 |
| **Mapk13** | mitogen-activated protein kinase 13 | 1.18 | 1.13 | 4.11E-04 | 1.20E-02 |
| **Lasp1** | LIM and SH3 protein 1 | 1.53 | 1.13 | 3.80E-03 | 4.88E-02 |
| **St3gal1** | ST3 beta-galactoside alpha-2,3-sialyltransferase 1 | 1.95 | 1.13 | 1.39E-03 | 2.61E-02 |
| **Npc2** | Niemann-Pick type C2 | 1.29 | 1.13 | 1.20E-03 | 2.36E-02 |
| **Mvp** | major vault protein | 2.10 | 1.13 | 3.24E-03 | 4.43E-02 |
| **Sept11** | septin 11 | 2.51 | 1.13 | 1.88E-03 | 3.17E-02 |
| **Coro1b** | coronin, actin binding protein 1B | 1.45 | 1.13 | 9.46E-04 | 2.03E-02 |
| **Eif6** | eukaryotic translation initiation factor 6 | 1.43 | 1.13 | 1.09E-03 | 2.22E-02 |
| **Sept2** | septin 2 | 1.24 | 1.12 | 1.70E-03 | 2.95E-02 |
| **Ezr** | ezrin | 1.19 | 1.12 | 2.61E-03 | 3.86E-02 |
| **Fkbp1a** | FK506 binding protein 1a | 1.56 | 1.12 | 2.81E-03 | 4.04E-02 |
| **Rpl18** | ribosomal protein L18 | 1.91 | 1.12 | 8.02E-04 | 1.81E-02 |
| **Adgra3** | adhesion G protein-coupled receptor A3 | 2.47 | 1.12 | 3.19E-03 | 4.38E-02 |
| **Fsd1l** | fibronectin type III and SPRY domain containing 1-like | 1.39 | 1.12 | 2.79E-03 | 4.04E-02 |
| **Rpl5** | ribosomal protein L5 | 1.27 | 1.12 | 3.73E-03 | 4.82E-02 |
| **Pag1** | phosphoprotein associated with glycosphingolipid microdomains 1 | 1.81 | 1.12 | 2.44E-03 | 3.70E-02 |
| **Pycard** | PYD and CARD domain containing | 1.74 | 1.12 | 3.43E-03 | 4.57E-02 |
| **Anxa4** | annexin A4 | 1.15 | 1.12 | 3.32E-03 | 4.50E-02 |
| **Pde4b** | phosphodiesterase 4B, cAMP specific | 1.46 | 1.12 | 3.90E-03 | 4.94E-02 |
| **Gdpd5** | glycerophosphodiester phosphodiesterase domain containing 5 | 1.44 | 1.12 | 3.22E-03 | 4.41E-02 |
| **Ltc4s** | leukotriene C4 synthase | 1.70 | 1.12 | 1.68E-03 | 2.93E-02 |
| **Tcf7** | transcription factor 7, T cell specific | 1.74 | 1.12 | 2.79E-03 | 4.04E-02 |
| **Scara5** | scavenger receptor class A, member 5 (putative) | 1.91 | 1.12 | 2.26E-03 | 3.56E-02 |
| **Fscn1** | fascin homolog 1, actin bundling protein (Strongylocentrotus purpuratus) | 1.68 | 1.12 | 3.97E-03 | 4.98E-02 |
| **Fam124a** | family with sequence similarity 124, member A | 1.65 | 1.12 | 3.34E-03 | 4.51E-02 |
| **Galnt12** | UDP-N-acetyl-alpha-D-galactosamine:polypeptide N-acetylgalactosaminyltransferase 12 | 1.13 | 1.12 | 2.01E-03 | 3.31E-02 |
| **Myl12b** | myosin, light chain 12B, regulatory | 1.12 | 1.12 | 1.03E-03 | 2.15E-02 |
| **Ptprj** | protein tyrosine phosphatase, receptor type, J | 1.82 | 1.12 | 1.10E-03 | 2.22E-02 |
| **Npdc1** | neural proliferation, differentiation and control 1 | 2.68 | 1.11 | 3.35E-03 | 4.51E-02 |

| | | | | | |
|---|---|---|---|---|---|
| **Rps5** | ribosomal protein S5 | 1.19 | 1.11 | 7.14E-04 | 1.69E-02 |
| **Fam171a1** | family with sequence similarity 171, member A1 | 1.01 | 1.11 | 3.74E-03 | 4.84E-02 |
| **Ddost** | dolichyl-di-phosphooligosaccharide-protein glycotransferase | 1.05 | 1.11 | 2.34E-03 | 3.63E-02 |
| **Adh6a** | alcohol dehydrogenase 6A (class V) | 1.01 | 1.11 | 1.89E-03 | 3.18E-02 |
| **Fnbp1** | formin binding protein 1 | 1.35 | 1.11 | 2.90E-03 | 4.11E-02 |
| **Sass6** | spindle assembly 6 homolog (C. elegans) | 1.57 | 1.11 | 3.33E-03 | 4.50E-02 |
| **Arpc3** | actin related protein 2/3 complex, subunit 3 | 1.27 | 1.11 | 3.67E-03 | 4.78E-02 |
| **Lpp** | LIM domain containing preferred translocation partner in lipoma | 1.84 | 1.10 | 3.42E-03 | 4.57E-02 |
| **Ptpn23** | protein tyrosine phosphatase, non-receptor type 23 | 1.39 | 1.10 | 3.91E-03 | 4.94E-02 |
| **Ptgr1** | prostaglandin reductase 1 | 1.38 | 1.10 | 1.13E-03 | 2.26E-02 |
| **Pabpc1** | poly(A) binding protein, cytoplasmic 1 | 1.25 | 1.10 | 2.26E-03 | 3.57E-02 |
| **Sptlc2** | serine palmitoyltransferase, long chain base subunit 2 | 1.32 | 1.10 | 2.86E-03 | 4.09E-02 |
| **Tnfrsf10b** | tumor necrosis factor receptor superfamily, member 10b | 1.29 | 1.10 | 3.71E-03 | 4.80E-02 |
| **Akt1** | thymoma viral proto-oncogene 1 | 1.29 | 1.10 | 3.31E-03 | 4.49E-02 |
| **Rpl6** | ribosomal protein L6 | 1.11 | 1.10 | 3.61E-03 | 4.72E-02 |
| **Surf2** | surfeit gene 2 | 1.03 | 1.10 | 3.94E-03 | 4.96E-02 |
| **Ccdc88c** | coiled-coil domain containing 88C | 1.71 | 1.10 | 3.78E-03 | 4.87E-02 |
| **Arpc5** | actin related protein 2/3 complex, subunit 5 | 1.32 | 1.09 | 3.31E-03 | 4.49E-02 |
| **Arpc2** | actin related protein 2/3 complex, subunit 2 | 1.39 | 1.09 | 3.81E-03 | 4.89E-02 |
| **Tecpr2** | tectonin beta-propeller repeat containing 2 | 0.99 | 0.92 | 2.69E-03 | 3.95E-02 |
| **Tmem246** | transmembrane protein 246 | 0.98 | 0.92 | 3.16E-03 | 4.35E-02 |
| **Wnt11** | wingless-type MMTV integration site family, member 11 | 0.87 | 0.91 | 3.63E-03 | 4.75E-02 |
| **Slc25a3** | solute carrier family 25 (mitochondrial carrier, phosphate carrier), member 3 | 0.63 | 0.91 | 2.29E-03 | 3.60E-02 |
| **Kank4os** | KN motif and ankyrin repeat domains 4, opposite strand | 0.77 | 0.91 | 2.44E-03 | 3.70E-02 |
| **Mdh1** | malate dehydrogenase 1, NAD (soluble) | 0.74 | 0.90 | 3.91E-03 | 4.94E-02 |
| **2610015P09Rik** | RIKEN cDNA 2610015P09 gene | 0.41 | 0.90 | 3.42E-03 | 4.57E-02 |
| **Ttn** | titin | 0.35 | 0.90 | 3.83E-03 | 4.89E-02 |
| **Nek8** | NIMA (never in mitosis gene a)-related expressed kinase 8 | 0.74 | 0.90 | 1.15E-03 | 2.29E-02 |
| **Glis1** | GLIS family zinc finger 1 | 0.82 | 0.90 | 3.09E-03 | 4.28E-02 |
| **Trmt61b** | tRNA methyltransferase 61B | 0.51 | 0.90 | 3.93E-03 | 4.95E-02 |
| **Snn** | stannin | 0.52 | 0.90 | 1.52E-03 | 2.75E-02 |
| **Ciapin1** | cytokine induced apoptosis inhibitor 1 | 0.75 | 0.90 | 2.43E-03 | 3.69E-02 |
| **Spata17** | spermatogenesis associated 17 | 0.68 | 0.90 | 8.90E-04 | 1.94E-02 |
| **Casd1** | CAS1 domain containing 1 | 0.88 | 0.90 | 2.74E-03 | 4.00E-02 |
| **Myrf** | myelin regulatory factor | 0.96 | 0.90 | 3.30E-03 | 4.49E-02 |
| **1700001C19Rik** | RIKEN cDNA 1700001C19 gene | 0.32 | 0.90 | 9.70E-04 | 2.07E-02 |
| **Atp5f1** | ATP synthase, H+ transporting, mitochondrial F0 complex, subunit B1 | 0.46 | 0.90 | 2.77E-03 | 4.02E-02 |
| **Ccl25** | chemokine (C-C motif) ligand 25 | 0.83 | 0.90 | 1.62E-03 | 2.88E-02 |
| **Sestd1** | SEC14 and spectrin domains 1 | 0.94 | 0.90 | 1.21E-03 | 2.36E-02 |
| **Wls** | wntless homolog (Drosophila) | 0.45 | 0.90 | 1.85E-03 | 3.13E-02 |
| **Dnah11** | dynein, axonemal, heavy chain 11 | 0.62 | 0.90 | 2.53E-03 | 3.79E-02 |
| **Ppp6r2** | protein phosphatase 6, regulatory subunit 2 | 0.60 | 0.89 | 3.31E-03 | 4.49E-02 |
| **Arfgef2** | ADP-ribosylation factor guanine nucleotide-exchange factor 2 (brefeldin A-inhibited) | 0.90 | 0.89 | 3.90E-03 | 4.94E-02 |
| **Atp5a1** | ATP synthase, H+ transporting, mitochondrial F1 complex, alpha subunit 1 | 0.76 | 0.89 | 1.31E-03 | 2.49E-02 |
| **Msi2** | musashi RNA-binding protein 2 | 0.32 | 0.89 | 2.79E-03 | 4.04E-02 |
| **Grin3a** | glutamate receptor ionotropic, NMDA3A | 0.86 | 0.89 | 3.84E-03 | 4.90E-02 |
| **Tbpl2** | TATA box binding protein like 2 | 0.24 | 0.89 | 2.17E-03 | 3.47E-02 |
| **Tmem35** | transmembrane protein 35 | 0.75 | 0.89 | 2.76E-03 | 4.02E-02 |
| **Dtd1** | D-tyrosyl-tRNA deacylase 1 | 0.70 | 0.89 | 2.81E-03 | 4.05E-02 |
| **Sod2** | superoxide dismutase 2, mitochondrial | 0.85 | 0.89 | 1.68E-03 | 2.93E-02 |
| **Pgbd1** | piggyBac transposable element derived 1 | 0.61 | 0.89 | 1.96E-03 | 3.25E-02 |
| **Vwa3a** | von Willebrand factor A domain containing 3A | 0.43 | 0.89 | 5.25E-04 | 1.38E-02 |
| **Prmt3** | protein arginine N-methyltransferase 3 | 0.79 | 0.89 | 8.30E-04 | 1.85E-02 |
| **Crispld1** | cysteine-rich secretory protein LCCL domain containing 1 | 0.90 | 0.89 | 3.42E-03 | 4.57E-02 |

| | | | | | |
|---|---|---|---|---|---|
| **Alad** | aminolevulinate, delta-, dehydratase | 0.98 | 0.89 | 3.84E-03 | 4.90E-02 |
| **Nek4** | NIMA (never in mitosis gene a)-related expressed kinase 4 | 0.84 | 0.89 | 3.01E-03 | 4.21E-02 |
| **Vpreb1** | pre-B lymphocyte gene 1 | 0.59 | 0.89 | 2.11E-03 | 3.41E-02 |
| **Galk2** | galactokinase 2 | 0.68 | 0.89 | 2.74E-03 | 4.00E-02 |
| **Plcd4** | phospholipase C, delta 4 | 0.93 | 0.89 | 2.16E-03 | 3.46E-02 |
| **Bbs9** | Bardet-Biedl syndrome 9 (human) | 0.88 | 0.89 | 2.96E-03 | 4.17E-02 |
| **Socs5** | suppressor of cytokine signaling 5 | 0.31 | 0.89 | 1.61E-03 | 2.87E-02 |
| **Vwa8** | von Willebrand factor A domain containing 8 | 0.71 | 0.89 | 3.40E-03 | 4.57E-02 |
| **Dld** | dihydrolipoamide dehydrogenase | 0.71 | 0.89 | 2.21E-03 | 3.52E-02 |
| **Dfna5** | deafness, autosomal dominant 5 (human) | 0.86 | 0.89 | 2.72E-03 | 3.98E-02 |
| **Sgsm1** | small G protein signaling modulator 1 | 0.98 | 0.89 | 1.49E-03 | 2.70E-02 |
| **Fars2** | phenylalanine-tRNA synthetase 2 (mitochondrial) | 0.83 | 0.89 | 3.14E-03 | 4.33E-02 |
| **Pfpl** | pore forming protein-like | 0.73 | 0.89 | 1.73E-03 | 2.98E-02 |
| **Kcnj11** | potassium inwardly rectifying channel, subfamily J, member 11 | 0.72 | 0.89 | 3.75E-03 | 4.84E-02 |
| **Dnajc13** | DnaJ (Hsp40) homolog, subfamily C, member 13 | 0.74 | 0.88 | 2.11E-03 | 3.41E-02 |
| **Ndufs2** | NADH dehydrogenase (ubiquinone) Fe-S protein 2 | 0.87 | 0.88 | 1.45E-03 | 2.66E-02 |
| **Mok** | MOK protein kinase | 0.86 | 0.88 | 3.48E-03 | 4.60E-02 |
| **Ift172** | intraflagellar transport 172 | 0.84 | 0.88 | 3.95E-03 | 4.97E-02 |
| **Vmn1r211** | vomeronasal 1 receptor 211 | 0.60 | 0.88 | 3.45E-03 | 4.58E-02 |
| **Aktip** | thymoma viral proto-oncogene 1 interacting protein | 0.75 | 0.88 | 2.77E-03 | 4.02E-02 |
| **Acat1** | acetyl-Coenzyme A acetyltransferase 1 | 0.61 | 0.88 | 1.47E-03 | 2.69E-02 |
| **Smad6** | SMAD family member 6 | 0.83 | 0.88 | 3.43E-03 | 4.57E-02 |
| **Whrn** | whirlin | 0.49 | 0.88 | 2.47E-03 | 3.74E-02 |
| **Mrps14** | mitochondrial ribosomal protein S14 | 0.72 | 0.88 | 1.19E-03 | 2.34E-02 |
| **Alox12** | arachidonate 12-lipoxygenase | 0.79 | 0.88 | 1.62E-03 | 2.88E-02 |
| **Atp5k** | ATP synthase, H+ transporting, mitochondrial F1F0 complex, subunit E | 0.85 | 0.88 | 3.43E-03 | 4.57E-02 |
| **Cog2** | component of oligomeric golgi complex 2 | 0.38 | 0.88 | 1.44E-03 | 2.65E-02 |
| **Cog6** | component of oligomeric golgi complex 6 | 0.92 | 0.88 | 2.96E-03 | 4.17E-02 |
| **Helb** | helicase (DNA) B | 0.66 | 0.88 | 3.70E-03 | 4.80E-02 |
| **Snx27** | sorting nexin family member 27 | 0.75 | 0.88 | 3.19E-03 | 4.39E-02 |
| **Ndufc2** | NADH dehydrogenase (ubiquinone) 1, subcomplex unknown, 2 | 0.79 | 0.88 | 8.29E-04 | 1.85E-02 |
| **Atp2c1** | ATPase, Ca++-sequestering | 0.94 | 0.88 | 9.88E-04 | 2.09E-02 |
| **Acadsb** | acyl-Coenzyme A dehydrogenase, short/branched chain | 0.68 | 0.88 | 3.64E-03 | 4.75E-02 |
| **4930430F08Rik** | RIKEN cDNA 4930430F08 gene | 0.95 | 0.88 | 2.48E-03 | 3.76E-02 |
| **Cdadc1** | cytidine and dCMP deaminase domain containing 1 | 0.80 | 0.88 | 1.72E-03 | 2.97E-02 |
| **Vamp8** | vesicle-associated membrane protein 8 | 0.96 | 0.88 | 5.86E-04 | 1.47E-02 |
| **Gk** | glycerol kinase | 0.55 | 0.88 | 1.88E-03 | 3.17E-02 |
| **Ccdc176** | coiled-coil domain containing 176 | 0.23 | 0.88 | 3.25E-03 | 4.44E-02 |
| **Celsr2** | cadherin, EGF LAG seven-pass G-type receptor 2 (flamingo homolog, Drosophila) | 0.79 | 0.88 | 1.93E-03 | 3.22E-02 |
| **Olfr1166** | olfactory receptor 1166 | 0.69 | 0.88 | 3.17E-03 | 4.37E-02 |
| **Rab11fip3** | RAB11 family interacting protein 3 (class II) | 0.72 | 0.88 | 3.12E-03 | 4.31E-02 |
| **Ghitm** | growth hormone inducible transmembrane protein | 0.88 | 0.88 | 3.71E-03 | 4.80E-02 |
| **Gabra4** | gamma-aminobutyric acid (GABA) A receptor, subunit alpha 4 | 0.21 | 0.87 | 5.22E-04 | 1.38E-02 |
| **Dlat** | dihydrolipoamide S-acetyltransferase (E2 component of pyruvate dehydrogenase complex) | 0.82 | 0.87 | 3.80E-03 | 4.88E-02 |
| **Pex7** | peroxisomal biogenesis factor 7 | 0.67 | 0.87 | 1.87E-03 | 3.15E-02 |
| **Wwc2** | WW, C2 and coiled-coil domain containing 2 | 0.52 | 0.87 | 1.23E-03 | 2.39E-02 |
| **Adck1** | aarF domain containing kinase 1 | 0.84 | 0.87 | 1.11E-03 | 2.23E-02 |
| **Coq9** | coenzyme Q9 homolog (yeast) | 0.83 | 0.87 | 1.53E-03 | 2.76E-02 |
| **Atp6v1a** | ATPase, H+ transporting, lysosomal V1 subunit A | 0.92 | 0.87 | 3.93E-03 | 4.95E-02 |
| **Park2** | Parkinson disease (autosomal recessive, juvenile) 2, parkin | 0.51 | 0.87 | 1.67E-03 | 2.93E-02 |
| **Car2** | carbonic anhydrase 2 | 0.77 | 0.87 | 8.29E-04 | 1.85E-02 |
| **Rnmtl1** | RNA methyltransferase like 1 | 0.95 | 0.87 | 2.83E-03 | 4.06E-02 |
| **Mtx2** | metaxin 2 | 0.95 | 0.87 | 1.05E-03 | 2.16E-02 |
| **Nt5dc1** | 5'-nucleotidase domain containing 1 | 0.96 | 0.87 | 2.53E-03 | 3.79E-02 |

| | | | | | |
|---|---|---|---|---|---|
| **Bbs1** | Bardet-Biedl syndrome 1 (human) | 0.99 | 0.87 | 3.74E-03 | 4.84E-02 |
| **Pnoc** | prepronociceptin | 0.71 | 0.87 | 3.53E-03 | 4.64E-02 |
| **Gfm1** | G elongation factor, mitochondrial 1 | 0.70 | 0.87 | 2.86E-03 | 4.08E-02 |
| **Trmu** | tRNA 5-methylaminomethyl-2-thiouridylate methyltransferase | 0.84 | 0.87 | 5.07E-04 | 1.36E-02 |
| **5730409K12Rik** | RIKEN cDNA 5730409K12 gene | 0.91 | 0.87 | 1.64E-03 | 2.90E-02 |
| **Smlr1** | small leucine-rich protein 1 | 0.44 | 0.87 | 2.67E-03 | 3.93E-02 |
| **Tm4sf1** | transmembrane 4 superfamily member 1 | 0.76 | 0.87 | 1.73E-03 | 2.98E-02 |
| **Peli2** | pellino 2 | 0.95 | 0.87 | 2.64E-03 | 3.90E-02 |
| **Fig4** | FIG4 homolog (S. cerevisiae) | 0.72 | 0.87 | 3.26E-03 | 4.45E-02 |
| **Pon3** | paraoxonase 3 | 1.00 | 0.87 | 6.22E-04 | 1.54E-02 |
| **Zfp839** | zinc finger protein 839 | 0.76 | 0.87 | 1.18E-03 | 2.33E-02 |
| **Nos1ap** | nitric oxide synthase 1 (neuronal) adaptor protein | 0.99 | 0.87 | 3.90E-03 | 4.94E-02 |
| **Cox4i1** | cytochrome c oxidase subunit IV isoform 1 | 0.83 | 0.87 | 2.87E-03 | 4.10E-02 |
| **Ndufv1** | NADH dehydrogenase (ubiquinone) flavoprotein 1 | 0.74 | 0.87 | 8.14E-04 | 1.83E-02 |
| **D830013O20Rik** | RIKEN cDNA D830013O20 gene | 0.80 | 0.87 | 2.16E-03 | 3.46E-02 |
| **Acads** | acyl-Coenzyme A dehydrogenase, short chain | 0.88 | 0.87 | 2.96E-03 | 4.17E-02 |
| **Fam160a2** | family with sequence similarity 160, member A2 | 0.57 | 0.87 | 3.32E-03 | 4.49E-02 |
| **Sh3tc1** | SH3 domain and tetratricopeptide repeats 1 | 0.80 | 0.87 | 3.25E-03 | 4.44E-02 |
| **Nxt2** | nuclear transport factor 2-like export factor 2 | 0.70 | 0.87 | 1.09E-03 | 2.22E-02 |
| **Tbcel** | tubulin folding cofactor E-like | 0.77 | 0.87 | 2.20E-03 | 3.51E-02 |
| **Clpp** | caseinolytic mitochondrial matrix peptidase proteolytic subunit | 0.96 | 0.87 | 3.12E-03 | 4.31E-02 |
| **Gm20555** | predicted gene, 20555 | 0.39 | 0.87 | 1.97E-03 | 3.26E-02 |
| **Ptgr2** | prostaglandin reductase 2 | 0.52 | 0.87 | 6.08E-04 | 1.52E-02 |
| **Acsf3** | acyl-CoA synthetase family member 3 | 0.88 | 0.87 | 3.67E-03 | 4.78E-02 |
| **Slc51b** | solute carrier family 51, beta subunit | 0.82 | 0.87 | 2.61E-03 | 3.86E-02 |
| **Abhd6** | abhydrolase domain containing 6 | 0.89 | 0.87 | 3.48E-03 | 4.60E-02 |
| **9430037G07Rik** | RIKEN cDNA 9430037G07 gene | 0.92 | 0.87 | 3.05E-03 | 4.24E-02 |
| **Mipep** | mitochondrial intermediate peptidase | 0.84 | 0.87 | 1.92E-03 | 3.21E-02 |
| **Gtpbp10** | GTP-binding protein 10 (putative) | 0.67 | 0.87 | 2.47E-03 | 3.74E-02 |
| **Atg4d** | autophagy related 4D, cysteine peptidase | 0.85 | 0.86 | 1.20E-03 | 2.36E-02 |
| **Lrp2** | low density lipoprotein receptor-related protein 2 | 0.68 | 0.86 | 3.79E-04 | 1.14E-02 |
| **Foxred1** | FAD-dependent oxidoreductase domain containing 1 | 0.83 | 0.86 | 1.94E-03 | 3.23E-02 |
| **Tmem245** | transmembrane protein 245 | 0.76 | 0.86 | 1.60E-03 | 2.86E-02 |
| **Agl** | amylo-1,6-glucosidase, 4-alpha-glucanotransferase | 0.69 | 0.86 | 3.78E-03 | 4.87E-02 |
| **Oat** | ornithine aminotransferase | 0.86 | 0.86 | 3.57E-03 | 4.69E-02 |
| **Ovol1** | OVO homolog-like 1 (Drosophila) | 0.75 | 0.86 | 2.51E-03 | 3.78E-02 |
| **Ahcyl2** | S-adenosylhomocysteine hydrolase-like 2 | 0.90 | 0.86 | 3.79E-03 | 4.87E-02 |
| **Wsb2** | WD repeat and SOCS box-containing 2 | 0.92 | 0.86 | 6.82E-04 | 1.64E-02 |
| **Tekt5** | tektin 5 | 0.63 | 0.86 | 4.85E-04 | 1.32E-02 |
| **Mto1** | mitochondrial translation optimization 1 homolog (S. cerevisiae) | 0.80 | 0.86 | 2.89E-03 | 4.11E-02 |
| **Thnsl2** | threonine synthase-like 2 (bacterial) | 0.58 | 0.86 | 3.24E-03 | 4.43E-02 |
| **Etfa** | electron transferring flavoprotein, alpha polypeptide | 0.76 | 0.86 | 5.82E-04 | 1.47E-02 |
| **Bphl** | biphenyl hydrolase-like (serine hydrolase, breast epithelial mucin-associated antigen) | 0.64 | 0.86 | 2.99E-03 | 4.19E-02 |
| **Ndufb5** | NADH dehydrogenase (ubiquinone) 1 beta subcomplex, 5 | 0.84 | 0.86 | 1.31E-03 | 2.49E-02 |
| **Entpd5** | ectonucleoside triphosphate diphosphohydrolase 5 | 0.81 | 0.86 | 1.42E-03 | 2.64E-02 |
| **Samm50** | sorting and assembly machinery component 50 homolog (S. cerevisiae) | 0.86 | 0.86 | 1.82E-03 | 3.09E-02 |
| **Fdx1** | ferredoxin 1 | 0.69 | 0.86 | 1.67E-03 | 2.93E-02 |
| **Rab28** | RAB28, member RAS oncogene family | 0.86 | 0.86 | 8.20E-04 | 1.84E-02 |
| **Smpd1** | sphingomyelin phosphodiesterase 1, acid lysosomal | 0.88 | 0.86 | 5.64E-04 | 1.44E-02 |
| **Acadvl** | acyl-Coenzyme A dehydrogenase, very long chain | 0.83 | 0.86 | 1.60E-03 | 2.86E-02 |
| **Palm3** | paralemmin 3 | 0.78 | 0.86 | 1.65E-03 | 2.91E-02 |
| **Zc2hc1c** | zinc finger, C2HC-type containing 1C | 0.81 | 0.86 | 1.39E-03 | 2.61E-02 |
| **Galnt11** | UDP-N-acetyl-alpha-D-galactosamine:polypeptide N-acetylgalactosaminyltransferase 11 | 0.52 | 0.86 | 9.74E-04 | 2.07E-02 |
| **Slc15a1** | solute carrier family 15 (oligopeptide transporter), member 1 | 0.51 | 0.86 | 6.35E-04 | 1.56E-02 |

| | | | | | |
|---|---|---|---|---|---|
| **Fgfr4** | fibroblast growth factor receptor 4 | 0.89 | 0.86 | 2.97E-03 | 4.17E-02 |
| **Wscd1** | WSC domain containing 1 | 0.96 | 0.86 | 6.89E-04 | 1.65E-02 |
| **Clcn3** | chloride channel 3 | 0.57 | 0.86 | 1.43E-03 | 2.65E-02 |
| **Ninl** | ninein-like | 0.93 | 0.86 | 2.58E-04 | 8.86E-03 |
| **Lonp1** | lon peptidase 1, mitochondrial | 0.96 | 0.86 | 3.45E-03 | 4.58E-02 |
| **Slc26a7** | solute carrier family 26, member 7 | 0.22 | 0.86 | 2.49E-03 | 3.76E-02 |
| **Tmem229a** | transmembrane protein 229A | 0.78 | 0.86 | 2.42E-03 | 3.68E-02 |
| **Adssl1** | adenylosuccinate synthetase like 1 | 0.50 | 0.85 | 3.99E-04 | 1.17E-02 |
| **Tmtc4** | transmembrane and tetratricopeptide repeat containing 4 | 0.40 | 0.85 | 1.04E-03 | 2.15E-02 |
| **Trpt1** | tRNA phosphotransferase 1 | 0.82 | 0.85 | 3.47E-04 | 1.09E-02 |
| **Ahi1** | Abelson helper integration site 1 | 0.41 | 0.85 | 1.18E-03 | 2.33E-02 |
| **Ecsit** | ECSIT homolog (Drosophila) | 0.81 | 0.85 | 1.63E-03 | 2.88E-02 |
| **P2rx3** | purinergic receptor P2X, ligand-gated ion channel, 3 | 0.60 | 0.85 | 1.29E-04 | 5.68E-03 |
| **Rap1gap2** | RAP1 GTPase activating protein 2 | 0.74 | 0.85 | 4.11E-04 | 1.20E-02 |
| **Pdpr** | pyruvate dehydrogenase phosphatase regulatory subunit | 0.65 | 0.85 | 2.90E-03 | 4.11E-02 |
| **Tinag** | tubulointerstitial nephritis antigen | 0.67 | 0.85 | 1.17E-03 | 2.31E-02 |
| **Acyp2** | acylphosphatase 2, muscle type | 0.84 | 0.85 | 3.91E-03 | 4.94E-02 |
| **Fam63b** | family with sequence similarity 63, member B | 0.94 | 0.85 | 4.99E-04 | 1.34E-02 |
| **Stard7** | START domain containing 7 | 0.78 | 0.85 | 1.31E-03 | 2.49E-02 |
| **1500035N22Rik** | RIKEN cDNA 1500035N22 gene | 0.72 | 0.85 | 1.78E-03 | 3.04E-02 |
| **Rp1** | retinitis pigmentosa 1 (human) | 0.70 | 0.85 | 1.03E-03 | 2.15E-02 |
| **Idh2** | isocitrate dehydrogenase 2 (NADP+), mitochondrial | 0.89 | 0.85 | 3.64E-04 | 1.12E-02 |
| **Pcsk7** | proprotein convertase subtilisin/kexin type 7 | 0.97 | 0.85 | 4.52E-04 | 1.27E-02 |
| **Tln2** | talin 2 | 0.57 | 0.85 | 4.65E-04 | 1.30E-02 |
| **Dusp15** | dual specificity phosphatase-like 15 | 0.72 | 0.85 | 1.46E-03 | 2.67E-02 |
| **Fastkd2** | FAST kinase domains 2 | 0.71 | 0.85 | 9.98E-04 | 2.10E-02 |
| **Tmem42** | transmembrane protein 42 | 0.87 | 0.85 | 1.96E-03 | 3.25E-02 |
| **Fn3k** | fructosamine 3 kinase | 0.70 | 0.85 | 1.60E-03 | 2.86E-02 |
| **Lace1** | lactation elevated 1 | 0.79 | 0.85 | 2.81E-03 | 4.04E-02 |
| **Mrpl11** | mitochondrial ribosomal protein L11 | 0.94 | 0.85 | 1.42E-03 | 2.64E-02 |
| **Arl6ip1** | ADP-ribosylation factor-like 6 interacting protein 1 | 0.82 | 0.85 | 3.79E-04 | 1.14E-02 |
| **Sucla2** | succinate-Coenzyme A ligase, ADP-forming, beta subunit | 0.76 | 0.85 | 1.47E-04 | 6.09E-03 |
| **Trmt10c** | tRNA methyltransferase 10C | 0.87 | 0.84 | 3.53E-03 | 4.64E-02 |
| **Kcnj3** | potassium inwardly-rectifying channel, subfamily J, member 3 | 0.94 | 0.84 | 2.12E-03 | 3.43E-02 |
| **Ube2ql1** | ubiquitin-conjugating enzyme E2Q family-like 1 | 0.49 | 0.84 | 5.35E-05 | 3.21E-03 |
| **Ift122** | intraflagellar transport 122 | 0.77 | 0.84 | 6.30E-04 | 1.55E-02 |
| **Tmem38a** | transmembrane protein 38A | 0.62 | 0.84 | 1.25E-03 | 2.41E-02 |
| **Ganc** | "glucosidase, alpha; neutral C" | 0.99 | 0.84 | 5.41E-04 | 1.41E-02 |
| **D3Ertd751e** | DNA segment, Chr 3, ERATO Doi 751, expressed | 0.50 | 0.84 | 3.64E-03 | 4.75E-02 |
| **Kazald1** | Kazal-type serine peptidase inhibitor domain 1 | 0.83 | 0.84 | 1.44E-03 | 2.65E-02 |
| **Gpr162** | G protein-coupled receptor 162 | 0.78 | 0.84 | 5.78E-04 | 1.46E-02 |
| **2900055J20Rik** | RIKEN cDNA 2900055J20 gene | 0.84 | 0.84 | 2.32E-03 | 3.62E-02 |
| **Olfr642** | olfactory receptor 642 | 0.48 | 0.84 | 3.92E-03 | 4.95E-02 |
| **Adcy6** | adenylate cyclase 6 | 0.98 | 0.84 | 2.41E-03 | 3.68E-02 |
| **Plekha7** | pleckstrin homology domain containing, family A member 7 | 0.36 | 0.84 | 3.66E-04 | 1.12E-02 |
| **Gm20187** | predicted gene, 20187 | 0.73 | 0.84 | 1.01E-03 | 2.12E-02 |
| **Nr3c2** | nuclear receptor subfamily 3, group C, member 2 | 0.51 | 0.84 | 2.87E-03 | 4.09E-02 |
| **Gatm** | glycine amidinotransferase (L-arginine:glycine amidinotransferase) | 0.39 | 0.84 | 2.82E-03 | 4.05E-02 |
| **Ckb** | creatine kinase, brain | 0.95 | 0.84 | 2.77E-03 | 4.02E-02 |
| **Acvr1b** | activin A receptor, type 1B | 0.70 | 0.84 | 1.39E-04 | 5.94E-03 |
| **Tenm2** | teneurin transmembrane protein 2 | 0.90 | 0.84 | 9.93E-05 | 4.75E-03 |
| **Lrpprc** | leucine-rich PPR-motif containing | 0.47 | 0.84 | 2.02E-03 | 3.32E-02 |
| **Pptc7** | PTC7 protein phosphatase homolog (S. cerevisiae) | 0.96 | 0.84 | 2.27E-03 | 3.57E-02 |
| **Pth1r** | parathyroid hormone 1 receptor | 0.78 | 0.84 | 1.13E-03 | 2.26E-02 |

| | | | | | |
|---|---|---|---|---|---|
| **Mmaa** | methylmalonic aciduria (cobalamin deficiency) type A | 0.92 | 0.84 | 1.65E-03 | 2.91E-02 |
| **Uqcrh** | ubiquinol-cytochrome c reductase hinge protein | 0.91 | 0.84 | 8.07E-04 | 1.82E-02 |
| **Slc4a5** | solute carrier family 4, sodium bicarbonate cotransporter, member 5 | 0.79 | 0.84 | 6.48E-06 | 9.08E-04 |
| **Kit** | kit oncogene | 0.93 | 0.84 | 6.55E-04 | 1.59E-02 |
| **Fitm2** | fat storage-inducing transmembrane protein 2 | 0.75 | 0.84 | 2.01E-03 | 3.31E-02 |
| **Slc9a4** | solute carrier family 9 (sodium/hydrogen exchanger), member 4 | 0.69 | 0.84 | 1.45E-03 | 2.66E-02 |
| **Phldb2** | pleckstrin homology-like domain, family B, member 2 | 0.60 | 0.84 | 2.80E-03 | 4.04E-02 |
| **Sema3b** | sema domain, immunoglobulin domain (Ig), short basic domain, secreted, (semaphorin) 3B | 0.91 | 0.84 | 1.70E-03 | 2.95E-02 |
| **1700019G24Rik** | RIKEN cDNA 1700019G24 gene | 0.47 | 0.84 | 3.60E-04 | 1.11E-02 |
| **Bckdha** | branched chain ketoacid dehydrogenase E1, alpha polypeptide | 0.77 | 0.84 | 1.32E-03 | 2.51E-02 |
| **Prkcq** | protein kinase C, theta | 0.95 | 0.84 | 1.23E-04 | 5.46E-03 |
| **Ndrg2** | N-myc downstream regulated gene 2 | 0.57 | 0.84 | 3.80E-04 | 1.14E-02 |
| **Mthfd1** | methylenetetrahydrofolate dehydrogenase (NADP+ dependent), methenyltetrahydrofolate cyclohydrolase, formyltetrahydrofolate synthase | 0.61 | 0.84 | 2.55E-03 | 3.81E-02 |
| **Tefm** | transcription elongation factor, mitochondrial | 0.73 | 0.84 | 2.87E-03 | 4.10E-02 |
| **Mtx3** | metaxin 3 | 0.72 | 0.84 | 3.09E-03 | 4.28E-02 |
| **A930033H14Rik** | RIKEN cDNA A930033H14 gene | 0.89 | 0.84 | 2.39E-04 | 8.47E-03 |
| **Tgds** | TDP-glucose 4,6-dehydratase | 0.86 | 0.84 | 5.39E-04 | 1.40E-02 |
| **E330009J07Rik** | RIKEN cDNA E330009J07 gene | 0.93 | 0.84 | 2.73E-03 | 3.99E-02 |
| **Atp5sl** | ATP5S-like | 0.60 | 0.84 | 1.88E-04 | 7.15E-03 |
| **C030037D09Rik** | RIKEN cDNA C030037D09 gene | 0.63 | 0.84 | 1.46E-03 | 2.67E-02 |
| **Tcaim** | T cell activation inhibitor, mitochondrial | 0.64 | 0.84 | 1.27E-03 | 2.45E-02 |
| **Pter** | phosphotriesterase related | 0.63 | 0.83 | 2.37E-03 | 3.65E-02 |
| **Plau** | plasminogen activator, urokinase | 0.62 | 0.83 | 1.18E-03 | 2.33E-02 |
| **Kcnn2** | potassium intermediate/small conductance calcium-activated channel, subfamily N, member 2 | 0.95 | 0.83 | 5.50E-05 | 3.28E-03 |
| **Cpeb2** | cytoplasmic polyadenylation element binding protein 2 | 0.91 | 0.83 | 4.77E-05 | 2.98E-03 |
| **Msrb2** | methionine sulfoxide reductase B2 | 0.90 | 0.83 | 4.49E-04 | 1.27E-02 |
| **Aspg** | asparaginase homolog (S. cerevisiae) | 0.88 | 0.83 | 1.31E-03 | 2.49E-02 |
| **Pdzk1** | PDZ domain containing 1 | 0.64 | 0.83 | 9.12E-04 | 1.98E-02 |
| **Mfsd1** | major facilitator superfamily domain containing 1 | 0.70 | 0.83 | 1.62E-03 | 2.88E-02 |
| **Rnase9** | ribonuclease, RNase A family, 9 (non-active) | 0.48 | 0.83 | 8.42E-04 | 1.87E-02 |
| **Tfam** | transcription factor A, mitochondrial | 0.98 | 0.83 | 9.81E-04 | 2.08E-02 |
| **Gm4890** | predicted gene 4890 | 0.76 | 0.83 | 2.74E-05 | 2.14E-03 |
| **Ndufa9** | NADH dehydrogenase (ubiquinone) 1 alpha subcomplex, 9 | 0.70 | 0.83 | 1.60E-03 | 2.86E-02 |
| **Stard8** | START domain containing 8 | 0.84 | 0.83 | 4.19E-04 | 1.21E-02 |
| **Slc6a12** | solute carrier family 6 (neurotransmitter transporter, betaine/GABA), member 12 | 0.54 | 0.83 | 3.91E-03 | 4.94E-02 |
| **P2ry1** | purinergic receptor P2Y, G-protein coupled 1 | 0.71 | 0.83 | 2.04E-03 | 3.34E-02 |
| **Zfp367** | zinc finger protein 367 | 0.94 | 0.83 | 7.72E-04 | 1.77E-02 |
| **Ptgfr** | prostaglandin F receptor | 0.89 | 0.83 | 2.71E-03 | 3.96E-02 |
| **Fzd5** | frizzled homolog 5 (Drosophila) | 0.66 | 0.83 | 2.53E-03 | 3.79E-02 |
| **Tm2d2** | TM2 domain containing 2 | 0.83 | 0.83 | 3.83E-04 | 1.15E-02 |
| **Scrn3** | secernin 3 | 0.57 | 0.83 | 2.66E-03 | 3.92E-02 |
| **Abcb8** | ATP-binding cassette, sub-family B (MDR/TAP), member 8 | 0.55 | 0.83 | 3.05E-03 | 4.24E-02 |
| **Bmp5** | bone morphogenetic protein 5 | 0.67 | 0.83 | 1.54E-04 | 6.25E-03 |
| **Tmem86b** | transmembrane protein 86B | 0.43 | 0.83 | 2.34E-03 | 3.63E-02 |
| **Sgms2** | sphingomyelin synthase 2 | 0.86 | 0.83 | 1.78E-03 | 3.04E-02 |
| **Tmem125** | transmembrane protein 125 | 0.83 | 0.83 | 2.20E-04 | 7.97E-03 |
| **Cysltr2** | cysteinyl leukotriene receptor 2 | 0.91 | 0.83 | 6.64E-04 | 1.60E-02 |
| **Sowaha** | sosondowah ankyrin repeat domain family member A | 0.94 | 0.83 | 5.78E-05 | 3.38E-03 |
| **Zxdb** | zinc finger, X-linked, duplicated B | 0.83 | 0.82 | 2.70E-03 | 3.95E-02 |
| **Odf4** | outer dense fiber of sperm tails 4 | 0.60 | 0.82 | 1.13E-04 | 5.15E-03 |
| **1700009J07Rik** | RIKEN cDNA 1700009J07 gene | 0.48 | 0.82 | 5.48E-05 | 3.27E-03 |
| **Map7** | microtubule-associated protein 7 | 0.86 | 0.82 | 3.67E-04 | 1.12E-02 |
| **Slc35a1** | solute carrier family 35 (CMP-sialic acid transporter), member 1 | 0.99 | 0.82 | 2.19E-03 | 3.50E-02 |
| **Ndufs1** | NADH dehydrogenase (ubiquinone) Fe-S protein 1 | 0.81 | 0.82 | 2.96E-04 | 9.82E-03 |

| | | | | | |
|---|---|---|---|---|---|
| **Slc35a5** | solute carrier family 35, member A5 | 0.97 | 0.82 | 1.79E-04 | 6.90E-03 |
| **Fut9** | fucosyltransferase 9 | 0.69 | 0.82 | 3.60E-04 | 1.11E-02 |
| **Pfkl** | phosphofructokinase, liver, B-type | 0.44 | 0.82 | 3.94E-04 | 1.17E-02 |
| **Tmem50b** | transmembrane protein 50B | 0.75 | 0.82 | 1.50E-03 | 2.71E-02 |
| **Hadhb** | hydroxyacyl-Coenzyme A dehydrogenase/3-ketoacyl-Coenzyme A thiolase/enoyl-Coenzyme A hydratase (trifunctional protein), beta subunit | 0.76 | 0.82 | 2.60E-04 | 8.93E-03 |
| **Notum** | notum pectinacetylesterase homolog (Drosophila) | 0.49 | 0.82 | 2.77E-04 | 9.31E-03 |
| **Map3k15** | mitogen-activated protein kinase kinase kinase 15 | 0.80 | 0.82 | 3.28E-03 | 4.47E-02 |
| **Spag5** | sperm associated antigen 5 | 0.60 | 0.82 | 4.34E-04 | 1.24E-02 |
| **Cldn17** | claudin 17 | 0.91 | 0.82 | 5.91E-04 | 1.48E-02 |
| **Tarsl2** | threonyl-tRNA synthetase-like 2 | 0.73 | 0.82 | 4.64E-04 | 1.29E-02 |
| **Aldh5a1** | aldhehyde dehydrogenase family 5, subfamily A1 | 0.76 | 0.82 | 2.84E-03 | 4.08E-02 |
| **Pah** | phenylalanine hydroxylase | 0.38 | 0.82 | 2.30E-03 | 3.60E-02 |
| **Ndufaf7** | NADH dehydrogenase (ubiquinone) 1 alpha subcomplex assembly factor 7 | 0.80 | 0.82 | 5.54E-04 | 1.42E-02 |
| **Ptpdc1** | protein tyrosine phosphatase domain containing 1 | 0.89 | 0.82 | 1.04E-04 | 4.88E-03 |
| **Stxbp4** | syntaxin binding protein 4 | 0.94 | 0.82 | 1.72E-03 | 2.97E-02 |
| **Pcyt2** | phosphate cytidylyltransferase 2, ethanolamine | 0.91 | 0.82 | 4.13E-04 | 1.20E-02 |
| **Syt17** | synaptotagmin XVII | 0.64 | 0.82 | 1.10E-03 | 2.22E-02 |
| **Fgf9** | fibroblast growth factor 9 | 0.69 | 0.82 | 3.65E-04 | 1.12E-02 |
| **5730403I07Rik** | RIKEN cDNA 5730403I07 gene | 0.39 | 0.82 | 2.52E-03 | 3.79E-02 |
| **1700028J19Rik** | RIKEN cDNA 1700028J19 gene | 0.83 | 0.81 | 1.37E-04 | 5.88E-03 |
| **Col4a4** | collagen, type IV, alpha 4 | 0.63 | 0.81 | 1.35E-05 | 1.43E-03 |
| **Gpc5** | glypican 5 | 0.85 | 0.81 | 4.35E-04 | 1.24E-02 |
| **D10Jhu81e** | DNA segment, Chr 10, Johns Hopkins University 81 expressed | 0.70 | 0.81 | 6.95E-04 | 1.66E-02 |
| **Slc30a5** | solute carrier family 30 (zinc transporter), member 5 | 0.89 | 0.81 | 7.80E-04 | 1.79E-02 |
| **Pepd** | peptidase D | 0.87 | 0.81 | 2.56E-03 | 3.81E-02 |
| **Rorb** | RAR-related orphan receptor beta | 0.54 | 0.81 | 1.15E-03 | 2.28E-02 |
| **Sh2d4a** | SH2 domain containing 4A | 0.95 | 0.81 | 7.86E-04 | 1.79E-02 |
| **Lrba** | LPS-responsive beige-like anchor | 0.82 | 0.81 | 2.86E-05 | 2.15E-03 |
| **Fam163a** | family with sequence similarity 163, member A | 0.90 | 0.81 | 2.80E-03 | 4.04E-02 |
| **Pdzrn4** | PDZ domain containing RING finger 4 | 0.67 | 0.81 | 8.68E-05 | 4.32E-03 |
| **Smim20** | small integral membrane protein 20 | 0.91 | 0.81 | 3.55E-04 | 1.11E-02 |
| **Smim24** | small integral membrane protein 24 | 0.63 | 0.81 | 9.29E-04 | 2.00E-02 |
| **Atp6v1h** | ATPase, H+ transporting, lysosomal V1 subunit H | 0.98 | 0.81 | 9.90E-05 | 4.74E-03 |
| **Tspan3** | tetraspanin 3 | 0.89 | 0.81 | 4.86E-04 | 1.32E-02 |
| **A1cf** | APOBEC1 complementation factor | 0.74 | 0.81 | 8.59E-04 | 1.89E-02 |
| **Smim5** | small integral membrane protein 5 | 0.89 | 0.81 | 8.56E-05 | 4.29E-03 |
| **P3h2** | prolyl 3-hydroxylase 2 | 0.69 | 0.81 | 4.70E-04 | 1.30E-02 |
| **Vldlr** | very low density lipoprotein receptor | 0.78 | 0.81 | 5.38E-04 | 1.40E-02 |
| **Bcar3** | breast cancer anti-estrogen resistance 3 | 0.64 | 0.80 | 1.86E-05 | 1.69E-03 |
| **Pfn4** | profilin family, member 4 | 0.91 | 0.80 | 1.55E-03 | 2.79E-02 |
| **Klk1** | kallikrein 1 | 0.81 | 0.80 | 8.36E-04 | 1.86E-02 |
| **Gphn** | gephyrin | 0.77 | 0.80 | 2.96E-04 | 9.83E-03 |
| **Pccb** | propionyl Coenzyme A carboxylase, beta polypeptide | 0.68 | 0.80 | 4.77E-04 | 1.31E-02 |
| **Tbc1d32** | TBC1 domain family, member 32 | 0.40 | 0.80 | 6.12E-05 | 3.50E-03 |
| **Slc47a1** | solute carrier family 47, member 1 | 0.75 | 0.80 | 4.58E-04 | 1.28E-02 |
| **Clcnkb** | chloride channel Kb | 0.85 | 0.80 | 3.97E-03 | 4.98E-02 |
| **Dio1** | deiodinase, iodothyronine, type I | 0.57 | 0.80 | 2.15E-03 | 3.46E-02 |
| **Aif1l** | allograft inflammatory factor 1-like | 0.91 | 0.80 | 2.76E-05 | 2.14E-03 |
| **Taco1** | translational activator of mitochondrially encoded cytochrome c oxidase I | 0.43 | 0.80 | 7.91E-04 | 1.79E-02 |
| **Dst** | dystonin | 0.86 | 0.80 | 5.82E-06 | 8.65E-04 |
| **Sim2** | single-minded homolog 2 (Drosophila) | 0.52 | 0.80 | 1.04E-03 | 2.15E-02 |
| **Tmem117** | transmembrane protein 117 | 0.97 | 0.80 | 5.10E-04 | 1.36E-02 |
| **Esrrg** | estrogen-related receptor gamma | 0.65 | 0.80 | 1.69E-05 | 1.61E-03 |
| **Pigg** | phosphatidylinositol glycan anchor biosynthesis, class G | 0.70 | 0.80 | 4.02E-04 | 1.18E-02 |

| | | | | | |
|---|---|---|---|---|---|
| **Lyrm4** | LYR motif containing 4 | 0.97 | 0.80 | 1.10E-04 | 5.05E-03 |
| **Rab11fip4** | RAB11 family interacting protein 4 (class II) | 0.79 | 0.80 | 1.40E-04 | 5.94E-03 |
| **Amacr** | alpha-methylacyl-CoA racemase | 0.59 | 0.80 | 1.67E-03 | 2.93E-02 |
| **Eml5** | echinoderm microtubule associated protein like 5 | 0.45 | 0.80 | 5.35E-04 | 1.40E-02 |
| **Lrrk2** | leucine-rich repeat kinase 2 | 0.39 | 0.80 | 1.56E-04 | 6.31E-03 |
| **Amy1** | amylase 1, salivary | 0.93 | 0.79 | 2.37E-03 | 3.65E-02 |
| **Efhd1** | EF hand domain containing 1 | 0.91 | 0.79 | 4.26E-04 | 1.23E-02 |
| **Gm15327** | predicted gene 15327 | 0.92 | 0.79 | 2.25E-03 | 3.56E-02 |
| **Pfn2** | profilin 2 | 0.94 | 0.79 | 9.56E-06 | 1.18E-03 |
| **Oxct1** | 3-oxoacid CoA transferase 1 | 0.72 | 0.79 | 8.35E-05 | 4.22E-03 |
| **Fabp3** | fatty acid binding protein 3, muscle and heart | 0.69 | 0.79 | 1.95E-04 | 7.35E-03 |
| **Fam189a2** | family with sequence similarity 189, member A2 | 0.87 | 0.79 | 3.53E-04 | 1.11E-02 |
| **Acss2** | acyl-CoA synthetase short-chain family member 2 | 0.54 | 0.79 | 3.90E-04 | 1.16E-02 |
| **Magix** | MAGI family member, X-linked | 0.40 | 0.79 | 4.46E-05 | 2.92E-03 |
| **Gm9938** | predicted gene 9938 | 0.57 | 0.79 | 5.18E-04 | 1.37E-02 |
| **Tmem163** | transmembrane protein 163 | 0.89 | 0.79 | 2.29E-04 | 8.20E-03 |
| **Pcca** | propionyl-Coenzyme A carboxylase, alpha polypeptide | 0.69 | 0.79 | 7.16E-04 | 1.69E-02 |
| **Plscr4** | phospholipid scramblase 4 | 0.98 | 0.79 | 1.05E-03 | 2.16E-02 |
| **Mep1b** | meprin 1 beta | 0.45 | 0.78 | 2.75E-03 | 4.00E-02 |
| **F5** | coagulation factor V | 0.81 | 0.78 | 5.17E-04 | 1.37E-02 |
| **Pclo** | piccolo (presynaptic cytomatrix protein) | 0.48 | 0.78 | 3.82E-06 | 7.22E-04 |
| **Slc22a12** | solute carrier family 22 (organic anion/cation transporter), member 12 | 0.59 | 0.78 | 3.21E-04 | 1.04E-02 |
| **Cyfip2** | cytoplasmic FMR1 interacting protein 2 | 0.66 | 0.78 | 3.04E-05 | 2.25E-03 |
| **Slc25a35** | solute carrier family 25, member 35 | 0.96 | 0.78 | 1.03E-03 | 2.14E-02 |
| **Mtfp1** | mitochondrial fission process 1 | 0.89 | 0.77 | 2.56E-04 | 8.86E-03 |
| **Larp1b** | La ribonucleoprotein domain family, member 1B | 0.57 | 0.77 | 4.78E-05 | 2.98E-03 |
| **Hhatl** | hedgehog acyltransferase-like | 0.95 | 0.77 | 5.86E-06 | 8.65E-04 |
| **Gm19537** | predicted gene, 19537 | 0.98 | 0.77 | 2.88E-03 | 4.10E-02 |
| **Myo3b** | myosin IIIB | 0.80 | 0.77 | 7.42E-05 | 3.97E-03 |
| **Tnfsf15** | tumor necrosis factor (ligand) superfamily, member 15 | 0.53 | 0.77 | 6.35E-04 | 1.56E-02 |
| **Rtn4ip1** | reticulon 4 interacting protein 1 | 0.40 | 0.77 | 1.44E-03 | 2.65E-02 |
| **Slc43a2** | solute carrier family 43, member 2 | 0.68 | 0.77 | 8.50E-04 | 1.88E-02 |
| **Wnk4** | WNK lysine deficient protein kinase 4 | 0.66 | 0.77 | 3.21E-05 | 2.31E-03 |
| **Lrrc3** | leucine rich repeat containing 3 | 0.62 | 0.77 | 1.42E-03 | 2.64E-02 |
| **Ifit1bl2** | interferon induced protein with tetratricopeptide repeats 1B like 2 | 0.44 | 0.77 | 8.90E-04 | 1.94E-02 |
| **Papss1** | 3'-phosphoadenosine 5'-phosphosulfate synthase 1 | 0.87 | 0.77 | 2.77E-05 | 2.14E-03 |
| **Itpr2** | inositol 1,4,5-triphosphate receptor 2 | 0.60 | 0.77 | 3.41E-05 | 2.43E-03 |
| **Ppp1r1a** | protein phosphatase 1, regulatory (inhibitor) subunit 1A | 0.82 | 0.77 | 7.77E-05 | 4.10E-03 |
| **Trabd2b** | TraB domain containing 2B | 0.45 | 0.77 | 7.10E-05 | 3.87E-03 |
| **Cwh43** | cell wall biogenesis 43 C-terminal homolog (S. cerevisiae) | 0.55 | 0.77 | 2.52E-03 | 3.79E-02 |
| **Htr1b** | 5-hydroxytryptamine (serotonin) receptor 1B | 0.97 | 0.77 | 2.59E-05 | 2.06E-03 |
| **Vstm2a** | V-set and transmembrane domain containing 2A | 0.59 | 0.76 | 8.90E-04 | 1.94E-02 |
| **Atp6v0e2** | ATPase, H+ transporting, lysosomal V0 subunit E2 | 0.47 | 0.76 | 9.03E-04 | 1.96E-02 |
| **Srgap3** | SLIT-ROBO Rho GTPase activating protein 3 | 0.57 | 0.76 | 8.84E-04 | 1.94E-02 |
| **Afmid** | arylformamidase | 0.46 | 0.76 | 7.70E-04 | 1.77E-02 |
| **Car12** | carbonic anyhydrase 12 | 0.69 | 0.76 | 6.24E-04 | 1.55E-02 |
| **Pfkm** | phosphofructokinase, muscle | 0.86 | 0.76 | 7.01E-05 | 3.86E-03 |
| **Afm** | afamin | 0.82 | 0.76 | 2.56E-03 | 3.81E-02 |
| **Tarm1** | T cell-interacting, activating receptor on myeloid cells 1 | 0.95 | 0.76 | 3.01E-06 | 6.32E-04 |
| **Nrip3** | nuclear receptor interacting protein 3 | 0.61 | 0.75 | 1.11E-03 | 2.24E-02 |
| **Sgpp1** | sphingosine-1-phosphate phosphatase 1 | 0.92 | 0.75 | 4.26E-05 | 2.81E-03 |
| **Kcnj1** | potassium inwardly-rectifying channel, subfamily J, member 1 | 0.84 | 0.75 | 3.12E-05 | 2.29E-03 |
| **Slc9a2** | solute carrier family 9 (sodium/hydrogen exchanger), member 2 | 0.95 | 0.75 | 7.85E-05 | 4.11E-03 |
| **Prcp** | prolylcarboxypeptidase (angiotensinase C) | 0.89 | 0.75 | 2.22E-05 | 1.87E-03 |

| | | | | | |
|---|---|---|---|---|---|
| **Slc13a3** | solute carrier family 13 (sodium-dependent dicarboxylate transporter), member 3 | 0.58 | 0.75 | 3.82E-07 | 2.14E-04 |
| **Slc22a4** | solute carrier family 22 (organic cation transporter), member 4 | 0.50 | 0.75 | 3.16E-04 | 1.03E-02 |
| **Idi1** | isopentenyl-diphosphate delta isomerase | 0.98 | 0.74 | 2.33E-04 | 8.31E-03 |
| **Scn4b** | sodium channel, type IV, beta | 0.89 | 0.74 | 2.20E-03 | 3.51E-02 |
| **Cpm** | carboxypeptidase M | 0.99 | 0.74 | 4.76E-04 | 1.31E-02 |
| **Chst9** | carbohydrate (N-acetylgalactosamine 4-0) sulfotransferase 9 | 0.71 | 0.74 | 5.11E-05 | 3.13E-03 |
| **4930556J24Rik** | RIKEN cDNA 4930556J24 gene | 0.90 | 0.74 | 5.02E-04 | 1.35E-02 |
| **P2rx5** | purinergic receptor P2X, ligand-gated ion channel, 5 | 0.79 | 0.74 | 4.47E-05 | 2.92E-03 |
| **Acmsd** | amino carboxymuconate semialdehyde decarboxylase | 0.41 | 0.74 | 2.25E-03 | 3.56E-02 |
| **Tmem64** | transmembrane protein 64 | 0.73 | 0.74 | 5.62E-05 | 3.32E-03 |
| **Defb1** | defensin beta 1 | 0.91 | 0.73 | 2.58E-05 | 2.06E-03 |
| **Impa2** | inositol (myo)-1(or 4)-monophosphatase 2 | 0.97 | 0.73 | 1.23E-05 | 1.35E-03 |
| **Prps2** | phosphoribosyl pyrophosphate synthetase 2 | 0.94 | 0.73 | 5.02E-05 | 3.09E-03 |
| **Cyp17a1** | cytochrome P450, family 17, subfamily a, polypeptide 1 | 0.68 | 0.73 | 3.92E-05 | 2.66E-03 |
| **Syn2** | synapsin II | 0.96 | 0.73 | 2.33E-07 | 1.86E-04 |
| **Asb9** | ankyrin repeat and SOCS box-containing 9 | 0.55 | 0.73 | 5.10E-04 | 1.36E-02 |
| **Igfbp5** | insulin-like growth factor binding protein 5 | 0.28 | 0.73 | 2.34E-03 | 3.63E-02 |
| **Tmem72** | transmembrane protein 72 | 0.94 | 0.72 | 1.12E-06 | 3.54E-04 |
| **Kl** | klotho | 0.82 | 0.72 | 5.35E-06 | 8.39E-04 |
| **Cpne4** | copine IV | 0.39 | 0.72 | 2.09E-05 | 1.80E-03 |
| **Slc22a22** | solute carrier family 22 (organic cation transporter), member 22 | 0.47 | 0.71 | 1.17E-05 | 1.31E-03 |
| **Clic6** | chloride intracellular channel 6 | 0.63 | 0.71 | 2.58E-04 | 8.86E-03 |
| **Fam81a** | family with sequence similarity 81, member A | 0.94 | 0.71 | 3.22E-04 | 1.04E-02 |
| **D630024D03Rik** | RIKEN cDNA D630024D03 gene | 0.95 | 0.71 | 1.60E-04 | 6.43E-03 |
| **Cldn10** | claudin 10 | 0.76 | 0.71 | 8.02E-06 | 1.05E-03 |
| **Asb15** | ankyrin repeat and SOCS box-containing 15 | 0.88 | 0.71 | 3.14E-04 | 1.02E-02 |
| **Slc6a17** | solute carrier family 6 (neurotransmitter transporter), member 17 | 0.86 | 0.71 | 4.60E-06 | 7.98E-04 |
| **Slc9a3** | solute carrier family 9 (sodium/hydrogen exchanger), member 3 | 0.79 | 0.71 | 6.85E-04 | 1.64E-02 |
| **Erc2** | ELKS/RAB6-interacting/CAST family member 2 | 0.54 | 0.70 | 1.18E-06 | 3.55E-04 |
| **Tmem61** | transmembrane protein 61 | 0.87 | 0.70 | 3.91E-07 | 2.14E-04 |
| **Sv2a** | synaptic vesicle glycoprotein 2 a | 0.87 | 0.70 | 5.89E-06 | 8.65E-04 |
| **Nnt** | nicotinamide nucleotide transhydrogenase | 0.69 | 0.70 | 1.95E-03 | 3.24E-02 |
| **Slc8a1** | solute carrier family 8 (sodium/calcium exchanger), member 1 | 0.98 | 0.69 | 1.21E-07 | 1.15E-04 |
| **Fmo1** | flavin containing monooxygenase 1 | 0.47 | 0.69 | 1.72E-05 | 1.62E-03 |
| **Sucnr1** | succinate receptor 1 | 0.23 | 0.69 | 2.99E-03 | 4.19E-02 |
| **Calb1** | calbindin 1 | 0.87 | 0.69 | 1.22E-06 | 3.59E-04 |
| **Slc7a13** | solute carrier family 7, (cationic amino acid transporter, y+ system) member 13 | 0.44 | 0.69 | 1.39E-05 | 1.46E-03 |
| **Mboat2** | membrane bound O-acyltransferase domain containing 2 | 0.80 | 0.68 | 1.16E-05 | 1.30E-03 |
| **Gp2** | glycoprotein 2 (zymogen granule membrane) | 0.66 | 0.68 | 4.88E-06 | 8.07E-04 |
| **Klk1b21** | kallikrein 1-related peptidase b21 | 0.83 | 0.67 | 5.89E-05 | 3.41E-03 |
| **Id2** | inhibitor of DNA binding 2 | 0.84 | 0.67 | 3.59E-04 | 1.11E-02 |
| **Cpt1b** | carnitine palmitoyltransferase 1b, muscle | 0.46 | 0.67 | 6.23E-06 | 8.88E-04 |
| **Slc2a13** | solute carrier family 2 (facilitated glucose transporter), member 13 | 0.57 | 0.67 | 2.03E-06 | 4.82E-04 |
| **Rnf152** | ring finger protein 152 | 0.61 | 0.67 | 5.39E-04 | 1.40E-02 |
| **Tmem86a** | transmembrane protein 86A | 0.56 | 0.67 | 6.61E-07 | 2.75E-04 |
| **Klhdc8a** | kelch domain containing 8A | 0.74 | 0.66 | 2.26E-06 | 5.17E-04 |
| **Pde1a** | phosphodiesterase 1A, calmodulin-dependent | 0.56 | 0.66 | 2.14E-07 | 1.78E-04 |
| **Slc34a3** | solute carrier family 34 (sodium phosphate), member 3 | 0.66 | 0.66 | 4.77E-04 | 1.31E-02 |
| **Slitrk6** | SLIT and NTRK-like family, member 6 | 0.44 | 0.65 | 3.93E-03 | 4.95E-02 |
| **Casr** | calcium-sensing receptor | 0.91 | 0.65 | 9.79E-06 | 1.19E-03 |
| **Nepn** | nephrocan | 0.54 | 0.65 | 2.85E-03 | 4.08E-02 |
| **Pygl** | liver glycogen phosphorylase | 0.77 | 0.65 | 3.08E-06 | 6.35E-04 |
| **Ppp1r1b** | protein phosphatase 1, regulatory (inhibitor) subunit 1B | 0.98 | 0.64 | 4.79E-07 | 2.40E-04 |
| **Mfsd4** | major facilitator superfamily domain containing 4 | 0.53 | 0.64 | 1.32E-06 | 3.67E-04 |

| | | | | | |
|---|---|---|---|---|---|
| **Pvalb** | parvalbumin | 0.38 | 0.64 | 1.57E-06 | 4.03E-04 |
| **Pla1a** | phospholipase A1 member A | 0.82 | 0.63 | 1.22E-06 | 3.59E-04 |
| **Clcnka** | chloride channel Ka | 0.92 | 0.63 | 3.45E-05 | 2.46E-03 |
| **Nccrp1** | non-specific cytotoxic cell receptor protein 1 homolog (zebrafish) | 0.57 | 0.63 | 4.15E-06 | 7.61E-04 |
| **5930403L14Rik** | RIKEN cDNA 5930403L14 gene | 0.82 | 0.63 | 4.72E-05 | 2.97E-03 |
| **Bckdhb** | branched chain ketoacid dehydrogenase E1, beta polypeptide | 0.60 | 0.62 | 5.55E-07 | 2.59E-04 |
| **Atp6v1c2** | ATPase, H+ transporting, lysosomal V1 subunit C2 | 0.86 | 0.61 | 3.60E-06 | 6.97E-04 |
| **Prox1** | prospero homeobox 1 | 0.66 | 0.61 | 7.43E-05 | 3.97E-03 |
| **Perm1** | PPARGC1 and ESRR induced regulator, muscle 1 | 0.77 | 0.61 | 1.44E-06 | 3.82E-04 |
| **Slc4a1** | solute carrier family 4 (anion exchanger), member 1 | 0.86 | 0.60 | 1.02E-07 | 1.15E-04 |
| **BC023202** | cDNA sequence BC023202 | 0.86 | 0.60 | 2.38E-07 | 1.86E-04 |
| **Cacnb4** | calcium channel, voltage-dependent, beta 4 subunit | 0.77 | 0.59 | 1.22E-07 | 1.15E-04 |
| **Fabp7** | fatty acid binding protein 7, brain | 0.80 | 0.58 | 9.59E-05 | 4.66E-03 |
| **Fa2h** | fatty acid 2-hydroxylase | 0.76 | 0.58 | 1.11E-06 | 3.54E-04 |
| **Slc16a7** | solute carrier family 16 (monocarboxylic acid transporters), member 7 | 0.76 | 0.57 | 8.72E-08 | 1.12E-04 |
| **Gys2** | glycogen synthase 2 | 0.64 | 0.57 | 1.68E-03 | 2.93E-02 |
| **Id1** | inhibitor of DNA binding 1 | 0.81 | 0.56 | 1.27E-06 | 3.67E-04 |
| **2410003L11Rik** | RIKEN cDNA 2410003L11 gene | 0.45 | 0.53 | 4.99E-08 | 8.30E-05 |
| **Kcnt1** | potassium channel, subfamily T, member 1 | 0.52 | 0.52 | 1.29E-08 | 4.01E-05 |
| **Egf** | epidermal growth factor | 0.49 | 0.51 | 1.52E-07 | 1.36E-04 |
| **Ugt2b38** | UDP glucuronosyltransferase 2 family, polypeptide B38 | 0.59 | 0.47 | 4.95E-04 | 1.34E-02 |
| **Pcsk6** | proprotein convertase subtilisin/kexin type 6 | 0.65 | 0.47 | 3.56E-09 | 1.48E-05 |
| **Slc26a4** | solute carrier family 26, member 4 | 0.65 | 0.46 | 9.69E-05 | 4.69E-03 |
| **Lrrc66** | leucine rich repeat containing 66 | 0.45 | 0.45 | 4.56E-06 | 7.98E-04 |
| **Wfdc15b** | WAP four-disulfide core domain 15B | 0.64 | 0.37 | 8.97E-08 | 1.12E-04 |
| **Gpx6** | glutathione peroxidase 6 | 0.67 | 0.34 | 5.57E-05 | 3.30E-03 |

**Supplementary Table 4: GSEA pathways (biological processes) upregulated for the 1,262 common regulated genes between *Glis2* $^{lacZ/lacZ}$ and *Lkb1* $^{\Delta Tub}$ mouse kidney datasets.**
ES: enrichment score; NES: normalized enrichment score; NOM: nominal; FDR: false discovery rate. Gene sets with FDR q-value < 0.05 are considered as significant.

| NAME | SIZE | ES | NES | NOM p-val | FDR q-val |
|---|---|---|---|---|---|
| GO_HUMORAL_IMMUNE_RESPONSE | 26 | 0.66621095 | 2.067855 | 0.0 | 0.007852228 |
| GO_EXTRACELLULAR_STRUCTURE_ORGANIZATION | 87 | 0.56073487 | 1.9892554 | 0.0 | 0.021128828 |
| GO_DEFENSE_RESPONSE | 195 | 0.5215497 | 1.9684628 | 0.0 | 0.020643592 |
| GO_DEFENSE_RESPONSE_TO_OTHER_ORGANISM | 129 | 0.53645766 | 1.9674906 | 0.0 | 0.015482694 |
| GO_REGULATION_OF_IMMUNE_RESPONSE | 129 | 0.53312606 | 1.9510323 | 0.0 | 0.018089907 |
| GO_CELL_KILLING | 17 | 0.6962832 | 1.9510143 | 0.0 | 0.015074923 |
| GO_REGULATION_OF_IMMUNE_EFFECTOR_PROCESS | 48 | 0.5731404 | 1.9456544 | 0.0 | 0.014187577 |
| GO_INNATE_IMMUNE_RESPONSE | 113 | 0.53209037 | 1.9413596 | 0.0 | 0.013512896 |
| GO_LYMPHOCYTE_MEDIATED_IMMUNITY | 29 | 0.61385214 | 1.9365278 | 0.0 | 0.012996295 |
| GO_SKELETAL_SYSTEM_DEVELOPMENT | 54 | 0.55610216 | 1.8967383 | 0.0 | 0.02074488 |
| GO_CELL_CHEMOTAXIS | 46 | 0.5634526 | 1.8960383 | 0.0 | 0.019126428 |
| GO_INFLAMMATORY_RESPONSE | 112 | 0.5211736 | 1.8955723 | 0.0 | 0.017858213 |
| GO_LEUKOCYTE_MIGRATION | 71 | 0.5269233 | 1.8658069 | 0.0 | 0.02661831 |
| GO_RESPONSE_TO_BIOTIC_STIMULUS | 173 | 0.49664843 | 1.8598448 | 0.0 | 0.027521139 |
| GO_ACTIVATION_OF_IMMUNE_RESPONSE | 78 | 0.5131366 | 1.8430433 | 0.0 | 0.033678357 |
| GO_CARTILAGE_DEVELOPMENT | 29 | 0.586406 | 1.8384608 | 0.0 | 0.034340322 |
| GO_MULTI_MULTICELLULAR_ORGANISM_PROCESS | 24 | 0.6068613 | 1.8353858 | 0.0 | 0.033476755 |
| GO_RESPONSE_TO_CYTOKINE | 159 | 0.49589044 | 1.8338768 | 0.0 | 0.032601755 |
| GO_CYTOKINE_MEDIATED_SIGNALING_PATHWAY | 113 | 0.49929243 | 1.8294133 | 0.0 | 0.03295453 |
| GO_POSITIVE_REGULATION_OF_IMMUNE_RESPONSE | 97 | 0.50678205 | 1.8249083 | 0.0 | 0.034305975 |
| GO_POSITIVE_REGULATION_OF_IMMUNE_SYSTEM_PROCESS | 137 | 0.49871063 | 1.8202782 | 0.0 | 0.034637894 |
| GO_ADAPTIVE_IMMUNE_RESPONSE | 58 | 0.5317506 | 1.8190666 | 0.0 | 0.033375505 |
| GO_COLLAGEN_FIBRIL_ORGANIZATION | 16 | 0.67008376 | 1.818461 | 0.0011587485 | 0.032094307 |
| GO_LEUKOCYTE_PROLIFERATION | 40 | 0.54940796 | 1.8114184 | 0.0 | 0.033988744 |
| GO_POSITIVE_REGULATION_OF_RESPONSE_TO_EXTERNAL_STIMULUS | 82 | 0.5072864 | 1.808564 | 0.0 | 0.033886746 |
| GO_SKELETAL_SYSTEM_MORPHOGENESIS | 20 | 0.61214364 | 1.8078674 | 0.0 | 0.032847222 |
| GO_LEUKOCYTE_CHEMOTAXIS | 37 | 0.55825764 | 1.8072802 | 0.0 | 0.031923257 |
| GO_ACUTE_INFLAMMATORY_RESPONSE | 15 | 0.6558295 | 1.8042324 | 0.0035799523 | 0.032571327 |
| GO_ADAPTIVE_IMMUNE_RESPONSE_BASED_ON_SOMATIC_RECOMBINATION_OF_IMMUNE_RECEPTORS_BUILT_FROM _IMMUNOGLOBULIN_SUPERFAMILY_DOMAINS | 37 | 0.5503824 | 1.8034061 | 0.0010615712 | 0.031720538 |
| GO_LEUKOCYTE_MEDIATED_IMMUNITY | 92 | 0.5024513 | 1.8023871 | 0.0 | 0.031187467 |
| GO_REGULATION_OF_DEFENSE_RESPONSE | 85 | 0.49317744 | 1.7970753 | 0.0010121458 | 0.032367725 |
| GO_NEGATIVE_REGULATION_OF_IMMUNE_RESPONSE | 19 | 0.61410946 | 1.789299 | 0.0022909509 | 0.035011288 |
| GO_REGULATION_OF_IMMUNE_SYSTEM_PROCESS | 183 | 0.4771025 | 1.7844775 | 0.0 | 0.03585723 |
| GO_REGULATION_OF_RESPONSE_TO_BIOTIC_STIMULUS | 54 | 0.51426196 | 1.7839221 | 0.0 | 0.035207093 |
| GO_REGULATION_OF_ADAPTIVE_IMMUNE_RESPONSE | 25 | 0.582107 | 1.78363 | 0.0010917031 | 0.034397963 |
| GO_POSITIVE_REGULATION_OF_INFLAMMATORY_RESPONSE | 20 | 0.60609365 | 1.7791643 | 0.0033860046 | 0.035054155 |
| GO_POSITIVE_REGULATION_OF_LEUKOCYTE_MIGRATION | 25 | 0.580036 | 1.7775105 | 0.0022123894 | 0.034878463 |
| GO_NEUTROPHIL_MIGRATION | 22 | 0.6017832 | 1.7745414 | 0.0033076075 | 0.035256065 |
| GO_TOLL_LIKE_RECEPTOR_SIGNALING_PATHWAY | 24 | 0.57859284 | 1.7689314 | 0.007633588 | 0.036592204 |
| GO_POSITIVE_REGULATION_OF_RESPONSE_TO_BIOTIC_STIMULUS | 43 | 0.53498745 | 1.7678666 | 0.0 | 0.036120072 |
| GO_REGULATION_OF_LEUKOCYTE_MEDIATED_IMMUNITY | 25 | 0.57391727 | 1.7658349 | 0.0032822757 | 0.0360085 |
| GO_POSITIVE_REGULATION_OF_MULTI_ORGANISM_PROCESS | 53 | 0.51404935 | 1.765294 | 0.0010373445 | 0.035454188 |
| GO_POSITIVE_REGULATION_OF_CELL_ACTIVATION | 50 | 0.5180487 | 1.7649782 | 0.0 | 0.034721445 |

| | | | | | |
|---|---|---|---|---|---|
| GO_FEMALE_PREGNANCY | 19 | 0.61670524 | 1.7647437 | 0.0011148272 | 0.03402204 |
| GO_POSITIVE_REGULATION_OF_DEFENSE_RESPONSE | 57 | 0.51009816 | 1.7609339 | 0.0 | 0.034793675 |
| GO_REGULATION_OF_RESPONSE_TO_EXTERNAL_STIMULUS | 129 | 0.4800708 | 1.7582253 | 0.0 | 0.034976516 |
| GO_BONE_DEVELOPMENT | 26 | 0.5693891 | 1.7563801 | 0.001112347 | 0.03498501 |
| GO_CONNECTIVE_TISSUE_DEVELOPMENT | 38 | 0.5298457 | 1.7549244 | 0.0031512605 | 0.034971733 |
| GO_NEGATIVE_REGULATION_OF_CYTOKINE_PRODUCTION | 40 | 0.5261624 | 1.7531017 | 0.0020768433 | 0.035219166 |
| GO_PATTERN_RECOGNITION_RECEPTOR_SIGNALING_PATHWAY | 28 | 0.5538741 | 1.7465814 | 0.005446623 | 0.03746395 |
| GO_ACTIVATION_OF_INNATE_IMMUNE_RESPONSE | 39 | 0.5296795 | 1.7433105 | 0.001059322 | 0.038021103 |
| GO_POSITIVE_REGULATION_OF_LEUKOCYTE_CHEMOTAXIS | 18 | 0.6042895 | 1.7415867 | 0.008045977 | 0.037949666 |
| GO_T_CELL_PROLIFERATION | 29 | 0.5589455 | 1.7402835 | 0.003215434 | 0.037826084 |
| GO_RESPONSE_TO_CHEMOKINE | 18 | 0.6078133 | 1.7356441 | 0.0045610033 | 0.039348092 |
| GO_BIOLOGICAL_ADHESION | 200 | 0.46077934 | 1.7309443 | 0.0 | 0.040866535 |
| GO_NEGATIVE_REGULATION_OF_CELL_ADHESION | 43 | 0.5121235 | 1.7298613 | 0.0041841003 | 0.04050643 |
| GO_POSITIVE_REGULATION_OF_TUMOR_NECROSIS_FACTOR_SUPERFAMILY_CYTOKINE_PRODUCTION | 24 | 0.5735702 | 1.7259356 | 0.0011086474 | 0.041778497 |
| GO_CYTOKINE_PRODUCTION | 103 | 0.4758575 | 1.7213104 | 0.0 | 0.04344863 |
| GO_IMMUNE_EFFECTOR_PROCESS | 139 | 0.46415687 | 1.7182915 | 0.0 | 0.04428009 |
| GO_REGULATION_OF_CELL_ADHESION | 107 | 0.4730925 | 1.7121757 | 0.0 | 0.04665549 |
| GO_REGULATION_OF_INFLAMMATORY_RESPONSE | 47 | 0.50560653 | 1.7119755 | 0.0010224949 | 0.04601984 |
| GO_REGULATION_OF_CELL_SUBSTRATE_ADHESION | 35 | 0.5278153 | 1.7089018 | 0.0010559662 | 0.04695909 |
| GO_INTERLEUKIN_2_PRODUCTION | 17 | 0.60971034 | 1.7086767 | 0.0034562212 | 0.046354588 |
| GO_REGULATION_OF_MULTI_ORGANISM_PROCESS | 77 | 0.47976625 | 1.7061187 | 0.0 | 0.047121096 |
| GO_POSITIVE_REGULATION_OF_CYTOKINE_PRODUCTION | 72 | 0.4829965 | 1.6966208 | 0.0010131713 | 0.05211432 |
| GO_INTERLEUKIN_6_PRODUCTION | 24 | 0.55946285 | 1.6958449 | 0.013172338 | 0.051755063 |
| GO_POSITIVE_REGULATION_OF_CHEMOTAXIS | 28 | 0.5427862 | 1.6953268 | 0.008695652 | 0.051217668 |
| GO_GRANULOCYTE_MIGRATION | 27 | 0.5499934 | 1.6951272 | 0.009771987 | 0.050608486 |
| GO_MYELOID_LEUKOCYTE_ACTIVATION | 86 | 0.47102153 | 1.6945473 | 0.0 | 0.05037353 |
| GO_TAXIS | 84 | 0.47158358 | 1.6903601 | 0.0 | 0.05237761 |
| GO_MYELOID_LEUKOCYTE_MIGRATION | 36 | 0.523802 | 1.68904 | 0.0020876827 | 0.0524286 |
| GO_REGULATION_OF_LEUKOCYTE_CHEMOTAXIS | 20 | 0.5853565 | 1.685618 | 0.011123471 | 0.053912792 |
| GO_CELLULAR_RESPONSE_TO_ACID_CHEMICAL | 26 | 0.55117327 | 1.6848837 | 0.004371585 | 0.05359102 |
| GO_REGULATION_OF_CHEMOTAXIS | 34 | 0.52116734 | 1.6844617 | 0.0053418805 | 0.053213175 |
| GO_POSITIVE_REGULATION_OF_CELL_ADHESION | 66 | 0.47422397 | 1.6799262 | 0.0 | 0.055349022 |
| GO_POSITIVE_REGULATION_OF_IMMUNE_EFFECTOR_PROCESS | 30 | 0.5340309 | 1.6795272 | 0.0087241 | 0.054879386 |
| GO_MYELOID_LEUKOCYTE_MEDIATED_IMMUNITY | 70 | 0.46890843 | 1.6759067 | 0.0 | 0.056271784 |
| GO_CELL_SUBSTRATE_ADHESION | 52 | 0.49191046 | 1.6710249 | 0.0020768433 | 0.05863848 |
| GO_REGULATION_OF_LEUKOCYTE_PROLIFERATION | 29 | 0.53055173 | 1.6681683 | 0.009667025 | 0.05973671 |
| GO_REGULATION_OF_CELL_ACTIVATION | 74 | 0.47413042 | 1.6662656 | 0.0 | 0.060228832 |
| GO_CELL_MATRIX_ADHESION | 32 | 0.52560127 | 1.6634274 | 0.0042283298 | 0.061413694 |
| GO_TUMOR_NECROSIS_FACTOR_SUPERFAMILY_CYTOKINE_PRODUCTION | 31 | 0.5276579 | 1.6591294 | 0.010810811 | 0.06366215 |
| GO_POSITIVE_REGULATION_OF_CELL_CELL_ADHESION | 46 | 0.49360088 | 1.6582862 | 0.0052246605 | 0.06358231 |
| GO_PHAGOCYTOSIS | 48 | 0.49339995 | 1.6570188 | 0.0 | 0.06351596 |
| GO_KERATINOCYTE_DIFFERENTIATION | 20 | 0.56312245 | 1.6565362 | 0.009090909 | 0.06310314 |
| GO_RESPONSE_TO_BACTERIUM | 83 | 0.4680798 | 1.6537218 | 0.0 | 0.06408375 |
| GO_POSITIVE_REGULATION_OF_CYTOKINE_BIOSYNTHETIC_PROCESS | 22 | 0.5465087 | 1.6530937 | 0.012127894 | 0.063788675 |
| GO_CELL_ACTIVATION | 172 | 0.4407701 | 1.6463608 | 0.0 | 0.06806724 |
| GO_POSITIVE_REGULATION_OF_LEUKOCYTE_PROLIFERATION | 22 | 0.5460353 | 1.6457249 | 0.0066740825 | 0.06770032 |
| GO_MULTI_ORGANISM_REPRODUCTIVE_PROCESS | 54 | 0.47716284 | 1.6443144 | 0.0040983604 | 0.06777745 |
| GO_ANATOMICAL_STRUCTURE_MATURATION | 17 | 0.5695059 | 1.6392533 | 0.018099548 | 0.0711711 |

| | | | | | |
|---|---|---|---|---|---|
| GO_POSITIVE_REGULATION_OF_PHAGOCYTOSIS | 15 | 0.59048617 | 1.6336101 | 0.021965317 | 0.074691184 |
| GO_CHEMOKINE_PRODUCTION | 18 | 0.5655363 | 1.6316646 | 0.01592719 | 0.0753137 |
| GO_PROTEIN_MATURATION | 33 | 0.50356776 | 1.6297717 | 0.012793177 | 0.07596485 |
| GO_REGULATION_OF_RESPONSE_TO_STRESS | 139 | 0.43919393 | 1.6257648 | 0.0 | 0.07831109 |
| GO_NEGATIVE_REGULATION_OF_RESPONSE_TO_WOUNDING | 15 | 0.59773076 | 1.6235441 | 0.014527845 | 0.07947012 |
| GO_DEFENSE_RESPONSE_TO_BACTERIUM | 25 | 0.52566946 | 1.6233519 | 0.008752735 | 0.078761935 |
| GO_COLLAGEN_METABOLIC_PROCESS | 26 | 0.52710205 | 1.6207104 | 0.015267176 | 0.08020781 |
| GO_NEGATIVE_REGULATION_OF_PEPTIDASE_ACTIVITY | 25 | 0.5233261 | 1.6182237 | 0.010787486 | 0.081292175 |
| GO_PROTEIN_PROCESSING | 32 | 0.5091953 | 1.6165082 | 0.008483564 | 0.08176706 |
| GO_MACROPHAGE_ACTIVATION | 19 | 0.5553411 | 1.6156855 | 0.016968327 | 0.08159013 |
| GO_MULTICELLULAR_ORGANISM_REPRODUCTION | 40 | 0.49033964 | 1.6144433 | 0.0063157897 | 0.08171646 |
| GO_POSITIVE_REGULATION_OF_VASCULATURE_DEVELOPMENT | 20 | 0.5586361 | 1.6122055 | 0.013729977 | 0.082793005 |
| GO_SUBSTRATE_ADHESION_DEPENDENT_CELL_SPREADING | 16 | 0.5871054 | 1.6105534 | 0.005800464 | 0.083310276 |
| GO_POSITIVE_REGULATION_OF_LEUKOCYTE_CELL_CELL_ADHESION | 36 | 0.4938203 | 1.6104839 | 0.009483667 | 0.082544945 |
| GO_NEGATIVE_REGULATION_OF_MULTICELLULAR_ORGANISMAL_PROCESS | 138 | 0.43732452 | 1.6069312 | 0.0 | 0.08509298 |
| GO_RESPONSE_TO_AMINO_ACID | 15 | 0.58569026 | 1.604765 | 0.014167651 | 0.08613587 |
| GO_IMMUNE_RESPONSE_REGULATING_SIGNALING_PATHWAY | 68 | 0.46478197 | 1.6034316 | 0.002038736 | 0.08642158 |
| GO_RESPONSE_TO_MOLECULE_OF_BACTERIAL_ORIGIN | 57 | 0.46331003 | 1.6024714 | 0.007106599 | 0.08633075 |
| GO_RESPONSE_TO_MECHANICAL_STIMULUS | 37 | 0.49400184 | 1.6014841 | 0.0063157897 | 0.08634043 |
| GO_POSITIVE_REGULATION_OF_T_CELL_PROLIFERATION | 21 | 0.5378327 | 1.5975326 | 0.011061947 | 0.08875235 |
| GO_NEGATIVE_REGULATION_OF_IMMUNE_SYSTEM_PROCESS | 52 | 0.46381512 | 1.5971988 | 0.004102564 | 0.08818745 |
| GO_REPRODUCTION | 83 | 0.4451463 | 1.5931427 | 0.0020120724 | 0.09127034 |
| GO_POSITIVE_REGULATION_OF_ADAPTIVE_IMMUNE_RESPONSE | 19 | 0.5568583 | 1.5924983 | 0.013714286 | 0.09114361 |
| GO_REGULATION_OF_LEUKOCYTE_MIGRATION | 39 | 0.48724595 | 1.5924565 | 0.01058201 | 0.090359606 |
| GO_RESPONSE_TO_LIPID | 105 | 0.4366926 | 1.5917192 | 0.0010050251 | 0.090233386 |
| GO_CELLULAR_RESPONSE_TO_LIPID | 68 | 0.45448402 | 1.5816286 | 0.0020283975 | 0.098762535 |
| GO_CELLULAR_RESPONSE_TO_OXYGEN_CONTAINING_COMPOUND | 128 | 0.42797247 | 1.5790076 | 0.0 | 0.10064164 |
| GO_EXTRACELLULAR_MATRIX_DISASSEMBLY | 19 | 0.5487866 | 1.5749276 | 0.025056947 | 0.10405863 |
| GO_REGULATION_OF_PHAGOCYTOSIS | 19 | 0.54410374 | 1.5747095 | 0.02283105 | 0.103379786 |
| GO_SKIN_DEVELOPMENT | 35 | 0.4897788 | 1.5744348 | 0.016949153 | 0.102728434 |
| GO_REGULATION_OF_LYMPHOCYTE_ACTIVATION | 51 | 0.45671117 | 1.5739751 | 0.014373717 | 0.10226435 |
| GO_LYMPHOCYTE_MIGRATION | 19 | 0.53837717 | 1.5706148 | 0.022522522 | 0.10441523 |
| GO_RESPONSE_TO_VITAMIN | 19 | 0.54564667 | 1.5690562 | 0.034168564 | 0.10518207 |
| GO_CELL_CELL_ADHESION | 119 | 0.42773744 | 1.5672095 | 0.001002004 | 0.10627466 |
| GO_REGULATION_OF_CELLULAR_COMPONENT_MOVEMENT | 137 | 0.42490056 | 1.566498 | 0.0 | 0.10616422 |
| GO_REGULATION_OF_RESPONSE_TO_WOUNDING | 37 | 0.4746701 | 1.5657206 | 0.015690377 | 0.10608716 |
| GO_REGULATION_OF_PEPTIDASE_ACTIVITY | 53 | 0.45907813 | 1.5648732 | 0.009268795 | 0.10617154 |
| GO_REGULATION_OF_VASCULATURE_DEVELOPMENT | 41 | 0.47340402 | 1.5618317 | 0.013415893 | 0.10870927 |
| GO_CELL_ACTIVATION_INVOLVED_IN_IMMUNE_RESPONSE | 91 | 0.4317183 | 1.5614451 | 0.0040241447 | 0.10823657 |
| GO_LEUKOCYTE_DIFFERENTIATION | 62 | 0.44570082 | 1.5595939 | 0.011282051 | 0.109271035 |
| GO_REGULATION_OF_PROTEOLYSIS | 69 | 0.4435201 | 1.5587044 | 0.0061791968 | 0.10933784 |
| GO_REGULATION_OF_CELL_CELL_ADHESION | 64 | 0.44537762 | 1.5579544 | 0.00814664 | 0.10949174 |
| GO_G_PROTEIN_COUPLED_RECEPTOR_SIGNALING_PATHWAY | 83 | 0.43541673 | 1.5569316 | 0.001010101 | 0.10964254 |
| GO_POSITIVE_REGULATION_OF_LEUKOCYTE_DIFFERENTIATION | 24 | 0.5099332 | 1.5568205 | 0.018660812 | 0.1089323 |
| GO_POSITIVE_REGULATION_OF_CELL_SUBSTRATE_ADHESION | 20 | 0.5296994 | 1.5563185 | 0.023014959 | 0.10868794 |
| GO_POSITIVE_REGULATION_OF_LYMPHOCYTE_DIFFERENTIATION | 15 | 0.56395906 | 1.5556462 | 0.020930232 | 0.108625874 |
| GO_LYMPHOCYTE_DIFFERENTIATION | 45 | 0.46117744 | 1.5542142 | 0.00935551 | 0.10926821 |
| GO_LYMPHOCYTE_ACTIVATION | 81 | 0.4407885 | 1.5536249 | 0.003033367 | 0.10911809 |

| | | | | | |
|---|---|---|---|---|---|
| GO_CELLULAR_RESPONSE_TO_BIOTIC_STIMULUS | 37 | 0.4769518 | 1.5533662 | 0.01590668 | 0.108675696 |
| GO_CARDIOVASCULAR_SYSTEM_DEVELOPMENT | 88 | 0.43283635 | 1.5524219 | 0.004052685 | 0.1088739 |
| GO_PATTERN_SPECIFICATION_PROCESS | 34 | 0.47995743 | 1.5471207 | 0.015974442 | 0.11360947 |
| GO_NEGATIVE_REGULATION_OF_PROTEOLYSIS | 29 | 0.49813306 | 1.543428 | 0.025695931 | 0.11670685 |
| GO_CELLULAR_COMPONENT_DISASSEMBLY | 40 | 0.461667 | 1.5390034 | 0.024819028 | 0.12091103 |
| GO_POSITIVE_REGULATION_OF_TRANSPORT | 120 | 0.42213592 | 1.5389471 | 0.0 | 0.12015159 |
| GO_REGULATION_OF_COAGULATION | 19 | 0.53064394 | 1.5375344 | 0.037078653 | 0.120795734 |
| GO_POSITIVE_REGULATION_OF_LOCOMOTION | 82 | 0.43005884 | 1.5312456 | 0.0070778565 | 0.126852 |
| GO_LEUKOCYTE_CELL_CELL_ADHESION | 49 | 0.44813713 | 1.5310514 | 0.016511869 | 0.1262006 |
| GO_CELL_MOTILITY | 194 | 0.4066234 | 1.5290997 | 0.0 | 0.12758851 |
| GO_REGULATION_OF_RESPONSE_TO_CYTOKINE_STIMULUS | 29 | 0.4898241 | 1.5280654 | 0.02967033 | 0.12795553 |
| GO_CYTOKINE_METABOLIC_PROCESS | 27 | 0.49478886 | 1.5279648 | 0.03504929 | 0.12723832 |
| GO_LYMPHOCYTE_ACTIVATION_INVOLVED_IN_IMMUNE_RESPONSE | 20 | 0.5224817 | 1.5247836 | 0.041426927 | 0.13046244 |
| GO_ENTRY_INTO_HOST | 18 | 0.53414816 | 1.5227982 | 0.04218928 | 0.13206291 |
| GO_BLOOD_VESSEL_MORPHOGENESIS | 75 | 0.43284664 | 1.5222986 | 0.0070921984 | 0.13184327 |
| GO_DEVELOPMENTAL_MATURATION | 27 | 0.48417336 | 1.5220497 | 0.035908595 | 0.13128483 |
| GO_RESPONSE_TO_WOUNDING | 100 | 0.41916972 | 1.52196 | 0.0020181634 | 0.13055038 |
| GO_POST_TRANSLATIONAL_PROTEIN_MODIFICATION | 34 | 0.47818583 | 1.519841 | 0.01910828 | 0.13202159 |
| GO_POSITIVE_REGULATION_OF_INTERLEUKIN_6_PRODUCTION | 17 | 0.55192876 | 1.5189724 | 0.032444958 | 0.13223672 |
| GO_REGULATION_OF_T_CELL_ACTIVATION | 40 | 0.4573137 | 1.518909 | 0.023983316 | 0.13146064 |
| GO_POSITIVE_REGULATION_OF_CELL_DEATH | 69 | 0.43574512 | 1.5183951 | 0.011235955 | 0.1311918 |
| GO_REGIONALIZATION | 26 | 0.49184313 | 1.5169146 | 0.020971302 | 0.13204394 |
| GO_INTERLEUKIN_8_PRODUCTION | 17 | 0.5349531 | 1.5163758 | 0.0314319 | 0.13186608 |
| GO_B_CELL_ACTIVATION | 24 | 0.50383604 | 1.5132011 | 0.0349345 | 0.13483834 |
| GO_CELLULAR_RESPONSE_TO_EXTERNAL_STIMULUS | 46 | 0.45006707 | 1.5119866 | 0.023858922 | 0.13541849 |
| GO_RESPONSE_TO_TUMOR_NECROSIS_FACTOR | 43 | 0.45905277 | 1.5092365 | 0.026123302 | 0.13770524 |
| GO_POSITIVE_REGULATION_OF_DEVELOPMENTAL_PROCESS | 138 | 0.4084279 | 1.5074544 | 0.004008016 | 0.1393088 |
| GO_REGULATION_OF_WOUND_HEALING | 33 | 0.47072712 | 1.505609 | 0.026766594 | 0.14079465 |
| GO_REGULATION_OF_CELL_DEATH | 138 | 0.40531185 | 1.502554 | 0.001003009 | 0.14347899 |
| GO_POSITIVE_REGULATION_OF_SECRETION | 49 | 0.44157356 | 1.4988748 | 0.024640657 | 0.14715272 |
| GO_POSITIVE_REGULATION_OF_LIPID_LOCALIZATION | 15 | 0.5423369 | 1.4976624 | 0.052873563 | 0.14776748 |
| GO_POSITIVE_REGULATION_OF_MULTICELLULAR_ORGANISMAL_PROCESS | 186 | 0.397975 | 1.4924983 | 0.0 | 0.15376648 |
| GO_POSITIVE_REGULATION_OF_CELL_DIFFERENTIATION | 100 | 0.41573346 | 1.4893477 | 0.007021063 | 0.15712342 |
| GO_REGULATION_OF_SECRETION | 87 | 0.41348037 | 1.4844035 | 0.0070635723 | 0.16297576 |
| GO_INTERACTION_WITH_HOST | 20 | 0.4988003 | 1.4759545 | 0.044843048 | 0.17346366 |
| GO_T_CELL_ACTIVATION | 63 | 0.42417064 | 1.4754769 | 0.015337423 | 0.17314042 |
| GO_OSSIFICATION | 65 | 0.42330956 | 1.4745574 | 0.011167512 | 0.17339729 |
| GO_NEGATIVE_REGULATION_OF_CELL_ACTIVATION | 30 | 0.47458872 | 1.4734434 | 0.030042918 | 0.17404959 |
| GO_CYTOKINE_SECRETION | 28 | 0.47669953 | 1.4721036 | 0.047878128 | 0.1750374 |
| GO_REGULATION_OF_LEUKOCYTE_DIFFERENTIATION | 34 | 0.46004438 | 1.4720439 | 0.04223865 | 0.17414725 |
| GO_PEPTIDE_SECRETION | 61 | 0.42566165 | 1.4714671 | 0.018348623 | 0.17396648 |
| GO_INTEGRIN_MEDIATED_SIGNALING_PATHWAY | 19 | 0.50470394 | 1.4708029 | 0.05473204 | 0.17399375 |
| GO_POSITIVE_REGULATION_OF_SIGNALING | 163 | 0.3963531 | 1.4697486 | 0.002 | 0.17451629 |
| GO_PROTEOLYSIS | 130 | 0.3983642 | 1.4642069 | 0.0020100502 | 0.18155423 |
| GO_RESPONSE_TO_CORTICOSTEROID | 19 | 0.5154021 | 1.4614165 | 0.056179777 | 0.18459569 |
| GO_POSITIVE_REGULATION_OF_RESPONSE_TO_WOUNDING | 16 | 0.52332723 | 1.4610062 | 0.072769955 | 0.18411219 |
| GO_POSITIVE_REGULATION_OF_HEMOPOIESIS | 27 | 0.47152275 | 1.4608517 | 0.069078945 | 0.18333398 |
| GO_DEFENSE_RESPONSE_TO_VIRUS | 25 | 0.46956545 | 1.4598194 | 0.0416222 | 0.18401572 |

| | | | | | |
|---|---|---|---|---|---|
| GO_SECOND_MESSENGER_MEDIATED_SIGNALING | 48 | 0.43098715 | 1.4591072 | 0.04140787 | 0.18406115 |
| GO_REPRODUCTIVE_SYSTEM_DEVELOPMENT | 33 | 0.4494553 | 1.4538778 | 0.04584222 | 0.1908894 |
| GO_VIRAL_LIFE_CYCLE | 29 | 0.46586505 | 1.4502544 | 0.040348966 | 0.19560622 |
| GO_SECRETION | 184 | 0.38552585 | 1.4498091 | 0.003 | 0.1952359 |
| GO_CYCLIC_NUCLEOTIDE_MEDIATED_SIGNALING | 19 | 0.50248164 | 1.4479524 | 0.05044843 | 0.19718319 |
| GO_WOUND_HEALING | 88 | 0.4035302 | 1.4450736 | 0.02016129 | 0.20055702 |
| GO_T_CELL_DIFFERENTIATION | 32 | 0.45041353 | 1.4446223 | 0.055853922 | 0.20020708 |
| GO_INTERFERON_GAMMA_PRODUCTION | 19 | 0.5066379 | 1.4425399 | 0.07386363 | 0.20220043 |
| GO_POSITIVE_REGULATION_OF_ERK1_AND_ERK2_CASCADE | 27 | 0.4707342 | 1.4418416 | 0.07189543 | 0.20236753 |
| GO_T_CELL_ACTIVATION_INVOLVED_IN_IMMUNE_RESPONSE | 16 | 0.51126194 | 1.4414847 | 0.0748538 | 0.20192851 |
| GO_RAS_PROTEIN_SIGNAL_TRANSDUCTION | 28 | 0.46238512 | 1.4414704 | 0.052801725 | 0.20092867 |
| GO_HORMONE_TRANSPORT | 26 | 0.47585315 | 1.4412497 | 0.059536934 | 0.20033818 |
| GO_CELL_CELL_ADHESION_VIA_PLASMA_MEMBRANE_ADHESION_MOLECULES | 35 | 0.44804442 | 1.4409956 | 0.046364594 | 0.19969083 |
| GO_RESPONSE_TO_INTERLEUKIN_1 | 26 | 0.4633444 | 1.4403998 | 0.067908 | 0.19958232 |
| GO_POSITIVE_REGULATION_OF_LEUKOCYTE_MEDIATED_IMMUNITY | 16 | 0.5063735 | 1.4398941 | 0.064177364 | 0.19938746 |
| GO_MONONUCLEAR_CELL_MIGRATION | 15 | 0.5186644 | 1.4397652 | 0.082932696 | 0.19859421 |
| GO_IMMUNE_SYSTEM_DEVELOPMENT | 89 | 0.402233 | 1.4391925 | 0.008056395 | 0.19860402 |
| GO_EXOCYTOSIS | 120 | 0.39386436 | 1.4369286 | 0.006006006 | 0.20124556 |
| GO_POSITIVE_REGULATION_OF_INTRACELLULAR_SIGNAL_TRANSDUCTION | 100 | 0.3993975 | 1.4362274 | 0.014112903 | 0.20132282 |
| GO_REGULATION_OF_ANATOMICAL_STRUCTURE_MORPHOGENESIS | 116 | 0.39204603 | 1.4362203 | 0.012060301 | 0.20035969 |
| GO_CELL_ADHESION_MEDIATED_BY_INTEGRIN | 17 | 0.5111911 | 1.4322207 | 0.069634706 | 0.20535553 |
| GO_CIRCULATORY_SYSTEM_DEVELOPMENT | 120 | 0.3879875 | 1.4310678 | 0.0070281127 | 0.20625389 |
| GO_INTERLEUKIN_1_PRODUCTION | 19 | 0.49858013 | 1.4289372 | 0.0776153 | 0.2086785 |
| GO_EPIDERMAL_CELL_DIFFERENTIATION | 27 | 0.45973042 | 1.428554 | 0.06832972 | 0.20824847 |
| GO_NEGATIVE_REGULATION_OF_DEFENSE_RESPONSE | 22 | 0.47835863 | 1.4275604 | 0.058628317 | 0.20887518 |
| GO_RESPONSE_TO_OXYGEN_CONTAINING_COMPOUND | 181 | 0.38073152 | 1.4274778 | 0.002 | 0.2080099 |
| GO_REGULATION_OF_VESICLE_MEDIATED_TRANSPORT | 65 | 0.4067531 | 1.4262128 | 0.025406504 | 0.20892617 |
| GO_RESPONSE_TO_INTERFERON_GAMMA | 40 | 0.43382362 | 1.4241894 | 0.054450262 | 0.21136953 |
| GO_CELLULAR_RESPONSE_TO_EXTRACELLULAR_STIMULUS | 34 | 0.43695748 | 1.4240776 | 0.06414301 | 0.21052259 |
| GO_REGULATION_OF_PEPTIDE_SECRETION | 53 | 0.4151236 | 1.4230454 | 0.049331963 | 0.21114261 |
| GO_EPITHELIAL_CELL_APOPTOTIC_PROCESS | 17 | 0.503453 | 1.4209905 | 0.06728538 | 0.2132392 |
| GO_RESPONSE_TO_VIRUS | 40 | 0.42902938 | 1.4181687 | 0.060732983 | 0.21673538 |
| GO_CELLULAR_COMPONENT_MORPHOGENESIS | 119 | 0.3832451 | 1.4153786 | 0.004004004 | 0.2206787 |
| GO_RESPONSE_TO_NUTRIENT | 39 | 0.42647234 | 1.414015 | 0.08559499 | 0.22185475 |
| GO_REGULATION_OF_ESTABLISHMENT_OF_PROTEIN_LOCALIZATION | 64 | 0.40100083 | 1.4124765 | 0.045408677 | 0.22346963 |
| GO_CHONDROCYTE_DIFFERENTIATION | 16 | 0.5127484 | 1.4121037 | 0.0811124 | 0.22308803 |
| GO_RESPONSE_TO_ESTRADIOL | 16 | 0.51058626 | 1.4119498 | 0.07385697 | 0.22230734 |
| GO_RHO_PROTEIN_SIGNAL_TRANSDUCTION | 18 | 0.49380243 | 1.4109136 | 0.08457143 | 0.2230972 |
| GO_POSITIVE_REGULATION_OF_CALCIUM_ION_TRANSPORT | 20 | 0.48686826 | 1.4067464 | 0.08896396 | 0.22891396 |
| GO_REGULATION_OF_HORMONE_SECRETION | 23 | 0.4646423 | 1.405527 | 0.091319054 | 0.23000841 |
| GO_SENSORY_PERCEPTION_OF_PAIN | 15 | 0.5253301 | 1.404693 | 0.085580304 | 0.23035748 |
| GO_PRODUCTION_OF_MOLECULAR_MEDIATOR_OF_IMMUNE_RESPONSE | 17 | 0.499163 | 1.4019932 | 0.08323831 | 0.2338139 |
| GO_OSTEOBLAST_DIFFERENTIATION | 34 | 0.4364591 | 1.4019231 | 0.07561235 | 0.23292989 |
| GO_REGULATION_OF_MYELOID_LEUKOCYTE_DIFFERENTIATION | 15 | 0.5104832 | 1.4007144 | 0.1005848 | 0.23390952 |
| GO_HOMEOSTASIS_OF_NUMBER_OF_CELLS | 29 | 0.44329044 | 1.3990396 | 0.077586204 | 0.23563774 |
| GO_RESPONSE_TO_EXTRACELLULAR_STIMULUS | 64 | 0.3947394 | 1.3987497 | 0.049847405 | 0.23516586 |
| GO_POSITIVE_REGULATION_OF_PEPTIDE_SECRETION | 35 | 0.4339599 | 1.3979691 | 0.07845188 | 0.23548703 |
| GO_SUPEROXIDE_METABOLIC_PROCESS | 15 | 0.5064195 | 1.397497 | 0.085946575 | 0.2353386 |

| | | | | | |
|---|---|---|---|---|---|
| GO_SMALL_GTPASE_MEDIATED_SIGNAL_TRANSDUCTION | 43 | 0.4186577 | 1.3937852 | 0.06735751 | 0.24090806 |
| GO_RECEPTOR_MEDIATED_ENDOCYTOSIS | 34 | 0.43260726 | 1.389896 | 0.075187966 | 0.24655023 |
| GO_POSITIVE_REGULATION_OF_PROTEIN_METABOLIC_PROCESS | 152 | 0.37283394 | 1.3888837 | 0.013026052 | 0.24730925 |
| GO_REGULATION_OF_CELL_POPULATION_PROLIFERATION | 153 | 0.37587672 | 1.3884181 | 0.015015015 | 0.24708864 |
| GO_ALPHA_BETA_T_CELL_ACTIVATION | 27 | 0.44395885 | 1.3858005 | 0.093247585 | 0.25067416 |
| GO_BIOMINERALIZATION | 21 | 0.46364117 | 1.3838856 | 0.09630459 | 0.25310126 |
| GO_POSITIVE_REGULATION_OF_CELL_POPULATION_PROLIFERATION | 87 | 0.38275048 | 1.3824794 | 0.032290615 | 0.2546816 |
| GO_MESENCHYMAL_CELL_DIFFERENTIATION | 24 | 0.45248887 | 1.382323 | 0.10792951 | 0.25391215 |
| GO_REGULATION_OF_EXOCYTOSIS | 27 | 0.44470122 | 1.3817971 | 0.08690987 | 0.25376138 |
| GO_APOPTOTIC_SIGNALING_PATHWAY | 57 | 0.39816964 | 1.3805313 | 0.06454918 | 0.25491223 |
| GO_INTERSPECIES_INTERACTION_BETWEEN_ORGANISMS | 79 | 0.38698483 | 1.3793148 | 0.044489384 | 0.25613335 |
| GO_ENDOCYTOSIS | 76 | 0.38900977 | 1.3781416 | 0.036400404 | 0.2571612 |
| GO_TISSUE_REMODELING | 26 | 0.4407634 | 1.3768628 | 0.0990099 | 0.25841153 |
| GO_NEGATIVE_REGULATION_OF_CATALYTIC_ACTIVITY | 58 | 0.3976394 | 1.3760499 | 0.05498982 | 0.25894037 |
| GO_TISSUE_HOMEOSTASIS | 22 | 0.45688286 | 1.3752246 | 0.10313901 | 0.25950828 |
| GO_NEURON_MIGRATION | 18 | 0.4774205 | 1.3711048 | 0.11374407 | 0.26594368 |
| GO_NEGATIVE_REGULATION_OF_RESPONSE_TO_EXTERNAL_STIMULUS | 42 | 0.41018483 | 1.3705462 | 0.077568136 | 0.2659261 |
| GO_POSITIVE_REGULATION_OF_TRANSMEMBRANE_TRANSPORT | 27 | 0.44757742 | 1.3701775 | 0.09219089 | 0.26553836 |
| GO_REGULATION_OF_MUSCLE_CELL_DIFFERENTIATION | 17 | 0.4811577 | 1.3659284 | 0.11587486 | 0.2722642 |
| GO_RESPONSE_TO_GROWTH_FACTOR | 73 | 0.38589257 | 1.365274 | 0.05172414 | 0.27239862 |
| GO_PEPTIDE_HORMONE_SECRETION | 21 | 0.46547592 | 1.3644329 | 0.10414334 | 0.27281678 |
| GO_POSITIVE_REGULATION_OF_CYTOKINE_SECRETION | 21 | 0.46139178 | 1.3618109 | 0.122847304 | 0.27669215 |
| GO_G_PROTEIN_COUPLED_RECEPTOR_SIGNALING_PATHWAY_COUPLED_TO_CYCLIC_NUCLEOTIDE_SECOND_MESSE | 22 | 0.46168834 | 1.3614354 | 0.109619685 | 0.2763585 |
| GO_INTERFERON_GAMMA_MEDIATED_SIGNALING_PATHWAY | 16 | 0.48620266 | 1.3610162 | 0.10502283 | 0.2761184 |
| GO_ANATOMICAL_STRUCTURE_FORMATION_INVOLVED_IN_MORPHOGENESIS | 122 | 0.3735093 | 1.3605659 | 0.02008032 | 0.27587357 |
| GO_PEPTIDYL_TYROSINE_MODIFICATION | 33 | 0.4230534 | 1.360153 | 0.09331919 | 0.27562672 |
| GO_REGULATION_OF_TRANSPORT | 180 | 0.36468297 | 1.3598007 | 0.008 | 0.27520937 |
| GO_RESPONSE_TO_STEROID_HORMONE | 37 | 0.41337654 | 1.359214 | 0.09437964 | 0.2752979 |
| GO_RESPONSE_TO_METAL_ION | 42 | 0.41332093 | 1.3578043 | 0.08012487 | 0.27693456 |
| GO_REGULATION_OF_INTRACELLULAR_SIGNAL_TRANSDUCTION | 159 | 0.3630199 | 1.3569808 | 0.025 | 0.27747232 |
| GO_REGULATION_OF_LYMPHOCYTE_DIFFERENTIATION | 23 | 0.4554407 | 1.355395 | 0.11758242 | 0.27930987 |
| GO_NEGATIVE_REGULATION_OF_HYDROLASE_ACTIVITY | 38 | 0.4143013 | 1.3534782 | 0.09513742 | 0.28166145 |
| GO_POSITIVE_REGULATION_OF_PEPTIDASE_ACTIVITY | 28 | 0.42747888 | 1.352208 | 0.111827955 | 0.2832117 |
| GO_REGENERATION | 22 | 0.45030957 | 1.3507036 | 0.11928651 | 0.28504202 |
| GO_NEURON_DEATH | 19 | 0.47684878 | 1.3494315 | 0.111751154 | 0.2864699 |
| GO_REGULATION_OF_CELL_DIFFERENTIATION | 160 | 0.3636854 | 1.3483653 | 0.019038076 | 0.2873812 |
| GO_ORGAN_GROWTH | 19 | 0.46797544 | 1.3468165 | 0.11649366 | 0.28938168 |
| GO_RESPONSE_TO_TRANSFORMING_GROWTH_FACTOR_BETA | 25 | 0.4434268 | 1.3462952 | 0.12967034 | 0.28924206 |
| GO_INTERLEUKIN_1_BETA_PRODUCTION | 17 | 0.48241463 | 1.3447838 | 0.11896349 | 0.29093164 |
| GO_POSITIVE_REGULATION_OF_CELL_DEVELOPMENT | 52 | 0.39373323 | 1.340906 | 0.07525773 | 0.2976689 |
| GO_NEGATIVE_REGULATION_OF_TRANSPORT | 55 | 0.39155343 | 1.3400495 | 0.08273009 | 0.29828873 |
| GO_POSITIVE_REGULATION_OF_ESTABLISHMENT_OF_PROTEIN_LOCALIZATION | 45 | 0.39985543 | 1.3372183 | 0.100926876 | 0.30263293 |
| GO_RESPONSE_TO_ACID_CHEMICAL | 43 | 0.40014103 | 1.3370581 | 0.098752595 | 0.30187994 |
| GO_COAGULATION | 47 | 0.40258208 | 1.3359253 | 0.07772021 | 0.3029365 |
| GO_REGULATION_OF_RAS_PROTEIN_SIGNAL_TRANSDUCTION | 17 | 0.4683137 | 1.3355895 | 0.1264637 | 0.30254865 |
| GO_CELL_MORPHOGENESIS_INVOLVED_IN_DIFFERENTIATION | 75 | 0.37981915 | 1.3351725 | 0.06281661 | 0.30225545 |
| GO_RESPONSE_TO_ABIOTIC_STIMULUS | 107 | 0.36657226 | 1.3342583 | 0.037111335 | 0.30302373 |
| GO_REGULATION_OF_CYTOSOLIC_CALCIUM_ION_CONCENTRATION | 43 | 0.40032926 | 1.3322551 | 0.10947368 | 0.3059635 |

| | | | | | |
|---|---|---|---|---|---|
| GO_REGULATION_OF_CELL_MORPHOGENESIS | 55 | 0.38741985 | 1.3317868 | 0.08790072 | 0.3057995 |
| GO_REGULATION_OF_T_CELL_DIFFERENTIATION | 16 | 0.4805787 | 1.3313121 | 0.13394919 | 0.30565488 |
| GO_TUBE_MORPHOGENESIS | 101 | 0.36780643 | 1.3312937 | 0.04321608 | 0.30463073 |
| GO_CALCIUM_MEDIATED_SIGNALING | 28 | 0.42424798 | 1.3304543 | 0.10560345 | 0.30536777 |
| GO_CELLULAR_RESPONSE_TO_ABIOTIC_STIMULUS | 25 | 0.43552947 | 1.3282993 | 0.12840043 | 0.30860478 |
| GO_POSITIVE_REGULATION_OF_CATALYTIC_ACTIVITY | 125 | 0.36190116 | 1.3273654 | 0.03203203 | 0.3093695 |
| GO_RESPONSE_TO_ENDOGENOUS_STIMULUS | 167 | 0.35606363 | 1.3262119 | 0.024 | 0.31063432 |
| GO_GLIOGENESIS | 43 | 0.40162066 | 1.3249382 | 0.12041885 | 0.31206006 |
| GO_POSITIVE_REGULATION_OF_MOLECULAR_FUNCTION | 153 | 0.35930175 | 1.3234898 | 0.031031031 | 0.31393224 |
| GO_NEUROTRANSMITTER_METABOLIC_PROCESS | 15 | 0.49196765 | 1.323323 | 0.12662722 | 0.31319577 |
| GO_DEVELOPMENTAL_CELL_GROWTH | 22 | 0.44201007 | 1.3230144 | 0.14333333 | 0.312696 |
| GO_AMEBOIDAL_TYPE_CELL_MIGRATION | 45 | 0.39721966 | 1.3218577 | 0.10330579 | 0.314045 |
| GO_TUBE_DEVELOPMENT | 122 | 0.3597987 | 1.3194561 | 0.044044044 | 0.31774822 |
| GO_REGULATION_OF_CELL_DEVELOPMENT | 83 | 0.36815223 | 1.3191727 | 0.06962664 | 0.3172179 |
| GO_B_CELL_DIFFERENTIATION | 17 | 0.47305107 | 1.3170319 | 0.13921113 | 0.32025912 |
| GO_RESPONSE_TO_HORMONE | 95 | 0.36407936 | 1.3156272 | 0.06363636 | 0.32183093 |
| GO_RESPONSE_TO_REACTIVE_OXYGEN_SPECIES | 25 | 0.42618886 | 1.3149424 | 0.14109743 | 0.32212463 |
| GO_ERK1_AND_ERK2_CASCADE | 43 | 0.3936569 | 1.3149055 | 0.12382934 | 0.3211166 |
| GO_ALPHA_BETA_T_CELL_DIFFERENTIATION | 19 | 0.4613614 | 1.3141977 | 0.14623173 | 0.32139122 |
| GO_REGULATION_OF_HYDROLASE_ACTIVITY | 120 | 0.3570982 | 1.3131387 | 0.05511022 | 0.32251105 |
| GO_REGULATION_OF_MYELOID_CELL_DIFFERENTIATION | 24 | 0.4346081 | 1.3130981 | 0.12513843 | 0.32154375 |
| GO_REGULATION_OF_PROTEIN_LOCALIZATION | 88 | 0.36383912 | 1.3117359 | 0.072289154 | 0.32336652 |
| GO_VASCULAR_PROCESS_IN_CIRCULATORY_SYSTEM | 24 | 0.4288923 | 1.3107595 | 0.14827202 | 0.3242984 |
| GO_MONOCARBOXYLIC_ACID_BIOSYNTHETIC_PROCESS | 22 | 0.4392484 | 1.3105797 | 0.14221725 | 0.32359383 |
| GO_POSITIVE_REGULATION_OF_HYDROLASE_ACTIVITY | 80 | 0.36406702 | 1.3100456 | 0.079878666 | 0.32358056 |
| GO_NEGATIVE_REGULATION_OF_CELL_POPULATION_PROLIFERATION | 77 | 0.36876473 | 1.3097366 | 0.08559919 | 0.32314062 |
| GO_NEGATIVE_REGULATION_OF_LYMPHOCYTE_ACTIVATION | 18 | 0.45753622 | 1.3083643 | 0.15261959 | 0.32490796 |
| GO_SENSORY_PERCEPTION_OF_LIGHT_STIMULUS | 15 | 0.48118106 | 1.3079082 | 0.15647058 | 0.3247417 |
| GO_REGULATION_OF_CELL_MORPHOGENESIS_INVOLVED_IN_DIFFERENTIATION | 27 | 0.4250172 | 1.3060735 | 0.14806867 | 0.32755366 |
| GO_PLATELET_DEGRANULATION | 30 | 0.41612968 | 1.3056093 | 0.13804348 | 0.3274265 |
| GO_REGULATION_OF_EXTRINSIC_APOPTOTIC_SIGNALING_PATHWAY | 20 | 0.45428962 | 1.3029956 | 0.1446613 | 0.331664 |
| GO_REGULATION_OF_LIPID_LOCALIZATION | 19 | 0.44909695 | 1.3018688 | 0.1513083 | 0.3328625 |
| GO_REGULATION_OF_HORMONE_LEVELS | 41 | 0.39666963 | 1.301636 | 0.12916666 | 0.33230984 |
| GO_REGULATION_OF_HEMOPOIESIS | 47 | 0.38480148 | 1.3013653 | 0.11258956 | 0.3317205 |
| GO_DEVELOPMENTAL_PROCESS_INVOLVED_IN_REPRODUCTION | 51 | 0.3811118 | 1.297759 | 0.1184346 | 0.33805236 |
| GO_SUPRAMOLECULAR_FIBER_ORGANIZATION | 96 | 0.35530165 | 1.2969973 | 0.06955645 | 0.33853292 |
| GO_NEGATIVE_REGULATION_OF_CELL_MOTILITY | 50 | 0.38828534 | 1.2967732 | 0.11408016 | 0.33790785 |
| GO_PLATELET_ACTIVATION | 30 | 0.41814554 | 1.295776 | 0.13224044 | 0.3389287 |
| GO_NEGATIVE_REGULATION_OF_MULTI_ORGANISM_PROCESS | 22 | 0.43394366 | 1.2946174 | 0.16444445 | 0.3401381 |
| GO_REACTIVE_OXYGEN_SPECIES_METABOLIC_PROCESS | 39 | 0.39771956 | 1.2933977 | 0.12970711 | 0.34157884 |
| GO_REGULATION_OF_BODY_FLUID_LEVELS | 68 | 0.36338216 | 1.2922095 | 0.0821501 | 0.3431009 |
| GO_REGULATION_OF_ENDOTHELIAL_CELL_MIGRATION | 19 | 0.45017448 | 1.291113 | 0.16400911 | 0.34433934 |
| GO_RESPONSE_TO_INORGANIC_SUBSTANCE | 61 | 0.3761597 | 1.2910903 | 0.11675127 | 0.34333155 |
| GO_METAL_ION_HOMEOSTASIS | 65 | 0.36819807 | 1.2906168 | 0.11405295 | 0.34327355 |
| GO_NEGATIVE_REGULATION_OF_MOLECULAR_FUNCTION | 78 | 0.35817623 | 1.2893679 | 0.1082996 | 0.34477323 |
| GO_PHOSPHATIDYLINOSITOL_3_KINASE_SIGNALING | 18 | 0.44886482 | 1.2881619 | 0.19435029 | 0.34637398 |
| GO_POSITIVE_REGULATION_OF_MYELOID_CELL_DIFFERENTIATION | 15 | 0.47009808 | 1.2880139 | 0.14906104 | 0.34561342 |
| GO_STEM_CELL_DIFFERENTIATION | 24 | 0.4259706 | 1.2861077 | 0.16870144 | 0.34835935 |

| | | | | | |
|---|---|---|---|---|---|
| GO_CELLULAR_RESPONSE_TO_NITROGEN_COMPOUND | 78 | 0.3645885 | 1.2850657 | 0.08924949 | 0.34937382 |
| GO_NEGATIVE_REGULATION_OF_LOCOMOTION | 51 | 0.37990347 | 1.2847762 | 0.13975155 | 0.34895888 |
| GO_NEGATIVE_REGULATION_OF_VASCULATURE_DEVELOPMENT | 17 | 0.45356464 | 1.2840917 | 0.18900344 | 0.34943718 |
| GO_REGULATION_OF_CELL_SIZE | 19 | 0.4485406 | 1.2827634 | 0.17687075 | 0.35115615 |
| GO_DEVELOPMENTAL_GROWTH | 52 | 0.3726147 | 1.2817906 | 0.13333334 | 0.35217786 |
| GO_DIVALENT_INORGANIC_CATION_HOMEOSTASIS | 55 | 0.37013435 | 1.2809736 | 0.118367344 | 0.3528773 |
| GO_CELLULAR_RESPONSE_TO_ENDOGENOUS_STIMULUS | 136 | 0.34809732 | 1.2808858 | 0.057057057 | 0.35201612 |
| GO_REGULATION_OF_CYSTEINE_TYPE_ENDOPEPTIDASE_ACTIVITY | 25 | 0.4160213 | 1.277798 | 0.18213059 | 0.35733646 |
| GO_POSITIVE_REGULATION_OF_PHOSPHORUS_METABOLIC_PROCESS | 98 | 0.35650685 | 1.2770988 | 0.078470826 | 0.35764658 |
| GO_POSITIVE_REGULATION_OF_PROTEOLYSIS | 42 | 0.38850972 | 1.276407 | 0.13972889 | 0.35806397 |
| GO_ENDOTHELIAL_CELL_PROLIFERATION | 15 | 0.47292465 | 1.2760861 | 0.18479532 | 0.35770434 |
| GO_RESPONSE_TO_FIBROBLAST_GROWTH_FACTOR | 18 | 0.44310313 | 1.2705513 | 0.1661017 | 0.36837494 |
| GO_EXTRINSIC_APOPTOTIC_SIGNALING_PATHWAY | 27 | 0.4099899 | 1.2691236 | 0.17576419 | 0.3701546 |
| GO_POSITIVE_REGULATION_OF_MAPK_CASCADE | 52 | 0.36978406 | 1.2643138 | 0.15653965 | 0.37930107 |
| GO_RESPONSE_TO_ORGANIC_CYCLIC_COMPOUND | 99 | 0.3531326 | 1.264127 | 0.10140562 | 0.37859672 |
| GO_REGULATION_OF_EPITHELIAL_CELL_MIGRATION | 25 | 0.409311 | 1.2638448 | 0.1899012 | 0.37810045 |
| GO_REGULATION_OF_SMALL_GTPASE_MEDIATED_SIGNAL_TRANSDUCTION | 28 | 0.40115234 | 1.2636894 | 0.16993465 | 0.37740424 |
| GO_REGULATION_OF_ALPHA_BETA_T_CELL_ACTIVATION | 19 | 0.43790925 | 1.2626716 | 0.19668508 | 0.37844905 |
| GO_RESPONSE_TO_CALCIUM_ION | 22 | 0.42436793 | 1.2617695 | 0.17920354 | 0.37914833 |
| GO_SENSORY_PERCEPTION | 40 | 0.38295057 | 1.2610321 | 0.1661442 | 0.37977928 |
| GO_CELL_GROWTH | 40 | 0.38077524 | 1.2599765 | 0.15295358 | 0.38099256 |
| GO_REGULATION_OF_REACTIVE_OXYGEN_SPECIES_METABOLIC_PROCESS | 26 | 0.40961263 | 1.2591352 | 0.1779476 | 0.38169786 |
| GO_CARBOHYDRATE_DERIVATIVE_CATABOLIC_PROCESS | 19 | 0.4342666 | 1.2534037 | 0.2073991 | 0.3934105 |
| GO_NEGATIVE_REGULATION_OF_PROTEIN_METABOLIC_PROCESS | 82 | 0.35114375 | 1.2532477 | 0.1160444 | 0.39265093 |
| GO_POSITIVE_REGULATION_OF_ION_TRANSPORT | 45 | 0.3764752 | 1.2530394 | 0.16024974 | 0.3919456 |
| GO_REGULATION_OF_CELLULAR_RESPONSE_TO_GROWTH_FACTOR_STIMULUS | 26 | 0.39917603 | 1.2518868 | 0.19496167 | 0.39338925 |
| GO_BONE_MINERALIZATION | 19 | 0.42750376 | 1.2516904 | 0.21274175 | 0.39276785 |
| GO_NEGATIVE_REGULATION_OF_CELLULAR_RESPONSE_TO_GROWTH_FACTOR_STIMULUS | 15 | 0.45150736 | 1.2512239 | 0.20689656 | 0.3927384 |
| GO_REGULATION_OF_TUBE_SIZE | 20 | 0.42786434 | 1.2501532 | 0.20382883 | 0.39399883 |
| GO_NEGATIVE_REGULATION_OF_RESPONSE_TO_STIMULUS | 150 | 0.337279 | 1.2466285 | 0.071 | 0.40066424 |
| GO_RESPONSE_TO_OXYGEN_LEVELS | 43 | 0.37676895 | 1.2460065 | 0.1682243 | 0.40096062 |
| GO_INOSITOL_LIPID_MEDIATED_SIGNALING | 24 | 0.4034173 | 1.2437388 | 0.22958058 | 0.4048614 |
| GO_NEGATIVE_REGULATION_OF_CELL_DEATH | 76 | 0.35288504 | 1.2412052 | 0.14213198 | 0.4094942 |
| GO_AGING | 45 | 0.36909017 | 1.2402472 | 0.17592593 | 0.4105415 |
| GO_REGULATION_OF_CALCIUM_ION_TRANSPORT | 36 | 0.37918893 | 1.2381885 | 0.19704953 | 0.4140251 |
| GO_NEGATIVE_REGULATION_OF_ION_TRANSPORT | 15 | 0.45551515 | 1.2378005 | 0.20693642 | 0.41376185 |
| GO_POSITIVE_REGULATION_OF_ION_TRANSMEMBRANE_TRANSPORT | 23 | 0.41791067 | 1.2358283 | 0.1942605 | 0.41717073 |
| GO_EPIDERMIS_DEVELOPMENT | 36 | 0.38329262 | 1.2357726 | 0.18124342 | 0.41619492 |
| GO_DNA_METABOLIC_PROCESS | 22 | 0.42333576 | 1.235524 | 0.22838138 | 0.4156098 |
| GO_POSITIVE_REGULATION_OF_PROTEIN_MODIFICATION_PROCESS | 96 | 0.34065354 | 1.2353566 | 0.12612613 | 0.4148847 |
| GO_ENDOTHELIAL_CELL_MIGRATION | 25 | 0.40788746 | 1.2345837 | 0.21081677 | 0.41542023 |
| GO_AXON_DEVELOPMENT | 45 | 0.3690008 | 1.2337713 | 0.18181819 | 0.41611785 |
| GO_NEGATIVE_REGULATION_OF_CELL_DEVELOPMENT | 27 | 0.3982768 | 1.2332933 | 0.20479302 | 0.4160387 |
| GO_GROWTH | 71 | 0.34750152 | 1.2323107 | 0.1605691 | 0.4171399 |
| GO_CELLULAR_RESPONSE_TO_ANTIBIOTIC | 16 | 0.44835997 | 1.2296419 | 0.2153667 | 0.42213655 |
| GO_PROTEIN_KINASE_B_SIGNALING | 35 | 0.3766137 | 1.2265626 | 0.222103 | 0.42790908 |
| GO_RESPONSE_TO_ANTIBIOTIC | 45 | 0.36523095 | 1.2209183 | 0.19186653 | 0.4395476 |
| GO_RESPONSE_TO_OXIDATIVE_STRESS | 44 | 0.3674488 | 1.220551 | 0.2104712 | 0.43928754 |

| | | | | | |
|---|---|---|---|---|---|
| GO_REGULATION_OF_CELL_MATRIX_ADHESION | 16 | 0.44431415 | 1.218157 | 0.24110219 | 0.44369116 |
| GO_REGULATION_OF_GLIOGENESIS | 18 | 0.42982587 | 1.2172092 | 0.225058 | 0.4447585 |
| GO_NEGATIVE_REGULATION_OF_LEUKOCYTE_CELL_CELL_ADHESION | 17 | 0.43945903 | 1.2161987 | 0.24942791 | 0.4459591 |
| GO_REGULATION_OF_PEPTIDYL_TYROSINE_PHOSPHORYLATION | 23 | 0.39962578 | 1.2095463 | 0.24347825 | 0.4600647 |
| GO_RESPONSE_TO_RADIATION | 25 | 0.3941847 | 1.2092742 | 0.2378902 | 0.45951185 |
| GO_FATTY_ACID_DERIVATIVE_METABOLIC_PROCESS | 17 | 0.42685 | 1.2078685 | 0.23650976 | 0.46160528 |
| GO_REGULATION_OF_GTPASE_ACTIVITY | 36 | 0.3658507 | 1.2068473 | 0.24207188 | 0.46284577 |
| GO_REGULATION_OF_NIK_NF_KAPPAB_SIGNALING | 15 | 0.44027194 | 1.2064477 | 0.25322393 | 0.46263692 |
| GO_POSITIVE_REGULATION_OF_CELLULAR_BIOSYNTHETIC_PROCESS | 118 | 0.32940668 | 1.2061044 | 0.1513026 | 0.46225512 |
| GO_LIPID_LOCALIZATION | 37 | 0.37264305 | 1.2054443 | 0.23354565 | 0.46256757 |
| GO_DIVALENT_INORGANIC_CATION_TRANSPORT | 56 | 0.3527088 | 1.2054154 | 0.2073922 | 0.46145204 |
| GO_NEURON_PROJECTION_GUIDANCE | 27 | 0.38512 | 1.2044946 | 0.22099447 | 0.4624609 |
| GO_RESPONSE_TO_TEMPERATURE_STIMULUS | 18 | 0.42123416 | 1.2039407 | 0.24716553 | 0.46260414 |
| GO_HORMONE_METABOLIC_PROCESS | 18 | 0.42080867 | 1.2039121 | 0.24857469 | 0.46149218 |
| GO_PROTEIN_LOCALIZATION_TO_NUCLEUS | 17 | 0.41978696 | 1.2001143 | 0.23577236 | 0.4693572 |
| GO_RECEPTOR_SIGNALING_PATHWAY_VIA_STAT | 19 | 0.41264603 | 1.1998888 | 0.28366446 | 0.46871105 |
| GO_TRANSMEMBRANE_RECEPTOR_PROTEIN_SERINE_THREONINE_KINASE_SIGNALING_PATHWAY | 31 | 0.37336436 | 1.1994183 | 0.24232456 | 0.46854305 |
| GO_NEGATIVE_REGULATION_OF_CELLULAR_COMPONENT_ORGANIZATION | 52 | 0.34871134 | 1.1981585 | 0.2179752 | 0.4704545 |
| GO_BRANCHING_MORPHOGENESIS_OF_AN_EPITHELIAL_TUBE | 18 | 0.41706014 | 1.1976353 | 0.2632184 | 0.47043535 |
| GO_I_KAPPAB_KINASE_NF_KAPPAB_SIGNALING | 34 | 0.3759981 | 1.1964581 | 0.23157895 | 0.47192138 |
| GO_UROGENITAL_SYSTEM_DEVELOPMENT | 41 | 0.35766777 | 1.1960086 | 0.24605678 | 0.4717567 |
| GO_DEVELOPMENTAL_GROWTH_INVOLVED_IN_MORPHOGENESIS | 23 | 0.3990254 | 1.1946046 | 0.23756906 | 0.47392678 |
| GO_IMPORT_INTO_CELL | 93 | 0.3309374 | 1.1944764 | 0.19095477 | 0.47302428 |
| GO_IMMUNE_RESPONSE_REGULATING_CELL_SURFACE_RECEPTOR_SIGNALING_PATHWAY | 42 | 0.35577342 | 1.194199 | 0.21991701 | 0.47248778 |
| GO_POSITIVE_REGULATION_OF_I_KAPPAB_KINASE_NF_KAPPAB_SIGNALING | 23 | 0.40157384 | 1.1898623 | 0.26977247 | 0.48161292 |
| GO_NEGATIVE_REGULATION_OF_DEVELOPMENTAL_PROCESS | 87 | 0.33080587 | 1.1897562 | 0.18365288 | 0.48066384 |
| GO_POSITIVE_REGULATION_OF_GTPASE_ACTIVITY | 33 | 0.36878115 | 1.18785 | 0.25957447 | 0.4840843 |
| GO_REGULATION_OF_BLOOD_PRESSURE | 17 | 0.42063358 | 1.1864223 | 0.25429553 | 0.48632622 |
| GO_MUSCLE_CELL_MIGRATION | 16 | 0.42357653 | 1.184415 | 0.26712328 | 0.4900466 |
| GO_RESPONSE_TO_KETONE | 24 | 0.39076972 | 1.1840245 | 0.2597826 | 0.48975697 |
| GO_POSITIVE_REGULATION_OF_CYSTEINE_TYPE_ENDOPEPTIDASE_ACTIVITY | 19 | 0.40989774 | 1.1837287 | 0.280543 | 0.48928672 |
| GO_HOMOTYPIC_CELL_CELL_ADHESION | 22 | 0.40199223 | 1.1824311 | 0.2860386 | 0.49109128 |
| GO_REGULATION_OF_PHOSPHORYLATION | 126 | 0.32329234 | 1.18132 | 0.17452358 | 0.4925884 |
| GO_NEGATIVE_REGULATION_OF_SECRETION | 35 | 0.368516 | 1.1811388 | 0.24708377 | 0.4918193 |
| GO_ANTIGEN_RECEPTOR_MEDIATED_SIGNALING_PATHWAY | 26 | 0.38780364 | 1.1800923 | 0.267101 | 0.49312955 |
| GO_MYELOID_LEUKOCYTE_DIFFERENTIATION | 28 | 0.38836125 | 1.1783293 | 0.27730194 | 0.4959332 |
| GO_REGULATION_OF_PHOSPHORUS_METABOLIC_PROCESS | 141 | 0.31717792 | 1.1771867 | 0.163 | 0.49755737 |
| GO_REGULATION_OF_APOPTOTIC_SIGNALING_PATHWAY | 37 | 0.36203402 | 1.175807 | 0.26282722 | 0.49965736 |
| GO_POSITIVE_REGULATION_OF_NEURON_PROJECTION_DEVELOPMENT | 23 | 0.39223903 | 1.1746901 | 0.269188 | 0.50113046 |
| GO_NEGATIVE_REGULATION_OF_APOPTOTIC_SIGNALING_PATHWAY | 22 | 0.39298925 | 1.1730369 | 0.26608506 | 0.5039205 |
| GO_GASTRULATION | 19 | 0.4082225 | 1.1713234 | 0.29954955 | 0.50678205 |
| GO_CELL_MATURATION | 20 | 0.40192443 | 1.1711506 | 0.2739726 | 0.5059985 |
| GO_POSITIVE_REGULATION_OF_REACTIVE_OXYGEN_SPECIES_METABOLIC_PROCESS | 17 | 0.4129301 | 1.1706336 | 0.27497062 | 0.50608724 |
| GO_SIGNAL_TRANSDUCTION_BY_PROTEIN_PHOSPHORYLATION | 90 | 0.32729393 | 1.1701559 | 0.20748988 | 0.50600135 |
| GO_NEGATIVE_REGULATION_OF_CATABOLIC_PROCESS | 21 | 0.40235743 | 1.1692452 | 0.29054055 | 0.5069663 |
| GO_GLIAL_CELL_DIFFERENTIATION | 31 | 0.37451628 | 1.1692176 | 0.27012128 | 0.5058312 |
| GO_REGULATION_OF_MAPK_CASCADE | 74 | 0.32778504 | 1.1680951 | 0.23517588 | 0.50720423 |
| GO_POSITIVE_REGULATION_OF_TRANSPORTER_ACTIVITY | 16 | 0.41624442 | 1.166978 | 0.3027523 | 0.5087863 |

| | | | | | |
|---|---|---|---|---|---|
| GO_NEURON_PROJECTION_EXTENSION | 17 | 0.40911403 | 1.1667974 | 0.2681764 | 0.5080113 |
| GO_POSITIVE_REGULATION_OF_GLIOGENESIS | 15 | 0.43144947 | 1.1665822 | 0.29142186 | 0.5073242 |
| GO_MESENCHYME_DEVELOPMENT | 30 | 0.37023604 | 1.1655791 | 0.27056277 | 0.5084849 |
| GO_CELLULAR_RESPONSE_TO_CHEMICAL_STRESS | 32 | 0.36957747 | 1.1651279 | 0.26220807 | 0.50838774 |
| GO_POSITIVE_REGULATION_OF_CELL_CYCLE | 17 | 0.41536218 | 1.164504 | 0.2962963 | 0.5086529 |
| GO_REGULATION_OF_REGULATED_SECRETORY_PATHWAY | 21 | 0.39568105 | 1.1633523 | 0.28425822 | 0.510181 |
| GO_CELLULAR_RESPONSE_TO_INORGANIC_SUBSTANCE | 21 | 0.39875224 | 1.1633354 | 0.2919955 | 0.5090487 |
| GO_PEPTIDYL_AMINO_ACID_MODIFICATION | 66 | 0.32678214 | 1.163274 | 0.25176945 | 0.5080456 |
| GO_SIGNAL_RELEASE | 37 | 0.35410202 | 1.1615592 | 0.29263157 | 0.5109946 |
| GO_POSITIVE_REGULATION_OF_PEPTIDYL_TYROSINE_PHOSPHORYLATION | 16 | 0.41625816 | 1.1572169 | 0.30490655 | 0.5200358 |
| GO_CALCIUM_ION_TRANSMEMBRANE_IMPORT_INTO_CYTOSOL | 19 | 0.39719015 | 1.155878 | 0.2918979 | 0.5219175 |
| GO_TISSUE_MIGRATION | 33 | 0.35891974 | 1.1552882 | 0.28479657 | 0.5220066 |
| GO_PROTEIN_PHOSPHORYLATION | 163 | 0.3112673 | 1.154831 | 0.2052052 | 0.5219245 |
| GO_REGULATION_OF_CELL_SHAPE | 24 | 0.3806957 | 1.1545266 | 0.30666667 | 0.5214001 |
| GO_CD4_POSITIVE_ALPHA_BETA_T_CELL_ACTIVATION | 18 | 0.4051304 | 1.1534011 | 0.31006864 | 0.5229763 |
| GO_REGULATION_OF_LIPID_TRANSPORT | 15 | 0.424011 | 1.1508515 | 0.30688447 | 0.5278947 |
| GO_TUMOR_NECROSIS_FACTOR_MEDIATED_SIGNALING_PATHWAY | 20 | 0.38989106 | 1.1471683 | 0.3105802 | 0.5357117 |
| GO_ANATOMICAL_STRUCTURE_HOMEOSTASIS | 29 | 0.37022942 | 1.1468999 | 0.2854054 | 0.5350863 |
| GO_REGULATION_OF_ENDOCYTOSIS | 24 | 0.37523156 | 1.1450776 | 0.328125 | 0.5382414 |
| GO_GAMETE_GENERATION | 25 | 0.3785786 | 1.1443776 | 0.3156146 | 0.5386707 |
| GO_NEGATIVE_REGULATION_OF_PEPTIDE_SECRETION | 18 | 0.40071088 | 1.1435263 | 0.30716723 | 0.53943765 |
| GO_MUCOPOLYSACCHARIDE_METABOLIC_PROCESS | 15 | 0.42083237 | 1.1421353 | 0.3121319 | 0.54163605 |
| GO_CELL_CELL_SIGNALING | 146 | 0.3104187 | 1.1405654 | 0.21421422 | 0.54421633 |
| GO_NEUROGENESIS | 141 | 0.3098592 | 1.1382971 | 0.218 | 0.5484017 |
| GO_ENZYME_LINKED_RECEPTOR_PROTEIN_SIGNALING_PATHWAY | 107 | 0.31540117 | 1.1365498 | 0.25527638 | 0.5514216 |
| GO_TRANSMEMBRANE_RECEPTOR_PROTEIN_TYROSINE_KINASE_SIGNALING_PATHWAY | 81 | 0.31725407 | 1.1335539 | 0.27208123 | 0.5573682 |
| GO_NEGATIVE_REGULATION_OF_CELL_CELL_ADHESION | 25 | 0.38015437 | 1.1329415 | 0.3131202 | 0.5575643 |
| GO_PLATELET_AGGREGATION | 20 | 0.38574666 | 1.1317666 | 0.34162897 | 0.5591083 |
| GO_NEGATIVE_REGULATION_OF_ESTABLISHMENT_OF_PROTEIN_LOCALIZATION | 20 | 0.3912283 | 1.12959 | 0.32794458 | 0.56297326 |
| GO_REGULATION_OF_SYMBIOSIS_ENCOMPASSING_MUTUALISM_THROUGH_PARASITISM | 19 | 0.3905304 | 1.1288189 | 0.34728506 | 0.563591 |
| GO_ORGANIC_HYDROXY_COMPOUND_METABOLIC_PROCESS | 46 | 0.33665723 | 1.1273232 | 0.313786 | 0.566044 |
| GO_NEGATIVE_REGULATION_OF_CELLULAR_CATABOLIC_PROCESS | 17 | 0.39735258 | 1.1270127 | 0.35327962 | 0.56553656 |
| GO_CELLULAR_RESPONSE_TO_ORGANIC_CYCLIC_COMPOUND | 61 | 0.32434058 | 1.1207834 | 0.3082401 | 0.5795384 |
| GO_POSITIVE_REGULATION_OF_CELLULAR_COMPONENT_ORGANIZATION | 112 | 0.3079517 | 1.1203717 | 0.29849246 | 0.57927436 |
| GO_CELL_JUNCTION_ASSEMBLY | 34 | 0.3493624 | 1.1201937 | 0.3268817 | 0.57846534 |
| GO_MAINTENANCE_OF_LOCATION | 38 | 0.34121025 | 1.1189615 | 0.3273305 | 0.5801708 |
| GO_REGULATION_OF_VIRAL_LIFE_CYCLE | 16 | 0.39368477 | 1.1187098 | 0.33060747 | 0.579547 |
| GO_CELLULAR_HOMEOSTASIS | 92 | 0.30933562 | 1.1182216 | 0.2893145 | 0.57960254 |
| GO_NEGATIVE_REGULATION_OF_SIGNALING | 127 | 0.30377105 | 1.1176319 | 0.26753506 | 0.5797805 |
| GO_REGULATION_OF_PROTEIN_MODIFICATION_PROCESS | 138 | 0.30403236 | 1.1169652 | 0.26379138 | 0.5801071 |
| GO_POSITIVE_REGULATION_OF_DNA_BINDING_TRANSCRIPTION_FACTOR_ACTIVITY | 23 | 0.36481652 | 1.1151845 | 0.36853686 | 0.5830904 |
| GO_RESPONSE_TO_ETHANOL | 23 | 0.37126693 | 1.1150866 | 0.35005453 | 0.5820779 |
| GO_REGULATION_OF_CELLULAR_LOCALIZATION | 64 | 0.32000604 | 1.1136477 | 0.3228745 | 0.5842486 |
| GO_STRIATED_MUSCLE_CELL_DIFFERENTIATION | 34 | 0.35182282 | 1.1089578 | 0.342918 | 0.5941508 |
| GO_DRUG_METABOLIC_PROCESS | 49 | 0.32290086 | 1.1082566 | 0.32917964 | 0.5945056 |
| GO_REGULATION_OF_CATION_CHANNEL_ACTIVITY | 19 | 0.379123 | 1.1008577 | 0.35722545 | 0.6109584 |
| GO_NEGATIVE_REGULATION_OF_INFLAMMATORY_RESPONSE | 16 | 0.40257365 | 1.1007664 | 0.36032864 | 0.6099038 |
| GO_CELLULAR_ION_HOMEOSTASIS | 73 | 0.30964473 | 1.0964787 | 0.33366936 | 0.61907244 |

| | | | | | |
|---|---|---|---|---|---|
| GO_CELLULAR_RESPONSE_TO_STEROID_HORMONE_STIMULUS | 16 | 0.39534175 | 1.0947069 | 0.36842105 | 0.62201333 |
| GO_CELLULAR_RESPONSE_TO_HORMONE_STIMULUS | 56 | 0.3191676 | 1.0944849 | 0.345957 | 0.6212102 |
| GO_RENAL_SYSTEM_PROCESS | 19 | 0.37745163 | 1.0939847 | 0.37330317 | 0.62106234 |
| GO_MUSCLE_ORGAN_DEVELOPMENT | 25 | 0.36008453 | 1.0933205 | 0.36553237 | 0.62131584 |
| GO_MYELOID_CELL_DIFFERENTIATION | 40 | 0.32942325 | 1.0896491 | 0.374477 | 0.6289831 |
| GO_POSITIVE_REGULATION_OF_CELL_PROJECTION_ORGANIZATION | 36 | 0.3368592 | 1.0893615 | 0.3834746 | 0.62837327 |
| GO_REGULATION_OF_ANION_TRANSPORT | 16 | 0.38930252 | 1.0888768 | 0.38416076 | 0.62829226 |
| GO_TYPE_I_INTERFERON_PRODUCTION | 16 | 0.39807388 | 1.088683 | 0.3814554 | 0.6274556 |
| GO_REGULATION_OF_TRANSMEMBRANE_RECEPTOR_PROTEIN_SERINE_THREONINE_KINASE_SIGNALING_PATHWAY | 24 | 0.3615274 | 1.0884435 | 0.37636763 | 0.626734 |
| GO_ALCOHOL_METABOLIC_PROCESS | 33 | 0.3406237 | 1.0866619 | 0.37806177 | 0.6296795 |
| GO_PROTEIN_COMPLEX_OLIGOMERIZATION | 42 | 0.3239339 | 1.0849731 | 0.3937173 | 0.6325055 |
| GO_MORPHOGENESIS_OF_A_BRANCHING_STRUCTURE | 25 | 0.3539896 | 1.0847261 | 0.37308535 | 0.631819 |
| GO_CIRCULATORY_SYSTEM_PROCESS | 64 | 0.3110288 | 1.0822297 | 0.40040857 | 0.63654876 |
| GO_ESTABLISHMENT_OF_ORGANELLE_LOCALIZATION | 27 | 0.34874588 | 1.0791229 | 0.37594798 | 0.6426166 |
| GO_RESPONSE_TO_DRUG | 102 | 0.29961446 | 1.0787193 | 0.35685483 | 0.6422585 |
| GO_ENSHEATHMENT_OF_NEURONS | 16 | 0.3875444 | 1.0786269 | 0.38973162 | 0.641189 |
| GO_POSITIVE_REGULATION_OF_ENDOCYTOSIS | 22 | 0.3582174 | 1.0784045 | 0.3903262 | 0.64046496 |
| GO_NEGATIVE_REGULATION_OF_GROWTH | 17 | 0.36988357 | 1.0783502 | 0.38559815 | 0.6392897 |
| GO_MUSCLE_TISSUE_DEVELOPMENT | 32 | 0.33697262 | 1.0769279 | 0.39331895 | 0.64126897 |
| GO_POSITIVE_REGULATION_OF_NEURON_DIFFERENTIATION | 31 | 0.34192038 | 1.0748812 | 0.40894568 | 0.6448815 |
| GO_REGULATION_OF_CELLULAR_PROTEIN_LOCALIZATION | 38 | 0.33051848 | 1.0735246 | 0.38794926 | 0.646741 |
| GO_CELLULAR_RESPONSE_TO_DRUG | 38 | 0.3270615 | 1.0708036 | 0.39556962 | 0.65198517 |
| GO_REGULATION_OF_MUSCLE_SYSTEM_PROCESS | 16 | 0.38200247 | 1.0698882 | 0.40606767 | 0.65283924 |
| GO_ANIMAL_ORGAN_MORPHOGENESIS | 114 | 0.29189125 | 1.0697409 | 0.36108324 | 0.65187025 |
| GO_HEMATOPOIETIC_PROGENITOR_CELL_DIFFERENTIATION | 15 | 0.38981238 | 1.069356 | 0.4158654 | 0.6514769 |
| GO_HOMEOSTATIC_PROCESS | 178 | 0.2866245 | 1.0693219 | 0.33633634 | 0.65025705 |
| GO_CYTOSOLIC_CALCIUM_ION_TRANSPORT | 24 | 0.35283884 | 1.0683645 | 0.40859032 | 0.6512819 |
| GO_CELL_JUNCTION_ORGANIZATION | 42 | 0.32199132 | 1.067446 | 0.3914405 | 0.6521531 |
| GO_MYELOID_CELL_HOMEOSTASIS | 19 | 0.3702803 | 1.0626725 | 0.42410198 | 0.6621374 |
| GO_REGULATION_OF_ANATOMICAL_STRUCTURE_SIZE | 57 | 0.3052428 | 1.0619782 | 0.41182467 | 0.66257244 |
| GO_CELL_PART_MORPHOGENESIS | 56 | 0.30422297 | 1.0615108 | 0.40449437 | 0.6623762 |
| GO_RESPONSE_TO_BMP | 18 | 0.37544012 | 1.0614374 | 0.4036281 | 0.66120744 |
| GO_RESPONSE_TO_NITROGEN_COMPOUND | 112 | 0.29190323 | 1.0611321 | 0.39278558 | 0.6606193 |
| GO_EPITHELIAL_CELL_DIFFERENTIATION | 74 | 0.2967886 | 1.0601754 | 0.39007092 | 0.6615081 |
| GO_AMINOGLYCAN_METABOLIC_PROCESS | 20 | 0.3668684 | 1.0562797 | 0.42519686 | 0.66946584 |
| GO_REGULATION_OF_INTRACELLULAR_TRANSPORT | 24 | 0.34906325 | 1.0561851 | 0.41694537 | 0.66839826 |
| GO_NEGATIVE_REGULATION_OF_INTRACELLULAR_SIGNAL_TRANSDUCTION | 43 | 0.31364492 | 1.0518719 | 0.42376447 | 0.6771921 |
| GO_POSITIVE_REGULATION_OF_KINASE_ACTIVITY | 44 | 0.3143109 | 1.050248 | 0.44827586 | 0.67970735 |
| GO_SULFUR_COMPOUND_BIOSYNTHETIC_PROCESS | 21 | 0.35633266 | 1.0499573 | 0.4254835 | 0.6791022 |
| GO_REGULATION_OF_MITOTIC_CELL_CYCLE | 29 | 0.33056983 | 1.0453769 | 0.45386267 | 0.6885512 |
| GO_MEMBRANE_ORGANIZATION | 68 | 0.29879662 | 1.0430295 | 0.4287169 | 0.69264877 |
| GO_NIK_NF_KAPPAB_SIGNALING | 23 | 0.35000935 | 1.0423542 | 0.44664466 | 0.69294435 |
| GO_POSITIVE_REGULATION_OF_CELLULAR_PROTEIN_LOCALIZATION | 25 | 0.34618086 | 1.0410463 | 0.44162995 | 0.6946383 |
| GO_NEURON_DIFFERENTIATION | 105 | 0.28568062 | 1.036115 | 0.4204204 | 0.7048323 |
| GO_REGULATION_OF_REPRODUCTIVE_PROCESS | 15 | 0.37524495 | 1.0357845 | 0.45847952 | 0.7041976 |
| GO_POSITIVE_REGULATION_OF_SMOOTH_MUSCLE_CELL_PROLIFERATION | 18 | 0.36099583 | 1.035704 | 0.4516496 | 0.7030306 |
| GO_EPITHELIAL_TO_MESENCHYMAL_TRANSITION | 15 | 0.3772518 | 1.0343978 | 0.43882352 | 0.7046939 |
| GO_REGULATION_OF_NERVOUS_SYSTEM_PROCESS | 15 | 0.37939617 | 1.0337055 | 0.45357144 | 0.7049746 |

| | | | | | |
|---|---|---|---|---|---|
| GO_RESPONSE_TO_PEPTIDE | 58 | 0.2950716 | 1.0328542 | 0.45407638 | 0.7054542 |
| GO_NEGATIVE_REGULATION_OF_CELL_DIFFERENTIATION | 56 | 0.3017194 | 1.031086 | 0.45482546 | 0.7083886 |
| GO_NEURON_DEVELOPMENT | 89 | 0.28785014 | 1.0303917 | 0.43473896 | 0.708731 |
| GO_AMINOGLYCAN_BIOSYNTHETIC_PROCESS | 16 | 0.37028685 | 1.0295084 | 0.4604811 | 0.70940167 |
| GO_REGULATION_OF_METAL_ION_TRANSPORT | 49 | 0.30578074 | 1.0284191 | 0.46169773 | 0.7104602 |
| GO_PROCESS_UTILIZING_AUTOPHAGIC_MECHANISM | 25 | 0.34075895 | 1.026496 | 0.4665203 | 0.71371526 |
| GO_CELLULAR_COMPONENT_ASSEMBLY_INVOLVED_IN_MORPHOGENESIS | 15 | 0.37580612 | 1.0255966 | 0.44934446 | 0.714422 |
| GO_PROTEIN_LOCALIZATION_TO_PLASMA_MEMBRANE | 22 | 0.342672 | 1.0250953 | 0.4510686 | 0.7143435 |
| GO_SYNAPSE_ORGANIZATION | 44 | 0.31255838 | 1.0237988 | 0.46715328 | 0.7159836 |
| GO_NERVOUS_SYSTEM_PROCESS | 74 | 0.28757107 | 1.0234946 | 0.47619048 | 0.71533567 |
| GO_ION_HOMEOSTASIS | 90 | 0.28666392 | 1.0206866 | 0.4653962 | 0.72038126 |
| GO_REGULATION_OF_PHOSPHATASE_ACTIVITY | 16 | 0.3642427 | 1.0192086 | 0.46064815 | 0.7224316 |
| GO_POSITIVE_REGULATION_OF_TRANSFERASE_ACTIVITY | 47 | 0.3050364 | 1.0189418 | 0.46907216 | 0.7217189 |
| GO_TRANSLATIONAL_INITIATION | 18 | 0.3536978 | 1.0181237 | 0.46215597 | 0.72213924 |
| GO_ORGANIC_ACID_BIOSYNTHETIC_PROCESS | 34 | 0.3151865 | 1.01747 | 0.47346073 | 0.72217226 |
| GO_REGULATION_OF_PROTEIN_LOCALIZATION_TO_MEMBRANE | 24 | 0.3323108 | 1.0163536 | 0.46187845 | 0.7234398 |
| GO_ACTIN_FILAMENT_BASED_PROCESS | 95 | 0.28130138 | 1.0148674 | 0.48185483 | 0.72533846 |
| GO_RESPIRATORY_SYSTEM_DEVELOPMENT | 24 | 0.3329605 | 1.0138106 | 0.46424642 | 0.7265042 |
| GO_MITOTIC_CELL_CYCLE | 42 | 0.30651906 | 1.0132599 | 0.47767395 | 0.7263714 |
| GO_GLYCOSYLATION | 17 | 0.35743147 | 1.0118127 | 0.46893317 | 0.7283863 |
| GO_POSITIVE_REGULATION_OF_PROTEIN_KINASE_B_SIGNALING | 25 | 0.33141738 | 1.0100737 | 0.4825708 | 0.7308984 |
| GO_EMBRYONIC_ORGAN_DEVELOPMENT | 34 | 0.31786793 | 1.009553 | 0.4815206 | 0.7306685 |
| GO_REGULATION_OF_PEPTIDE_HORMONE_SECRETION | 16 | 0.36319166 | 1.0076184 | 0.4729412 | 0.73354745 |
| GO_CELLULAR_RESPONSE_TO_TOXIC_SUBSTANCE | 30 | 0.31450245 | 1.0066777 | 0.4748663 | 0.7342985 |
| GO_ANTIGEN_PROCESSING_AND_PRESENTATION | 17 | 0.35571408 | 1.006167 | 0.47630057 | 0.7341472 |
| GO_RNA_CATABOLIC_PROCESS | 27 | 0.32445022 | 1.0056927 | 0.4913232 | 0.73384535 |
| GO_PROTEIN_LOCALIZATION_TO_CELL_PERIPHERY | 26 | 0.32701477 | 1.0050439 | 0.47783783 | 0.73395777 |
| GO_REGULATION_OF_BLOOD_CIRCULATION | 30 | 0.31618538 | 1.0046558 | 0.48857453 | 0.7334816 |
| GO_NEGATIVE_REGULATION_OF_EPITHELIAL_CELL_PROLIFERATION | 17 | 0.35229245 | 1.0043789 | 0.4919355 | 0.7327185 |
| GO_ESTABLISHMENT_OF_LOCALIZATION_IN_CELL | 41 | 0.30613497 | 1.0008104 | 0.48329854 | 0.7394915 |
| GO_MUSCLE_CELL_DIFFERENTIATION | 40 | 0.30773693 | 1.0006812 | 0.50840336 | 0.73840684 |
| GO_MUSCLE_STRUCTURE_DEVELOPMENT | 57 | 0.2887643 | 0.9990268 | 0.5118191 | 0.7406546 |
| GO_POLYOL_METABOLIC_PROCESS | 15 | 0.3710624 | 0.99572057 | 0.5168801 | 0.74680305 |
| GO_RESPONSE_TO_ALCOHOL | 36 | 0.3093323 | 0.9945788 | 0.4947589 | 0.7480062 |
| GO_VIRAL_GENE_EXPRESSION | 16 | 0.36276087 | 0.9940244 | 0.5069606 | 0.74781317 |
| GO_NEGATIVE_REGULATION_OF_NERVOUS_SYSTEM_DEVELOPMENT | 26 | 0.32235637 | 0.99370575 | 0.5114504 | 0.74714524 |
| GO_REGULATION_OF_PROTEIN_BINDING | 15 | 0.3662949 | 0.9935979 | 0.48300117 | 0.74601454 |
| GO_PROTEIN_POLYMERIZATION | 26 | 0.32148963 | 0.9883967 | 0.49836066 | 0.7559804 |
| GO_TISSUE_MORPHOGENESIS | 73 | 0.27817237 | 0.9881312 | 0.5196771 | 0.7552177 |
| GO_RESPONSE_TO_HYDROGEN_PEROXIDE | 15 | 0.36286554 | 0.9860577 | 0.5106132 | 0.7583975 |
| GO_RESPONSE_TO_XENOBIOTIC_STIMULUS | 25 | 0.32299942 | 0.98493683 | 0.51746726 | 0.7594646 |
| GO_DIGESTIVE_SYSTEM_DEVELOPMENT | 16 | 0.35293826 | 0.9839527 | 0.5094787 | 0.76033807 |
| GO_NUCLEAR_TRANSCRIBED_MRNA_CATABOLIC_PROCESS | 16 | 0.35313 | 0.9836603 | 0.5074456 | 0.7596547 |
| GO_RESPONSE_TO_TOXIC_SUBSTANCE | 65 | 0.28123966 | 0.9827736 | 0.5254583 | 0.76015216 |
| GO_DEVELOPMENT_OF_PRIMARY_SEXUAL_CHARACTERISTICS | 16 | 0.34895766 | 0.9822534 | 0.51184833 | 0.75991446 |
| GO_INTRINSIC_APOPTOTIC_SIGNALING_PATHWAY | 19 | 0.3392752 | 0.98126036 | 0.5293454 | 0.7606295 |
| GO_DETECTION_OF_STIMULUS | 19 | 0.3372857 | 0.98018426 | 0.50851303 | 0.7616797 |
| GO_REGULATION_OF_CELLULAR_RESPONSE_TO_STRESS | 41 | 0.29485643 | 0.97743213 | 0.53541666 | 0.7663012 |

| | | | | | |
|---|---|---|---|---|---|
| GO_REGULATION_OF_CELL_PROJECTION_ORGANIZATION | 59 | 0.2807553 | 0.9758485 | 0.5426516 | 0.76829827 |
| GO_MUSCLE_CELL_DEVELOPMENT | 18 | 0.33808222 | 0.97328717 | 0.4954023 | 0.7723703 |
| GO_POSITIVE_REGULATION_OF_PROTEIN_LOCALIZATION_TO_MEMBRANE | 16 | 0.35017124 | 0.97316897 | 0.51121604 | 0.77131826 |
| GO_GOLGI_VESICLE_TRANSPORT | 17 | 0.3435766 | 0.9699731 | 0.5289444 | 0.7766023 |
| GO_REGULATION_OF_TRANSMEMBRANE_TRANSPORT | 55 | 0.28182465 | 0.96945226 | 0.5534079 | 0.77632594 |
| GO_RENAL_SYSTEM_DEVELOPMENT | 38 | 0.30098113 | 0.9676274 | 0.53974897 | 0.7787662 |
| GO_T_CELL_RECEPTOR_SIGNALING_PATHWAY | 21 | 0.33023527 | 0.9671256 | 0.5302691 | 0.7785311 |
| GO_MONOCARBOXYLIC_ACID_METABOLIC_PROCESS | 57 | 0.28100348 | 0.96586794 | 0.55419225 | 0.7798017 |
| GO_NOTCH_SIGNALING_PATHWAY | 27 | 0.3167735 | 0.9645467 | 0.5484222 | 0.7812242 |
| GO_MALE_GAMETE_GENERATION | 19 | 0.32801083 | 0.9635085 | 0.5337143 | 0.78209096 |
| GO_EMBRYO_DEVELOPMENT | 80 | 0.2700984 | 0.96277595 | 0.56180906 | 0.78232396 |
| GO_MULTICELLULAR_ORGANISMAL_HOMEOSTASIS | 50 | 0.28350893 | 0.95900863 | 0.56804127 | 0.7887194 |
| GO_NUCLEAR_TRANSCRIBED_MRNA_CATABOLIC_PROCESS_NONSENSE_MEDIATED_DECAY | 15 | 0.35284686 | 0.95895696 | 0.54319525 | 0.78746146 |
| GO_POSITIVE_REGULATION_OF_NUCLEOBASE_CONTAINING_COMPOUND_METABOLIC_PROCESS | 92 | 0.2650806 | 0.9584156 | 0.58787876 | 0.78718364 |
| GO_REGULATION_OF_NEURON_PROJECTION_DEVELOPMENT | 39 | 0.29182813 | 0.953588 | 0.5520505 | 0.7956191 |
| GO_ORGANONITROGEN_COMPOUND_CATABOLIC_PROCESS | 76 | 0.26882917 | 0.9530907 | 0.57315844 | 0.79519403 |
| GO_ACID_SECRETION | 15 | 0.35447526 | 0.95222527 | 0.5428907 | 0.7957698 |
| GO_POSITIVE_REGULATION_OF_NF_KAPPAB_TRANSCRIPTION_FACTOR_ACTIVITY | 15 | 0.35058665 | 0.95216894 | 0.5389151 | 0.794499 |
| GO_REGULATION_OF_CELLULAR_COMPONENT_SIZE | 40 | 0.2912356 | 0.95184946 | 0.5735294 | 0.7937918 |
| GO_PROTEIN_CATABOLIC_PROCESS | 43 | 0.28647313 | 0.9507438 | 0.5705394 | 0.79472095 |
| GO_POSITIVE_REGULATION_OF_MAP_KINASE_ACTIVITY | 24 | 0.32024843 | 0.9505072 | 0.5622933 | 0.79377735 |
| GO_REGULATION_OF_ION_TRANSPORT | 82 | 0.26696435 | 0.95044047 | 0.5931174 | 0.79256004 |
| GO_REGULATION_OF_CYTOSKELETON_ORGANIZATION | 51 | 0.28055322 | 0.949713 | 0.56584364 | 0.79263896 |
| GO_REGULATION_OF_ACTIN_FILAMENT_BASED_PROCESS | 52 | 0.28081927 | 0.9478595 | 0.5836777 | 0.7949932 |
| GO_EPITHELIUM_DEVELOPMENT | 127 | 0.25610805 | 0.9471931 | 0.6056056 | 0.7950596 |
| GO_REGULATION_OF_DEPHOSPHORYLATION | 18 | 0.32843688 | 0.94712365 | 0.556582 | 0.7938681 |
| GO_REGULATION_OF_CELL_PROJECTION_ASSEMBLY | 18 | 0.33315828 | 0.9445212 | 0.5692666 | 0.79765034 |
| GO_SEXUAL_REPRODUCTION | 34 | 0.29307246 | 0.94425267 | 0.5720384 | 0.7968193 |
| GO_REGULATION_OF_CALCIUM_ION_TRANSMEMBRANE_TRANSPORT | 17 | 0.33251935 | 0.94316286 | 0.556713 | 0.7976778 |
| GO_REGULATION_OF_PROTEIN_CONTAINING_COMPLEX_ASSEMBLY | 40 | 0.28122953 | 0.9418264 | 0.58158994 | 0.7990252 |
| GO_CELLULAR_RESPONSE_TO_PEPTIDE | 43 | 0.27880117 | 0.9390576 | 0.58798736 | 0.8033876 |
| GO_POSITIVE_REGULATION_OF_CELLULAR_CATABOLIC_PROCESS | 27 | 0.3005719 | 0.93874216 | 0.5770065 | 0.80269736 |
| GO_CYTOSKELETON_ORGANIZATION | 118 | 0.2557237 | 0.9370831 | 0.6292585 | 0.80469525 |
| GO_CELLULAR_RESPONSE_TO_OXYGEN_LEVELS | 20 | 0.32092765 | 0.93584794 | 0.5576497 | 0.8058006 |
| GO_REGULATION_OF_PROTEIN_CATABOLIC_PROCESS | 27 | 0.30330324 | 0.9350462 | 0.5656009 | 0.8060089 |
| GO_REGULATION_OF_CELLULAR_COMPONENT_BIOGENESIS | 88 | 0.2609508 | 0.93412656 | 0.6157735 | 0.806492 |
| GO_CELL_CELL_JUNCTION_ORGANIZATION | 23 | 0.31355807 | 0.9340924 | 0.5821545 | 0.805235 |
| GO_REGULATION_OF_MAP_KINASE_ACTIVITY | 31 | 0.2939105 | 0.93382263 | 0.5689655 | 0.80439514 |
| GO_INTRACELLULAR_RECEPTOR_SIGNALING_PATHWAY | 17 | 0.33059856 | 0.9328972 | 0.5677868 | 0.8048631 |
| GO_REGULATION_OF_ORGANELLE_ORGANIZATION | 77 | 0.26446536 | 0.93094414 | 0.62336355 | 0.807449 |
| GO_EMBRYONIC_MORPHOGENESIS | 51 | 0.2738881 | 0.93058777 | 0.5967078 | 0.80689174 |
| GO_FAT_CELL_DIFFERENTIATION | 23 | 0.30670583 | 0.9305419 | 0.57298476 | 0.80569226 |
| GO_CELL_MORPHOGENESIS_INVOLVED_IN_NEURON_DIFFERENTIATION | 51 | 0.27288744 | 0.9289221 | 0.61625516 | 0.8075272 |
| GO_REGULATION_OF_ANIMAL_ORGAN_MORPHOGENESIS | 28 | 0.29886708 | 0.9263979 | 0.5883621 | 0.8111083 |
| GO_POSITIVE_REGULATION_OF_NERVOUS_SYSTEM_DEVELOPMENT | 50 | 0.2751548 | 0.925093 | 0.61944157 | 0.81234515 |
| GO_CELL_CELL_JUNCTION_ASSEMBLY | 15 | 0.3364934 | 0.9203502 | 0.5985998 | 0.8201521 |
| GO_POSITIVE_REGULATION_OF_RNA_METABOLIC_PROCESS | 83 | 0.25590464 | 0.91791105 | 0.6481855 | 0.82351214 |
| GO_REGULATION_OF_DNA_BINDING_TRANSCRIPTION_FACTOR_ACTIVITY | 39 | 0.2806693 | 0.91740257 | 0.62198955 | 0.8231155 |

| | | | | | |
|---|---|---|---|---|---|
| GO_SMOOTH_MUSCLE_CELL_PROLIFERATION | 27 | 0.2994135 | 0.91549987 | 0.6146288 | 0.82528466 |
| GO_ADAPTIVE_THERMOGENESIS | 17 | 0.3214525 | 0.9154913 | 0.591224 | 0.82397 |
| GO_NEGATIVE_REGULATION_OF_PHOSPHORUS_METABOLIC_PROCESS | 43 | 0.27218774 | 0.9144001 | 0.63569164 | 0.8247227 |
| GO_POSITIVE_REGULATION_OF_CELLULAR_PROTEIN_CATABOLIC_PROCESS | 15 | 0.33568782 | 0.9097757 | 0.60188454 | 0.83209246 |
| GO_REGULATION_OF_TRANSFERASE_ACTIVITY | 63 | 0.25952348 | 0.9093624 | 0.6364562 | 0.83154327 |
| GO_POSITIVE_REGULATION_OF_TRANSCRIPTION_BY_RNA_POLYMERASE_II | 60 | 0.26507998 | 0.9086827 | 0.65123457 | 0.8314474 |
| GO_EMBRYO_DEVELOPMENT_ENDING_IN_BIRTH_OR_EGG_HATCHING | 48 | 0.26675114 | 0.90301335 | 0.6369231 | 0.8404728 |
| GO_ACTIVATION_OF_PROTEIN_KINASE_ACTIVITY | 30 | 0.2864123 | 0.9024107 | 0.6257928 | 0.8402786 |
| GO_REGULATION_OF_KINASE_ACTIVITY | 60 | 0.26149356 | 0.90180504 | 0.6595529 | 0.8400239 |
| GO_MUSCLE_CONTRACTION | 40 | 0.2741222 | 0.90124613 | 0.637122 | 0.839743 |
| GO_RESPONSE_TO_ORGANOPHOSPHORUS | 26 | 0.28973725 | 0.9005276 | 0.6231263 | 0.83969223 |
| GO_CATION_TRANSPORT | 118 | 0.24254145 | 0.8935146 | 0.7251755 | 0.8509697 |
| GO_POSITIVE_REGULATION_OF_CATABOLIC_PROCESS | 30 | 0.28466365 | 0.8924289 | 0.626753 | 0.8515628 |
| GO_CELL_CYCLE | 68 | 0.25308233 | 0.8919757 | 0.680203 | 0.85100484 |
| GO_CELL_PROJECTION_ORGANIZATION | 123 | 0.24407983 | 0.8908731 | 0.71414244 | 0.8515614 |
| GO_REGULATION_OF_NERVOUS_SYSTEM_DEVELOPMENT | 79 | 0.2502257 | 0.8905568 | 0.6791498 | 0.8507685 |
| GO_CALCIUM_ION_TRANSMEMBRANE_TRANSPORT | 32 | 0.28356147 | 0.89044 | 0.6490486 | 0.8496276 |
| GO_IN_UTERO_EMBRYONIC_DEVELOPMENT | 23 | 0.29744962 | 0.88898253 | 0.6391639 | 0.8509615 |
| GO_GLYCOPROTEIN_BIOSYNTHETIC_PROCESS | 20 | 0.30342335 | 0.88728446 | 0.6379691 | 0.8526454 |
| GO_REGULATION_OF_STRESS_ACTIVATED_PROTEIN_KINASE_SIGNALING_CASCADE | 16 | 0.3116383 | 0.88518304 | 0.63786983 | 0.8549712 |
| GO_POSITIVE_REGULATION_OF_PROTEIN_CONTAINING_COMPLEX_ASSEMBLY | 27 | 0.28790936 | 0.8822798 | 0.6369565 | 0.85852265 |
| GO_HEART_DEVELOPMENT | 55 | 0.25585803 | 0.88187957 | 0.6934827 | 0.8578998 |
| GO_REGULATION_OF_GROWTH | 46 | 0.260933 | 0.8808278 | 0.66875 | 0.85833824 |
| GO_POSITIVE_REGULATION_OF_PROTEIN_CATABOLIC_PROCESS | 18 | 0.3062333 | 0.88080895 | 0.63253695 | 0.8570339 |
| GO_MUSCLE_CELL_PROLIFERATION | 29 | 0.27921107 | 0.88030475 | 0.6453362 | 0.8566195 |
| GO_RESPONSE_TO_PURINE_CONTAINING_COMPOUND | 27 | 0.27795932 | 0.88003194 | 0.65334773 | 0.8557561 |
| GO_PROTEIN_HOMOOLIGOMERIZATION | 37 | 0.27205393 | 0.8794942 | 0.65918803 | 0.85532624 |
| GO_ORGANELLE_LOCALIZATION | 40 | 0.26935437 | 0.8793086 | 0.66528064 | 0.8543211 |
| GO_REGULATION_OF_CATABOLIC_PROCESS | 61 | 0.25303984 | 0.8791312 | 0.69325155 | 0.8532934 |
| GO_FOREBRAIN_DEVELOPMENT | 32 | 0.27754247 | 0.8779909 | 0.66631246 | 0.8538454 |
| GO_FC_RECEPTOR_SIGNALING_PATHWAY | 24 | 0.28267854 | 0.8774279 | 0.6484375 | 0.8534776 |
| GO_GLYCEROLIPID_BIOSYNTHETIC_PROCESS | 15 | 0.3158096 | 0.8765242 | 0.637455 | 0.8537132 |
| GO_SULFUR_COMPOUND_METABOLIC_PROCESS | 28 | 0.28260255 | 0.87627876 | 0.6457883 | 0.85276663 |
| GO_REGULATION_OF_BINDING | 25 | 0.28979817 | 0.875977 | 0.6728665 | 0.8519667 |
| GO_GLIAL_CELL_DEVELOPMENT | 17 | 0.31149423 | 0.8734394 | 0.6312719 | 0.85492235 |
| GO_RESPONSE_TO_CAMP | 18 | 0.30266452 | 0.8732962 | 0.6465116 | 0.8538769 |
| GO_REGULATION_OF_CATION_TRANSMEMBRANE_TRANSPORT | 35 | 0.26806194 | 0.8731206 | 0.67395836 | 0.85287654 |
| GO_TEMPERATURE_HOMEOSTASIS | 19 | 0.30310872 | 0.87089163 | 0.6617647 | 0.8552701 |
| GO_REGULATION_OF_TRANS_SYNAPTIC_SIGNALING | 34 | 0.2720766 | 0.8698133 | 0.6736402 | 0.8556828 |
| GO_ORGANIC_HYDROXY_COMPOUND_TRANSPORT | 23 | 0.2871079 | 0.8646831 | 0.66519827 | 0.862747 |
| GO_CELLULAR_PROTEIN_CATABOLIC_PROCESS | 32 | 0.2742732 | 0.8626603 | 0.685441 | 0.8647993 |
| GO_STRIATED_MUSCLE_CONTRACTION | 17 | 0.30450714 | 0.8602785 | 0.6670547 | 0.8673898 |
| GO_POSITIVE_REGULATION_OF_INTRACELLULAR_TRANSPORT | 17 | 0.30869046 | 0.860245 | 0.6674365 | 0.8661241 |
| GO_FATTY_ACID_METABOLIC_PROCESS | 38 | 0.26837137 | 0.8592474 | 0.6962963 | 0.86641467 |
| GO_MACROMOLECULE_CATABOLIC_PROCESS | 78 | 0.23938899 | 0.8588758 | 0.7474645 | 0.8656827 |
| GO_REGULATION_OF_OSSIFICATION | 27 | 0.27635634 | 0.85705334 | 0.67907995 | 0.86728567 |
| GO_CARDIAC_MUSCLE_TISSUE_DEVELOPMENT | 20 | 0.29251987 | 0.85391724 | 0.6815643 | 0.87092984 |
| GO_ACTOMYOSIN_STRUCTURE_ORGANIZATION | 24 | 0.28234798 | 0.8512578 | 0.66372657 | 0.87395394 |

| | | | | | |
|---|---|---|---|---|---|
| GO_NEGATIVE_REGULATION_OF_MAPK_CASCADE | 19 | 0.29421017 | 0.8501389 | 0.686636 | 0.8743814 |
| GO_ESTABLISHMENT_OF_PROTEIN_LOCALIZATION_TO_MEMBRANE | 31 | 0.26695424 | 0.8471163 | 0.70010906 | 0.8778958 |
| GO_POSITIVE_REGULATION_OF_DEVELOPMENTAL_GROWTH | 16 | 0.3007157 | 0.8458054 | 0.6770115 | 0.8785932 |
| GO_POSITIVE_REGULATION_OF_GROWTH | 21 | 0.29101697 | 0.8450514 | 0.68350166 | 0.8783798 |
| GO_PROTEIN_AUTOPHOSPHORYLATION | 18 | 0.2893314 | 0.83965945 | 0.68607306 | 0.88537216 |
| GO_REGULATION_OF_SYSTEM_PROCESS | 53 | 0.2441143 | 0.8386166 | 0.7252066 | 0.8856247 |
| GO_MRNA_METABOLIC_PROCESS | 32 | 0.26288217 | 0.8376181 | 0.7171825 | 0.88584137 |
| GO_RESPONSE_TO_ENDOPLASMIC_RETICULUM_STRESS | 17 | 0.29198846 | 0.8350969 | 0.68213457 | 0.88845646 |
| GO_POSITIVE_REGULATION_OF_CYTOSKELETON_ORGANIZATION | 26 | 0.27205622 | 0.83282727 | 0.7263736 | 0.8905413 |
| GO_RESPONSE_TO_PEPTIDE_HORMONE | 45 | 0.24842647 | 0.8298993 | 0.7530992 | 0.89357144 |
| GO_ORGANIC_ACID_METABOLIC_PROCESS | 95 | 0.23017015 | 0.82806903 | 0.79376256 | 0.8950195 |
| GO_CHEMICAL_HOMEOSTASIS | 121 | 0.22667453 | 0.8277243 | 0.8198198 | 0.8942181 |
| GO_REGULATION_OF_CELLULAR_PROTEIN_CATABOLIC_PROCESS | 18 | 0.285769 | 0.82713974 | 0.687067 | 0.8938416 |
| GO_REGULATION_OF_TRANSPORTER_ACTIVITY | 28 | 0.26677036 | 0.8258897 | 0.70614034 | 0.894385 |
| GO_REGULATION_OF_SUPRAMOLECULAR_FIBER_ORGANIZATION | 42 | 0.24970677 | 0.8257809 | 0.72129434 | 0.8932449 |
| GO_METAL_ION_TRANSPORT | 93 | 0.22911017 | 0.825038 | 0.8068411 | 0.8930406 |
| GO_PROTEIN_TARGETING_TO_MEMBRANE | 22 | 0.2755899 | 0.8217671 | 0.7081967 | 0.8966164 |
| GO_ENDOTHELIUM_DEVELOPMENT | 16 | 0.3000096 | 0.82063746 | 0.6913295 | 0.8969335 |
| GO_LIPID_METABOLIC_PROCESS | 123 | 0.22492646 | 0.8203397 | 0.8214644 | 0.8960387 |
| GO_PROTEIN_LOCALIZATION_TO_ENDOPLASMIC_RETICULUM | 16 | 0.29356998 | 0.8172821 | 0.70422536 | 0.89913845 |
| GO_REGULATION_OF_PROTEIN_SERINE_THREONINE_KINASE_ACTIVITY | 35 | 0.25060496 | 0.81556773 | 0.75 | 0.90017915 |
| GO_REGULATION_OF_NEURON_DIFFERENTIATION | 49 | 0.240162 | 0.8135653 | 0.7507724 | 0.90166444 |
| GO_LIPID_CATABOLIC_PROCESS | 35 | 0.25440988 | 0.8119656 | 0.7340425 | 0.9025944 |
| GO_MUSCLE_SYSTEM_PROCESS | 48 | 0.24124077 | 0.81182986 | 0.77662873 | 0.90148306 |
| GO_POSITIVE_REGULATION_OF_SUPRAMOLECULAR_FIBER_ORGANIZATION | 26 | 0.26290193 | 0.8107222 | 0.75376344 | 0.90170294 |
| GO_ACTIN_MEDIATED_CELL_CONTRACTION | 17 | 0.29029518 | 0.8091495 | 0.7118644 | 0.90259963 |
| GO_REGULATION_OF_CELLULAR_CATABOLIC_PROCESS | 52 | 0.23520881 | 0.80246747 | 0.76954734 | 0.9105153 |
| GO_LIPID_BIOSYNTHETIC_PROCESS | 55 | 0.23363003 | 0.80139387 | 0.7833676 | 0.91066283 |
| GO_NEGATIVE_REGULATION_OF_ORGANELLE_ORGANIZATION | 25 | 0.25802925 | 0.8003084 | 0.74336284 | 0.91081053 |
| GO_HEAD_DEVELOPMENT | 64 | 0.23075762 | 0.79959416 | 0.7823887 | 0.9104674 |
| GO_PROTEIN_DEPHOSPHORYLATION | 22 | 0.26818684 | 0.7963211 | 0.7357456 | 0.9136333 |
| GO_ACTIN_FILAMENT_BASED_MOVEMENT | 21 | 0.26680973 | 0.79463255 | 0.73460245 | 0.9144919 |
| GO_MESONEPHROS_DEVELOPMENT | 16 | 0.2898242 | 0.7939965 | 0.75 | 0.9140848 |
| GO_CHROMOSOME_ORGANIZATION | 15 | 0.29138017 | 0.789931 | 0.72843826 | 0.9181669 |
| GO_ION_TRANSPORT | 164 | 0.21142903 | 0.7884153 | 0.891 | 0.9188541 |
| GO_REGULATION_OF_MORPHOGENESIS_OF_AN_EPITHELIUM | 23 | 0.2608717 | 0.78657895 | 0.7511161 | 0.9199159 |
| GO_OXIDATION_REDUCTION_PROCESS | 88 | 0.22003095 | 0.78490734 | 0.8415742 | 0.9207895 |
| GO_NEGATIVE_REGULATION_OF_SUPRAMOLECULAR_FIBER_ORGANIZATION | 18 | 0.27071857 | 0.7819302 | 0.7580275 | 0.9232515 |
| GO_POSITIVE_REGULATION_OF_PROTEIN_SERINE_THREONINE_KINASE_ACTIVITY | 26 | 0.25901777 | 0.78060865 | 0.77292573 | 0.92361504 |
| GO_EPITHELIAL_CELL_PROLIFERATION | 43 | 0.23371424 | 0.7787962 | 0.79249215 | 0.9245349 |
| GO_REGULATION_OF_SYNAPSE_STRUCTURE_OR_ACTIVITY | 16 | 0.28343078 | 0.77740175 | 0.76583034 | 0.9250284 |
| GO_RHYTHMIC_PROCESS | 20 | 0.2672009 | 0.7761616 | 0.7595376 | 0.92521435 |
| GO_CELLULAR_RESPONSE_TO_DNA_DAMAGE_STIMULUS | 17 | 0.27579954 | 0.7757595 | 0.7523256 | 0.9244051 |
| GO_RECEPTOR_METABOLIC_PROCESS | 19 | 0.26990607 | 0.77454066 | 0.75678736 | 0.9245616 |
| GO_REGULATION_OF_ION_TRANSMEMBRANE_TRANSPORT | 47 | 0.230228 | 0.77427894 | 0.8068536 | 0.92358404 |
| GO_CELLULAR_MACROMOLECULE_LOCALIZATION | 125 | 0.20941699 | 0.7737327 | 0.8668669 | 0.9229646 |
| GO_REGULATION_OF_LIPID_BIOSYNTHETIC_PROCESS | 17 | 0.2735041 | 0.77269757 | 0.7727811 | 0.92289096 |
| GO_POSITIVE_REGULATION_OF_ORGANELLE_ORGANIZATION | 44 | 0.23085657 | 0.7724519 | 0.80877745 | 0.92192936 |

| | | | | | |
|---|---|---|---|---|---|
| GO_NEGATIVE_REGULATION_OF_CELL_PROJECTION_ORGANIZATION | 15 | 0.28421283 | 0.7722112 | 0.76367867 | 0.92092043 |
| GO_ERBB_SIGNALING_PATHWAY | 15 | 0.28512123 | 0.77173275 | 0.760514 | 0.9201977 |
| GO_TELENCEPHALON_DEVELOPMENT | 15 | 0.2821396 | 0.77166146 | 0.7587822 | 0.9190038 |
| GO_HEART_PROCESS | 25 | 0.25198755 | 0.768597 | 0.7697368 | 0.92151225 |
| GO_SYNAPTIC_SIGNALING | 54 | 0.22171594 | 0.7669504 | 0.82032853 | 0.9220984 |
| GO_MODIFICATION_DEPENDENT_MACROMOLECULE_CATABOLIC_PROCESS | 18 | 0.26998943 | 0.7666788 | 0.7599545 | 0.92117274 |
| GO_NEGATIVE_REGULATION_OF_PHOSPHORYLATION | 34 | 0.23454496 | 0.76613635 | 0.798722 | 0.92055607 |
| GO_GLAND_MORPHOGENESIS | 15 | 0.28275594 | 0.7617028 | 0.7912844 | 0.92455614 |
| GO_REGULATION_OF_NEUROTRANSMITTER_LEVELS | 29 | 0.24556315 | 0.7610788 | 0.78177965 | 0.9240445 |
| GO_GLAND_DEVELOPMENT | 36 | 0.23720098 | 0.7602702 | 0.80021596 | 0.9236594 |
| GO_PLASMA_MEMBRANE_ORGANIZATION | 15 | 0.28060967 | 0.75973797 | 0.77922076 | 0.92300296 |
| GO_REGULATION_OF_OSTEOBLAST_DIFFERENTIATION | 16 | 0.2696039 | 0.7587361 | 0.7867133 | 0.9228782 |
| GO_PEPTIDYL_SERINE_MODIFICATION | 17 | 0.26337516 | 0.7544045 | 0.79186046 | 0.9265473 |
| GO_REGULATION_OF_PROTEIN_POLYMERIZATION | 22 | 0.2524448 | 0.75344616 | 0.77034557 | 0.926374 |
| GO_SMALL_MOLECULE_BIOSYNTHETIC_PROCESS | 61 | 0.21496454 | 0.75136757 | 0.8400413 | 0.927507 |
| GO_PROTEIN_LOCALIZATION_TO_MEMBRANE | 61 | 0.21621227 | 0.7504313 | 0.8494405 | 0.9272751 |
| GO_SMALL_MOLECULE_METABOLIC_PROCESS | 158 | 0.19885863 | 0.7449271 | 0.9248497 | 0.93218315 |
| GO_REGULATION_OF_LIPID_METABOLIC_PROCESS | 38 | 0.22779973 | 0.7437466 | 0.8277311 | 0.93218505 |
| GO_DENDRITE_DEVELOPMENT | 21 | 0.25242224 | 0.73780435 | 0.7967033 | 0.9373101 |
| GO_CARBOHYDRATE_DERIVATIVE_BIOSYNTHETIC_PROCESS | 53 | 0.21684258 | 0.7355614 | 0.8543388 | 0.9384734 |
| GO_CENTRAL_NERVOUS_SYSTEM_DEVELOPMENT | 85 | 0.20449662 | 0.7344643 | 0.8780242 | 0.93843246 |
| GO_REGULATION_OF_DEVELOPMENTAL_GROWTH | 25 | 0.24073401 | 0.73224574 | 0.83093923 | 0.9395119 |
| GO_MITOCHONDRION_ORGANIZATION | 38 | 0.22252145 | 0.7318046 | 0.82172996 | 0.93872327 |
| GO_CELLULAR_LIPID_METABOLIC_PROCESS | 96 | 0.20224966 | 0.73170954 | 0.9007021 | 0.9375369 |
| GO_STRESS_ACTIVATED_PROTEIN_KINASE_SIGNALING_CASCADE | 25 | 0.23840708 | 0.729912 | 0.8093682 | 0.9381401 |
| GO_CARDIAC_MUSCLE_CONTRACTION | 15 | 0.2640991 | 0.7203856 | 0.8203593 | 0.9467542 |
| GO_CELLULAR_MACROMOLECULE_CATABOLIC_PROCESS | 57 | 0.2094992 | 0.71710855 | 0.8599382 | 0.94871587 |
| GO_REGULATION_OF_AUTOPHAGY | 18 | 0.24831422 | 0.71664417 | 0.819222 | 0.9478875 |
| GO_SEX_DIFFERENTIATION | 20 | 0.24542329 | 0.71620107 | 0.8076923 | 0.9470861 |
| GO_MORPHOGENESIS_OF_AN_EPITHELIUM | 58 | 0.2036738 | 0.7140231 | 0.89014375 | 0.94786626 |
| GO_VESICLE_ORGANIZATION | 18 | 0.24992093 | 0.71282786 | 0.82578397 | 0.9476833 |
| GO_REGULATION_OF_CELL_CYCLE | 48 | 0.21439764 | 0.71194494 | 0.8679245 | 0.9472863 |
| GO_NEGATIVE_REGULATION_OF_NEURON_DIFFERENTIATION | 15 | 0.26081157 | 0.71080166 | 0.8160652 | 0.9471506 |
| GO_CELL_CYCLE_PHASE_TRANSITION | 25 | 0.23001799 | 0.7029066 | 0.8308026 | 0.9534743 |
| GO_NON_CANONICAL_WNT_SIGNALING_PATHWAY | 16 | 0.25361326 | 0.6989339 | 0.83566433 | 0.9560048 |
| GO_HEART_MORPHOGENESIS | 28 | 0.22381243 | 0.69418496 | 0.85761225 | 0.9590111 |
| GO_POSITIVE_REGULATION_OF_EPITHELIAL_CELL_PROLIFERATION | 19 | 0.23678666 | 0.6935064 | 0.8447277 | 0.9583455 |
| GO_SENSORY_SYSTEM_DEVELOPMENT | 37 | 0.21125047 | 0.6928179 | 0.8819149 | 0.9577148 |
| GO_ACTIN_POLYMERIZATION_OR_DEPOLYMERIZATION | 24 | 0.22645667 | 0.6844684 | 0.8620309 | 0.9638086 |
| GO_DEPHOSPHORYLATION | 33 | 0.21344143 | 0.68388635 | 0.8854719 | 0.963048 |
| GO_NEGATIVE_REGULATION_OF_PROTEIN_MODIFICATION_PROCESS | 45 | 0.20295013 | 0.68356234 | 0.8896982 | 0.96205276 |
| GO_MORPHOGENESIS_OF_EMBRYONIC_EPITHELIUM | 15 | 0.24881843 | 0.6780677 | 0.8611435 | 0.9655871 |
| GO_SYNAPSE_ASSEMBLY | 15 | 0.24397016 | 0.67086565 | 0.8574713 | 0.97040206 |
| GO_ACTIN_FILAMENT_ORGANIZATION | 56 | 0.19103952 | 0.6705976 | 0.90759754 | 0.9693337 |
| GO_ORGANONITROGEN_COMPOUND_BIOSYNTHETIC_PROCESS | 139 | 0.1796222 | 0.6691386 | 0.961 | 0.9692782 |
| GO_POSITIVE_REGULATION_OF_CANONICAL_WNT_SIGNALING_PATHWAY | 16 | 0.2400383 | 0.6678537 | 0.8485549 | 0.96907264 |
| GO_NEGATIVE_REGULATION_OF_RNA_BIOSYNTHETIC_PROCESS | 61 | 0.19111265 | 0.6677895 | 0.9100205 | 0.96786827 |
| GO_PROTEIN_CONTAINING_COMPLEX_ASSEMBLY | 130 | 0.18158922 | 0.6668219 | 0.9649299 | 0.9673746 |

| | | | | | |
|---|---|---|---|---|---|
| GO_NEPHRON_DEVELOPMENT | 23 | 0.21357703 | 0.6568954 | 0.87068963 | 0.974041 |
| GO_CELL_SURFACE_RECEPTOR_SIGNALING_PATHWAY_INVOLVED_IN_CELL_CELL_SIGNALING | 58 | 0.18849656 | 0.65434384 | 0.9117949 | 0.9747842 |
| GO_CATION_TRANSMEMBRANE_TRANSPORT | 83 | 0.1794679 | 0.64774585 | 0.9481707 | 0.97847724 |
| GO_ACTIN_FILAMENT_BUNDLE_ORGANIZATION | 22 | 0.21551968 | 0.6443051 | 0.879558 | 0.9797626 |
| GO_POSITIVE_REGULATION_OF_CELLULAR_COMPONENT_BIOGENESIS | 60 | 0.18543744 | 0.6405435 | 0.9270298 | 0.98121935 |
| GO_CARDIAC_CHAMBER_DEVELOPMENT | 19 | 0.21877499 | 0.63463986 | 0.8868145 | 0.9841668 |
| GO_DETOXIFICATION | 15 | 0.22751147 | 0.6307553 | 0.88836664 | 0.98552704 |
| GO_NEGATIVE_REGULATION_OF_CYTOSKELETON_ORGANIZATION | 15 | 0.22761852 | 0.6304568 | 0.89634866 | 0.9844607 |
| GO_CARDIAC_CHAMBER_MORPHOGENESIS | 19 | 0.21877499 | 0.6295796 | 0.8901345 | 0.98380244 |
| GO_EPITHELIAL_TUBE_MORPHOGENESIS | 37 | 0.19480425 | 0.6276169 | 0.91534394 | 0.98387027 |
| GO_REGULATION_OF_ACTIN_FILAMENT_ORGANIZATION | 33 | 0.19699283 | 0.62672526 | 0.9139785 | 0.9832163 |
| GO_ORGANIC_HYDROXY_COMPOUND_BIOSYNTHETIC_PROCESS | 24 | 0.20588088 | 0.62412626 | 0.89709175 | 0.9836789 |
| GO_NEGATIVE_REGULATION_OF_BIOSYNTHETIC_PROCESS | 80 | 0.17591105 | 0.6232145 | 0.95238096 | 0.98301923 |
| GO_KIDNEY_EPITHELIUM_DEVELOPMENT | 21 | 0.2052132 | 0.61934537 | 0.8969072 | 0.984266 |
| GO_INTRACELLULAR_PROTEIN_TRANSPORT | 60 | 0.18360531 | 0.61894035 | 0.961264 | 0.9832875 |
| GO_NEURAL_TUBE_DEVELOPMENT | 19 | 0.21123078 | 0.6134879 | 0.90909094 | 0.9855001 |
| GO_REGULATION_OF_CELL_CYCLE_PROCESS | 32 | 0.1934779 | 0.6128678 | 0.92068595 | 0.9846481 |
| GO_CELLULAR_NITROGEN_COMPOUND_CATABOLIC_PROCESS | 39 | 0.1851399 | 0.6083801 | 0.9309623 | 0.9861087 |
| GO_BLOOD_VESSEL_ENDOTHELIAL_CELL_MIGRATION | 15 | 0.21877614 | 0.6065187 | 0.9178744 | 0.9860135 |
| GO_NEGATIVE_REGULATION_OF_CANONICAL_WNT_SIGNALING_PATHWAY | 15 | 0.21644942 | 0.60163957 | 0.92614305 | 0.98765385 |
| GO_RENAL_TUBULE_DEVELOPMENT | 15 | 0.21957672 | 0.60010076 | 0.9226328 | 0.9872584 |
| GO_ERYTHROCYTE_HOMEOSTASIS | 15 | 0.21981166 | 0.59939754 | 0.9115566 | 0.9864353 |
| GO_ORGANIC_CYCLIC_COMPOUND_CATABOLIC_PROCESS | 44 | 0.17754997 | 0.5967255 | 0.9387331 | 0.98668075 |
| GO_REGULATION_OF_MEMBRANE_POTENTIAL | 41 | 0.1807335 | 0.59547704 | 0.9386056 | 0.986142 |
| GO_REGULATION_OF_CELL_CYCLE_PHASE_TRANSITION | 21 | 0.20140454 | 0.59385926 | 0.92542374 | 0.98582137 |
| GO_CARBOHYDRATE_DERIVATIVE_METABOLIC_PROCESS | 86 | 0.16684343 | 0.5929266 | 0.95445347 | 0.98508394 |
| GO_ESTABLISHMENT_OR_MAINTENANCE_OF_CELL_POLARITY | 22 | 0.1965586 | 0.5924334 | 0.92367256 | 0.98408777 |
| GO_EMBRYONIC_ORGAN_MORPHOGENESIS | 23 | 0.19977158 | 0.58946884 | 0.92809737 | 0.9845107 |
| GO_CELL_CYCLE_PROCESS | 45 | 0.1796515 | 0.58444136 | 0.9523316 | 0.9860296 |
| GO_INSULIN_SECRETION | 15 | 0.2121748 | 0.5836847 | 0.9314421 | 0.9851672 |
| GO_PROTEIN_LOCALIZATION_TO_ORGANELLE | 60 | 0.16623114 | 0.58226407 | 0.96247464 | 0.98463655 |
| GO_REGULATION_OF_CARBOHYDRATE_METABOLIC_PROCESS | 15 | 0.21026556 | 0.5816329 | 0.94145197 | 0.9837293 |
| GO_JNK_CASCADE | 16 | 0.20660064 | 0.57629126 | 0.94062865 | 0.9851702 |
| GO_NEGATIVE_REGULATION_OF_NUCLEOBASE_CONTAINING_COMPOUND_METABOLIC_PROCESS | 66 | 0.1627906 | 0.5655616 | 0.9695122 | 0.98918605 |
| GO_ACTIN_FILAMENT_POLYMERIZATION | 20 | 0.19114141 | 0.56409615 | 0.924594 | 0.98864627 |
| GO_ELECTRON_TRANSPORT_CHAIN | 18 | 0.19614156 | 0.56333905 | 0.93942857 | 0.98775786 |
| GO_CELLULAR_RESPONSE_TO_PEPTIDE_HORMONE_STIMULUS | 31 | 0.17896397 | 0.5621586 | 0.9544962 | 0.9870976 |
| GO_NEGATIVE_REGULATION_OF_TRANSCRIPTION_BY_RNA_POLYMERASE_II | 42 | 0.16955325 | 0.55266875 | 0.971875 | 0.9900654 |
| GO_INTRACELLULAR_TRANSPORT | 100 | 0.15219602 | 0.55124736 | 0.979859 | 0.9894934 |
| GO_CELLULAR_LIPID_CATABOLIC_PROCESS | 22 | 0.18503375 | 0.5493605 | 0.9590909 | 0.98904735 |
| GO_REGULATION_OF_CELLULAR_AMIDE_METABOLIC_PROCESS | 22 | 0.18540053 | 0.5482199 | 0.94150734 | 0.9882473 |
| GO_SENSORY_ORGAN_DEVELOPMENT | 49 | 0.16066378 | 0.5437963 | 0.96498454 | 0.98883796 |
| GO_MEMBRANE_LIPID_METABOLIC_PROCESS | 18 | 0.18321298 | 0.52475107 | 0.9516129 | 0.99482566 |
| GO_CANONICAL_WNT_SIGNALING_PATHWAY | 31 | 0.15666157 | 0.5026436 | 0.9717698 | 1.0 |
| GO_PROTEIN_TARGETING | 33 | 0.15790506 | 0.49950013 | 0.9815217 | 1.0 |
| GO_NEGATIVE_REGULATION_OF_KINASE_ACTIVITY | 15 | 0.17925933 | 0.49222192 | 0.9775148 | 1.0 |
| GO_LIPID_MODIFICATION | 22 | 0.16793114 | 0.49109447 | 0.979615 | 1.0 |
| GO_CELL_CELL_SIGNALING_BY_WNT | 48 | 0.14495811 | 0.48923424 | 0.98654246 | 0.99963695 |

| | | | | | |
|---|---|---|---|---|---|
| GO_REGULATION_OF_WNT_SIGNALING_PATHWAY | 34 | 0.15236975 | 0.48844692 | 0.97639483 | 0.99861914 |
| GO_POSITIVE_REGULATION_OF_WNT_SIGNALING_PATHWAY | 20 | 0.16523926 | 0.47866815 | 0.979615 | 0.9998992 |
| GO_RESPONSE_TO_CARBOHYDRATE | 22 | 0.15921144 | 0.47553235 | 0.98318386 | 0.99941117 |
| GO_CELLULAR_AMIDE_METABOLIC_PROCESS | 78 | 0.13314764 | 0.47311646 | 0.99697274 | 0.9987534 |
| GO_MONOSACCHARIDE_METABOLIC_PROCESS | 19 | 0.15756696 | 0.4649801 | 0.98504025 | 0.9992338 |
| GO_ORGANELLE_ASSEMBLY | 53 | 0.13481815 | 0.45687908 | 0.9917355 | 0.99960345 |
| GO_NEGATIVE_REGULATION_OF_TRANSFERASE_ACTIVITY | 16 | 0.16373956 | 0.45249578 | 0.985023 | 0.9991578 |
| GO_MICROTUBULE_CYTOSKELETON_ORGANIZATION | 24 | 0.14636274 | 0.44701993 | 0.9855234 | 0.9989119 |
| GO_SPHINGOLIPID_METABOLIC_PROCESS | 15 | 0.16297576 | 0.44541997 | 0.9847775 | 0.9979298 |
| GO_PEPTIDE_METABOLIC_PROCESS | 56 | 0.12904884 | 0.44087526 | 0.99591005 | 0.9974473 |
| GO_CELL_PROJECTION_ASSEMBLY | 46 | 0.12801892 | 0.43296003 | 0.99484 | 0.99733245 |
| GO_REGULATION_OF_PROTEIN_STABILITY | 20 | 0.14480446 | 0.42026994 | 0.9954597 | 0.9977922 |
| GO_RESPONSE_TO_INSULIN | 28 | 0.12787673 | 0.40181682 | 0.99349946 | 0.998373 |
| GO_CELLULAR_RESPONSE_TO_INSULIN_STIMULUS | 23 | 0.10907424 | 0.32166678 | 1.0 | 1.0 |
| GO_CELLULAR_PROTEIN_CONTAINING_COMPLEX_ASSEMBLY | 67 | 0.07729677 | 0.26710054 | 1.0 | 0.999956 |

**Supplementary Table 5: GSEA pathways (biological processes) downregulated for the 1,262 common regulated genes between *Glis2* $^{lacZ/lacZ}$ and *Lkb1* $^{\Delta Tub}$ mouse kidney**
ES: enrichment score; NES: normalized enrichment score; NOM: nominal; FDR: false discovery rate. Gene sets with FDR q-value < 0.05 are considered as significant.

| NAME | SIZE | ES | NES | NOM p-val | FDR q-val |
|---|---|---|---|---|---|
| GO_ENERGY_DERIVATION_BY_OXIDATION_OF_ORGANIC_COMPOUNDS | 26 | -0.63349515 | -3.3528178 | 0.0 | 0.0 |
| GO_CELLULAR_RESPIRATION | 18 | -0.63022506 | -2.6741097 | 0.0 | 0.0 |
| GO_COENZYME_BIOSYNTHETIC_PROCESS | 15 | -0.56466544 | -2.3351736 | 0.0 | 0.0052489946 |
| GO_ANION_TRANSMEMBRANE_TRANSPORT | 37 | -0.4012176 | -2.1459675 | 0.0 | 0.017747791 |
| GO_ORGANIC_ACID_CATABOLIC_PROCESS | 28 | -0.41107604 | -2.0819454 | 0.0 | 0.02221537 |
| GO_MONOVALENT_INORGANIC_CATION_TRANSPORT | 47 | -0.32601765 | -1.9480371 | 0.0 | 0.041142363 |
| GO_MONOVALENT_INORGANIC_CATION_HOMEOSTASIS | 17 | -0.46807086 | -1.9465358 | 0.008474576 | 0.035493456 |
| GO_COENZYME_METABOLIC_PROCESS | 20 | -0.436605 | -1.8925304 | 0.0 | 0.04434427 |
| GO_COFACTOR_BIOSYNTHETIC_PROCESS | 21 | -0.3943273 | -1.8058313 | 0.0 | 0.06627838 |
| GO_VESICLE_MEDIATED_TRANSPORT_IN_SYNAPSE | 18 | -0.41340137 | -1.7974292 | 0.022727273 | 0.06304202 |
| GO_MULTICELLULAR_ORGANISMAL_SIGNALING | 15 | -0.44049096 | -1.7775297 | 0.007751938 | 0.065661095 |
| GO_GENERATION_OF_PRECURSOR_METABOLITES_AND_ENERGY | 45 | -0.28848022 | -1.6926657 | 0.0 | 0.10017364 |
| GO_SMALL_MOLECULE_CATABOLIC_PROCESS | 41 | -0.2731912 | -1.624169 | 0.018181818 | 0.13441616 |
| GO_PROTON_TRANSMEMBRANE_TRANSPORT | 17 | -0.3787842 | -1.6174095 | 0.031496063 | 0.12968306 |
| GO_INORGANIC_ANION_TRANSPORT | 25 | -0.32328567 | -1.5986445 | 0.010869565 | 0.13531911 |
| GO_POTASSIUM_ION_TRANSPORT | 17 | -0.37086862 | -1.5932577 | 0.030303031 | 0.1313101 |
| GO_REGULATION_OF_CELLULAR_KETONE_METABOLIC_PROCESS | 15 | -0.39465693 | -1.578296 | 0.057142857 | 0.13476701 |
| GO_INORGANIC_ION_TRANSMEMBRANE_TRANSPORT | 86 | -0.21596226 | -1.5668279 | 0.0 | 0.13523461 |
| GO_ORGANIC_ANION_TRANSPORT | 39 | -0.2802466 | -1.5651087 | 0.037037037 | 0.12961411 |
| GO_CHLORIDE_TRANSPORT | 19 | -0.34336168 | -1.5472827 | 0.045045044 | 0.13587607 |
| GO_INORGANIC_ANION_TRANSMEMBRANE_TRANSPORT | 19 | -0.33933124 | -1.4992915 | 0.060344826 | 0.16887866 |
| GO_ORGANIC_ACID_TRANSPORT | 30 | -0.2869893 | -1.4821357 | 0.04761905 | 0.17526779 |
| GO_CELLULAR_KETONE_METABOLIC_PROCESS | 21 | -0.32627645 | -1.4802635 | 0.056603774 | 0.16943383 |
| GO_NEUROTRANSMITTER_TRANSPORT | 20 | -0.32786936 | -1.4369673 | 0.069565214 | 0.20491686 |
| GO_SODIUM_ION_TRANSPORT | 23 | -0.2962114 | -1.3934697 | 0.06521739 | 0.2464507 |
| GO_IMPORT_ACROSS_PLASMA_MEMBRANE | 16 | -0.3300577 | -1.3769759 | 0.055555556 | 0.25981882 |
| GO_CAMERA_TYPE_EYE_MORPHOGENESIS | 15 | -0.33188826 | -1.3621584 | 0.1056338 | 0.269162 |
| GO_COTRANSLATIONAL_PROTEIN_TARGETING_TO_MEMBRANE | 15 | -0.329591 | -1.3597755 | 0.124223605 | 0.2632726 |
| GO_RETINA_DEVELOPMENT_IN_CAMERA_TYPE_EYE | 16 | -0.31873092 | -1.3223126 | 0.0942029 | 0.3075261 |
| GO_ANION_TRANSPORT | 65 | -0.2203763 | -1.318343 | 0.0 | 0.3036991 |
| GO_CILIUM_ORGANIZATION | 20 | -0.30084622 | -1.312728 | 0.15517241 | 0.3014863 |
| GO_TRANSMEMBRANE_TRANSPORT | 143 | -0.18609025 | -1.3020679 | 0.0 | 0.3069768 |
| GO_ION_TRANSMEMBRANE_TRANSPORT | 115 | -0.21280995 | -1.3002279 | 0.0 | 0.30013055 |
| GO_CARBOHYDRATE_BIOSYNTHETIC_PROCESS | 19 | -0.28169286 | -1.2951041 | 0.14634146 | 0.29806408 |
| GO_ORGANOPHOSPHATE_BIOSYNTHETIC_PROCESS | 49 | -0.21885423 | -1.2856225 | 0.074074075 | 0.30250603 |
| GO_GLYCEROPHOSPHOLIPID_METABOLIC_PROCESS | 23 | -0.2501836 | -1.2198063 | 0.20224719 | 0.4041879 |
| GO_EAR_DEVELOPMENT | 23 | -0.24404702 | -1.187994 | 0.18987341 | 0.45673707 |
| GO_CELLULAR_AMINO_ACID_METABOLIC_PROCESS | 24 | -0.23598032 | -1.1838996 | 0.18292683 | 0.45306996 |
| GO_ATP_METABOLIC_PROCESS | 22 | -0.2496397 | -1.1697907 | 0.21782178 | 0.47146505 |
| GO_CARBOHYDRATE_HOMEOSTASIS | 18 | -0.25568512 | -1.1232127 | 0.25396827 | 0.5628007 |
| GO_PHOSPHOLIPID_BIOSYNTHETIC_PROCESS | 17 | -0.26911643 | -1.1230816 | 0.259542 | 0.5493531 |
| GO_BEHAVIOR | 34 | -0.20885406 | -1.1214039 | 0.26666668 | 0.54012656 |
| GO_NEGATIVE_REGULATION_OF_CELL_CYCLE | 17 | -0.26907632 | -1.1091206 | 0.30935252 | 0.55453587 |
| GO_PROTEIN_MODIFICATION_BY_SMALL_PROTEIN_CONJUGATION | 27 | -0.22756737 | -1.1026298 | 0.34246576 | 0.5556745 |
| GO_SENSORY_ORGAN_MORPHOGENESIS | 27 | -0.21730429 | -1.1024073 | 0.30769232 | 0.5439224 |
| GO_NUCLEOSIDE_PHOSPHATE_BIOSYNTHETIC_PROCESS | 21 | -0.24093473 | -1.0804358 | 0.3364486 | 0.58382064 |
| GO_STEROID_METABOLIC_PROCESS | 22 | -0.23497671 | -1.0666208 | 0.3298969 | 0.6044748 |
| GO_CELL_DIVISION | 21 | -0.23552513 | -1.0405071 | 0.3962264 | 0.6518778 |
| GO_RIBOSE_PHOSPHATE_BIOSYNTHETIC_PROCESS | 16 | -0.2399679 | -1.0132737 | 0.43037975 | 0.7045951 |
| GO_CARBOHYDRATE_METABOLIC_PROCESS | 49 | -0.16545348 | -1.0078117 | 0.5 | 0.7035611 |
| GO_PURINE_CONTAINING_COMPOUND_BIOSYNTHETIC_PROCESS | 18 | -0.24035369 | -1.004932 | 0.36764705 | 0.69720244 |
| GO_CELLULAR_CARBOHYDRATE_METABOLIC_PROCESS | 22 | -0.20794007 | -0.98588336 | 0.52380955 | 0.73083436 |
| GO_MONOCARBOXYLIC_ACID_TRANSPORT | 17 | -0.23482212 | -0.98524934 | 0.46052632 | 0.718351 |
| GO_RIBOSE_PHOSPHATE_METABOLIC_PROCESS | 28 | -0.19911776 | -0.97528356 | 0.5050505 | 0.7294064 |
| GO_ORGANOPHOSPHATE_METABOLIC_PROCESS | 83 | -0.15452436 | -0.96614444 | 0.41666666 | 0.7376826 |
| GO_GLYCOPROTEIN_METABOLIC_PROCESS | 26 | -0.19736508 | -0.9609649 | 0.5362319 | 0.7363802 |
| GO_PROTEIN_MODIFICATION_BY_SMALL_PROTEIN_CONJUGATION_OR_REMOVAL | 30 | -0.19049175 | -0.9551728 | 0.48 | 0.7366565 |
| GO_ESTABLISHMENT_OF_PROTEIN_LOCALIZATION_TO_ORGANELLE | 37 | -0.16816327 | -0.9346235 | 0.5 | 0.76856536 |
| GO_EPITHELIAL_CELL_DEVELOPMENT | 25 | -0.18558615 | -0.9284559 | 0.5841584 | 0.7695343 |
| GO_PHOSPHOLIPID_METABOLIC_PROCESS | 38 | -0.17393175 | -0.92123985 | 0.5714286 | 0.77209336 |
| GO_LOCOMOTORY_BEHAVIOR | 17 | -0.22165604 | -0.9199372 | 0.54471546 | 0.76180106 |
| GO_EYE_MORPHOGENESIS | 19 | -0.20790392 | -0.9025995 | 0.5785124 | 0.78590405 |
| GO_AMMONIUM_ION_METABOLIC_PROCESS | 16 | -0.21090157 | -0.88754314 | 0.5971223 | 0.803842 |
| GO_PEPTIDE_BIOSYNTHETIC_PROCESS | 39 | -0.15122662 | -0.8596384 | 0.6923077 | 0.846813 |
| GO_NEGATIVE_REGULATION_OF_WNT_SIGNALING_PATHWAY | 21 | -0.19258662 | -0.8558489 | 0.6764706 | 0.8407868 |
| GO_NUCLEOBASE_CONTAINING_SMALL_MOLECULE_METABOLIC_PROCESS | 37 | -0.15733533 | -0.83805424 | 0.76363635 | 0.8621247 |
| GO_GLYCEROLIPID_METABOLIC_PROCESS | 32 | -0.16210335 | -0.8304251 | 0.69736844 | 0.8638472 |
| GO_RNA_PROCESSING | 18 | -0.19112638 | -0.828797 | 0.7518248 | 0.8543894 |
| GO_ALCOHOL_BIOSYNTHETIC_PROCESS | 15 | -0.20840186 | -0.82482743 | 0.74050635 | 0.84884405 |
| GO_MORPHOGENESIS_OF_A_POLARIZED_EPITHELIUM | 15 | -0.20274901 | -0.8240985 | 0.7692308 | 0.83800745 |
| GO_REGULATION_OF_SMALL_MOLECULE_METABOLIC_PROCESS | 34 | -0.15472312 | -0.82209563 | 0.6923077 | 0.83020353 |
| GO_NEPHRON_EPITHELIUM_DEVELOPMENT | 17 | -0.1871486 | -0.8046845 | 0.76153845 | 0.8482806 |
| GO_KIDNEY_MORPHOGENESIS | 16 | -0.1869984 | -0.77267915 | 0.8064516 | 0.8854121 |
| GO_COFACTOR_METABOLIC_PROCESS | 35 | -0.1350689 | -0.7037476 | 0.8666667 | 0.96022046 |
| GO_ENDOMEMBRANE_SYSTEM_ORGANIZATION | 32 | -0.12895359 | -0.70293105 | 0.9027778 | 0.94833934 |
| GO_PURINE_CONTAINING_COMPOUND_METABOLIC_PROCESS | 32 | -0.13085379 | -0.6775096 | 0.9402985 | 0.9627876 |
| GO_POSTTRANSCRIPTIONAL_REGULATION_OF_GENE_EXPRESSION | 24 | -0.13812602 | -0.65796494 | 0.9583333 | 0.96767616 |
| GO_AMIDE_BIOSYNTHETIC_PROCESS | 57 | -0.09792531 | -0.63223547 | 0.93333334 | 0.9736754 |
| GO_CELL_FATE_COMMITMENT | 15 | -0.13508505 | -0.5326666 | 1.0 | 1.0 |
| GO_MICROTUBULE_BASED_PROCESS | 32 | -0.09384175 | -0.5007692 | 1.0 | 0.99453914 |

**Supplementary Table 6: Common secreted cytokine-coding genes linked to immune response/inflammation among the 823 common upregulated genes between *Glis2* $^{lacZ/lacZ}$ and *Lkb1* $^{\Delta Tub}$ mouse kidney datasets.**

| Gene_ID | Gene_Name | Dataset *Glis2* $^{lacZ/lacZ}$ Fold change | Dataset *Lkb1* $^{\Delta Tub}$ Fold change |
|---|---|---|---|
| **Ccl2** | chemokine (C-C motif) ligand 2 | 2.402 | 1.293 |
| **Ccl5** | chemokine (C-C motif) ligand 5 | 8.537 | 1.622 |
| **Ccl6** | chemokine (C-C motif) ligand 6 | 4.266 | 1.447 |
| **Ccl9** | chemokine (C-C motif) ligand 9 | 13.654 | 2.120 |
| **Ccl12** | chemokine (C-C motif) ligand 12 | 7.932 | 1.376 |
| **Ccl19** | chemokine (C-C motif) ligand 19 | 3.422 | 1.160 |
| **Cx3cl1** | chemokine (C-X3-C motif) ligand 1 | 3.552 | 1.295 |
| **Cxcl1** | chemokine (C-X-C motif) ligand 1 | 10.290 | 1.405 |
| **Cxcl9** | chemokine (C-X-C motif) ligand 9 | 3.872 | 1.610 |
| **Cxcl10** | chemokine (C-X-C motif) ligand 10 | 5.078 | 1.482 |
| **Cxcl12** | chemokine (C-X-C motif) ligand 12 | 1.694 | 1.460 |
| **Cxcl14** | chemokine (C-X-C motif) ligand 14 | 3.973 | 1.314 |
| **Cxcl16** | chemokine (C-X-C motif) ligand 16 | 4.729 | 1.507 |
| **Cxcl17** | chemokine (C-X-C motif) ligand 17 | 1.394 | 1.261 |
| **Il1rn** | interleukin 1 receptor antagonist | 1.932 | 1.346 |
| **Il33** | interleukin 33 | 2.016 | 1.836 |
| **Il34** | interleukin 34 | 1.845 | 1.636 |
| **Lgals9** | lectin, galactose binding, soluble 9 | 2.682 | 1.463 |

**Supplementary Table 7: Expression matrix of the 17 identified cytokines from *Lkb1*$^{\Delta Tub}$ and *Pkd2*$^{\Delta Tub}$ mouse kidney datasets.**

**Dataset *Lkb1*$^{\Delta Tub}$**, GSE86011[16]

| | Control mice | | | | | *Lkb1*$^{\Delta Tub}$ mice | | | | |
|---|---|---|---|---|---|---|---|---|---|---|
| **Gene_ID** | **1** | **2** | **3** | **4** | **5** | **1** | **2** | **3** | **4** | **5** |
| **Ccl5** | 7.125414338 | 6.810792727 | 6.810518472 | 6.914446434 | 6.830401241 | 7.821037108 | 7.456506398 | 8.000980245 | 7.368121218 | 7.33204312 |
| **Ccl6** | 5.691444269 | 5.732864051 | 5.74740808 | 5.570908934 | 5.776654447 | 6.336702504 | 6.418158985 | 6.356774613 | 6.131350874 | 5.940501273 |
| **Ccl9** | 5.790140908 | 5.730285329 | 5.791112887 | 5.656808225 | 5.793988094 | 6.973383335 | 6.913629929 | 7.311428351 | 6.774880909 | 6.208024126 |
| **Ccl12** | 3.896517587 | 3.721551801 | 3.704315479 | 3.801552439 | 3.705581423 | 4.421131143 | 4.094147491 | 4.288536339 | 4.132169986 | 4.193527138 |
| **Ccl19** | 7.048546789 | 6.993994075 | 6.892443016 | 6.764709943 | 6.853125747 | 7.152032751 | 7.055386148 | 7.121918077 | 7.145622605 | 7.14979819 |
| **Cx3cl1** | 8.470048929 | 8.397033197 | 8.370659762 | 8.350374807 | 8.266709271 | 8.817573896 | 8.931806828 | 8.708812819 | 8.633539062 | 8.625419416 |
| **Cxcl1** | 5.828130812 | 5.366751375 | 6.003815006 | 5.85430624 | 5.768670173 | 6.313616873 | 6.042332029 | 6.632991539 | 5.994373084 | 6.290383606 |
| **Cxcl9** | 5.71559903 | 5.940239081 | 5.432509059 | 5.659786812 | 5.239894685 | 6.449370831 | 6.465804898 | 6.768322569 | 5.948751386 | 5.788997367 |
| **Cxcl10** | 5.042581975 | 5.101911189 | 5.145766005 | 5.114525717 | 5.121302668 | 5.607661155 | 5.629850223 | 5.835290906 | 5.556016595 | 5.737290537 |
| **Cxcl12** | 7.990506583 | 8.104861096 | 8.148232079 | 8.177523426 | 7.809948923 | 8.628041931 | 8.812213977 | 8.638768108 | 8.441688923 | 8.441707237 |
| **Cxcl14** | 7.738075798 | 7.939140368 | 8.016163721 | 7.992385656 | 7.971007784 | 8.233873424 | 8.53652988 | 8.525357077 | 8.069691235 | 8.263517422 |
| **Cxcl16** | 7.922921037 | 7.567276725 | 7.720400061 | 7.807455943 | 7.834017309 | 8.566734025 | 8.473978304 | 8.447159166 | 8.127791886 | 8.195358623 |
| **Cxcl17** | 6.240168685 | 6.267132194 | 6.396902684 | 6.200994542 | 6.395175818 | 6.69392813 | 6.633382408 | 6.791748953 | 6.566333373 | 6.487556866 |
| **Il1rn** | 4.800941634 | 4.614863543 | 4.666594473 | 4.596809472 | 4.774817412 | 5.171382344 | 5.053071 | 5.237811814 | 4.947963826 | 5.18771665 |
| **Il33** | 7.461720622 | 7.746553608 | 7.418368367 | 7.440092828 | 7.334218016 | 8.598889893 | 8.585700362 | 8.514461363 | 8.262042392 | 7.823176554 |
| **Il34** | 6.819519761 | 6.618807175 | 6.581003954 | 6.491501323 | 6.454088418 | 7.492094361 | 7.484028492 | 7.480226959 | 7.044630134 | 7.013739973 |
| **Lgals9** | 7.830970596 | 7.767922326 | 7.715137574 | 7.763849441 | 7.753696867 | 8.398397149 | 8.623995549 | 8.427007495 | 8.271282056 | 7.857647871 |

**Dataset *Pkd2*$^{\Delta Tub}$**, GSE149739[25]

| | Control mice | | | *Pkd2*$^{\Delta Tub}$ mice | | |
|---|---|---|---|---|---|---|
| **Gene_ID** | **1** | **2** | **3** | **1** | **2** | **3** |
| **Ccl5** | 15.23947979 | 19.08251621 | 28.17089081 | 88.38501491 | 71.01889552 | 33.14251035 |
| **Ccl6** | 92.52541299 | 127.5515557 | 86.7663437 | 180.74239 | 190.6296669 | 123.7970239 |
| **Ccl9** | 56.60378206 | 47.20411905 | 42.81975403 | 185.7078403 | 111.200639 | 54.5876641 |
| **Ccl12** | 11.97387697 | 5.021714792 | 2.253671265 | 54.61995303 | 24.29593794 | 13.64691603 |
| **Ccl19** | 3.265602811 | 7.030400709 | 3.380506897 | 4.965450276 | 8.410132364 | 0.974779716 |
| **Cx3cl1** | 3498.549145 | 3134.554373 | 3428.960829 | 3188.812167 | 3642.521773 | 2494.461294 |
| **Cxcl1** | 75.10886466 | 139.6036712 | 215.2256058 | 164.8529492 | 168.2026473 | 110.1501079 |
| **Cxcl9** | 19.59361687 | 50.21714792 | 54.08811036 | 107.253726 | 100.9215884 | 50.68854524 |
| **Cxcl10** | 59.86938487 | 70.30400709 | 118.3177414 | 243.3070635 | 192.4985852 | 175.4603489 |
| **Cxcl12** | 7036.285524 | 6540.281345 | 6493.95375 | 4216.660374 | 4297.577638 | 4615.581956 |
| **Cxcl14** | 936.1394726 | 539.3321687 | 414.6755127 | 307.8579171 | 445.7370153 | 452.2977883 |
| **Cxcl16** | 1454.281785 | 1271.498185 | 1424.320239 | 1690.239274 | 1654.927157 | 1595.714395 |
| **Cxcl17** | 5.442671352 | 1.004342958 | 4.50734253 | 0.993090055 | 1.868918303 | 0 |
| **Il1rn** | 5.442671352 | 4.017371834 | 6.761013794 | 58.59231325 | 26.16485624 | 11.69735659 |
| **Il33** | 211.1756485 | 370.6025517 | 273.8210587 | 291.9684762 | 467.2295758 | 251.4931668 |
| **Il34** | 1183.236752 | 954.1258105 | 997.2495347 | 1592.916448 | 1330.669832 | 966.9814784 |
| **Lgals9** | 410.37742 | 307.3289453 | 343.6848679 | 352.5469696 | 402.7518943 | 282.6861177 |
